# Identifying genomic data use with the Data Citation Explorer

**DOI:** 10.1101/2024.01.26.577091

**Authors:** Neil Byers, Charles Parker, Chris Beecroft, T.B.K. Reddy, Hugh Salamon, George Garrity, Kjiersten Fagnan

## Abstract

Increases in sequencing capacity, combined with rapid accumulation of publications and associated data resources, have increased the complexity of maintaining associations between literature and genomic data. As the volume of literature and data have exceeded the capacity of manual curation, automated approaches to maintaining and confirming associations among these resources have become necessary. Here we present the Data Citation Explorer (DCE), which discovers literature incorporating genomic data whether or not provenance was clearly indicated. This service provides advantages over manual curation methods including consistent resource coverage, metadata enrichment, documentation of new use cases, and identification of conflicting metadata. The service reduces labor costs associated with manual review, improves the quality of genome metadata maintained by the U.S. Department of Energy Joint Genome Institute (JGI), and increases the number of known publications that incorporate its data products. The DCE facilitates an understanding of JGI impact, improves credit attribution for data generators, and can encourage data sharing by allowing scientists to see how reuse amplifies the impact of their original studies.

## Introduction

The Department of Energy’s (DOE) Joint Genome Institute (JGI, jgi.doe.gov) is a national User Facility that provides state-of-the-art environmental genomics capabilities to the scientific community. JGI has directly supported more than 4,000 researchers and generated over 15 petabytes of data in its 25-year history. Following delivery to the primary investigators (PIs) and a short embargo period, data produced through JGI proposals is made available for public use through a variety of external and JGI-maintained systems. These include Integrated Microbial Genomes & Microbiomes (IMG/M)^1^, Phytozome^2^, MycoCosm^3^, PhycoCosm^4^, and the JGI Data Portal (data.jgi.doe.gov), as well as the National Center for Biotechnology Information’s (NCBI) Sequence Read Archive (SRA)^5^ and GenBank^6^ databases. This data is made public with the understanding that it may have impact beyond the PIs’ original intended uses. A wider appreciation of this concept has led in recent years to greater community emphasis on the importance not just of initial publication and the findings of original data generators, but also of the downstream reuse of public scientific data. Many efforts are underway to make public scientific data more Findable, Accessible, Interoperable, and Reusable (FAIR)^7^, in order to encourage and facilitate downstream impact.

### Motivation: Understanding institutional and individual impact

Organizations providing services or products to a specific community can benefit from a better understanding of its impact within that community. This understanding is essential for directing service improvements that can benefit the organization and the community it serves. JGI thus has strong motivations for capturing citations of its products. Doing so enables the organization to better serve its users by identifying which data and thematic areas are heavily cited as well as those that are underutilized. This can inform researchers as to which topics may be ripe for innovative analysis and uncover ways in which older data can be reused. Citation capture can also direct improvements in operational efficiency and inform policy decisions. Knowing how specific workflows and product offerings contribute to downstream publications facilitates resource allocation for activities that have the greatest scientific impact.

A more complete understanding of data use is essential for appreciating the extent to which the activities of JGI users align with its policies, initiatives, and strategic goals. JGI also has an interest in identifying researchers, regions, institutions, or specific research fields that make heavy use of its products but with which it has little direct engagement or towards which it has not yet made concerted outreach efforts. Extending searches to include data citations in patents and other intellectual property provides a window into potential commercialization of JGI products and can demonstrate contributions to economic growth and innovation. Building a comprehensive picture of product utilization across science and industry demonstrates to key stakeholders (i.e., taxpayers, elected officials, and the scientific community) that funds granted represent a worthwhile investment with compounding returns over time.

Many organizations, including nonprofit entities like Research Data Alliance^8^ and government entities like NSF, NIH, and DOE^9^, have long advocated for more transparent links among data producers and data consumers. By building connections between citations of JGI data and the datasets themselves, JGI can attribute credit to the primary investigators and JGI personnel who contributed to the creation of a given dataset. Doing so preserves the provenance of an individual’s work and demonstrates their impact on downstream research. Systematically preserving contributor relationships and roles can inform funders, reviewers, and institutional evaluators of significant contributions that are currently missed when using traditional metrics such as the h-index^10^ as indicators of research productivity.

New metrics can account for deficiencies in existing methods for research evaluation that rely heavily on publication authorship, the standards for which are uneven at best^11,12^. For example, an individual’s overall citation count or h-index may not be high, or their inclusion in authorship lists may not be extensive, but their contributions to high-value, frequently utilized data may be significant (i.e., workflow managers and personnel responsible for sample processing). A more granular methodology of research evaluation can also benefit researchers with publications that may not see heavy citation activity, but whose overall contributions as data producers have significant impact across the community. Adopting a contributor evaluation model based on data citations (for example, the “data-index” proposed by Hood et al.^13^) thus offers improvements in equity and transparency over more traditional metrics grounded in the authorship of scientific publications.

### The Problem: Too much literature, too much data

Through anecdotal evidence, the U.S. Department of Energy Joint Genome Institute (JGI) recognizes that a significant body of literature exists that incorporates JGI data products but that does not directly cite either JGI or the individual contributors who produced that data. These publications are difficult for JGI to identify for two reasons. First, as these data are hosted by JGI as well as the US National Institutes for Health (NIH) National Center for Biological Information (NCBI), researchers frequently cite NCBI^56^ and other external identifiers (e.g., Human Oral Microbiome Database, Human Microbiome Project) for a given dataset rather than JGI identifiers. The relationship between these external metadata and JGI data to which they refer are often complex and difficult to traverse. Second, the context within individual publications in which citations of JGI-linked identifiers occur can often be ambiguous or left unindexed by full-text search tools. For example, a JGI-linked identifier could be mistaken for some other identifier by naïve text matching, or could be buried within the supplemental materials of a publication beyond the reach of most searches.

With the rates at which new literature is published and the scale of data production by JGI, it is not feasible for humans to identify all citations of JGI data without automated assistance. Publications associated with the field of “Genetics”’ published from 2011 to 2023 were found to number between 206,974 and 693,491 using research area or MeSH term queries in Web of Science (Clarivate), PubMed (National Library of Medicine)^14^, Dimensions (Digital Science)^15,16^, and SciVal (Elsevier). Though growth in the yearly volume of Genetics publications during this time has slowed in recent years, all four sources show steady increases throughout much of this period (Figure 1). During the same period, the yearly JGI output of raw sequence data increased from just over 30,000 gigabases per year to over 700,000 gigabases per year (Figure 2).

**Figure 1.**
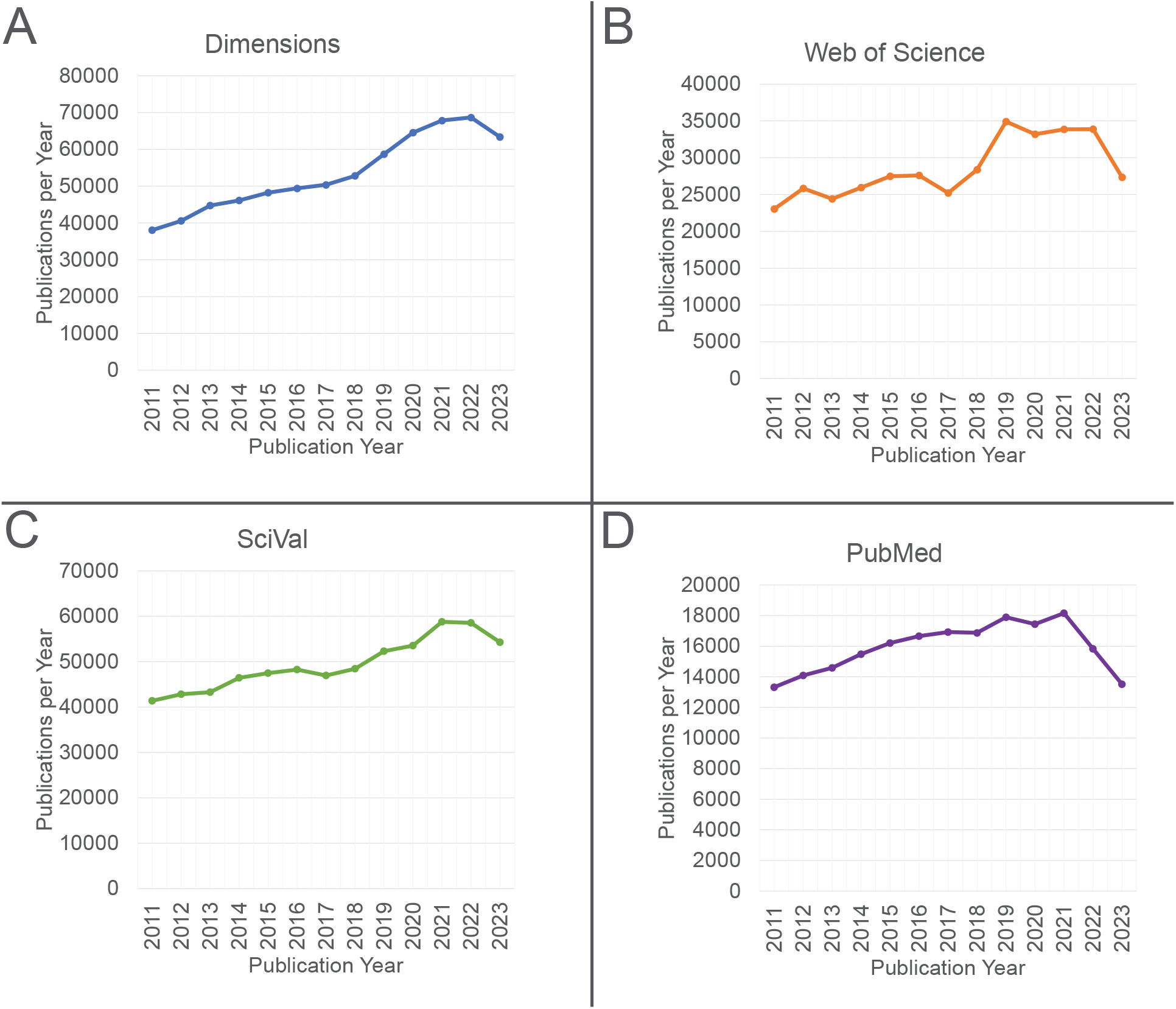
Yearly genetics-associated publications from 2011-2023 in Dimensions (A), Web of Science (B), SciVal (C), and PubMed (D). The net increases in yearly publications were as follows: Dimensions, 66.4% (693,491 total publications). Web of Science, 18.65% (371,079 total publications). SciVal, 31.24% (642,528 total publications), PubMed, 1.48% (206,974 total publications). The research area or MeSH terms used to generate each publication set were “Genetics” (PubMed, SciVal, Dimensions) and “Genetics & Heredity” (Web of Science).

**Figure 2.**
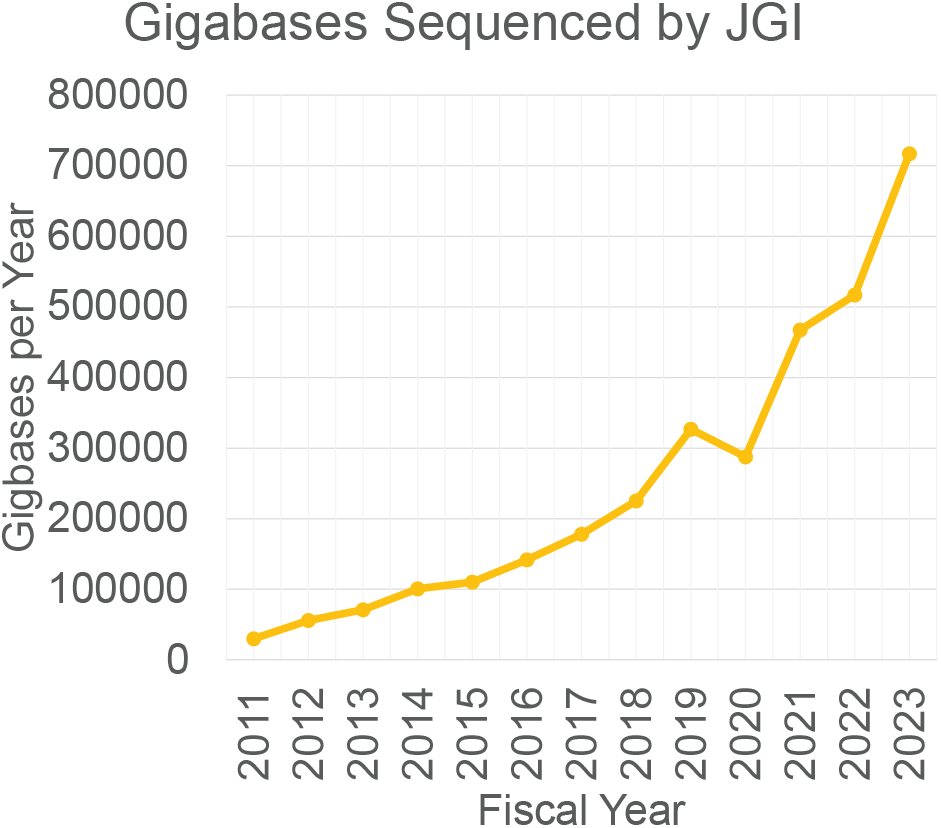
Yearly sequencing production at the DOE Joint Genome Institute (JGI). Note that the fiscal year begins on October 1st of the preceding calendar year. The decrease in FY20 sequencing output was due to the COVID-19 pandemic.

Positive identification of literature citing JGI products is a labor-intensive process when performed manually. This manual process relies on an individual’s expert knowledge within a specific field, which may have specialized literature, subject language terminology, and data resources. This process often involves repetitive searches across numerous sources using distinct queries intended to capture a limited set of JGI products. The timeliness, scope, comprehensiveness, and repeatability of the manual search process is limited by the availability of skilled data curators. Consistency and accuracy of manually compiled citation data varies not only among those performing the searches but also individually over time. The accuracy of even well-trained curators may decline over the course of a single work day. Thus, at present, verification and validation of highly accurate citation data requires a duplication of effort among multiple reviewers in order to identify and correct errors. Even with redundant human reviewers, this form of precise manual tracking of information sources is impractical if not impossible when branching paths are discovered through multiple online resources. Nonetheless, manual literature searches are important for collecting initial data and verifying citations, but there are many repetitive components of this process that are amenable to automation. The DCE automates those repetitive tasks, freeing skilled personnel to focus their attention on tasks that resist automation.

### An automated solution: The Data Citation Explorer

The overarching goal of the DCE is to systematically and reproducibly identify literature that relies on data stored in JGI repositories. The system was designed to encapsulate the expertise of data curators into the business logic of a web service. Using a selected subset of metadata fields from JGI genome projects as input, this web service can consistently apply curatorial methods to incrementally discover and traverse additional resources and metadata that are directly associated with a specific genome. Starting with a limited set of validated genome metadata stored in JGI systems, the service automatically performs an exhaustive search of targeted literature and online resources to discover any uses of that associated data. The service provides users with an audit trail that provides a precise explanation of how each additional resource was discovered. In testing, the DCE has been shown to uncover multiple paths to new information about a genome. The service can also re-process genomes at any time to discover new uses and citations, whether or not the data was consistently referenced.

## Results

Using results returned via searches in PubMed and Pubmed Central, 998 audit trails linking 576 unique publications to the 300 sampled JAMO records were manually evaluated by checking the nature of the hits in the citing publication as well as by verifying the feasibility of relationship between keys in the larger audit trail. Only 10 of the 998 audit trails accounting for 10 out of 576 publications led to invalid connections. Examples of true and false positive hits can be seen in Figures 3 and 4. From the perspective of individual publications, the precision value for connections between publications and JAMO records in this sample was 0.983.

**Figure 3.**
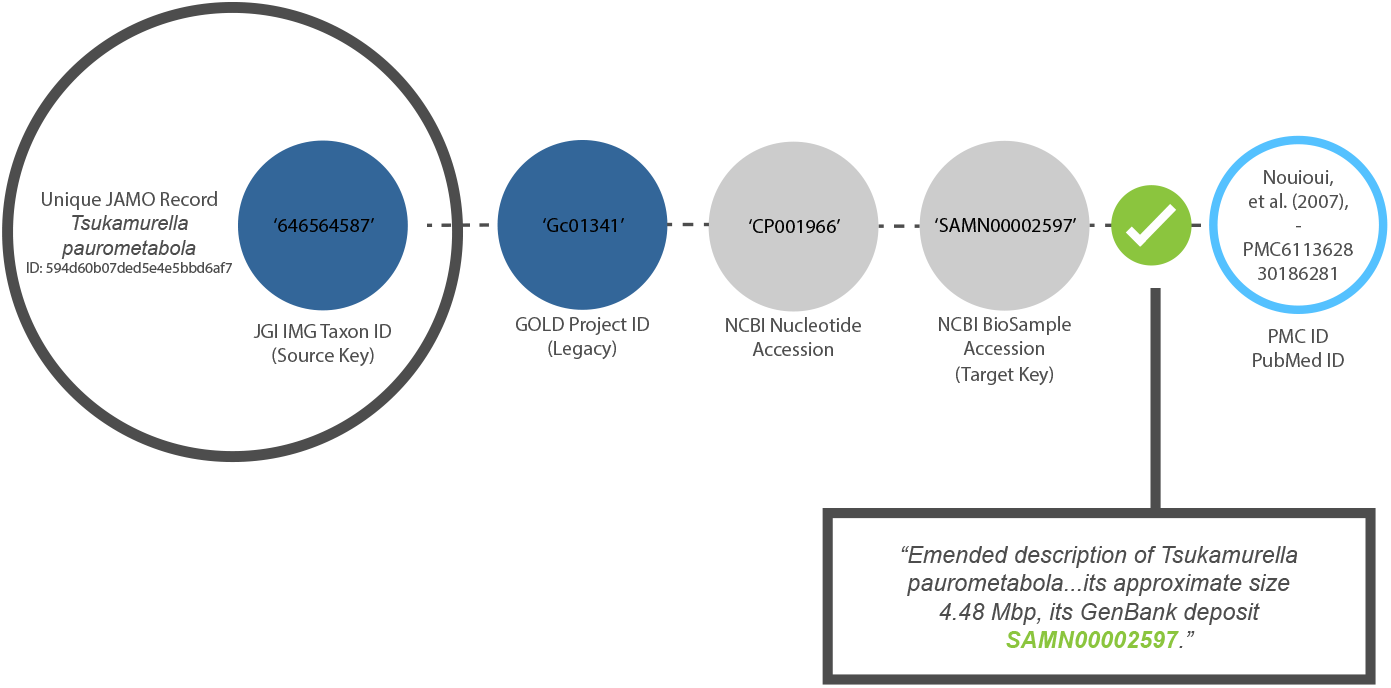
Example of a valid audit trail. The target key refers unambiguously to the data in the original JAMO record.

**Figure 4.**
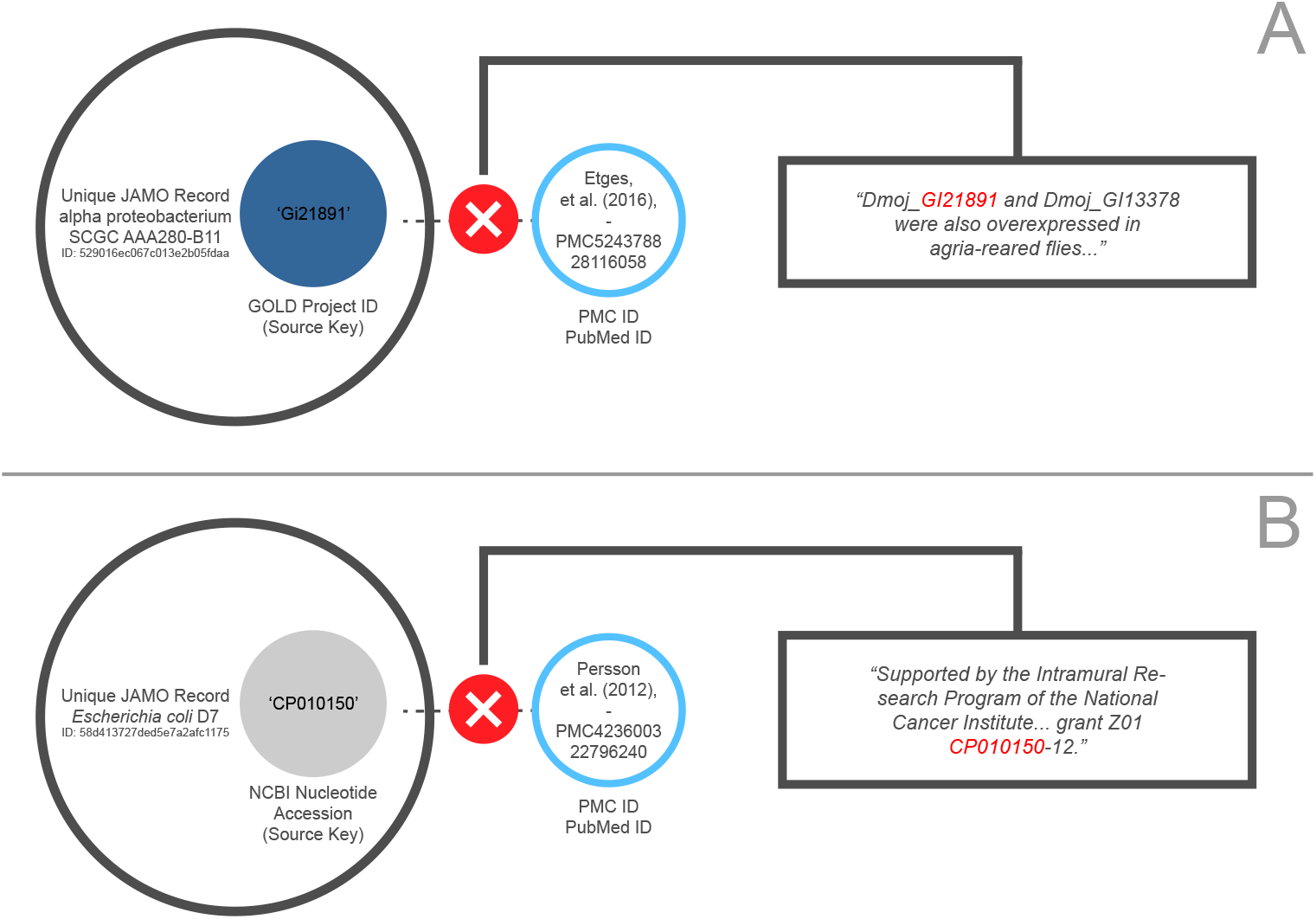
Examples of invalid audit trails. The error in example A results from a namespace collision with a grant number, while the error in example B results from a namespace collision between a GOLD Project ID and a FlyBase gene identifier. Some namespace collisions could only be resolved by retrieving and parsing full GenBank records, which degrades performance but improves accuracy.

Of the 489 keys used for searches in Dimensions, 348 (71.2%) resulted in hits. The total number of individual publications returned was 1,027, substantially greater than that returned through NCBI (Figure 5). The precision value for individual publications returned via Dimensions was evaluated using the same methods as with the initial sample and was determined to be 0.991. Combining these results with those from public indexing sources, the test set of keys generated hits on 1,234 unique publications. Over half of these were only identifiable through the proprietary Dimensions database.

**Figure 5.**
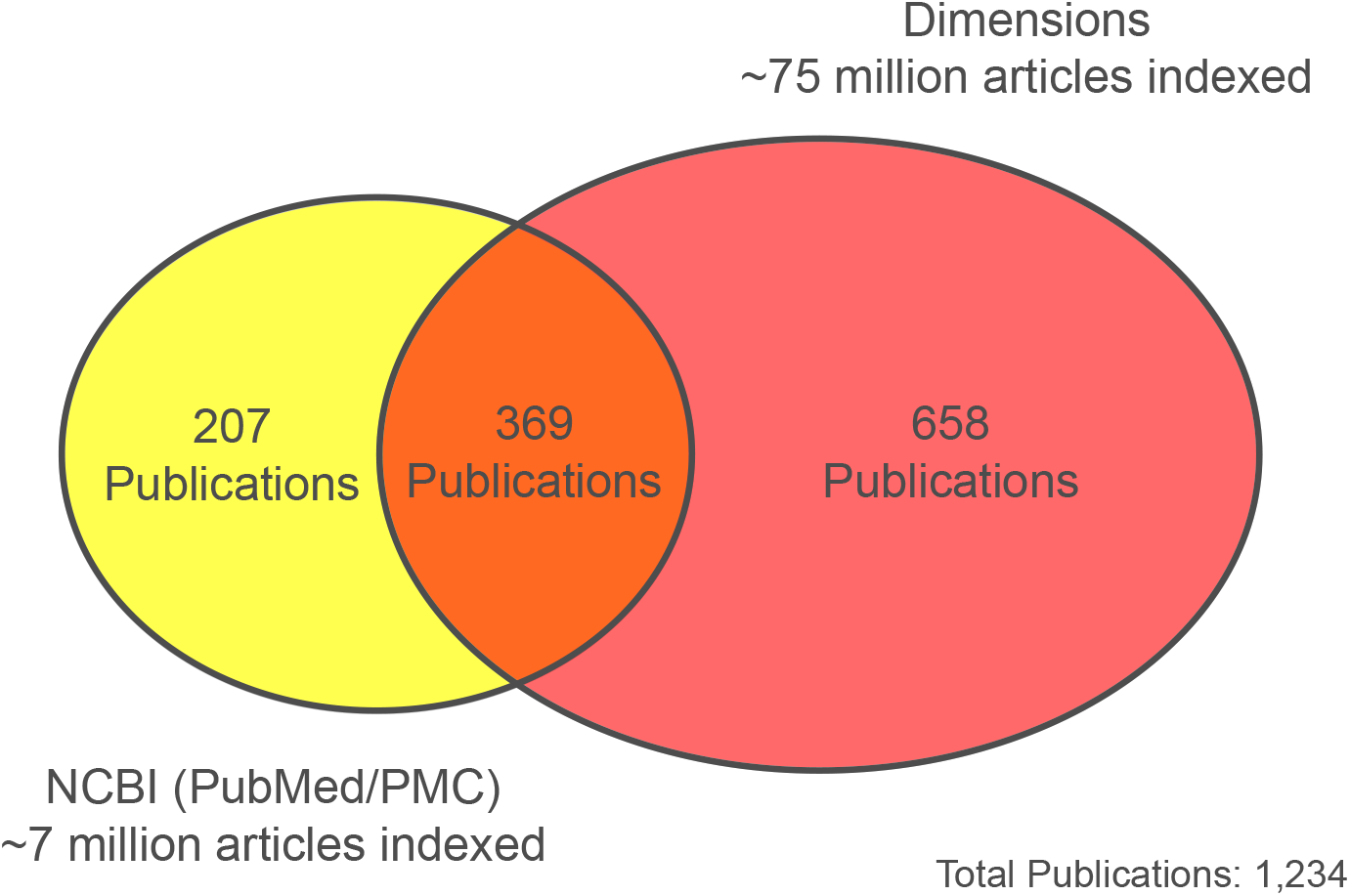
Comparison of publication results returned through the Data Citation Explorer using two distinct publication indexes and a stratified random sample of 300 JAMO records. The precision value for both sets of results was over 0.98.

## Discussion

### Corpus expansion and context analysis

Comparing results between the two publication data sources indicates the degree to which access to a larger corpus of full-text literature could increase the number of hits returned by the Data Citation Explorer’s search feature, particularly for disciplines not indexed by PubMed or PubMed Central. As all of the 207 publications returned by searching NCBI sources and not returned by Dimensions were later found to be indexed by Dimensions, it is likely that the differences in results returned by each were due to variations in search functionalities and full-text document indexing between the two providers. Investigation of these differences is beyond the scope of this work but represents an opportunity for future work. In addition to expanding the body of literature available to the DCE, further exploration of strategies for parsing non-standard supplemental material files included with scientific articles may increase this number still further. Internally, JGI can improve the comprehensiveness and quality of citable metadata indexed by JAMO for any given dataset to account for a wider range of possible citation methods.

Though the results of this evaluation indicate high levels of reliability, the system currently does not feature means for distinguishing the significance of one positively identified citation from another. Previous work has shown the efficacy of applying natural language processing (NLP) techniques to the textual context surrounding individual citations^17,18^. Similar techniques could be applied to evaluate DCE results beyond a simple validity assessment.

### Generalization

Expanding the service model embodied by the DCE to other disciplines (e.g. physics, earth sciences) is the next goal for future development and collaboration. Generalization of the service for use by other public data resources could maximize its impact and greatly improve wider knowledge of how public data is being used and by whom, supporting a Data Ecosystem that encourages connections among cross-organizational resources. While these results validate the conceptual underpinnings of the service’s architecture, much work remains to maximize the usefulness of the DCE’s service model to users beyond JGI and the biosciences field.

### Accessibility and collaboration

The citation results returned by the DCE are not currently accessible to the public, and much development work remains to be done to determine how best to make them accessible. The citations and audit paths are back-propagated into the JAMO database, which could serve as the integration point for making this data available via the JGI Data Portal (Figure 6).

**Figure 6.**
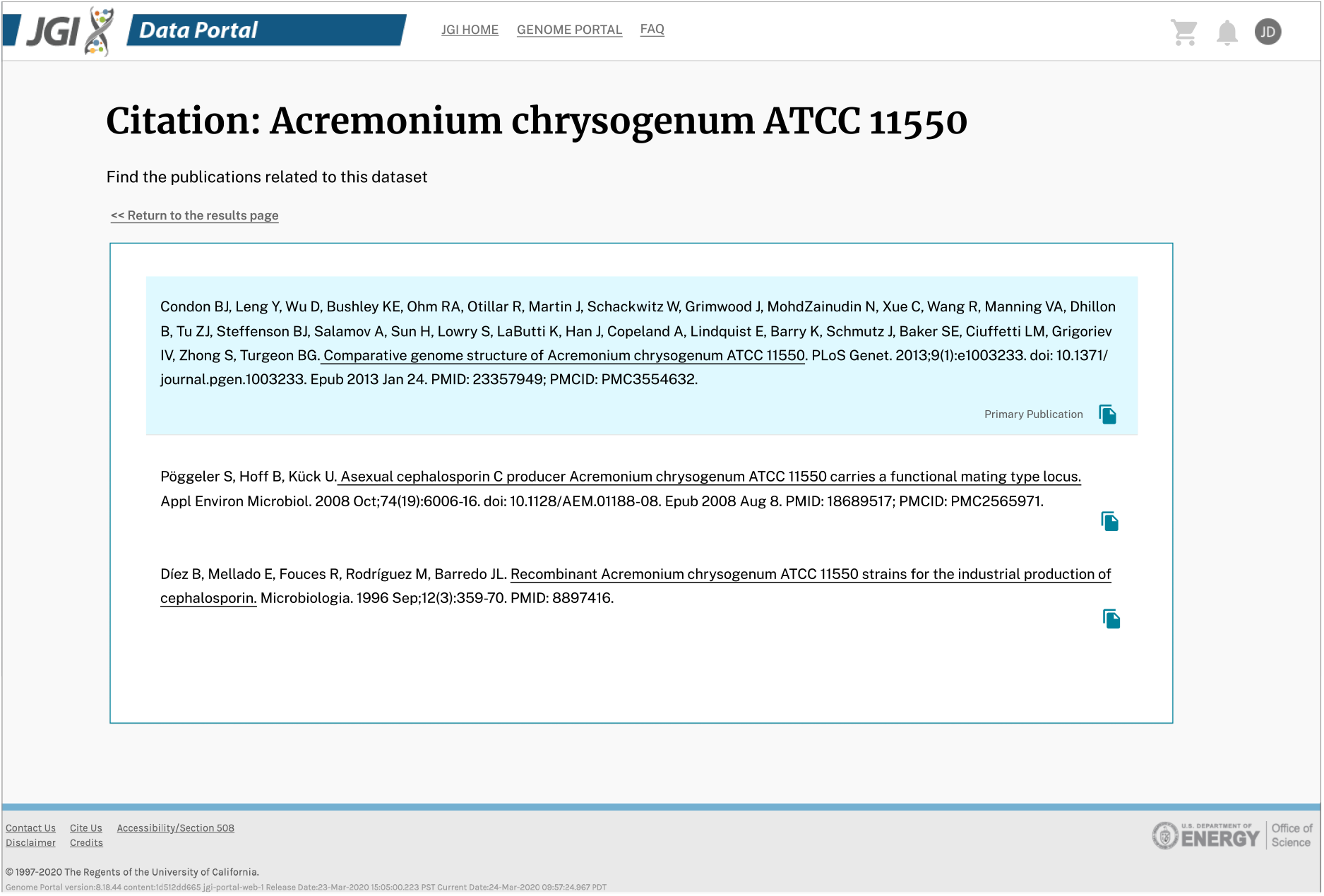
A mockup of how data citations discovered by the DCE might be presented to a user via the JGI Data Portal.

Some examples of potentially desirable features could include research-facilitating search tools for the Joint Genome Institute and associated DOE organizations like the National Microbiome Data Collaborative (NMDC)^19^ and the Systems Biology Knowledgebase (KBase)^20^. Generally available reports could describe who is using the work products produced by a given researcher and provide public-facing views that increase the visibility of JGI work products. From a user’s perspective, such features can illustrate previous use cases (or lack thereof) of any dataset of interest. The lessons learned from this project could be applied in other organizations that have an interest in mining highly focused genomic literature.

### FAIRness

The metadata and literature connections established by the DCE enable JGI to more equitably attribute credit to individual contributors for downstream outcomes of individual work products, though the specific metrics and methodology for doing so remain undetermined. Automated and standardized means for determining the extent of the service’s contributions to the findability, accessibility, interoperability, and reusability (FAIR) of public genomic data, similar to the framework developed by Wilkinson et al.^21^, would allow for continuous reassessment of the DCE for potential future updates.

## Methods

### JGI source data: JAMO

The data source for the DCE is the JGI Archive and Metadata Organizer (JAMO), which is JGI’s primary data management system. It manages most of the data assets the organization produces and caches associated metadata from the other data support systems, for example the Genomes On-Line Database (GOLD)^22^. JAMO also archives data to several geographically dispersed high-performance tape systems, manages the restore and purge policies of files on spinning disk, provides publish/subscribe services to internal pipelines, and creates a single connection point for all internal data systems to communicate with each other with regard to metadata services.

Metadata in JAMO are organized by JGI sequencing product/pipeline and file type, each of which has a defined dictionary of required and optional metadata. There are over 3,000 distinct metadata fields in JAMO across approximately 400 product, pipeline, and file types. The metadata in JAMO can be broken down into several classes: operational data, project and proposal information, genomic classifications, data ownership, data usability, internally produced public identifiers, and external public identifiers. Data provenance is also captured in the operational and project metadata.

The DCE pilot project focused on the metadata likely to be present in publications: NCBI GenBank Accessions, NCBI BioProject and BioSample Accessions, references to the NCBI taxonomy (from GOLD), IMG Taxon Object IDs (from JGI’s Integrated Microbial Genomes & Microbiomes system), SRA IDs (from NCBI’s Sequence Read Archive), and contact information (from JGI’s proposal system). After evaluation and testing, the initial production run of the DCE included 1.7 million selected JAMO records. These records were those that either were published at NCBI’s SRA or were downloaded by two or more distinct non-JGI users from JGI systems between February 22, 2019 and August 30, 2022.

### Citation discovery process

The citation discovery process runs in two phases: a metadata collection phase and a citation search phase (Figure 7). The metadata collection phase crawls genomic data repositories to accumulate new metadata that has been produced in downstream resources. This phase starts with an initial metadata registry from JGI’s JAMO database, then incrementally adds newly discovered metadata to the registry by crawling other data repositories via unique identifiers (e.g., genome assemblies, sequence data, sequence reads, bioprojects, biosamples) while respecting identifier hierarchies and cardinality. This process continues iteratively until no additional resources are discovered. As a new accession is discovered, its source is stored with the accession in a relational database in a form that preserves the graph representation of the DCE’s traversal through all discovered resources. The resulting graph may be used for evaluating the correctness of the system. Note that while some classes of identifier are ambiguous, these cases are generally mitigated by the addition of identifier namespaces that are respected by the relational database schema and associated business logic (Figure 8). In the second phase of the process, the DCE searches the Open Access literature for occurrences of any of the collected identifiers that identify the project of interest. The results of each search are added to the metadata registry and stored in the database. At this point, the citation discovery process is complete (Figures 9 and 10).

**Figure 7.**
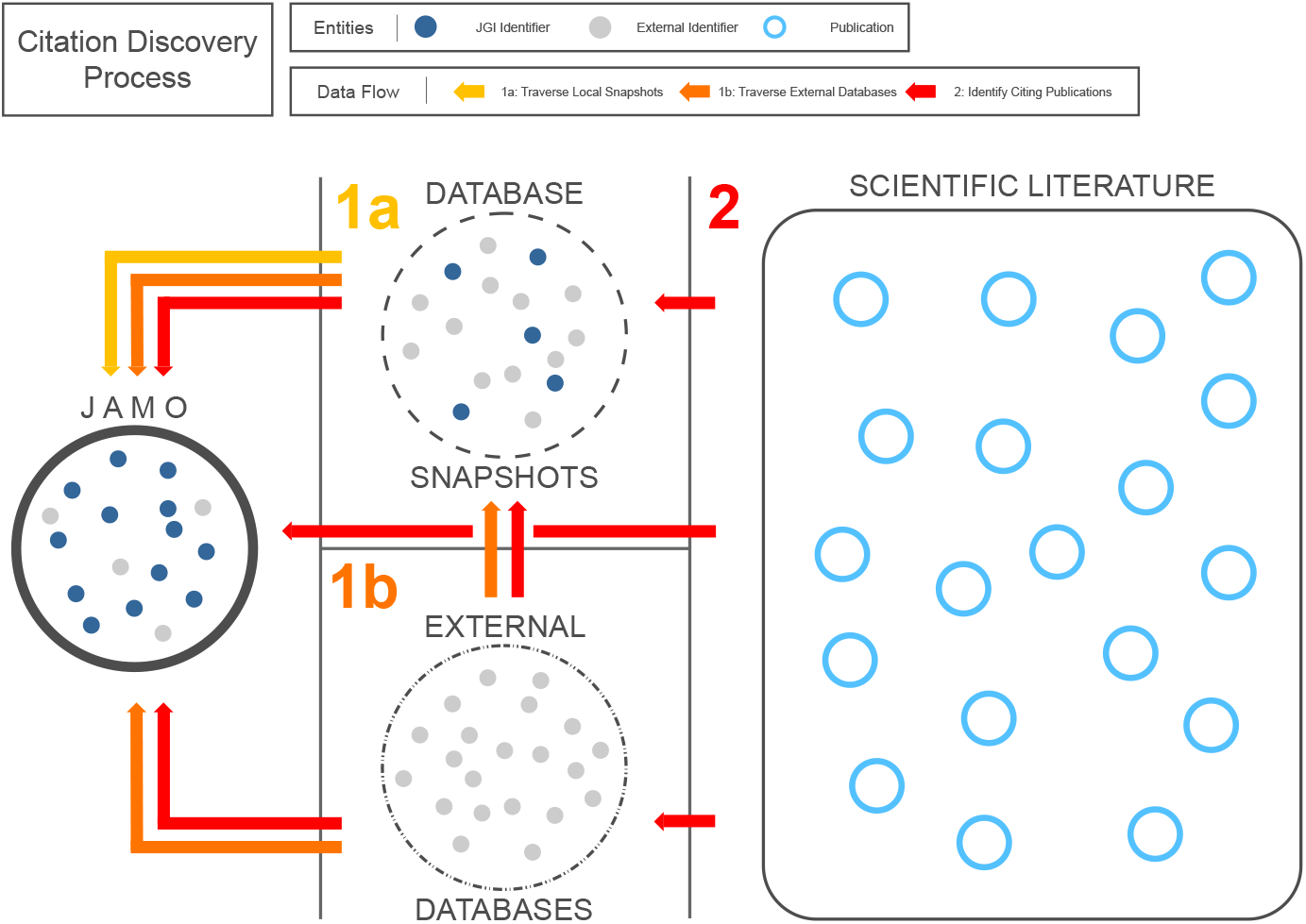
The citation discovery process for an individual JAMO record (the target genome) happens in two phases: (1) crawl genomic data repositories to accumulate new metadata that has been produced in downstream resources, and (2) search for occurrences of unique identifiers in publicly available literature.

**Figure 8.**
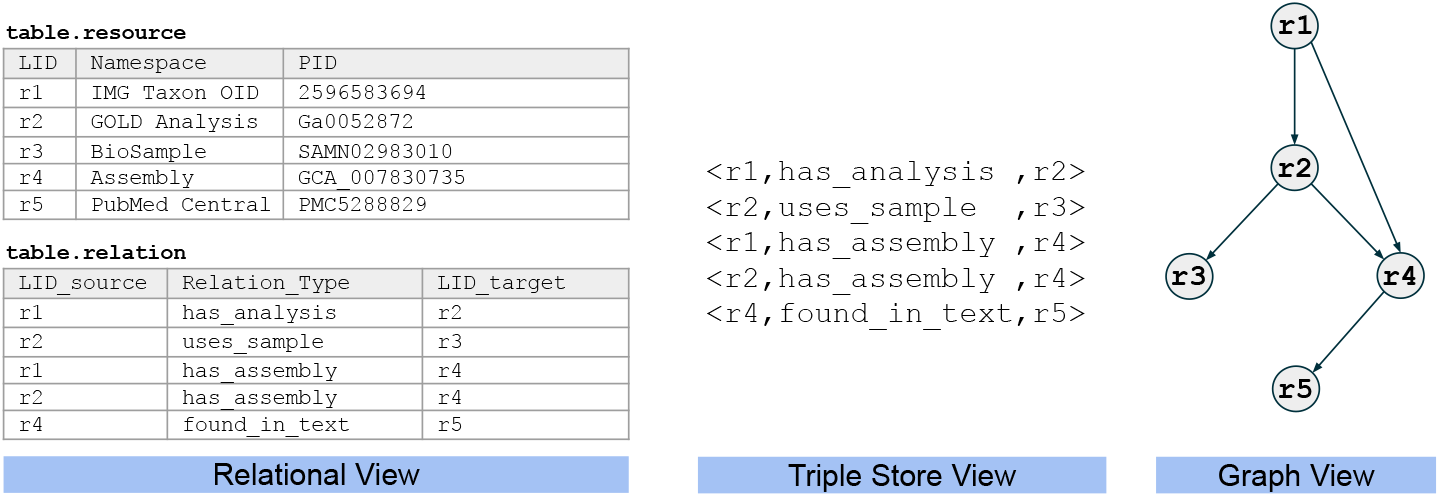
A sample audit trail for a set of connected genomic data resources as they are stored in the underlying namespace-aware relational database, retrieved as a set of triples via a SQL view, and reconstructed as a directed, acyclic graph. These audit trails support the validation of discovered citations. A depth-first traversal of the graph can identify all known paths between any two identifiers in the audit trail, as well as the shortest path between any two identifiers.

**Figure 9.**
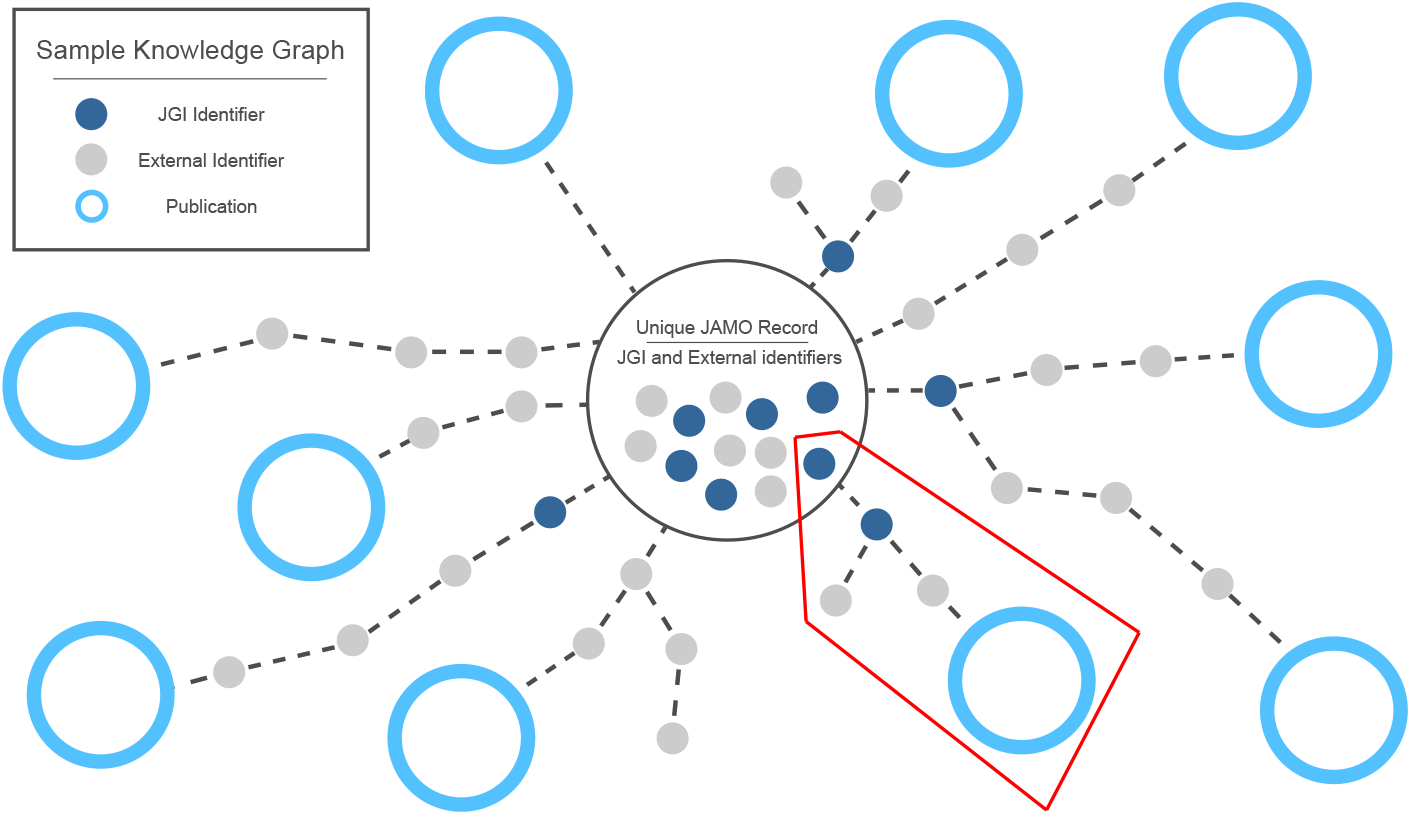
High-level depiction of a DCE knowledge graph as it branches out from a single JAMO record. Some data citations are connected via a single identifier, but others have more complex paths, often branching or forming redundant connections to publications. In this simplified image, for example, the highlighted branching path indicates that two separate JGI-external identifiers are both linked via a common JGI-internal identifier. One of these links to a publication, while the other does not.

**Figure 10.**
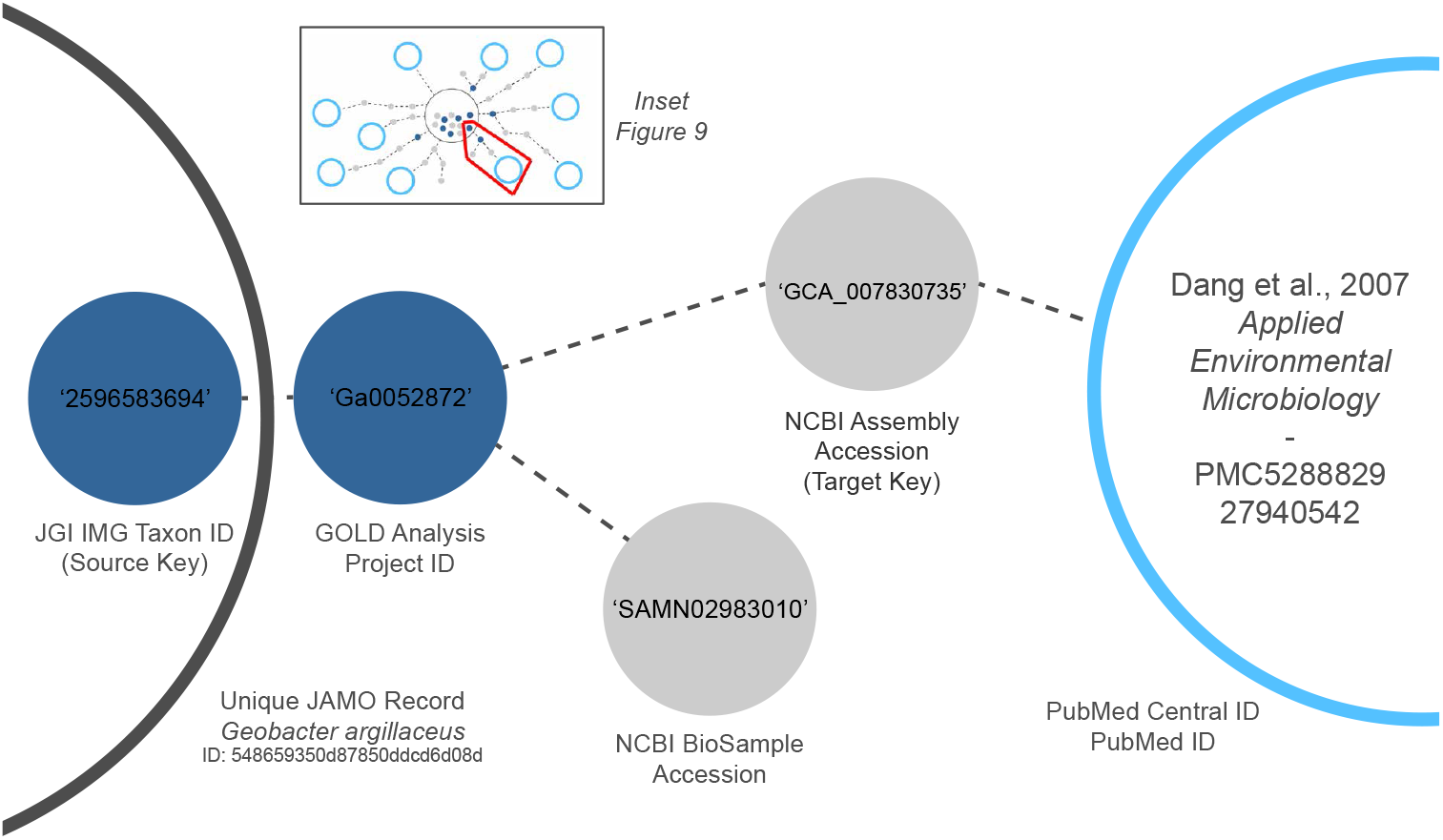
Sample audit trail for a single path to a publication that relies on a genome of interest (see inset). In this example, a JGI-internal identifier from the source JAMO record links to another JGI-internal identifier from a second resource. This second identifier, in turn, is linked through an external NCBI identifier to the citing publication, represented here by a PubMed Central identifier. The source and target resource from which each accession is discovered is stored as a triple, providing a traceable path that can explain how each resource was discovered. This is an important improvement over manual searches, as tracking the source of each individual metadata entry would be burdensome and error-prone for a human data curator. Note that a linkage to a separate external NCBI identifier forms a branch of this audit trail but does not link to any publications. These identifiers are also stored, as additional connections may be discovered at a later time.

### Evaluation procedures

In order to evaluate the system’s performance prior to large-scale production batches, an initial trial was performed by processing *∼*228,000 JAMO metadata records with the Data Citation Explorer. Records were selected if they represented genomic data downloaded by external users of JGI’s Genome Portal^23^ four or more times between the dates of February 22, 2019 and May 31, 2020. These dates span the period between when internal file request information became available and the start of the DCE trial. These metadata records describe publicly available genomic data that was produced by JGI or uploaded to its systems between January 2009 and May 2020. Of these *∼* 228,000 records, approximately*∼* 57,000 were linked to publications during the initial DCE trial.

As manual evaluation of all *∼*57,000 records with linked publications would be impossible, publications associated with a subset of records from the initial test run were selected. These were extracted from a stratified random sample of records to avoid overrepresenting JGI projects and data that have disproportionately high numbers of associated JAMO records. This would ensure that a diverse set of citations was available for evaluation. The stratified sample consisted of three groups of 100 records distinguished by particular characteristics of interest:

1. All records for data that were associated with a NCBI BioProject ID in the GOLD system. This is data that already had a set of linked publications indexed by GOLD and could thus be used for comparative purposes.
2. All records generated from JGI-sequenced data that were not registered at NCBI. The inclusion of this group would determine whether the DCE could find citations where external metadata IDs were limited or non-existent in the JAMO records.
3. All JAMO records generated from sequence data that were not produced by JGI, primarily gene annotations and similar data from external sources uploaded to JGI’s IMG/M system. The goal for this group was similar to Group 2 with the added benefit of determining whether or not the DCE would reject citations from upstream data sources (i.e., publications from the original sequence data).

The 300 records in the resulting sample were linked to 576 unique publications by the DCE (Figure 11).

**Figure 11.**
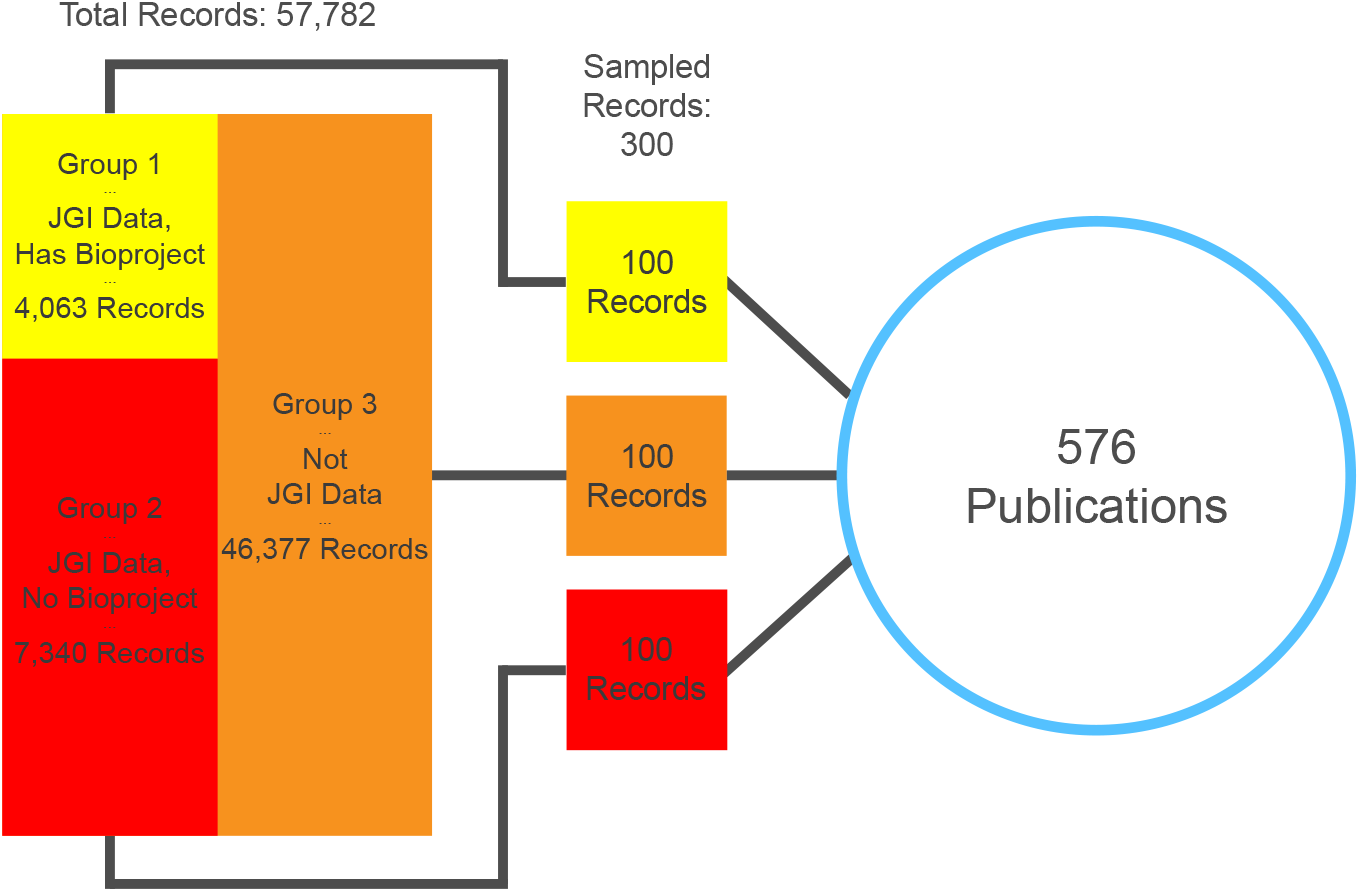
JAMO record sampling for manual evaluation. 300 JAMO records were selected from three stratified groups. In total, there were 998 unique audit trails connecting these records to 576 unique publications.

Following sample generation, the validity of each connection between a JAMO record and a publication was evaluated manually (Supplementary Tables S2, S3). A connection was considered “valid” only if the following conditions are met:

1. The key tied directly to or found within a target publication is an actual data identifier and not a false positive
2. The key tied directly to or found within a target publication unambiguously refers to the JGI data entity represented by the source JAMO record.

The initial proof of concept for the Data Citation Explorer was based on a core full-text corpus composed of the primary taxonomic literature of prokaryotes (primarily the *International Journal of Systematic and Evolutionary Microbiology* from 2005 through 2018 and *Standards in Genomic Sciences* v1-9). The corpus was expanded prior to the initial evaluation to include literature searches in two publicly available indexing services, NCBI’s PubMed and PubMed Central. To understand how the Data Citation Explorer performs with access to a larger corpus of publications, search results from the original corpus were compared with results from Digital Science’s Dimensions^15,16^ platform. This was done by using Dimensions’ public search interface (app.dimensions.ai) to run individual searches for each of the keys that led to hits in the initial sample. This process returned hits for a total of 1,027 publications.

## Data availability

Materials used for manual evaluation of DCE results can be found in Supplementary Tables S2, S3. Genomic metadata can be found at the Genomes OnLine Database (gold.jgi.doe.gov), the JGI Genome Portal (genome.jgi.doe.gov), the JGI Data Portal (data.jgi.doe.gov), Integrated Microbial Genomes & Microbiomes (img.jgi.doe.gov), Phytozome (phytozome.jgi.doe.gov), PhycoCosm (phycocosm.jgi.doe.gov), MyCosm (mycocosm.jgi.doe.gov), GenBank (ncbi.nlm.nih.gov/genbank), and the Sequence Read Archive (ncbi.nlm.nih.gov/sra). Publication data can be found at PubMed (pubmed.ncbi.nlm.nih.gov) and PubMed Central (www.ncbi.nlm.nih.gov/pmc). Additionally, the analyses include proprietary data sourced with permission from the Dimensions database (app.dimensions.ai), as well as information extracted from non-Open Access literature from text and data mining APIs that cannot be redistributed in raw form.

## Code availability

Researchers interested in access to the source code for the Data Citation Explorer may contact Charles Parker at. The authors will assist with any reasonable replication attempts for two years following publication.

## Acknowledgements

The authors would like to thank Tatyana Smirnova at JGI for contributing JGI Data Portal mockup images. The work conducted by the U.S. Department of Energy Joint Genome Institute (https://ror.org/04xm1d337), a DOE Office of Science User Facility, is supported by the Office of Science of the U.S. Department of Energy operated under Contract No. DE-AC02-05CH11231. Certain data included herein are derived from Clarivate Web of Science. Copyright Clarivate 2021. All rights reserved. This paper was written using data obtained on September 8, 2021 (DCE results) and January 22, 2024 (Genetics publications) from Digital Science’s Dimensions platform, available at https://app.dimensions.ai.

## Supplementary Information

Supplementary File 1

## Author contributions statement

N.B. designed and conducted DCE evaluation experiments. C.P. designed and developed the DCE. C.B. generated the evaluation sample and supervised DCE integration with JAMO. T.B.K.R. advised on JGI metadata usage, facilitated access to GOLD metadata, and provided guidance on NCBI and PubMed integration. H.S. supervised initial JAMO integration activities. G.G. and K.F. conceived of the presented idea and supervised the project. N.B., C.P., C.B., G.G. and K.F. contributed to the writing and revision of the manuscript.

