## Supplementary File 1 for "Identifying genomic data use with the Data Citation Explorer"

### Supplementary Information

**Table S1**: Column descriptions for Tables S2 and S3.
**Table S2**: Evaluation materials for DCE results retrieved through Dimensions
**Table S3**: Evaluation materials for DCE results retrieved through PubMed & PubMed Central

**Acknowledgement:**
Data in this file was collected from the Dimensions Software Platform. Digital Science. (2018-) Dimensions [Software] available from https://app.dimensions.ai. Accessed on September 8, 2021, under license agreement.

#### Table S1: Column descriptions for Tables S2 and S3.

| Table | Column | Description |
|---|---|---|
| S2 | jamo_id | unique hexidecimal JAMO record ID |
| S2 | publication_type | Type of linked publication |
| S2 | doi | publication doi |
| S2 | pmid | PubMed ID |
| S2 | pmcid | PubMed Central ID |
| S2 | t_key | Target key found in or directly linked to citing publication |
| S2 | valid | Validity (1=yes, 0=no) |
| S2 | pub_year | Year of publication |
| S3 | jamo_id | unique hexidecimal JAMO record ID |
| S3 | pmid | Title of JAMO record |
| S3 | valid | Validity (1=yes, 0=no) |
| S3 | s_key | Source key found within source JAMO record |
| S3 | s_key_type | Source key type |
| S3 | t_key | Target key found in or directly linked to citing publication |
| S3 | t_key_type | Target key type |
| S3 | final_db | Database in which the final hit in an audit trail was made |
| S3 | sample_group | Stratified JAMO record sample group |

Note: Each row in S2 and S3 refers to a pair between a JAMO record and a linked publication.

#### Table S2: Evaluation materials for DCE results retrieved through Dimensions

| jamo_id | publication_type | doi | pmid | pmcid | t_key | valid | pub_year |
|---|---|---|---|---|---|---|---|
| 51d4caf8067c014cd6eb0566 | Article | 10.1128/aem.06027-11 | 21856825 | PMC3194852 | ABDF00000000 | 1 | 2011 |
| 51d4caf8067c014cd6eb0566 | Article | 10.1186/s12864-017-4181-9 | 29025413 | PMC5639747 | PRJNA19983 | 1 | 2017 |
| 51d4caf8067c014cd6eb0566 | Article | 10.1186/s12864-020-07158-w | 33138770 | PMC7607812 | PRJNA19983 | 1 | 2020 |
| 51d4caf8067c014cd6eb0566 | Article | 10.1186/s12864-020-6653-6 | 32216757 | PMC7099791 | SAMN02744059 | 1 | 2020 |
| 51d4caf8067c014cd6eb0566 | Article | 10.3389/fmicb.2020.00200 | 32226413 | PMC7080844 | PRJNA19983 | 1 | 2020 |
| 51d4caf8067c014cd6eb0566 | Article | 10.1016/j.fgb.2014.02.002 | 24530791 | | PRJNA19983 | 1 | 2014 |
| 51d4caf8067c014cd6eb0566 | Article | 10.1186/gb-2011-12-4-r40 | 21501500 | PMC3218866 | ABDF00000000 | 1 | 2011 |
| 51d4caf8067c014cd6eb0566 | Article | 10.1080/21501203.2018.1478333 | 30181924 | PMC6115877 | ABDF00000000 | 1 | 2018 |
| 51d4c149067c014cd6ea7ede | Article | 10.1128/aem.00611-16 | 27235430 | PMC4984305 | CP002345 | 1 | 2016 |
| 51d4c149067c014cd6ea7ede | Article | 10.13057/biodiv/d200417 | | | CP002345 | 1 | 2019 |
| 51d4c149067c014cd6ea7ede | Article | 10.1016/j.biortech.2012.08.040 | 22989636 | | CP002345 | 1 | 2012 |
| 51d4c149067c014cd6ea7ede | Article | 10.1016/j.biortech.2017.06.053 | 28637165 | PMC7362340 | CP002345 | 1 | 2017 |
| 51d4c149067c014cd6ea7ede | Article | 10.4056/sigs.1503846 | 21475585 | PMC3072083 | CP002345 | 1 | 2011 |
| 51d4c149067c014cd6ea7ede | Article | 10.1093/femsec/fiv030 | 25873605 | | CP002345 | 1 | 2015 |
| 51d4c149067c014cd6ea7ede | Article | 10.1093/femsec/fiz001 | 30649346 | | CP002345 | 1 | 2019 |
| 51d4c149067c014cd6ea7ede | Article | 10.3390/genes11080878 | 32756341 | PMC7465726 | CP002345 | 1 | 2020 |
| 51d4c149067c014cd6ea7ede | Article | 10.1099/ijs.0.000250 | 25866024 | | CP002345 | 1 | 2015 |
| 51d4c149067c014cd6ea7ede | Article | 10.1099/ijs.0.032508-0 | 22180609 | | CP002345 | 1 | 2011 |
| 51d4c149067c014cd6ea7ede | Article | 10.1099/ijs.0.056812-0 | 24523447 | | CP002345 | 1 | 2014 |
| 51d4c149067c014cd6ea7ede | Article | 10.1099/ijs.0.066902-0 | 25048210 | | CP002345 | 1 | 2014 |
| 51d4c149067c014cd6ea7ede | Article | 10.1099/ijsem.0.000617 | 26377180 | | CP002345 | 1 | 2015 |
| 51d4c149067c014cd6ea7ede | Article | 10.1093/jac/dkr251 | 21680581 | | CP002345 | 1 | 2011 |
| 51d4c149067c014cd6ea7ede | Article | 10.1016/s2095-3119(15)61272-2 | | | CP002345 | 1 | 2016 |
| 51d4c149067c014cd6ea7ede | Article | 10.1016/s2095-3119(19)62609-2 | | | CP002345 | 1 | 2019 |
| 51d4c149067c014cd6ea7ede | Article | 10.1264/jsme2.me14142 | 25736980 | PMC4356463 | CP002345 | 1 | 2015 |
| 51d4c149067c014cd6ea7ede | Chapter | 10.1007/978-3-642-38954-2_132 | | | CP002345 | 1 | 2014 |
| 51d4c149067c014cd6ea7ede | Article | 10.11606/d.11.2013.tde-16122013-172059 | | | CP002345 | 1 | 2013 |
| 5a8386c364d0b326cdd1dd2d | Article | 10.1186/s13068-019-1569-6 | 31572496 | PMC6757388 | PRJNA250595 | 1 | 2019 |
| 51d4c3e3067c014cd6eaa5e7 | Chapter | 10.1007/978-3-642-38954-2_162 | | | PRJNA45817 | 1 | 2014 |
| 51d4bf7c067c014cd6ea6afe | Article | 10.14201/gredos.127301 | | | PRJNA65273 | 1 | 2014 |
| 51d4bdc6067c014cd6ea5e5c | Article | 10.1007/s00253-012-4285-8 | 22854892 | | ACXX00000000 | 1 | 2012 |
| 51d4bdc6067c014cd6ea5e5c | Article | 10.1099/ijsem.0.002966 | 30124399 | | ACXX00000000 | 1 | 2018 |
| 51d4bdc6067c014cd6ea5e5c | Article | 10.1128/jb.01064-10 | 20889752 | PMC3008519 | ACXX00000000 | 1 | 2010 |
| 51d4bdc6067c014cd6ea5e5c | Chapter | 10.1128/9781555816827.ch17 | | | ACXX00000000 | 1 | 2010 |
| 5621c9320d878540fd70755e | Article | 10.1128/jb.01398-12 | 23045486 | PMC3486092 | CP003704 | 1 | 2012 |
| 5621c9320d878540fd70755e | Article | 10.1128/jb.01398-12 | 23045486 | PMC3486092 | CP003707 | 1 | 2012 |
| 5621c9320d878540fd70755e | Chapter | 10.1007/978-981-13-8739-5_14 | | | CP003704 | 1 | 2019 |
| 51d4c5d0067c014cd6eac2bd | Article | 10.1007/s10482-013-9970-9 | 23851717 | | CP001688 | 1 | 2013 |
| 51d4c5d0067c014cd6eac2bd | Article | 10.1155/2010/485051 | 20672053 | PMC2905702 | CP001688 | 1 | 2010 |
| 51d4c5d0067c014cd6eac2bd | Article | 10.1007/s00705-013-1970-6 | 24395078 | | CP001688 | 1 | 2014 |
| 51d4c5d0067c014cd6eac2bd | Article | 10.1186/1471-2164-13-478 | 22978470 | PMC3528665 | CP001688 | 1 | 2012 |
| 51d4c5d0067c014cd6eac2bd | Article | 10.4056/sigs.23264 | 21304651 | PMC3035229 | CP001688 | 1 | 2009 |
| 51d4c5d0067c014cd6eac2bd | Article | 10.4056/sigs.42644 | 21304667 | PMC3035239 | CP001688 | 1 | 2009 |
| 51d4c5d0067c014cd6eac2bd | Article | 10.1099/ijs.0.029298-0 | 21296924 | | CP001688 | 1 | 2011 |
| 51d4c5d0067c014cd6eac2bd | Article | 10.1099/ijs.0.058164-0 | 24425819 | | CP001688 | 1 | 2013 |
| 51d4c5d0067c014cd6eac2bd | Article | 10.1099/ijsem.0.000781 | 26596884 | | CP001688 | 1 | 2015 |
| 51d4c5d0067c014cd6eac2bd | Article | 10.1099/ijsem.0.001663 | 28475027 | | CP001688 | 1 | 2017 |
| 51d4c5d0067c014cd6eac2bd | Article | 10.1128/jb.00124-10 | 20400546 | PMC2901701 | CP001688 | 1 | 2010 |
| 51d4c5d0067c014cd6eac2bd | Article | 10.1016/j.micres.2010.07.002 | 20869220 | | CP001688 | 1 | 2010 |
| 51d4c5d0067c014cd6eac2bd | Article | 10.1111/mmi.13204 | 26331239 | | CP001688 | 1 | 2015 |
| 51d4c5d0067c014cd6eac2bd | Article | 10.1093/nar/gks215 | 22396526 | PMC3384331 | CP001688 | 1 | 2012 |
| 51d4c5d0067c014cd6eac2bd | Chapter | 10.1002/9781118960608.fbm00293 | | | CP001688 | 1 | 2017 |
| 51d4c5d0067c014cd6eac2bd | Chapter | 10.1002/9781118960608.gbm00481.pub2 | | | CP001688 | 1 | 2019 |
| 51d4c5d0067c014cd6eac2bd | Chapter | 10.1002/9781118960608.gbm01342 | | | CP001688 | 1 | 2016 |
| 51d490e8067c014cd6e9fdbd | Article | 10.1111/efp.12304 | | | PRJNA74753 | 1 | 2016 |
| 51d490e8067c014cd6e9fdbd | Article | 10.3390/microorganisms7100420 | 31590374 | PMC6843257 | PRJNA74753 | 1 | 2019 |
| 51d4bdac067c014cd6ea5d94 | Article | 10.1093/aob/mcs206 | 22989463 | PMC3503493 | CP001622 | 1 | 2012 |
| 51d4bdac067c014cd6ea5d94 | Article | 10.1007/s00253-015-6515-3 | 25776061 | | Gc01039 | 1 | 2015 |
| 51d4bdac067c014cd6ea5d94 | Article | 10.1016/j.apsoil.2017.06.030 | 29104370 | PMC5637928 | CP001622 | 1 | 2017 |
| 51d4bdac067c014cd6ea5d94 | Article | 10.5194/bg-10-8269-2013 | | | CP001622 | 1 | 2013 |

| | | | | | | | |
|---|---|---|---|---|---|---|---|
| 51d4bdac067c014cd6ea5d94 | Article | 10.1111/j.1462-2920.2010.02393.x | 21176055 | | CP001622 | 1 | 2010 |
| 51d4bdac067c014cd6ea5d94 | Article | 10.1186/s40793-017-0220-z | 28163823 | PMC5278577 | CP001622 | 1 | 2017 |
| 51d4bdac067c014cd6ea5d94 | Article | 10.4056/sigs.852027 | 21304718 | PMC3035295 | CP001622 | 1 | 2010 |
| 51d4bdac067c014cd6ea5d94 | Article | 10.4056/sigs.2675953 | 23409217 | PMC3558968 | Gc01039 | 1 | 2012 |
| 51d4bdac067c014cd6ea5d94 | Article | 10.4056/sigs.852027 | 21304718 | PMC3035295 | Gc01039 | 1 | 2010 |
| 51d4bdac067c014cd6ea5d94 | Article | 10.1093/gbe/evx255 | 29220487 | PMC5739047 | CP001622 | 1 | 2017 |
| 51d4bdac067c014cd6ea5d94 | Article | 10.1016/j.meegid.2012.06.009 | 22771358 | | CP001622 | 1 | 2012 |
| 51d4bdac067c014cd6ea5d94 | Article | 10.1099/ijs.0.033555-0 | 22081714 | | CP001622 | 1 | 2011 |
| 51d4bdac067c014cd6ea5d94 | Article | 10.1128/jb.06192-11 | 22037403 | PMC3256650 | CP001622 | 1 | 2011 |
| 51d4bdac067c014cd6ea5d94 | Article | 10.1016/j.ympev.2014.06.006 | 24952318 | | CP001622 | 1 | 2014 |
| 51d4bdac067c014cd6ea5d94 | Article | 10.3955/046.094.0205 | | | CP001622 | 1 | 2020 |
| 51d4bdac067c014cd6ea5d94 | Article | 10.1016/j.plasmid.2015.02.003 | 25752994 | | CP001622 | 1 | 2015 |
| 51d4bdac067c014cd6ea5d94 | Article | 10.1016/j.resmic.2013.05.002 | 23764913 | | CP001622 | 1 | 2013 |
| 51d4bdac067c014cd6ea5d94 | Article | 10.1134/s2079059712020025 | | | CP001622 | 1 | 2012 |
| 51d4bdac067c014cd6ea5d94 | Article | 10.1016/j.soilbio.2013.01.009 | | | Gc01039 | 1 | 2013 |
| 51d4bdac067c014cd6ea5d94 | Article | 10.1007/s13199-015-0365-8 | | | CP001622 | 1 | 2015 |
| 51d4bdac067c014cd6ea5d94 | Article | 10.1016/j.syapm.2010.11.015 | 21306854 | | CP001622 | 1 | 2011 |
| 51d4bdac067c014cd6ea5d94 | Article | 10.1016/j.syapm.2012.02.003 | 22429391 | | CP001622 | 1 | 2012 |
| 51d4bdac067c014cd6ea5d94 | Article | 10.1016/j.syapm.2020.126127 | 32847793 | | CP001622 | 1 | 2020 |
| 51d4bdac067c014cd6ea5d94 | Chapter | 10.1002/9780470015902.a0021157 | | | CP001622 | 1 | 2010 |
| 51d4bdac067c014cd6ea5d94 | Chapter | 10.1007/978-3-642-21680-0_5 | | | CP001622 | 1 | 2011 |
| 51d4bdac067c014cd6ea5d94 | Chapter | 10.1007/978-3-642-30194-0_73 | | | CP001622 | 1 | 2013 |
| 51d4c514067c014cd6eab6e6 | Chapter | 10.1007/978-3-642-38954-2_384 | | | PRJNA33371 | 1 | 2014 |
| 5592b1fb0d87854fee5ff0e4 | Article | 10.1128/genomea.00605-13 | 23929491 | PMC3738907 | CM001240 | 1 | 2013 |
| 51d4c81b067c014cd6eae393 | Chapter | 10.1007/978-3-642-38954-2_162 | | | PRJNA29431 | 1 | 2014 |
| 53a2653d0d878514c2d0f1b3 | Article | 10.3389/fmicb.2018.02007 | 30186281 | PMC6113628 | SAMN02745412 | 1 | 2018 |
| 53a2653d0d878514c2d0f1b3 | Article | 10.3390/genes11101166 | 33022985 | PMC7601586 | PRJNA234788 | 1 | 2020 |
| 51d4bc76067c014cd6ea5585 | Article | 10.1146/annurev-ento-031616-035509 | 27860524 | | PRJNA67455 | 1 | 2016 |
| 51d4bc76067c014cd6ea5585 | Article | 10.3390/v11040351 | 30999558 | PMC6520771 | PRJNA67455 | 1 | 2019 |
| 51d4bf8a067c014cd6ea6b8e | Article | 10.4056/sigs.3476977 | 23991262 | PMC3746429 | CP001819 | 1 | 2013 |
| 51d4bf8a067c014cd6ea6b8e | Article | 10.3389/fmicb.2018.02007 | 30186281 | PMC6113628 | SAMN02598426 | 1 | 2018 |
| 51d4bf8a067c014cd6ea6b8e | Article | 10.1099/ijsem.0.001261 | 27373977 | | CP001819 | 1 | 2016 |
| 51d4bf8a067c014cd6ea6b8e | Article | 10.1099/ijsem.0.001701 | 27902298 | | CP001819 | 1 | 2017 |
| 51d4bf8a067c014cd6ea6b8e | Article | 10.1099/ijsem.0.002584 | 29458505 | | CP001819 | 1 | 2018 |
| 51d4bf8a067c014cd6ea6b8e | Article | 10.1371/journal.pone.0033800 | 22479445 | PMC3315585 | CP001819 | 1 | 2012 |
| 51d4bf8a067c014cd6ea6b8e | Article | 10.1016/j.syapm.2018.02.003 | 29567394 | | CP001819 | 1 | 2018 |
| 51d4bf8a067c014cd6ea6b8e | Article | 10.1038/ja.2011.57 | 21792208 | | CP001819 | 1 | 2011 |
| 51d4ca02067c014cd6eaf9a9 | Article | 10.18946/jssm.72.1_7 | | | CP002629 | 1 | 2018 |
| 51d4ca02067c014cd6eaf9a9 | Article | 10.4056/sigs.2064705 | 21886866 | PMC3156406 | CP002629 | 1 | 2011 |
| 51d4ca02067c014cd6eaf9a9 | Article | 10.3389/fmicb.2012.00026 | 22347219 | PMC3271277 | CP002629 | 1 | 2012 |
| 51d4ca02067c014cd6eaf9a9 | Article | 10.3389/fmicb.2012.00404 | 23316187 | PMC3541049 | CP002629 | 1 | 2012 |
| 51d4ca02067c014cd6eaf9a9 | Article | 10.1099/ijs.0.064360-0 | 24944334 | | CP002629 | 1 | 2014 |
| 51d4ca02067c014cd6eaf9a9 | Article | 10.1099/ijsem.0.001083 | 27082267 | | CP002629 | 1 | 2016 |
| 51d4ca02067c014cd6eaf9a9 | Article | 10.1007/s12275-012-1342-z | 22538648 | | CP002629 | 1 | 2012 |
| 51d4ca02067c014cd6eaf9a9 | Article | 10.1134/s002626171404016x | | | CP002629 | 1 | 2014 |
| 51d4ca02067c014cd6eaf9a9 | Article | 10.1007/s11104-016-2851-z | | | CP002629 | 1 | 2016 |
| 51d4ca02067c014cd6eaf9a9 | Chapter | 10.1002/9781118960608.fbm00192.pub2 | | | CP002629 | 1 | 2020 |
| 51d4ca02067c014cd6eaf9a9 | Chapter | 10.1002/9781118960608.fbm00193.pub2 | | | CP002629 | 1 | 2020 |
| 51d4ca02067c014cd6eaf9a9 | Chapter | 10.1002/9781118960608.fbm00194.pub2 | | | CP002629 | 1 | 2020 |
| 51d4ca02067c014cd6eaf9a9 | Chapter | 10.1002/9781118960608.fbm00196.pub2 | | | CP002629 | 1 | 2020 |
| 51d4ca02067c014cd6eaf9a9 | Chapter | 10.1002/9781118960608.fbm00197.pub2 | | | CP002629 | 1 | 2020 |
| 51d4ca02067c014cd6eaf9a9 | Chapter | 10.1002/9781118960608.fbm00198.pub2 | | | CP002629 | 1 | 2020 |
| 51d4ca02067c014cd6eaf9a9 | Chapter | 10.1002/9781118960608.fbm00324 | | | CP002629 | 1 | 2019 |
| 51d4ca02067c014cd6eaf9a9 | Chapter | 10.1002/9781118960608.fbm00331 | | | CP002629 | 1 | 2020 |
| 51d4ca02067c014cd6eaf9a9 | Chapter | 10.1002/9781118960608.fbm00334 | | | CP002629 | 1 | 2020 |
| 51d4ca02067c014cd6eaf9a9 | Chapter | 10.1002/9781118960608.fbm00336 | | | CP002629 | 1 | 2020 |
| 51d4ca02067c014cd6eaf9a9 | Chapter | 10.1002/9781118960608.fbm00341 | | | CP002629 | 1 | 2020 |
| 51d4ca02067c014cd6eaf9a9 | Chapter | 10.1002/9781118960608.fbm00365 | | | CP002629 | 1 | 2020 |
| 51d4ca02067c014cd6eaf9a9 | Chapter | 10.1002/9781118960608.gbm01061.pub2 | | | CP002629 | 1 | 2019 |
| 51d4ca02067c014cd6eaf9a9 | Chapter | 10.1007/978-3-642-39044-9_269 | | | CP002629 | 1 | 2014 |
| 51d4c6fe067c014cd6ead5e1 | Article | 10.1007/s00284-019-01823-4 | 31792570 | | CP002305 | 1 | 2019 |
| 51d4c6fe067c014cd6ead5e1 | Article | 10.4056/sigs.1413518 | 21475582 | PMC3072089 | CP002305 | 1 | 2011 |
| 51d4c6fe067c014cd6ead5e1 | Article | 10.1099/ijs.0.041533-0 | 22544798 | | CP002305 | 1 | 2012 |

| | | | | | | | |
|---|---|---|---|---|---|---|---|
| 51d4c6fe067c014cd6ead5e1 | Article | 10.1099/ijs.0.052423-0 | 23687060 | | CP002305 | 1 | 2013 |
| 51d4c6fe067c014cd6ead5e1 | Article | 10.1099/ijs.0.053439-0 | 23907222 | | CP002305 | 1 | 2013 |
| 51d4c6fe067c014cd6ead5e1 | Article | 10.1099/ijsem.0.000577 | 26346054 | | CP002305 | 1 | 2015 |
| 51d4c6fe067c014cd6ead5e1 | Article | 10.1099/ijsem.0.000582 | 26341781 | | CP002305 | 1 | 2015 |
| 51d4c6fe067c014cd6ead5e1 | Article | 10.1099/ijsem.0.001038 | 27031260 | | CP002305 | 1 | 2016 |
| 51d4c6fe067c014cd6ead5e1 | Article | 10.1099/ijsem.0.001205 | 27264529 | | CP002305 | 1 | 2016 |
| 51d4c6fe067c014cd6ead5e1 | Article | 10.1099/ijsem.0.001490 | 27613103 | | CP002305 | 1 | 2016 |
| 51d4c6fe067c014cd6ead5e1 | Article | 10.1099/ijsem.0.001690 | 27902275 | | CP002305 | 1 | 2017 |
| 51d4c6fe067c014cd6ead5e1 | Article | 10.1099/ijsem.0.001840 | 28150574 | | CP002305 | 1 | 2017 |
| 51d4c6fe067c014cd6ead5e1 | Article | 10.1099/ijsem.0.001899 | 28809153 | | CP002305 | 1 | 2017 |
| 51d4c6fe067c014cd6ead5e1 | Article | 10.1099/ijsem.0.002041 | 28820097 | | CP002305 | 1 | 2017 |
| 51d4c6fe067c014cd6ead5e1 | Article | 10.1007/s12649-017-0122-8 | | | CP002305 | 1 | 2017 |
| 51d4be7a067c014cd6ea6341 | Article | 10.1007/s10482-014-0311-4 | 25344421 | | CP002536 | 1 | 2014 |
| 51d4be7a067c014cd6ea6341 | Article | 10.1002/arch.21190 | 25195523 | | CP002536 | 1 | 2014 |
| 51d4be7a067c014cd6ea6341 | Article | 10.4056/sigs.2756060 | 22768367 | PMC3387796 | CP002536 | 1 | 2012 |
| 51d4be7a067c014cd6ea6341 | Article | 10.4056/sigs.2836114 | | PMC3387800 | CP002536 | 0 | 2012 |
| 51d4be7a067c014cd6ea6341 | Article | 10.3389/fmicb.2019.02083 | 31608019 | PMC6767994 | SAMN00016729 | 1 | 2019 |
| 51d4be7a067c014cd6ea6341 | Article | 10.1099/ijs.0.066324-0 | 25351880 | | CP002536 | 1 | 2014 |
| 51d4be7a067c014cd6ea6341 | Article | 10.1099/ijs.0.066555-0 | 25256704 | | CP002536 | 1 | 2014 |
| 51d4be7a067c014cd6ea6341 | Article | 10.1099/ijsem.0.000439 | 26297659 | | CP002536 | 1 | 2015 |
| 51d4be7a067c014cd6ea6341 | Article | 10.1099/ijsem.0.002683 | 29504923 | | CP002536 | 1 | 2018 |
| 51d4be7a067c014cd6ea6341 | Article | 10.1371/journal.pone.0034458 | 22470573 | PMC3314630 | CP002536 | 1 | 2012 |
| 51d4910b067c014cd6e9ff55 | Article | 10.3390/proceedings2110650 | | | CP002530 | 1 | 2018 |
| 51d4910b067c014cd6e9ff55 | Article | 10.4056/sigs.1704212 | 21677856 | PMC3111984 | CP002530 | 1 | 2011 |
| 51d4910b067c014cd6e9ff55 | Article | 10.1371/journal.pone.0206484 | 31509535 | PMC6738582 | CP002530 | 1 | 2019 |
| 59138c6e7ded5e1e49fff62e | Article | 10.3389/fmicb.2016.00363 | 27047476 | PMC4797314 | Gs0090290 | 1 | 2016 |
| 59138c6e7ded5e1e49fff62e | Article | 10.7717/peerj.3134 | 28396823 | PMC5385130 | PRJNA269163 | 1 | 2017 |
| 59138c6e7ded5e1e49fff62e | Article | 10.7717/peerj.3134 | 28396823 | PMC5385130 | SAMN03166137 | 1 | 2017 |
| 59138c6e7ded5e1e49fff62e | Article | 10.1038/sdata.2017.37 | 28350381 | PMC5369317 | PRJNA269163 | 1 | 2017 |
| 59138c6e7ded5e1e49fff62e | Article | 10.1038/sdata.2017.37 | 28350381 | PMC5369317 | SAMN03166137 | 1 | 2017 |
| 51d4ce4c067c014cd6eb3334 | Article | 10.1007/s10482-020-01424-3 | 32399714 | PMC7716859 | CP002353 | 1 | 2020 |
| 51d4ce4c067c014cd6eb3334 | Article | 10.1007/s10482-020-01471-w | 32936355 | | CP002353 | 1 | 2020 |
| 51d4ce4c067c014cd6eb3334 | Article | 10.1186/1944-3277-9-10 | 25780503 | PMC4334474 | CP002353 | 1 | 2014 |
| 51d4ce4c067c014cd6eb3334 | Article | 10.4056/sigs.1533840 | 21475588 | PMC3072084 | CP002353 | 1 | 2011 |
| 51d4ce4c067c014cd6eb3334 | Article | 10.3389/fmicb.2017.00412 | 28360896 | PMC5352709 | PRJNA32825 | 1 | 2017 |
| 51d4ce4c067c014cd6eb3334 | Article | 10.1099/ijs.0.000009 | 25479950 | | CP002353 | 1 | 2014 |
| 51d4ce4c067c014cd6eb3334 | Article | 10.1099/ijsem.0.000767 | 26559645 | | CP002353 | 1 | 2015 |
| 51d4ce4c067c014cd6eb3334 | Article | 10.1099/ijsem.0.002846 | 29873630 | | CP002353 | 1 | 2018 |
| 51d4ce4c067c014cd6eb3334 | Article | 10.1038/s41429-018-0035-1 | 29467380 | | CP002353 | 1 | 2018 |
| 59cd8a497ded5e2f18696152 | Article | 10.1080/00275514.2019.1670018 | 31750788 | | PRJNA234811 | 1 | 2019 |
| 51d55e7a067c014cd6f12f7c | Article | 10.1007/s00253-015-6515-3 | 25776061 | | Gc00590 | 1 | 2015 |
| 51d55e7a067c014cd6f12f7c | Article | 10.1016/j.apsoil.2017.06.030 | 29104370 | PMC5637928 | CP000738 | 1 | 2017 |
| 51d55e7a067c014cd6f12f7c | Article | 10.1186/1471-2164-10-437 | 19758436 | PMC2761423 | CP000738 | 1 | 2009 |
| 51d55e7a067c014cd6f12f7c | Article | 10.1139/w10-086 | 21164569 | | CP000738 | 1 | 2010 |
| 51d55e7a067c014cd6f12f7c | Article | 10.1007/s00284-016-1149-y | 27770191 | | CP000738 | 1 | 2016 |
| 51d55e7a067c014cd6f12f7c | Article | 10.1111/j.1462-2920.2007.01434.x | 17803764 | PMC2040194 | CP000738 | 1 | 2007 |
| 51d55e7a067c014cd6f12f7c | Article | 10.4056/sigs.43526 | 21304680 | PMC3035259 | CP000738 | 1 | 2010 |
| 51d55e7a067c014cd6f12f7c | Article | 10.4056/sigs.2675953 | 23409217 | PMC3558968 | Gc00590 | 1 | 2012 |
| 51d55e7a067c014cd6f12f7c | Article | 10.4056/sigs.43526 | 21304680 | PMC3035259 | Gc00590 | 1 | 2010 |
| 51d55e7a067c014cd6f12f7c | Article | 10.1111/j.1574-6968.2012.02648.x | 22846039 | | CP000738 | 1 | 2012 |
| 51d55e7a067c014cd6f12f7c | Article | 10.3389/fpls.2020.560768 | 33519831 | PMC7840509 | CP000738 | 1 | 2021 |
| 51d55e7a067c014cd6f12f7c | Article | 10.1093/gbe/evx255 | 29220487 | PMC5739047 | CP000738 | 1 | 2017 |
| 51d55e7a067c014cd6f12f7c | Article | 10.1038/hdy.2014.32 | 24736785 | PMC4181065 | CP000738 | 1 | 2014 |
| 51d55e7a067c014cd6f12f7c | Article | 10.1099/ijs.0.008508-0 | 19567586 | | CP000738 | 1 | 2009 |
| 51d55e7a067c014cd6f12f7c | Article | 10.1099/ijsem.0.001510 | 27653171 | | CP000738 | 1 | 2016 |
| 51d55e7a067c014cd6f12f7c | Article | 10.1099/ijsem.0.001951 | 28771124 | | CP000738 | 1 | 2017 |
| 51d55e7a067c014cd6f12f7c | Article | 10.1099/ijsem.0.002332 | 28945538 | | CP000738 | 1 | 2017 |
| 51d55e7a067c014cd6f12f7c | Chapter | 10.1007/978-1-84800-255-5_6 | | | CP000738 | 1 | 2009 |
| 51d55e7a067c014cd6f12f7c | Article | 10.1016/j.jbiotec.2011.01.011 | 21329739 | | CP000738 | 1 | 2011 |
| 51d55e7a067c014cd6f12f7c | Article | 10.1007/s10295-016-1762-6 | 27021845 | | CP000738 | 1 | 2016 |
| 51d55e7a067c014cd6f12f7c | Article | 10.1007/s00248-010-9685-7 | 20521039 | | CP000738 | 1 | 2010 |
| 51d55e7a067c014cd6f12f7c | Article | 10.1590/s0100-204x2009000400008 | | | CP000738 | 1 | 2009 |
| 51d55e7a067c014cd6f12f7c | Article | 10.1016/j.resmic.2009.03.009 | 19403105 | | CP000738 | 1 | 2009 |

| | | | | | | | |
|---|---|---|---|---|---|---|---|
| 51d55e7a067c014cd6f12f7c | Article | 10.1134/s1022795418080045 | | | CP000738 | 1 | 2018 |
| 51d55e7a067c014cd6f12f7c | Article | 10.1016/j.syapm.2012.02.003 | 22429391 | | CP000738 | 1 | 2012 |
| 51d55e7a067c014cd6f12f7c | Article | 10.1016/j.tim.2009.07.004 | 19766492 | | CP000738 | 1 | 2009 |
| 51d55e7a067c014cd6f12f7c | Chapter | 10.1002/9780470015902.a0021157 | | | CP000738 | 1 | 2010 |
| 51d55e7a067c014cd6f12f7c | Chapter | 10.1007/978-3-642-21680-0_5 | | | CP000738 | 1 | 2011 |
| 51d55e7a067c014cd6f12f7c | Chapter | 10.1007/978-3-642-30194-0_73 | | | CP000738 | 1 | 2013 |
| 51d55e7a067c014cd6f12f7c | Article | 10.14201/gredos.83338 | | | CP000738 | 1 | 2010 |
| 51d68ef5067c014cd6012202 | Article | 10.18946/jssm.72.1_7 | | | CP000769 | 1 | 2018 |
| 51d68ef5067c014cd6012202 | Article | 10.18946/jssm.74.1_2 | | | CP000769 | 1 | 2020 |
| 51d68ef5067c014cd6012202 | Article | 10.1128/aem.01854-09 | 19897758 | PMC2798628 | CP000769 | 1 | 2009 |
| 51d68ef5067c014cd6012202 | Article | 10.1128/aem.02473-08 | 19346346 | PMC2687297 | CP000769 | 1 | 2009 |
| 51d68ef5067c014cd6012202 | Article | 10.1016/j.apsoil.2011.07.002 | | | CP000769 | 1 | 2011 |
| 51d68ef5067c014cd6012202 | Article | 10.1016/j.biocontrol.2011.02.012 | | | CP000769 | 1 | 2011 |
| 51d68ef5067c014cd6012202 | Article | 10.1111/j.1462-2920.2007.01434.x | 17803764 | PMC2040194 | CP000769 | 1 | 2007 |
| 51d68ef5067c014cd6012202 | Article | 10.1111/j.1758-2229.2009.00021.x | 23765745 | | CP000769 | 1 | 2009 |
| 51d68ef5067c014cd6012202 | Article | 10.1093/femsec/fiv030 | 25873605 | | CP000769 | 1 | 2015 |
| 51d68ef5067c014cd6012202 | Article | 10.1099/ijs.0.65806-0 | 18984709 | | CP000769 | 1 | 2008 |
| 51d68ef5067c014cd6012202 | Chapter | 10.1007/978-1-84800-255-5_6 | | | CP000769 | 1 | 2009 |
| 51d68ef5067c014cd6012202 | Article | 10.1371/journal.pone.0108877 | 25280065 | PMC4184826 | CP000769 | 1 | 2014 |
| 51d68ef5067c014cd6012202 | Article | 10.1016/j.soilbio.2010.11.005 | | | CP000769 | 1 | 2011 |
| 51d68ef5067c014cd6012202 | Chapter | 10.1002/9783527807796.ch12 | | | CP000769 | 1 | 2016 |
| 51d4a5f6067c014cd6ea22aa | Article | 10.4014/jmb.1604.04070 | 27381337 | | PRJNA51139 | 1 | 2016 |
| 59cad6347ded5e2f186918a7 | Article | 10.1038/ncomms12662 | 27601008 | PMC5023957 | PRJNA196014 | 1 | 2016 |
| 51d4c2ca067c014cd6ea9493 | Article | 10.1007/s12010-014-1079-8 | 25099371 | | CP002352 | 1 | 2014 |
| 51d4c2ca067c014cd6ea9493 | Article | 10.4056/sigs.1513795 | 21475586 | PMC3072090 | CP002352 | 1 | 2011 |
| 51d4c2ca067c014cd6ea9493 | Article | 10.1016/j.nmni.2016.06.002 | 27408745 | PMC4932483 | CP002352 | 1 | 2016 |
| 51d4c2ca067c014cd6ea9493 | Article | 10.1016/j.nmni.2017.11.009 | 29321938 | PMC5751998 | CP002352 | 1 | 2017 |
| 51d4c2ca067c014cd6ea9493 | Article | 10.1371/journal.pone.0206484 | 31509535 | PMC6738582 | CP002352 | 1 | 2019 |
| 523b41ce067c01393707f440 | Article | 10.1186/s40793-015-0128-4 | 26767090 | PMC4711178 | PRJNA163045 | 1 | 2016 |
| 51d4cec5067c014cd6eb3988 | Chapter | 10.1007/978-3-642-38954-2_384 | | | PRJNA33163 | 1 | 2014 |
| 51d4cec6067c014cd6eb3991 | Article | 10.4056/sigs.1273360 | 21304735 | PMC3035301 | CP002281 | 1 | 2010 |
| 51d4cec6067c014cd6eb3991 | Article | 10.1099/ijs.0.061242-0 | 24912824 | | CP002281 | 1 | 2014 |
| 51d4cec6067c014cd6eb3991 | Article | 10.1016/j.nmni.2016.12.003 | 28116104 | PMC5225283 | CP002281 | 1 | 2016 |
| 51d4cec6067c014cd6eb3991 | Chapter | 10.1007/978-3-642-30120-9_213 | | | CP002281 | 1 | 2014 |
| 51d4ccae067c014cd6eb1d56 | Chapter | 10.1007/978-3-642-38954-2_384 | | | PRJNA20741 | 1 | 2014 |
| 51d4caba067c014cd6eb0239 | Article | 10.1128/aem.00810-12 | 22660709 | PMC3406152 | CP002361 | 1 | 2012 |
| 51d4caba067c014cd6eb0239 | Article | 10.4056/sigs.1563919 | 21475591 | PMC3072082 | CP002361 | 1 | 2011 |
| 51d4caba067c014cd6eb0239 | Article | 10.4056/sigs.1734292 | 21677858 | PMC3111992 | CP002361 | 1 | 2011 |
| 51d4caba067c014cd6eb0239 | Article | 10.1007/s00792-018-1055-2 | 30219948 | | CP002361 | 1 | 2018 |
| 51d4caba067c014cd6eb0239 | Article | 10.1007/s00792-019-01131-6 | 31535211 | | CP002361 | 1 | 2019 |
| 51d4caba067c014cd6eb0239 | Article | 10.3389/fmicb.2019.02083 | 31608019 | PMC6767994 | SAMN00138957 | 1 | 2019 |
| 51d4caba067c014cd6eb0239 | Chapter | 10.1002/9781118960608.gbm00476.pub2 | | | CP002361 | 1 | 2018 |
| 51d4caba067c014cd6eb0239 | Chapter | 10.1002/9781118960608.gbm00477.pub2 | | | CP002361 | 1 | 2018 |
| 51d4caba067c014cd6eb0239 | Chapter | 10.1002/9781118960608.gbm01328 | | | CP002361 | 1 | 2018 |
| 51d493b9067c014cd6ea1912 | Article | 10.1128/aem.00611-16 | 27235430 | PMC4984305 | CP002364 | 1 | 2016 |
| 51d493b9067c014cd6ea1912 | Article | 10.1128/aem.01064-15 | 26116678 | PMC4551263 | CP002364 | 1 | 2015 |
| 51d493b9067c014cd6ea1912 | Article | 10.11233/aquaculturesci.60.519 | | | CP002364 | 1 | 2012 |
| 51d493b9067c014cd6ea1912 | Article | 10.1016/j.bej.2014.05.023 | | | CP002364 | 1 | 2014 |
| 51d493b9067c014cd6ea1912 | Article | 10.1111/1462-2920.12388 | 24428801 | PMC4262008 | CP002364 | 1 | 2014 |
| 51d493b9067c014cd6ea1912 | Article | 10.1111/1462-2920.14343 | 29968357 | | CP002364 | 1 | 2018 |
| 51d493b9067c014cd6ea1912 | Article | 10.1111/1758-2229.12136 | 25646534 | | CP002364 | 1 | 2014 |
| 51d493b9067c014cd6ea1912 | Article | 10.4056/sigs.1613929 | 21475592 | PMC3072085 | CP002364 | 1 | 2011 |
| 51d493b9067c014cd6ea1912 | Article | 10.3389/fmicb.2017.00152 | 28217124 | PMC5289976 | CP002364 | 1 | 2017 |
| 51d493b9067c014cd6ea1912 | Article | 10.1099/ijsem.0.000828 | 26646853 | | CP002364 | 1 | 2015 |
| 51d493b9067c014cd6ea1912 | Article | 10.1099/ijsem.0.001083 | 27082267 | | CP002364 | 1 | 2016 |
| 51d493b9067c014cd6ea1912 | Article | 10.1007/s12275-012-1342-z | 22538648 | | CP002364 | 1 | 2012 |
| 51d493b9067c014cd6ea1912 | Article | 10.1134/s002626171404016x | | | CP002364 | 1 | 2014 |
| 51d493b9067c014cd6ea1912 | Article | 10.1038/s41564-019-0432-7 | 31036911 | PMC6697534 | CP002364 | 1 | 2019 |
| 51d493b9067c014cd6ea1912 | Chapter | 10.1002/9781118960608.fbm00194.pub2 | | | CP002364 | 1 | 2020 |
| 51d493b9067c014cd6ea1912 | Chapter | 10.1002/9781118960608.gbm01023.pub2 | | | CP002364 | 1 | 2019 |
| 51d4c31b067c014cd6ea9999 | Article | 10.1637/9770-050411-reg.1 | 22856189 | | CP002346 | 1 | 2012 |
| 51d4c31b067c014cd6ea9999 | Article | 10.1080/03079457.2012.752066 | 23391177 | | CP002346 | 1 | 2012 |
| 51d4c31b067c014cd6ea9999 | Article | 10.1007/s00284-021-02465-1 | 33813642 | | CP002346 | 1 | 2021 |

| | | | | | | | |
|---|---|---|---|---|---|---|---|
| 51d4c31b067c014cd6ea9999 | Article | 10.1016/j.diagmicrobio.2014.12.001 | 25544000 | | CP002346 | 1 | 2014 |
| 51d4c31b067c014cd6ea9999 | Article | 10.4056/sigs.1553862 | 21677851 | PMC3111989 | CP002346 | 1 | 2011 |
| 51d4c31b067c014cd6ea9999 | Article | 10.4056/sigs.1553865 | | | CP002346 | 1 | 2011 |
| 51d4c31b067c014cd6ea9999 | Article | 10.1099/ijs.0.052779-0 | 24271213 | | CP002346 | 1 | 2013 |
| 51d4c31b067c014cd6ea9999 | Article | 10.1099/ijs.0.054585-0 | 24014625 | | CP002346 | 1 | 2013 |
| 51d4c31b067c014cd6ea9999 | Article | 10.1099/ijs.0.060178-0 | 24453231 | | CP002346 | 1 | 2014 |
| 51d4c31b067c014cd6ea9999 | Article | 10.1099/ijs.0.063115-0 | 24844262 | | CP002346 | 1 | 2014 |
| 51d4c31b067c014cd6ea9999 | Article | 10.1099/ijsem.0.000757 | 26554606 | | CP002346 | 1 | 2015 |
| 51d4c31b067c014cd6ea9999 | Article | 10.1099/ijsem.0.001841 | 28150572 | | CP002346 | 1 | 2017 |
| 51d4c31b067c014cd6ea9999 | Article | 10.1099/mic.0.000123 | 26293113 | | CP002346 | 1 | 2015 |
| 51d4c31b067c014cd6ea9999 | Article | 10.1371/journal.pone.0039805 | 22768127 | PMC3387259 | CP002346 | 1 | 2012 |
| 51d4c31b067c014cd6ea9999 | Article | 10.1371/journal.pone.0131078 | 26107936 | PMC4481100 | CP002346 | 1 | 2015 |
| 51d4c31b067c014cd6ea9999 | Article | 10.1016/j.vetmic.2011.11.002 | 22112855 | | CP002346 | 1 | 2011 |
| 51d4c31b067c014cd6ea9999 | Article | 10.1016/j.vetmic.2012.01.009 | 22317978 | | CP002346 | 1 | 2012 |
| 51d4c31b067c014cd6ea9999 | Article | 10.1016/j.vetmic.2013.11.027 | 24345412 | | CP002346 | 1 | 2013 |
| 51d4c31b067c014cd6ea9999 | Article | 10.1016/j.vetmic.2015.03.003 | 25804836 | | CP002346 | 1 | 2015 |
| 51d4c31b067c014cd6ea9999 | Article | 10.1016/j.vetmic.2016.04.014 | 27259827 | | CP002346 | 1 | 2016 |
| 51d4c31b067c014cd6ea9999 | Article | 10.1186/s13567-021-00900-6 | 33579370 | PMC7881567 | CP002346 | 1 | 2021 |
| 51d4ca46067c014cd6eafc58 | Article | 10.4028/www.scientific.net/amr.641-642.684 | | | CP002534 | 1 | 2013 |
| 51d4ca46067c014cd6eafc58 | Article | 10.1007/s10482-014-0196-2 | 24906660 | | CP002534 | 1 | 2014 |
| 51d4ca46067c014cd6eafc58 | Article | 10.1128/aem.07339-11 | 22267664 | PMC3302594 | CP002534 | 1 | 2012 |
| 51d4ca46067c014cd6eafc58 | Article | 10.1007/s00284-018-1496-y | 29761217 | | CP002534 | 1 | 2018 |
| 51d4ca46067c014cd6eafc58 | Article | 10.4056/sigs.1774329 | 21677859 | PMC3111997 | CP002534 | 1 | 2011 |
| 51d4ca46067c014cd6eafc58 | Article | 10.1099/ijs.0.000138 | 25713047 | | CP002534 | 1 | 2015 |
| 51d4ca46067c014cd6eafc58 | Article | 10.1099/ijs.0.030254-0 | 22753526 | | CP002534 | 1 | 2012 |
| 51d4ca46067c014cd6eafc58 | Article | 10.1099/ijs.0.041509-0 | 22771683 | | CP002534 | 1 | 2012 |
| 51d4ca46067c014cd6eafc58 | Article | 10.1099/ijs.0.041889-0 | 22707529 | | CP002534 | 1 | 2012 |
| 51d4ca46067c014cd6eafc58 | Article | 10.1099/ijs.0.044305-0 | 22843717 | | CP002534 | 1 | 2012 |
| 51d4ca46067c014cd6eafc58 | Article | 10.1099/ijs.0.048413-0 | 23416571 | | CP002534 | 1 | 2013 |
| 51d4ca46067c014cd6eafc58 | Article | 10.1099/ijs.0.054627-0 | 24126635 | | CP002534 | 1 | 2013 |
| 51d4ca46067c014cd6eafc58 | Article | 10.1099/ijs.0.055517-0 | 24030690 | | CP002534 | 1 | 2013 |
| 51d4ca46067c014cd6eafc58 | Article | 10.1099/ijsem.0.000757 | 26554606 | | CP002534 | 1 | 2015 |
| 51d4ca46067c014cd6eafc58 | Article | 10.1099/ijsem.0.002095 | 28829014 | | CP002534 | 1 | 2017 |
| 51d4ca46067c014cd6eafc58 | Article | 10.1099/ijsem.0.002298 | 28857039 | | CP002534 | 1 | 2017 |
| 51d4ca46067c014cd6eafc58 | Article | 10.1099/ijsem.0.002340 | 28945544 | | CP002534 | 1 | 2017 |
| 51d4ca46067c014cd6eafc58 | Article | 10.1099/ijsem.0.002489 | 29205128 | | CP002534 | 1 | 2017 |
| 51d4ca46067c014cd6eafc58 | Article | 10.1099/ijsem.0.002576 | 29458460 | | CP002534 | 1 | 2018 |
| 51d4ca46067c014cd6eafc58 | Article | 10.1099/ijsem.0.002607 | 29458481 | | CP002534 | 1 | 2018 |
| 51d4ca46067c014cd6eafc58 | Article | 10.1099/ijsem.0.002730 | 29570445 | | CP002534 | 1 | 2018 |
| 51d4ca46067c014cd6eafc58 | Article | 10.1099/ijsem.0.002808 | 29781798 | | CP002534 | 1 | 2018 |
| 51d4ca46067c014cd6eafc58 | Article | 10.1128/jb.05241-11 | 21622754 | PMC3147531 | CP002534 | 1 | 2011 |
| 51d4ca46067c014cd6eafc58 | Article | 10.1371/journal.pone.0146307 | 26745366 | PMC4720170 | CP002534 | 1 | 2016 |
| 51d4ca46067c014cd6eafc58 | Article | 10.1016/j.syapm.2013.02.004 | 23623798 | | CP002534 | 1 | 2013 |
| 51d4ca46067c014cd6eafc58 | Article | 10.1038/ismej.2013.156 | 24048225 | PMC3906817 | CP002534 | 1 | 2013 |
| 51d4ca46067c014cd6eafc58 | Chapter | 10.1002/9781118960608.gbm00300.pub2 | | | CP002534 | 1 | 2016 |
| 548659000d87850ddcd6cdb1 | Article | 10.1186/s40793-017-0283-x | 29225730 | PMC5717998 | PRJNA255603 | 1 | 2017 |
| 55e07f090d878556782d9b3f | Article | 10.1038/ncomms10613 | 26837824 | PMC4742961 | SAMN03203005 | 1 | 2016 |
| 51d492be067c014cd6ea1224 | Article | 10.1007/s00203-018-1608-x | 30519708 | PMC6514085 | NC_009042 | 1 | 2018 |
| 51d492be067c014cd6ea1224 | Article | 10.1016/j.sjbs.2020.12.039 | 33732074 | PMC7938122 | NC_009042 | 1 | 2020 |
| 51d492be067c014cd6ea1224 | Chapter | 10.1201/9781420076813.ch7 | | | NC_009042 | 1 | 2009 |
| 51d4cebc067c014cd6eb3913 | Article | 10.3389/fmicb.2018.02007 | 30186281 | PMC6113628 | SAMN00713569 | 1 | 2018 |
| 51d4c72f067c014cd6ead843 | Article | 10.1155/2017/7039245 | 28497061 | PMC5405348 | CP002175 | 1 | 2017 |
| 51d4c72f067c014cd6ead843 | Article | 10.4056/sigs.1824509 | 21886858 | PMC3156398 | CP002175 | 1 | 2011 |
| 51d4c72f067c014cd6ead843 | Article | 10.1007/s00792-012-0455-y | 22527048 | | CP002175 | 1 | 2012 |
| 51d4c72f067c014cd6ead843 | Article | 10.1007/s00792-012-0476-6 | 22907126 | | CP002175 | 1 | 2012 |
| 51d4c72f067c014cd6ead843 | Chapter | 10.1002/9781118960608.gbm01725 | | | CP002175 | 1 | 2019 |
| 51d4c72f067c014cd6ead843 | Chapter | 10.1007/978-3-642-30120-9_218 | | | CP002175 | 1 | 2014 |
| 51d4a62a067c014cd6ea2340 | Article | 10.1007/s00253-016-7294-1 | 26762388 | | CP002416 | 1 | 2016 |
| 51d4a62a067c014cd6ea2340 | Article | 10.1007/s00253-017-8438-7 | 28779289 | | CP002416 | 1 | 2017 |
| 51d4a62a067c014cd6ea2340 | Article | 10.1016/j.biortech.2012.11.048 | 23262013 | | CP002416 | 1 | 2012 |
| 51d4a62a067c014cd6ea2340 | Article | 10.1186/1754-6834-6-31 | 23448304 | PMC3598825 | CP002416 | 1 | 2013 |
| 51d4a62a067c014cd6ea2340 | Article | 10.1186/s13068-016-0684-x | 28053665 | PMC5209896 | CP002416 | 1 | 2017 |
| 51d4a62a067c014cd6ea2340 | Article | 10.1186/s13068-018-1245-2 | 30202437 | PMC6125887 | CP002416 | 1 | 2018 |

| | | | | | | | |
|---|---|---|---|---|---|---|---|
| 51d4a62a067c014cd6ea2340 | Article | 10.1186/s13068-019-1524-6 | 31367231 | PMC6652007 | CP002416 | 1 | 2019 |
| 51d4a62a067c014cd6ea2340 | Article | 10.4056/sigs.2044675 | | | CP002416 | 1 | 2011 |
| 51d4a62a067c014cd6ea2340 | Article | 10.1111/jam.12112 | 23279216 | | CP002416 | 1 | 2013 |
| 51d4a62a067c014cd6ea2340 | Article | 10.1128/jb.00232-15 | 26013492 | PMC4518838 | CP002416 | 1 | 2015 |
| 51d4a62a067c014cd6ea2340 | Article | 10.1128/jb.00322-11 | 21460082 | PMC3133140 | CP002416 | 1 | 2011 |
| 51d4a62a067c014cd6ea2340 | Article | 10.1016/j.jhazmat.2011.10.087 | 22137177 | | CP002416 | 1 | 2011 |
| 51d4a62a067c014cd6ea2340 | Article | 10.1007/s10295-018-2073-x | 30187243 | | CP002416 | 1 | 2018 |
| 51d4a62a067c014cd6ea2340 | Article | 10.1016/j.ymben.2017.04.002 | 28400329 | | CP002416 | 1 | 2017 |
| 51d4a62a067c014cd6ea2340 | Article | 10.1016/j.ymben.2017.06.011 | 28663138 | | CP002416 | 1 | 2017 |
| 51d4a62a067c014cd6ea2340 | Article | 10.1016/j.mec.2019.e00116 | 31890588 | PMC6926293 | CP002416 | 1 | 2019 |
| 51d4a62a067c014cd6ea2340 | Article | 10.1016/j.meteno.2016.04.001 | 29142822 | PMC5678826 | CP002416 | 1 | 2016 |
| 51d4a62a067c014cd6ea2340 | Article | 10.1038/nrmicro2729 | 22266780 | | CP002416 | 1 | 2012 |
| 51d4a62a067c014cd6ea2340 | Article | 10.1016/j.renene.2013.06.047 | | | CP002416 | 1 | 2014 |
| 51d4a62a067c014cd6ea2340 | Article | 10.1038/s41598-018-37979-5 | 30741948 | PMC6370804 | CP002416 | 1 | 2019 |
| 51d4a62a067c014cd6ea2340 | Article | 10.1016/j.syapm.2014.10.002 | 25467556 | | CP002416 | 1 | 2014 |
| 51d4a62a067c014cd6ea2340 | Article | 10.1016/j.syapm.2020.126154 | 33227632 | | CP002416 | 1 | 2020 |
| 523b4d61067c01393707fddd | Article | 10.4056/sigs.5449586 | 25197480 | PMC4148996 | Gi11554 | 1 | 2014 |
| 52f3e4c0067c011a2113fd1c | Article | 10.1371/journal.pone.0183007 | 28832647 | PMC5568408 | Gi11069 | 0 | 2017 |
| 523b429a067c01393707f602 | Article | 10.1186/s40793-015-0036-7 | 26380638 | PMC4572677 | Gp0013740 | 1 | 2015 |
| 540648ec0d878557fd3be7ef | Article | 10.3389/fmicb.2018.02007 | 30186281 | PMC6113628 | SAMN02440801 | 1 | 2018 |
| 523b41da067c01393707f45c | Article | 10.1186/s40793-016-0191-5 | 27721915 | PMC5052931 | Gp0013295 | 1 | 2016 |
| 52e04cc1067c017f29056847 | Article | 10.1186/s12864-017-3955-4 | 28764658 | PMC5540593 | JHWM00000000 | 1 | 2017 |
| 52e04cc1067c017f29056847 | Article | 10.1099/ijsem.0.002243 | 28893355 | | JHWM00000000 | 1 | 2017 |
| 52e04cc1067c017f29056847 | Article | 10.1099/ijsem.0.003306 | 30789326 | | JHWM00000000 | 1 | 2019 |
| 530d650849607a1be0054c26 | Article | 10.3389/fmicb.2018.02007 | 30186281 | PMC6113628 | SAMN02745916 | 1 | 2018 |
| 58ea58357ded5e52c4b0447d | Article | 10.1186/s40168-018-0541-1 | 30219103 | PMC6138922 | Gs0110119 | 1 | 2018 |
| 57d6ee757ded5e3135ba4f33 | Article | 10.3389/fmicb.2019.02083 | 31608019 | PMC6767994 | SAMN05443667 | 1 | 2019 |
| 51d4cb53067c014cd6eb0adf | Article | 10.1099/ijsem.0.001888 | 28742009 | | CP002637 | 1 | 2017 |
| 51d4cb53067c014cd6eb0adf | Article | 10.1002/mbo3.50 | 23239474 | PMC3584212 | CP002637 | 1 | 2012 |
| 51d4cb53067c014cd6eb0adf | Article | 10.1371/journal.pone.0060120 | 23577086 | PMC3618517 | CP002637 | 1 | 2013 |
| 51d4cb53067c014cd6eb0adf | Article | 10.1371/journal.pone.0069076 | 23950883 | PMC3741291 | CP002637 | 1 | 2013 |
| 58ccefca7ded5e7a2afbbe64 | Article | 10.1186/s40168-018-0541-1 | 30219103 | PMC6138922 | Gs0110119 | 1 | 2018 |
| 51d53736067c014cd6efefa1 | Article | 10.3389/fmicb.2019.02083 | 31608019 | PMC6767994 | SAMN02440754 | 1 | 2019 |
| 59122b2c7ded5e1e49ffdd4e | Article | 10.1186/s40168-018-0541-1 | 30219103 | PMC6138922 | Gs0110119 | 1 | 2018 |
| 53051c9349607a64d3cac728 | Article | 10.1186/s40793-016-0168-4 | 27471578 | PMC4964001 | Gp0006506 | 1 | 2016 |
| 51d53185067c014cd6efa827 | Article | 10.1016/j.nmni.2017.11.003 | 29348922 | PMC5767839 | AUJC00000000 | 1 | 2017 |
| 5b35710464d0b3516a5eb4ad | Article | 10.3389/fmicb.2018.02007 | 30186281 | PMC6113628 | SAMN04489716 | 1 | 2018 |
| 51edff15067c014b2c663b35 | Article | 10.1016/j.anaerobe.2014.06.007 | 24969840 | | AXVL00000000 | 1 | 2014 |
| 51edff15067c014b2c663b35 | Article | 10.1080/20002297.2017.1368848 | 29081911 | PMC5646626 | AXVL00000000 | 1 | 2017 |
| 52901e3b067c013e2b060c9d | Article | 10.1038/s42003-019-0365-y | 30993215 | PMC6465285 | NZ_KB899704 | 1 | 2019 |
| 52901e3b067c013e2b060c9d | Article | 10.3389/fmicb.2019.02083 | 31608019 | PMC6767994 | SAMN02440995 | 1 | 2019 |
| 58010ebd7ded5e3135bc4d29 | Article | 10.1007/s00284-021-02467-z | 33797566 | | LT670845 | 1 | 2021 |
| 58010ebd7ded5e3135bc4d29 | Article | 10.1264/jsme2.me17188 | 29709896 | PMC6031400 | LT670845 | 1 | 2018 |
| 52901805067c013e2b060034 | Article | 10.1016/j.actatropica.2017.05.018 | 28535905 | | CP003341 | 1 | 2017 |
| 52901805067c013e2b060034 | Article | 10.1128/aem.03463-16 | 28213544 | PMC5394329 | CP003341 | 1 | 2017 |
| 52901805067c013e2b060034 | Article | 10.1128/cmr.00032-13 | 24092850 | PMC3811236 | CP003341 | 1 | 2013 |
| 52901805067c013e2b060034 | Article | 10.3201/eid2010.140175 | 25271771 | PMC4193168 | CP003341 | 1 | 2014 |
| 52901805067c013e2b060034 | Article | 10.1007/s10493-019-00343-x | 30771038 | PMC6402582 | CP003341 | 1 | 2019 |
| 52901805067c013e2b060034 | Article | 10.1007/s10493-020-00524-z | 33025238 | | CP003341 | 1 | 2020 |
| 52901805067c013e2b060034 | Article | 10.1007/s10493-020-00524-z | 33025238 | | NC_017044 | 1 | 2020 |
| 52901805067c013e2b060034 | Article | 10.1016/j.meegid.2014.03.014 | 24662440 | | NC_017044 | 1 | 2014 |
| 52901805067c013e2b060034 | Article | 10.1016/j.meegid.2014.05.028 | 24924907 | | NC_017044 | 1 | 2014 |
| 52901805067c013e2b060034 | Article | 10.1099/ijsem.0.003057 | 30307387 | | NC_017044 | 1 | 2018 |
| 52901805067c013e2b060034 | Article | 10.1080/07391102.2021.1898473 | 33719856 | | NC_017044 | 1 | 2021 |
| 52901805067c013e2b060034 | Article | 10.1128/jcm.00457-15 | 26135877 | PMC4540899 | CP003341 | 1 | 2015 |
| 52901805067c013e2b060034 | Article | 10.18474/jes16-12.1 | | | CP003341 | 1 | 2016 |
| 52901805067c013e2b060034 | Article | 10.1093/jme/tjaa059 | 32249319 | | CP003341 | 1 | 2020 |
| 52901805067c013e2b060034 | Article | 10.1093/jme/tjaa130 | 32647878 | PMC7780643 | CP003341 | 1 | 2020 |
| 52901805067c013e2b060034 | Article | 10.1093/jme/tjv251 | 26819330 | | CP003341 | 1 | 2016 |
| 52901805067c013e2b060034 | Article | 10.1093/jme/tjw121 | 27480099 | | CP003341 | 1 | 2016 |
| 52901805067c013e2b060034 | Article | 10.1093/jme/tjx138 | 28981813 | | CP003341 | 1 | 2017 |
| 52901805067c013e2b060034 | Article | 10.1093/jme/tjx141 | 28981693 | | CP003341 | 1 | 2017 |
| 52901805067c013e2b060034 | Article | 10.1093/jme/tjx194 | 29045695 | | CP003341 | 1 | 2017 |

| | | | | | | | |
|---|---|---|---|---|---|---|---|
| 52901805067c013e2b060034 | Article | 10.1093/jme/tjy169 | 30312440 | | NC_017044 | 1 | 2018 |
| 52901805067c013e2b060034 | Article | 10.1292/jvms.12-0143 | 22986299 | | CP003341 | 1 | 2012 |
| 52901805067c013e2b060034 | Article | 10.1111/mve.12315 | 29972600 | | CP003341 | 1 | 2018 |
| 52901805067c013e2b060034 | Article | 10.1111/mve.12480 | 32936461 | | CP003341 | 1 | 2020 |
| 52901805067c013e2b060034 | Article | 10.1016/j.micinf.2015.09.012 | 26423020 | | CP003341 | 1 | 2015 |
| 52901805067c013e2b060034 | Article | 10.1016/j.micinf.2015.09.012 | 26423020 | | NC_017044 | 1 | 2015 |
| 52901805067c013e2b060034 | Article | 10.1016/j.micinf.2017.11.009 | 29287988 | | NC_017044 | 1 | 2017 |
| 52901805067c013e2b060034 | Article | 10.1002/mbo3.527 | 29047217 | PMC5822307 | NC_017044 | 1 | 2017 |
| 52901805067c013e2b060034 | Article | 10.3103/s0891416818020088 | | | CP003341 | 1 | 2018 |
| 52901805067c013e2b060034 | Article | 10.1016/j.nmni.2018.02.012 | 29692912 | PMC5913362 | NC_017044 | 1 | 2018 |
| 52901805067c013e2b060034 | Article | 10.1186/1756-3305-7-318 | 25011617 | PMC4230250 | CP003341 | 1 | 2014 |
| 52901805067c013e2b060034 | Article | 10.1186/s13071-015-1242-2 | 26652857 | PMC4675064 | CP003341 | 1 | 2015 |
| 52901805067c013e2b060034 | Article | 10.1007/s00436-016-5091-5 | 27130322 | | CP003341 | 1 | 2016 |
| 52901805067c013e2b060034 | Article | 10.1007/s00436-021-07128-5 | 33830363 | | CP003341 | 1 | 2021 |
| 52901805067c013e2b060034 | Article | 10.1371/journal.pntd.0007734 | 31490924 | PMC6750615 | CP003341 | 1 | 2019 |
| 52901805067c013e2b060034 | Article | 10.1371/journal.pone.0037310 | 22624012 | PMC3356282 | CP003341 | 1 | 2012 |
| 52901805067c013e2b060034 | Article | 10.1371/journal.pone.0071861 | 24039725 | PMC3767692 | CP003341 | 1 | 2013 |
| 52901805067c013e2b060034 | Article | 10.1371/journal.pone.0147492 | 26866478 | PMC4750851 | CP003341 | 1 | 2016 |
| 52901805067c013e2b060034 | Article | 10.1371/journal.pone.0197012 | 29723287 | PMC5933787 | CP003341 | 1 | 2018 |
| 52901805067c013e2b060034 | Article | 10.11158/saa.26.2.4 | | | CP003341 | 1 | 2021 |
| 52901805067c013e2b060034 | Article | 10.1016/j.ttbdis.2013.07.001 | 24331642 | | CP003341 | 1 | 2013 |
| 52901805067c013e2b060034 | Article | 10.1016/j.ttbdis.2014.07.010 | 25108786 | | CP003341 | 1 | 2014 |
| 52901805067c013e2b060034 | Article | 10.1016/j.ttbdis.2015.02.011 | 25800099 | | CP003341 | 1 | 2015 |
| 52901805067c013e2b060034 | Article | 10.1016/j.ttbdis.2015.04.011 | 25958197 | | CP003341 | 1 | 2015 |
| 52901805067c013e2b060034 | Article | 10.1016/j.ttbdis.2016.08.006 | 27554852 | | CP003341 | 1 | 2016 |
| 52901805067c013e2b060034 | Article | 10.1016/j.ttbdis.2017.11.002 | 29174365 | | CP003341 | 1 | 2017 |
| 52901805067c013e2b060034 | Article | 10.1016/j.ttbdis.2018.02.015 | 29483057 | | CP003341 | 1 | 2018 |
| 52901805067c013e2b060034 | Article | 10.1016/j.ttbdis.2018.03.022 | 29636236 | | CP003341 | 1 | 2018 |
| 52901805067c013e2b060034 | Article | 10.1016/j.ttbdis.2018.12.010 | 30611725 | | CP003341 | 1 | 2018 |
| 52901805067c013e2b060034 | Article | 10.1016/j.ttbdis.2019.05.013 | 31176663 | | CP003341 | 1 | 2019 |
| 52901805067c013e2b060034 | Article | 10.1016/j.ttbdis.2020.101639 | 33360385 | | CP003341 | 1 | 2020 |
| 52901805067c013e2b060034 | Article | 10.1016/j.ttbdis.2021.101672 | 33561680 | | CP003341 | 1 | 2021 |
| 52901805067c013e2b060034 | Article | 10.1111/tbed.13525 | 32090496 | | CP003341 | 1 | 2020 |
| 52901805067c013e2b060034 | Article | 10.1111/tbed.14027 | 33667026 | | CP003341 | 1 | 2021 |
| 52901805067c013e2b060034 | Article | 10.1016/j.tmaid.2018.10.007 | 30312734 | | CP003341 | 1 | 2018 |
| 52901805067c013e2b060034 | Article | 10.1016/j.vprsr.2020.100448 | 33308714 | | CP003341 | 1 | 2020 |
| 52901805067c013e2b060034 | Article | 10.1111/zph.12749 | 32697888 | PMC7496946 | CP003341 | 1 | 2020 |
| 52901805067c013e2b060034 | Chapter | 10.1017/cbo9781139794749.012 | | | CP003341 | 1 | 2015 |
| 52901805067c013e2b060034 | Article | 10.11606/t.10.2016.tde-23032016-115746 | | | CP003341 | 1 | 2016 |
| 52901805067c013e2b060034 | Chapter | 10.1007/978-3-319-46859-4_20 | | | NC_017044 | 1 | 2016 |
| 530d64d049607a1be0054be2 | Article | 10.1111/jam.12784 | 25727916 | | CP007055 | 1 | 2015 |
| 530d64d049607a1be0054be2 | Article | 10.1016/j.mimet.2015.06.012 | 26122310 | | CP007055 | 1 | 2015 |
| 530d64d049607a1be0054be2 | Article | 10.3390/life5010949 | 25789552 | PMC4390887 | CP007055 | 1 | 2015 |
| 530d64d049607a1be0054be2 | Article | 10.1111/mmi.13204 | 26331239 | | CP007055 | 1 | 2015 |
| 530d64d049607a1be0054be2 | Chapter | 10.1002/9781118960608.gbm01530 | | | CP007055 | 1 | 2018 |
| 530d649f49607a1be0054bac | Article | 10.1007/s10482-015-0396-4 | 25636945 | | CP007034 | 1 | 2015 |
| 530d649f49607a1be0054bac | Article | 10.1007/s10482-021-01521-x | 33566238 | | CP007034 | 1 | 2021 |
| 530d649f49607a1be0054bac | Article | 10.3390/proceedings2110650 | | | CP007034 | 1 | 2018 |
| 530d649f49607a1be0054bac | Article | 10.1099/ijsem.0.000617 | 26377180 | | CP007034 | 1 | 2015 |
| 530d649f49607a1be0054bac | Article | 10.1371/journal.pone.0206484 | 31509535 | PMC6738582 | CP007034 | 1 | 2019 |
| 594d609a7ded5e4e5bbd6ade | Article | 10.1186/s40793-016-0165-7 | 27340512 | PMC4918011 | CP001825 | 1 | 2016 |
| 594d609a7ded5e4e5bbd6ade | Article | 10.4056/sigs.1153107 | 21304745 | PMC3035366 | CP001825 | 1 | 2010 |
| 594d609a7ded5e4e5bbd6ade | Article | 10.4056/sigs.2806097 | 22768368 | PMC3387798 | CP001825 | 1 | 2012 |
| 594d609a7ded5e4e5bbd6ade | Article | 10.4056/sigs.1153107 | 21304745 | PMC3035366 | Gc01150 | 1 | 2010 |
| 594d609a7ded5e4e5bbd6ade | Article | 10.1099/ijs.0.046532-0 | 23378114 | | CP001825 | 1 | 2013 |
| 594d609a7ded5e4e5bbd6ade | Article | 10.1099/ijs.0.060327-0 | 24814334 | | CP001825 | 1 | 2014 |
| 594d609a7ded5e4e5bbd6ade | Article | 10.1099/ijsem.0.002141 | 28741992 | | CP001825 | 1 | 2017 |
| 594d609a7ded5e4e5bbd6ade | Chapter | 10.1002/9781118960608.gbm01563 | | | CP001825 | 1 | 2018 |
| 530d68b749607a1be0054ee5 | Article | 10.1186/s40793-016-0165-7 | 27340512 | PMC4918011 | JAGA00000000 | 1 | 2016 |
| 530d68b749607a1be0054ee5 | Article | 10.3389/fmicb.2018.02007 | 30186281 | PMC6113628 | SAMN02584935 | 1 | 2018 |
| 55791c510d878529e7cb084b | Article | 10.3389/fmicb.2018.02453 | 30364313 | PMC6193092 | LIPW00000000 | 1 | 2018 |
| 55791c510d878529e7cb084b | Article | 10.1128/genomea.00831-16 | 27516518 | PMC4982297 | LIPW00000000 | 1 | 2016 |
| 5513329c0d878525404e829e | Article | 10.1016/j.cub.2015.01.014 | 25702576 | | CP010426 | 1 | 2015 |

| | | | | | | | |
|---|---|---|---|---|---|---|---|
| 5513329c0d878525404e829e | Article | 10.1186/s40168-018-0488-2 | 29884244 | PMC5994134 | CP010426 | 1 | 2018 |
| 5513329c0d878525404e829e | Article | 10.1038/s41579-018-0076-2 | 30181663 | | CP010426 | 1 | 2018 |
| 530d63a049607a1be0054a72 | Article | 10.1007/s10482-020-01400-x | 32193664 | | CP002279 | 1 | 2020 |
| 530d63a049607a1be0054a72 | Article | 10.1007/s00253-015-6515-3 | 25776061 | | Gc01853 | 1 | 2015 |
| 530d63a049607a1be0054a72 | Article | 10.1007/s00203-021-02189-7 | 33611634 | | CP002279 | 1 | 2021 |
| 530d63a049607a1be0054a72 | Article | 10.4056/sigs.3486982 | 24019991 | PMC3764936 | CP002279 | 1 | 2013 |
| 530d63a049607a1be0054a72 | Article | 10.4056/sigs.4538264 | 24976886 | PMC4062634 | CP002279 | 1 | 2013 |
| 530d63a049607a1be0054a72 | Article | 10.1186/1944-3277-9-6 | 25780499 | PMC4334631 | Gc01853 | 1 | 2014 |
| 530d63a049607a1be0054a72 | Article | 10.1186/1944-3277-9-7 | 25780500 | PMC4334872 | Gc01853 | 1 | 2014 |
| 530d63a049607a1be0054a72 | Article | 10.4056/sigs.4538264 | 24976886 | PMC4062634 | Gc01853 | 1 | 2013 |
| 530d63a049607a1be0054a72 | Article | 10.1007/s11356-017-9319-4 | 28593540 | | CP002279 | 1 | 2017 |
| 530d63a049607a1be0054a72 | Article | 10.1111/1574-6941.12411 | 25112496 | | CP002279 | 1 | 2014 |
| 530d63a049607a1be0054a72 | Article | 10.1111/j.1574-6941.2012.01476.x | 22928867 | | CP002279 | 1 | 2012 |
| 530d63a049607a1be0054a72 | Article | 10.1111/j.1574-6968.2012.02648.x | 22846039 | | CP002279 | 1 | 2012 |
| 530d63a049607a1be0054a72 | Article | 10.1093/gbe/evx255 | 29220487 | PMC5739047 | CP002279 | 1 | 2017 |
| 530d63a049607a1be0054a72 | Article | 10.1099/ijs.0.000164 | 25736411 | | CP002279 | 1 | 2015 |
| 530d63a049607a1be0054a72 | Article | 10.1099/ijsem.0.000430 | 26296780 | | CP002279 | 1 | 2015 |
| 530d63a049607a1be0054a72 | Article | 10.1099/ijsem.0.000796 | 26610329 | | CP002279 | 1 | 2015 |
| 530d63a049607a1be0054a72 | Article | 10.1099/ijsem.0.001448 | 27565417 | | CP002279 | 1 | 2016 |
| 530d63a049607a1be0054a72 | Article | 10.1099/ijsem.0.002770 | 29676730 | | CP002279 | 1 | 2018 |
| 530d63a049607a1be0054a72 | Article | 10.1099/ijsem.0.002924 | 30063199 | | NC_015675 | 1 | 2018 |
| 530d63a049607a1be0054a72 | Article | 10.1007/s11104-016-2851-z | | | CP002279 | 1 | 2016 |
| 530d63a049607a1be0054a72 | Article | 10.1007/s11104-016-2987-x | | | CP002279 | 1 | 2016 |
| 530d63a049607a1be0054a72 | Article | 10.1007/s11104-018-3830-3 | | | CP002279 | 1 | 2018 |
| 530d63a049607a1be0054a72 | Article | 10.1134/s1022795418080045 | | | CP002279 | 1 | 2018 |
| 530d63a049607a1be0054a72 | Article | 10.1007/s13199-015-0368-5 | | | CP002279 | 1 | 2015 |
| 530d63a049607a1be0054a72 | Article | 10.1007/s13199-016-0381-3 | | | CP002279 | 1 | 2015 |
| 530d63a049607a1be0054a72 | Article | 10.1016/j.syapm.2016.09.003 | 27712916 | | CP002279 | 1 | 2016 |
| 530d63a049607a1be0054a72 | Article | 10.1016/j.syapm.2017.01.004 | 28238475 | | CP002279 | 1 | 2017 |
| 530d63a049607a1be0054a72 | Article | 10.1016/j.syapm.2017.01.004 | 28238475 | | NC_015675 | 1 | 2017 |
| 530d63a049607a1be0054a72 | Article | 10.14201/gredos.137084 | | | CP002279 | 1 | 2017 |
| 530d651d49607a1be0054c40 | Article | 10.1186/s40793-015-0018-9 | 26203342 | PMC4511699 | Gp0010091 | 1 | 2015 |
| 530d652b49607a1be0054c54 | Article | 10.1099/mgen.0.000254 | 30777812 | PMC6421345 | ARRT00000000 | 1 | 2019 |
| 537cf6b90d87852a04c5a36c | Article | 10.1016/j.jbiotec.2014.10.024 | 25449545 | | JHYX00000000 | 1 | 2014 |
| 537cf6b90d87852a04c5a36c | Article | 10.1007/s12275-018-7549-x | 29948825 | | JHYX00000000 | 1 | 2018 |
| 5340e53b49607a0614355d27 | Article | 10.1371/journal.pone.0132660 | 26172151 | PMC4501809 | JHZN00000000 | 1 | 2015 |
| 53bc8ab50d8785134026e064 | Article | 10.1007/s00284-016-0995-y | 26858132 | | JQMU00000000 | 1 | 2016 |
| 5314a2eb49607a1be005664f | Article | 10.1007/s10482-020-01422-5 | 32361957 | | AUDI00000000 | 1 | 2020 |
| 5314a2eb49607a1be005664f | Article | 10.1007/s00284-020-02104-1 | 32601835 | | AUDI00000000 | 1 | 2020 |
| 5314a2eb49607a1be005664f | Article | 10.3389/fmicb.2016.01668 | 27833590 | PMC5080296 | AUDI00000000 | 1 | 2016 |
| 52902808067c013e2b0621fb | Article | 10.1186/s40793-016-0216-0 | 28074122 | PMC5217420 | Gi11889 | 1 | 2017 |
| 5938bbc87ded5e4e5bbbeffd | Article | 10.1002/ece3.5254 | 31380022 | PMC6662431 | SRX3939697 | 1 | 2019 |
| 5702184a7ded5e7f7b944d9f | Article | 10.3389/fmicb.2018.02458 | 30459722 | PMC6232825 | SAMN04883176 | 1 | 2018 |
| 535290280d87855a8277bc67 | Article | 10.1186/s12862-019-1447-7 | 31215393 | PMC6582537 | JMLM00000000 | 1 | 2019 |
| 55132fc60d878525404e72db | Article | 10.1146/annurev-marine-120308-081034 | 21141667 | | BX548174 | 1 | 2010 |
| 55132fc60d878525404e72db | Article | 10.1128/aem.01178-18 | 29915114 | PMC6102989 | BX548174 | 1 | 2018 |
| 55132fc60d878525404e72db | Article | 10.1128/aem.02480-07 | 18586962 | PMC2546643 | BX548174 | 1 | 2008 |
| 55132fc60d878525404e72db | Article | 10.1093/bioinformatics/btp609 | 19850757 | PMC2796815 | BX548174 | 1 | 2009 |
| 55132fc60d878525404e72db | Article | 10.1271/bbb.100216 | 20834156 | | BX548174 | 1 | 2010 |
| 55132fc60d878525404e72db | Article | 10.1080/13102818.2008.10817533 | | | NC_005072 | 1 | 2008 |
| 55132fc60d878525404e72db | Article | 10.1186/1471-2148-7-85 | 17550603 | PMC1904183 | NC_005072 | 1 | 2007 |
| 55132fc60d878525404e72db | Article | 10.1186/1471-2164-8-437 | 18045455 | PMC2242806 | BX548174 | 1 | 2007 |
| 55132fc60d878525404e72db | Article | 10.1186/1471-2164-9-274 | 18534010 | PMC2442094 | BX548174 | 1 | 2008 |
| 55132fc60d878525404e72db | Article | 10.1186/1471-2164-8-375 | 17941988 | PMC2190773 | NC_005072 | 1 | 2007 |
| 55132fc60d878525404e72db | Article | 10.1186/1471-2164-8-437 | 18045455 | PMC2242806 | NC_005072 | 1 | 2007 |
| 55132fc60d878525404e72db | Article | 10.1186/s12864-018-5019-9 | 30134835 | PMC6106888 | NC_005072 | 1 | 2018 |
| 55132fc60d878525404e72db | Article | 10.1186/1471-2180-14-11 | 24438106 | PMC3898218 | NC_005072 | 1 | 2014 |
| 55132fc60d878525404e72db | Article | 10.1002/cbdv.200890180 | 18972533 | | NC_005072 | 1 | 2008 |
| 55132fc60d878525404e72db | Article | 10.1016/j.compbiolchem.2005.11.001 | 16439185 | | BX548174 | 1 | 2006 |
| 55132fc60d878525404e72db | Article | 10.1016/j.copbio.2004.03.007 | 15193326 | | BX548174 | 1 | 2004 |
| 55132fc60d878525404e72db | Article | 10.1111/j.1462-2920.2007.01471.x | 18028413 | | BX548174 | 1 | 2007 |
| 55132fc60d878525404e72db | Article | 10.1111/j.1462-2920.2010.02203.x | 20345942 | PMC2955971 | BX548174 | 1 | 2010 |
| 55132fc60d878525404e72db | Article | 10.1111/1462-2920.12742 | 25522910 | | NC_005072 | 1 | 2015 |

| | | | | | | | |
|---|---|---|---|---|---|---|---|
| 55132fc60d878525404e72db | Article | 10.4056/sigs.3416907 | | PMC3570808 | BX548174 | 1 | 2012 |
| 55132fc60d878525404e72db | Article | 10.4056/sigs.3597227 | | PMC3569392 | BX548174 | 1 | 2012 |
| 55132fc60d878525404e72db | Article | 10.1186/s40793-015-0034-9 | 26380633 | PMC4572445 | NC_005072 | 1 | 2015 |
| 55132fc60d878525404e72db | Article | 10.1177/117693430700300001 | 19468314 | PMC2684143 | NC_005072 | 1 | 2007 |
| 55132fc60d878525404e72db | Article | 10.12688/f1000research.9416.1 | | PMC5031134 | GCA_000011465 | 1 | 2016 |
| 55132fc60d878525404e72db | Article | 10.12688/f1000research.9416.3 | 27703668 | PMC5031134 | GCA_000011465 | 1 | 2016 |
| 55132fc60d878525404e72db | Article | 10.1186/gb-2008-9-5-r90 | 18507822 | PMC2441476 | BX548174 | 1 | 2008 |
| 55132fc60d878525404e72db | Article | 10.1186/gb-2004-5-4-r27 | 15059260 | PMC395786 | NC_005072 | 1 | 2004 |
| 55132fc60d878525404e72db | Article | 10.1186/gb-2005-6-2-r14 | 15693943 | PMC551534 | NC_005072 | 1 | 2005 |
| 55132fc60d878525404e72db | Article | 10.1186/gb-2012-13-3-r23 | 22455878 | PMC3439974 | NC_005072 | 1 | 2012 |
| 55132fc60d878525404e72db | Proceeding | 10.1142/9781860949920_0001 | 18546470 | | BX548174 | 1 | 2007 |
| 55132fc60d878525404e72db | Article | 10.3390/ijms20010152 | 30609821 | PMC6337551 | NC_005072 | 1 | 2019 |
| 55132fc60d878525404e72db | Article | 10.1128/jb.188.8.3012-3023.2006 | 16585762 | PMC1446974 | BX548174 | 1 | 2006 |
| 55132fc60d878525404e72db | Article | 10.1177/0748730408316040 | 18487411 | | NC_005072 | 1 | 2008 |
| 55132fc60d878525404e72db | Article | 10.1007/s00239-011-9473-0 | 22210457 | | BX548174 | 1 | 2011 |
| 55132fc60d878525404e72db | Article | 10.1007/s00239-004-0073-0 | 15696373 | | NC_005072 | 1 | 2005 |
| 55132fc60d878525404e72db | Article | 10.1017/s0025315406013403 | | | BX548174 | 1 | 2006 |
| 55132fc60d878525404e72db | Article | 10.4319/lo.2008.53.6.2485 | | | BX548174 | 1 | 2008 |
| 55132fc60d878525404e72db | Article | 10.1111/j.1751-7915.2010.00204.x | 21255351 | PMC3815765 | Gc00152 | 1 | 2010 |
| 55132fc60d878525404e72db | Article | 10.1099/mic.0.27014-0 | 15133090 | | BX548174 | 1 | 2004 |
| 55132fc60d878525404e72db | Article | 10.1099/mic.0.27032-0 | 14993295 | | BX548174 | 1 | 2004 |
| 55132fc60d878525404e72db | Article | 10.1099/mic.0.28400-0 | 16385117 | | NC_005072 | 1 | 2006 |
| 55132fc60d878525404e72db | Article | 10.1128/mmbr.00035-08 | 19487728 | PMC2698417 | BX548174 | 1 | 2009 |
| 55132fc60d878525404e72db | Article | 10.1093/molbev/msl096 | 16926242 | | BX548174 | 1 | 2006 |
| 55132fc60d878525404e72db | Article | 10.1093/molbev/msr081 | 21531921 | PMC3203624 | BX548174 | 1 | 2011 |
| 55132fc60d878525404e72db | Article | 10.1111/mec.12000 | 22989289 | | BX548174 | 1 | 2012 |
| 55132fc60d878525404e72db | Article | 10.1039/b717118h | 18414733 | | BX548174 | 1 | 2008 |
| 55132fc60d878525404e72db | Article | 10.1038/nature01929 | 12944965 | | BX548174 | 1 | 2003 |
| 55132fc60d878525404e72db | Article | 10.1038/nature01947 | 12917642 | | BX548174 | 1 | 2003 |
| 55132fc60d878525404e72db | Article | 10.1093/nar/gks823 | 22941652 | PMC3488255 | BX548174 | 1 | 2012 |
| 55132fc60d878525404e72db | Article | 10.1093/nar/gkh510 | 15096577 | PMC407844 | NC_005072 | 1 | 2004 |
| 55132fc60d878525404e72db | Article | 10.1093/nar/gkh829 | 15371551 | PMC519117 | NC_005072 | 1 | 2004 |
| 55132fc60d878525404e72db | Article | 10.1093/nar/gkm1181 | 18203741 | PMC2241905 | NC_005072 | 1 | 2008 |
| 55132fc60d878525404e72db | Article | 10.1007/s11120-009-9455-x | 19557544 | | NC_005072 | 1 | 2009 |
| 55132fc60d878525404e72db | Article | 10.1007/s11120-020-00762-7 | 32556852 | | NC_005072 | 1 | 2020 |
| 55132fc60d878525404e72db | Article | 10.1093/pcp/pcl026 | 17071624 | | BX548174 | 1 | 2006 |
| 55132fc60d878525404e72db | Article | 10.1093/pcp/pcs126 | 22968452 | | NC_005072 | 1 | 2012 |
| 55132fc60d878525404e72db | Article | 10.1371/journal.pbio.0030144 | 15828858 | PMC1079782 | BX548174 | 1 | 2005 |
| 55132fc60d878525404e72db | Article | 10.1371/journal.pgen.0030231 | 18159947 | PMC2151091 | BX548174 | 1 | 2007 |
| 55132fc60d878525404e72db | Article | 10.1371/journal.pgen.1000173 | 18769676 | PMC2518516 | BX548174 | 1 | 2008 |
| 55132fc60d878525404e72db | Article | 10.1371/journal.pone.0022705 | 21799937 | PMC3142192 | NC_005072 | 1 | 2011 |
| 55132fc60d878525404e72db | Article | 10.1371/journal.pone.0050274 | 23185592 | PMC3502338 | NC_005072 | 1 | 2012 |
| 55132fc60d878525404e72db | Proceeding | 10.1117/12.929653 | | | NC_005072 | 1 | 2012 |
| 55132fc60d878525404e72db | Article | 10.1073/pnas.1420347112 | 25922520 | PMC4418883 | NC_005072 | 1 | 2015 |
| 55132fc60d878525404e72db | Article | 10.1016/j.protis.2015.11.003 | 26709891 | | BX548174 | 1 | 2015 |
| 55132fc60d878525404e72db | Article | 10.1261/rna.2193110 | 20843985 | PMC2957045 | NC_005072 | 1 | 2010 |
| 55132fc60d878525404e72db | Article | 10.4161/rna.24160 | 23535141 | PMC3737342 | BX548174 | 1 | 2013 |
| 55132fc60d878525404e72db | Article | 10.1126/science.1122050 | 16556843 | | BX548174 | 1 | 2006 |
| 55132fc60d878525404e72db | Article | 10.1126/science.1123933 | 16497887 | | BX548174 | 1 | 2006 |
| 55132fc60d878525404e72db | Article | 10.1007/s11430-020-9651-0 | | | BX548174 | 1 | 2020 |
| 55132fc60d878525404e72db | Article | 10.1360/sste-2020-0063 | | | BX548174 | 1 | 2020 |
| 55132fc60d878525404e72db | Article | 10.1038/sdata.2014.34 | 25977791 | PMC4421930 | BX548174 | 1 | 2014 |
| 55132fc60d878525404e72db | Article | 10.1038/srep00486 | 22768379 | PMC3390001 | BX548174 | 1 | 2012 |
| 55132fc60d878525404e72db | Article | 10.1038/s41396-020-0658-7 | 32346084 | PMC7368042 | BX548174 | 1 | 2020 |
| 55132fc60d878525404e72db | Article | 10.1038/ismej.2008.46 | 18509382 | | NC_005072 | 1 | 2008 |
| 55132fc60d878525404e72db | Article | 10.1038/ismej.2010.43 | 20410936 | | NC_005072 | 1 | 2010 |
| 55132fc60d878525404e72db | Chapter | 10.1002/9780470015902.a0022840 | | | BX548174 | 1 | 2010 |
| 55132fc60d878525404e72db | Chapter | 10.1007/978-1-4614-3348-4_22 | | | BX548174 | 1 | 2012 |
| 55132fc60d878525404e72db | Chapter | 10.1007/978-3-7091-0218-3_13 | | | NC_005072 | 1 | 2012 |
| 55132fc60d878525404e72db | Chapter | 10.1007/978-94-007-0388-9_1 | | | NC_005072 | 1 | 2011 |
| 55132fc60d878525404e72db | Chapter | 10.1201/b13853-5 | | | NC_005072 | 1 | 2013 |
| 530d650249607a1be0054c1f | Article | 10.1007/s00253-015-6515-3 | 25776061 | | Gi08825 | 1 | 2015 |
| 530d650249607a1be0054c1f | Article | 10.1186/1944-3277-9-6 | 25780499 | PMC4334631 | AZAM00000000 | 1 | 2014 |

| | | | | | | | |
|---|---|---|---|---|---|---|---|
| 530d650249607a1be0054c1f | Article | 10.1186/1944-3277-9-6 | 25780499 | PMC4334631 | Gi08825 | 1 | 2014 |
| 5314a25f49607a1be00565d3 | Article | 10.1093/gbe/evv220 | 26568374 | PMC4700951 | AQVC00000000 | 1 | 2015 |
| 5314a25f49607a1be00565d3 | Article | 10.1099/ijsem.0.000617 | 26377180 | | AQVC00000000 | 1 | 2015 |
| 5726d57c7ded5e4fbeb8283f | Article | 10.22456/1679-9216.103176 | | | FOJP00000000 | 1 | 2020 |
| 5726d57c7ded5e4fbeb8283f | Article | 10.1016/j.jgar.2021.01.020 | 33577996 | | FOJP00000000 | 1 | 2021 |
| 5726d57c7ded5e4fbeb8283f | Article | 10.1002/mbo3.1101 | 32657018 | PMC7520993 | FOJP00000000 | 1 | 2020 |
| 5726d57c7ded5e4fbeb8283f | Article | 10.1128/msystems.00455-19 | 31506266 | PMC6739104 | GCF_900111835 | 1 | 2019 |
| 5bcb97c046d1e64cf8dadc28 | Article | 10.1038/s41597-019-0222-3 | 31619684 | PMC6795848 | SRP185225 | 1 | 2019 |
| 5af1795664d0b3374773f264 | Article | 10.1128/aem.02012-16 | 27590813 | PMC5086546 | SRX2014391 | 1 | 2016 |
| 5af1795664d0b3374773f264 | Article | 10.3389/fmicb.2019.01883 | 31474963 | PMC6707425 | Gs0085736 | 1 | 2019 |
| 5290433f067c013e2b066087 | Article | 10.1128/genomea.00381-14 | 24874669 | PMC4038874 | JDWH00000000 | 1 | 2014 |
| 52901eb8067c013e2b060dd5 | Article | 10.1007/s10482-013-0017-z | 24022396 | | CP000606 | 1 | 2013 |
| 52901eb8067c013e2b060dd5 | Article | 10.1111/j.1365-2109.2010.02719.x | | | CP000606 | 1 | 2010 |
| 52901eb8067c013e2b060dd5 | Article | 10.1186/s12859-018-2268-1 | 30367596 | PMC6101096 | NC_009092 | 1 | 2018 |
| 52901eb8067c013e2b060dd5 | Article | 10.1186/1471-2164-10-437 | 19758436 | PMC2761423 | CP000606 | 1 | 2009 |
| 52901eb8067c013e2b060dd5 | Article | 10.1186/1471-2164-12-237 | 21569439 | PMC3107185 | NC_009092 | 1 | 2011 |
| 52901eb8067c013e2b060dd5 | Article | 10.1186/1471-2164-9-274 | 18534010 | PMC2442094 | NC_009092 | 1 | 2008 |
| 52901eb8067c013e2b060dd5 | Article | 10.1111/cla.12020 | | | NC_009092 | 1 | 2013 |
| 52901eb8067c013e2b060dd5 | Article | 10.1111/j.1462-2920.2007.01328.x | 17504473 | PMC1974809 | CP000606 | 1 | 2007 |
| 52901eb8067c013e2b060dd5 | Article | 10.1111/j.1574-6968.2007.00757.x | 17490428 | | CP000606 | 1 | 2007 |
| 52901eb8067c013e2b060dd5 | Article | 10.1099/ijs.0.000313 | 25957050 | | CP000606 | 1 | 2015 |
| 52901eb8067c013e2b060dd5 | Article | 10.1099/ijsem.0.001075 | 27064664 | | CP000606 | 1 | 2016 |
| 52901eb8067c013e2b060dd5 | Article | 10.1099/ijsem.0.001878 | 28629494 | | CP000606 | 1 | 2017 |
| 52901eb8067c013e2b060dd5 | Article | 10.1093/jac/dkr335 | 21846670 | | CP000606 | 1 | 2011 |
| 52901eb8067c013e2b060dd5 | Article | 10.1128/jb.00873-08 | 19060147 | PMC2632075 | NC_009092 | 1 | 2008 |
| 52901eb8067c013e2b060dd5 | Article | 10.1016/j.mimet.2011.03.017 | 21477623 | | CP000606 | 1 | 2011 |
| 52901eb8067c013e2b060dd5 | Article | 10.1016/j.marenvres.2011.11.008 | 22197479 | | CP000606 | 1 | 2011 |
| 52901eb8067c013e2b060dd5 | Article | 10.1016/j.margen.2021.100846 | | | CP000606 | 1 | 2021 |
| 52901eb8067c013e2b060dd5 | Article | 10.1111/j.1751-7915.2009.00092.x | 21261910 | PMC3815836 | NC_009092 | 1 | 2009 |
| 52901eb8067c013e2b060dd5 | Article | 10.1007/s00438-009-0439-5 | 19283410 | | CP000606 | 1 | 2009 |
| 52901eb8067c013e2b060dd5 | Article | 10.1371/journal.pone.0048280 | 23110226 | PMC3480492 | CP000606 | 1 | 2012 |
| 52901eb8067c013e2b060dd5 | Article | 10.1371/journal.pone.0117663 | 25768732 | PMC4358887 | CP000606 | 1 | 2015 |
| 52901eb8067c013e2b060dd5 | Article | 10.1371/journal.pone.0058669 | 23516532 | PMC3597637 | NC_009092 | 1 | 2013 |
| 52901eb8067c013e2b060dd5 | Article | 10.1038/ismej.2014.201 | 25350157 | PMC4409154 | NC_009092 | 1 | 2014 |
| 52901eb8067c013e2b060dd5 | Article | 10.1016/j.tim.2010.05.002 | 20646925 | | NC_009092 | 1 | 2010 |
| 533c5e8e49607a0614353b33 | Article | 10.3389/fmicb.2018.02007 | 30186281 | PMC6113628 | SAMN02743880 | 1 | 2018 |
| 530d646049607a1be0054b64 | Article | 10.4056/sigs.3126494 | | PMC3558956 | CP003153 | 1 | 2012 |
| 530d646049607a1be0054b64 | Article | 10.1111/1574-6941.12346 | 24784780 | | CP003153 | 1 | 2014 |
| 530d646049607a1be0054b64 | Article | 10.1128/jb.00124-12 | 22535943 | PMC3347210 | CP003153 | 1 | 2012 |
| 530d646049607a1be0054b64 | Article | 10.1016/j.mimet.2013.06.013 | 23806694 | | CP003153 | 1 | 2013 |
| 530d646049607a1be0054b64 | Article | 10.1128/msystems.00190-16 | 28293682 | PMC5347186 | CP003153 | 1 | 2017 |
| 530d646049607a1be0054b64 | Article | 10.1016/j.syapm.2013.07.001 | 23972399 | | CP003153 | 1 | 2013 |
| 530d646049607a1be0054b64 | Article | 10.1007/s11274-019-2690-1 | 31332532 | | CP003153 | 1 | 2019 |
| 530d63e849607a1be0054acd | Article | 10.3389/fmicb.2016.00919 | 27379049 | PMC4911352 | CP003029 | 1 | 2016 |
| 530d63e849607a1be0054acd | Article | 10.3389/fmicb.2016.01109 | 27471502 | PMC4943939 | CP003029 | 1 | 2016 |
| 530d63dd49607a1be0054abe | Article | 10.4056/sigs.2014648 | 21886865 | PMC3156397 | CP002838 | 1 | 2011 |
| 530d63dd49607a1be0054abe | Article | 10.1007/s00792-013-0616-7 | 24366681 | | CP002838 | 1 | 2013 |
| 530d63dd49607a1be0054abe | Article | 10.3390/life5010949 | 25789552 | PMC4390887 | NC_015931 | 1 | 2015 |
| 530d63dd49607a1be0054abe | Chapter | 10.1002/9781118960608.gbm01325 | | | CP002838 | 1 | 2016 |
| 530d63dd49607a1be0054abe | Chapter | 10.1002/9781118960608.obm00125 | | | CP002838 | 1 | 2016 |
| 530d63dd49607a1be0054abe | Chapter | 10.1002/9781118960608.obm00126 | | | CP002838 | 1 | 2016 |
| 530d659749607a1be0054cdd | Article | 10.3389/fmicb.2019.01457 | 31333602 | PMC6624747 | LANG00000000 | 1 | 2019 |
| 530d659749607a1be0054cdd | Article | 10.1128/genomea.00889-15 | 26251504 | PMC4541282 | LANG00000000 | 1 | 2015 |
| 530d659749607a1be0054cdd | Chapter | 10.1007/978-981-10-4862-3_12 | | | LANG00000000 | 1 | 2017 |
| 530d659749607a1be0054cdd | Article | 10.1007/s13199-016-0390-2 | | | LANG00000000 | 1 | 2016 |
| 5314a28749607a1be00565f9 | Article | 10.1007/s10482-014-0368-0 | 25707905 | | ARJT00000000 | 1 | 2015 |
| 5314a28749607a1be00565f9 | Article | 10.3389/fmicb.2015.00281 | 25914684 | PMC4391266 | ARJT00000000 | 1 | 2015 |
| 5314a28749607a1be00565f9 | Article | 10.3389/fmicb.2018.01849 | 30147685 | PMC6096048 | NZ_KB899391 | 1 | 2018 |
| 5314a28749607a1be00565f9 | Article | 10.1099/ijsem.0.003055 | 30362935 | | ARJT00000000 | 1 | 2018 |
| 530d63ee49607a1be0054ad4 | Article | 10.1007/s10482-014-0213-5 | 24952743 | | CP002771 | 1 | 2014 |
| 530d63ee49607a1be0054ad4 | Article | 10.1177/1177932217733422 | 28989277 | PMC5624349 | NC_015559 | 1 | 2017 |
| 530d63ee49607a1be0054ad4 | Article | 10.4056/sigs.2976373 | 23458837 | PMC3577112 | NC_015559 | 1 | 2012 |
| 530d63ee49607a1be0054ad4 | Article | 10.1099/ijs.0.054809-0 | 24105944 | | CP002771 | 1 | 2013 |

| | | | | | | | |
|---|---|---|---|---|---|---|---|
| 530d63ee49607a1be0054ad4 | Article | 10.1099/ijsem.0.001006 | 26957484 | | CP002771 | 1 | 2016 |
| 530d63ee49607a1be0054ad4 | Article | 10.1099/ijsem.0.002014 | 28771118 | | CP002771 | 1 | 2017 |
| 530d63ee49607a1be0054ad4 | Article | 10.1099/ijsem.0.002374 | 28984571 | | CP002771 | 1 | 2017 |
| 530d63ee49607a1be0054ad4 | Article | 10.1099/ijsem.0.003227 | 30648946 | | CP002771 | 1 | 2019 |
| 530d63ee49607a1be0054ad4 | Article | 10.1099/ijsem.0.003241 | 30688631 | | CP002771 | 1 | 2019 |
| 530d63ee49607a1be0054ad4 | Article | 10.1099/ijsem.0.003631 | 31380736 | | CP002771 | 1 | 2019 |
| 56362f3f0d87852aea5e72b2 | Article | 10.1038/s41598-019-41368-x | 30894631 | PMC6427011 | LT629699 | 1 | 2019 |
| 529029e6067c013e2b0625f6 | Article | 10.1371/journal.pone.0081086 | 24363793 | PMC3867191 | Gi21295 | 0 | 2013 |
| 55e07de50d878556782d8fad | Article | 10.3389/fmicb.2015.00872 | 26379647 | PMC4549626 | Gp0087948 | 1 | 2015 |
| 52901b6c067c013e2b0606cd | Article | 10.1128/aem.00521-17 | 28314726 | PMC5411500 | ASWY00000000 | 1 | 2017 |
| 52901b6c067c013e2b0606cd | Article | 10.1111/1758-2229.12234 | 25382584 | | ASWY00000000 | 1 | 2014 |
| 52901b6c067c013e2b0606cd | Article | 10.1038/nature12352 | 23851394 | | ASWY00000000 | 1 | 2013 |
| 5290253a067c013e2b061bc0 | Article | 10.1186/s12864-017-3955-4 | 28764658 | PMC5540593 | ATVB00000000 | 1 | 2017 |
| 5290253a067c013e2b061bc0 | Article | 10.3389/fmicb.2018.02007 | 30186281 | PMC6113628 | SAMN02441248 | 1 | 2018 |
| 537d25820d87852a04c5b200 | Article | 10.1111/1462-2920.12173 | 23834245 | PMC4056668 | AAWL00000000 | 1 | 2013 |
| 537d25820d87852a04c5b200 | Article | 10.1371/journal.pone.0060120 | 23577086 | PMC3618517 | AAWL00000000 | 1 | 2013 |
| 5314a28749607a1be00565fa | Article | 10.1016/j.nmni.2016.12.013 | 28070336 | PMC5219628 | ARJO00000000 | 1 | 2016 |
| 52901d73067c013e2b060ae4 | Article | 10.3389/fmicb.2017.00048 | 28184216 | PMC5266723 | ARJN00000000 | 1 | 2017 |
| 52cb210e067c0120babfc21c | Article | 10.3389/fgene.2019.00957 | 31749830 | PMC6843070 | SRR3989263 | 1 | 2019 |
| 545aa66d0d878552848903a1 | Article | 10.1099/acmi.0.000019 | 32974515 | PMC7471780 | FTPU01000000 | 1 | 2019 |
| 545aa66d0d878552848903a1 | Article | 10.1186/s40793-017-0242-6 | 28491240 | PMC5422911 | FTPU01000000 | 1 | 2017 |
| 545aa66d0d878552848903a1 | Article | 10.1186/s40793-017-0242-6 | 28491240 | PMC5422911 | Gp0103631 | 1 | 2017 |
| 594d60937ded5e4e5bbd6ad8 | Article | 10.1007/s10532-012-9561-x | 22684212 | PMC3553412 | CP001823 | 1 | 2012 |
| 594d60937ded5e4e5bbd6ad8 | Article | 10.1186/s40793-016-0165-7 | 27340512 | PMC4918011 | CP001823 | 1 | 2016 |
| 594d60937ded5e4e5bbd6ad8 | Article | 10.4056/sigs.2806097 | 22768368 | PMC3387798 | CP001823 | 1 | 2012 |
| 594d60937ded5e4e5bbd6ad8 | Article | 10.4056/sigs.601105 | 21304677 | PMC3035262 | CP001823 | 1 | 2010 |
| 594d60937ded5e4e5bbd6ad8 | Article | 10.1111/1574-6941.12346 | 24784780 | | CP001823 | 1 | 2014 |
| 594d60937ded5e4e5bbd6ad8 | Article | 10.3389/fmicb.2018.02007 | 30186281 | PMC6113628 | SAMN02598446 | 1 | 2018 |
| 594d60937ded5e4e5bbd6ad8 | Article | 10.3389/fmicb.2019.01612 | 31354692 | PMC6640209 | SAMN02598446 | 1 | 2019 |
| 594d60937ded5e4e5bbd6ad8 | Article | 10.1099/ijs.0.059675-0 | 24425739 | | CP001823 | 1 | 2014 |
| 594d60937ded5e4e5bbd6ad8 | Article | 10.1099/ijs.0.060327-0 | 24814334 | | CP001823 | 1 | 2014 |
| 594d60937ded5e4e5bbd6ad8 | Article | 10.1099/ijsem.0.002141 | 28741992 | | CP001823 | 1 | 2017 |
| 594d60937ded5e4e5bbd6ad8 | Article | 10.1038/ismej.2012.150 | 23190731 | PMC3603402 | CP001823 | 1 | 2012 |
| 594d60937ded5e4e5bbd6ad8 | Article | 10.1038/s41396-019-0530-9 | 31624340 | PMC6976673 | CP001823 | 1 | 2019 |
| 594d60937ded5e4e5bbd6ad8 | Article | 10.1111/j.1365-313x.2011.04684.x | 21699590 | | CP001823 | 1 | 2011 |
| 594d60937ded5e4e5bbd6ad8 | Chapter | 10.1002/9781118960608.gbm01563 | | | CP001823 | 1 | 2018 |
| 559c853b0d87852b215059d1 | Article | 10.3389/fmicb.2019.02083 | 31608019 | PMC6767994 | SAMN04488087 | 1 | 2019 |
| 52902786067c013e2b0620e1 | Article | 10.1186/s40793-015-0074-1 | 26478786 | PMC4609095 | AUFE00000000 | 1 | 2015 |
| 52902786067c013e2b0620e1 | Article | 10.1186/s40793-015-0074-1 | 26478786 | PMC4609095 | Gp0009812 | 1 | 2015 |
| 52901d77067c013e2b060aec | Article | 10.1186/1944-3277-10-1 | 25678942 | PMC4315136 | ARKK00000000 | 1 | 2015 |
| 52901d77067c013e2b060aec | Article | 10.1186/1944-3277-10-1 | 25678942 | PMC4315136 | Gi11553 | 1 | 2015 |
| 52901d77067c013e2b060aec | Article | 10.1099/ijsem.0.001282 | 27381261 | | ARKK00000000 | 1 | 2016 |
| 52901d77067c013e2b060aec | Chapter | 10.1007/978-3-642-38922-1_235 | | | ARKK00000000 | 1 | 2014 |
| 5314a27c49607a1be00565ed | Article | 10.1016/j.nmni.2017.07.004 | 28912952 | PMC5583396 | ARGO00000000 | 1 | 2017 |
| 5314a27c49607a1be00565ed | Chapter | 10.1007/978-3-319-44409-3_11 | | | ARGO00000000 | 1 | 2016 |
| 56abd2900d878559e286f3ae | Article | 10.3389/fmicb.2019.02081 | 31551998 | PMC6746948 | SRP117915 | 1 | 2019 |
| 594d60b07ded5e4e5bbd6af7 | Article | 10.1186/1756-0500-6-482 | 24266988 | PMC4222082 | CP001966 | 1 | 2013 |
| 594d60b07ded5e4e5bbd6af7 | Article | 10.4056/sigs.1894556 | 21886861 | PMC3156402 | CP001966 | 1 | 2011 |
| 594d60b07ded5e4e5bbd6af7 | Article | 10.4056/sigs.1894556 | 21886861 | PMC3156402 | Gc01341 | 1 | 2011 |
| 594d60b07ded5e4e5bbd6af7 | Article | 10.3389/fmicb.2018.02007 | 30186281 | PMC6113628 | SAMN00002597 | 1 | 2018 |
| 594d60b07ded5e4e5bbd6af7 | Article | 10.1099/ijsem.0.000963 | 26868112 | | CP001966 | 1 | 2016 |
| 594d60b07ded5e4e5bbd6af7 | Article | 10.1099/ijsem.0.002033 | 28869002 | | CP001966 | 1 | 2017 |
| 594d60b07ded5e4e5bbd6af7 | Article | 10.1099/ijsem.0.003039 | 30300123 | | CP001966 | 1 | 2018 |
| 594d60b07ded5e4e5bbd6af7 | Article | 10.1099/ijsem.0.003069 | 30355399 | | CP001966 | 1 | 2018 |
| 594d60b07ded5e4e5bbd6af7 | Article | 10.1099/ijsem.0.003580 | 31310200 | | CP001966 | 1 | 2019 |
| 594d60b07ded5e4e5bbd6af7 | Article | 10.1128/jcm.02545-14 | 25520439 | PMC4298536 | CP001966 | 1 | 2014 |
| 594d60b07ded5e4e5bbd6af7 | Chapter | 10.1007/978-3-030-11461-9_1 | | | CP001966 | 1 | 2019 |
| 594d60b07ded5e4e5bbd6af7 | Article | 10.1038/ja.2011.57 | 21792208 | | CP001966 | 1 | 2011 |
| 594d60b07ded5e4e5bbd6af7 | Chapter | 10.1007/978-3-642-30138-4_189 | | | CP001966 | 1 | 2014 |
| 594d60b07ded5e4e5bbd6af7 | Article | 10.11606/t.11.2017.tde-06012017-113512 | | | CP001966 | 1 | 2016 |
| 594d609c7ded5e4e5bbd6ae0 | Article | 10.1128/aem.02821-13 | 24242241 | PMC3911101 | CP001820 | 1 | 2013 |
| 594d609c7ded5e4e5bbd6ae0 | Article | 10.1186/s13068-016-0549-3 | 27340431 | PMC4918077 | CP001820 | 1 | 2016 |
| 594d609c7ded5e4e5bbd6ae0 | Article | 10.1111/1462-2920.12173 | 23834245 | PMC4056668 | CP001820 | 1 | 2013 |

| | | | | | | | |
|---|---|---|---|---|---|---|---|
| 594d609c7ded5e4e5bbd6ae0 | Article | 10.4056/sigs.521107 | 21304678 | PMC3035260 | CP001820 | 1 | 2010 |
| 594d609c7ded5e4e5bbd6ae0 | Article | 10.4056/sigs.521107 | 21304678 | PMC3035260 | Gc01152 | 1 | 2010 |
| 594d609c7ded5e4e5bbd6ae0 | Article | 10.2169/internalmedicine.5120-20 | 32963153 | PMC7925290 | CP001820 | 1 | 2020 |
| 594d609c7ded5e4e5bbd6ae0 | Article | 10.1099/ijs.0.063875-0 | 25061065 | | CP001820 | 1 | 2014 |
| 594d609c7ded5e4e5bbd6ae0 | Article | 10.1099/ijsem.0.001888 | 28742009 | | CP001820 | 1 | 2017 |
| 594d609c7ded5e4e5bbd6ae0 | Article | 10.1099/ijsem.0.002966 | 30124399 | | CP001820 | 1 | 2018 |
| 594d609c7ded5e4e5bbd6ae0 | Article | 10.1371/journal.pone.0060120 | 23577086 | PMC3618517 | CP001820 | 1 | 2013 |
| 594d609c7ded5e4e5bbd6ae0 | Article | 10.1371/journal.pone.0086428 | 24466089 | PMC3899262 | CP001820 | 1 | 2014 |
| 594d609c7ded5e4e5bbd6ae0 | Article | 10.1371/journal.pone.0206484 | 31509535 | PMC6738582 | CP001820 | 1 | 2019 |
| 594d609c7ded5e4e5bbd6ae0 | Chapter | 10.1128/9781555817107.ch6 | | | CP001820 | 1 | 2011 |
| 530d650d49607a1be0054c2c | Article | 10.1007/s00253-015-6515-3 | 25776061 | | Gi08905 | 1 | 2015 |
| 530d650d49607a1be0054c2c | Article | 10.1186/1944-3277-9-4 | 25780497 | PMC4334989 | ATTL00000000 | 1 | 2014 |
| 530d650d49607a1be0054c2c | Article | 10.1186/1944-3277-9-4 | 25780497 | PMC4334989 | Gi08905 | 1 | 2014 |
| 569c14480d8785737066d8b5 | Article | 10.1155/2014/148276 | | | CP002297 | 1 | 2014 |
| 569c14480d8785737066d8b5 | Article | 10.1016/j.bej.2014.05.023 | | | CP002297 | 1 | 2014 |
| 569c14480d8785737066d8b5 | Article | 10.1016/j.ibiod.2017.06.014 | | | NC_017310 | 1 | 2018 |
| 569c14480d8785737066d8b5 | Article | 10.1007/s00248-012-0014-1 | 22349905 | | CP002297 | 1 | 2012 |
| 569c14480d8785737066d8b5 | Article | 10.1371/journal.pone.0061215 | 23577209 | PMC3618111 | CP002297 | 1 | 2013 |
| 569c14480d8785737066d8b5 | Article | 10.1371/journal.ppat.1003204 | 23468637 | PMC3585131 | NC_017310 | 1 | 2013 |
| 529018c4067c013e2b060210 | Article | 10.1186/s40793-016-0191-5 | 27721915 | PMC5052931 | Gp0013295 | 1 | 2016 |
| 529018c4067c013e2b060210 | Article | 10.1099/ijsem.0.001006 | 26957484 | | ATZE00000000 | 1 | 2016 |
| 55c388220d87855594c278db | Article | 10.3389/fmicb.2018.00771 | 29765358 | PMC5938378 | SAMN04487782 | 1 | 2018 |
| 551331c60d878525404e7d24 | Article | 10.1007/s10482-007-9144-8 | 17356928 | | NC_007298 | 1 | 2007 |
| 551331c60d878525404e7d24 | Article | 10.1128/aem.03840-12 | 23811518 | PMC3753938 | CP000089 | 1 | 2013 |
| 551331c60d878525404e7d24 | Article | 10.1128/aem.00457-11 | 21666016 | PMC3147481 | NC_007298 | 1 | 2011 |
| 551331c60d878525404e7d24 | Article | 10.1128/aem.04160-13 | 24682302 | PMC4018867 | NC_007298 | 1 | 2014 |
| 551331c60d878525404e7d24 | Article | 10.1007/s00253-010-2731-z | 20582411 | | CP000089 | 1 | 2010 |
| 551331c60d878525404e7d24 | Article | 10.1016/j.biortech.2008.06.063 | 18752938 | | CP000089 | 1 | 2008 |
| 551331c60d878525404e7d24 | Article | 10.1016/j.biortech.2011.01.069 | 21333531 | PMC3081540 | CP000089 | 1 | 2011 |
| 551331c60d878525404e7d24 | Article | 10.1016/j.cej.2008.05.028 | | | CP000089 | 1 | 2009 |
| 551331c60d878525404e7d24 | Article | 10.1016/j.cej.2017.08.095 | | | CP000089 | 1 | 2018 |
| 551331c60d878525404e7d24 | Article | 10.1007/s00284-006-0474-y | 17486405 | | CP000089 | 1 | 2007 |
| 551331c60d878525404e7d24 | Article | 10.1016/j.cbpa.2007.02.027 | 17349816 | | NC_007298 | 1 | 2007 |
| 551331c60d878525404e7d24 | Article | 10.1111/j.1462-2920.2005.00923.x | 16156723 | | CP000089 | 1 | 2005 |
| 551331c60d878525404e7d24 | Article | 10.1111/j.1462-2920.2008.01662.x | 18484997 | | CP000089 | 1 | 2008 |
| 551331c60d878525404e7d24 | Article | 10.1111/j.1462-2920.2006.01087.x | 17107552 | | NC_007298 | 1 | 2006 |
| 551331c60d878525404e7d24 | Article | 10.1111/j.1462-2920.2010.02248.x | 20545743 | | NC_007298 | 1 | 2010 |
| 551331c60d878525404e7d24 | Article | 10.1111/1758-2229.12004 | 23584966 | | CP000089 | 1 | 2012 |
| 551331c60d878525404e7d24 | Article | 10.1177/117693430700300001 | 19468314 | PMC2684143 | NC_007298 | 1 | 2007 |
| 551331c60d878525404e7d24 | Article | 10.1177/117693430700300006 | 19461976 | PMC2684130 | NC_007298 | 1 | 2007 |
| 551331c60d878525404e7d24 | Article | 10.1007/s00792-011-0428-6 | 22212659 | | CP000089 | 1 | 2012 |
| 551331c60d878525404e7d24 | Article | 10.1111/1574-6941.12090 | 23398624 | | CP000089 | 1 | 2013 |
| 551331c60d878525404e7d24 | Article | 10.1111/j.1574-6941.2010.00996.x | 21073490 | | NC_007298 | 1 | 2010 |
| 551331c60d878525404e7d24 | Article | 10.1111/j.1574-6968.2009.01646.x | 19459951 | | NC_007298 | 1 | 2009 |
| 551331c60d878525404e7d24 | Article | 10.1007/s12223-008-0076-0 | 19381472 | | NC_007298 | 1 | 2008 |
| 551331c60d878525404e7d24 | Chapter | 10.1007/978-3-319-50433-9_16 | | | NC_007298 | 1 | 2019 |
| 551331c60d878525404e7d24 | Article | 10.1016/j.heliyon.2019.e03089 | 31922045 | PMC6948241 | NC_007298 | 1 | 2020 |
| 551331c60d878525404e7d24 | Article | 10.1099/ijs.0.057281-0 | 24480906 | | CP000089 | 1 | 2014 |
| 551331c60d878525404e7d24 | Chapter | 10.1007/978-1-84800-255-5_6 | | | CP000089 | 1 | 2009 |
| 551331c60d878525404e7d24 | Article | 10.1074/jbc.m700886200 | 17439954 | | CP000089 | 1 | 2007 |
| 551331c60d878525404e7d24 | Article | 10.1016/j.jbiosc.2013.08.010 | 24095212 | | CP000089 | 1 | 2013 |
| 551331c60d878525404e7d24 | Article | 10.1080/10934529.2011.586266 | 21847790 | | CP000089 | 1 | 2011 |
| 551331c60d878525404e7d24 | Article | 10.1007/s10295-016-1853-4 | 27796612 | | CP000089 | 1 | 2016 |
| 551331c60d878525404e7d24 | Article | 10.1007/s00239-008-9151-z | 18696026 | | NC_007298 | 1 | 2008 |
| 551331c60d878525404e7d24 | Article | 10.7857/jsge.2012.17.6.023 | | | NC_007298 | 1 | 2012 |
| 551331c60d878525404e7d24 | Article | 10.1080/10962247.2012.672396 | 22866576 | | CP000089 | 1 | 2012 |
| 551331c60d878525404e7d24 | Article | 10.1007/s11814-010-0124-8 | | | CP000089 | 1 | 2010 |
| 551331c60d878525404e7d24 | Article | 10.1128/mbio.00044-11 | 21750120 | PMC3132874 | CP000089 | 1 | 2011 |
| 551331c60d878525404e7d24 | Article | 10.1007/s00248-007-9330-2 | 18034358 | | CP000089 | 1 | 2007 |
| 551331c60d878525404e7d24 | Article | 10.1159/000121324 | 18685265 | | CP000089 | 1 | 2008 |
| 551331c60d878525404e7d24 | Article | 10.1099/mic.0.000117 | 25998264 | | NC_007298 | 1 | 2015 |
| 551331c60d878525404e7d24 | Article | 10.1099/mic.0.045344-0 | 21511765 | | NC_007298 | 1 | 2011 |
| 551331c60d878525404e7d24 | Article | 10.1371/journal.pone.0066971 | 23825601 | PMC3692508 | CP000089 | 1 | 2013 |

| | | | | | | | |
|---|---|---|---|---|---|---|---|
| 551331c60d878525404e7d24 | Article | 10.1371/journal.pone.0084000 | 24475028 | PMC3901652 | NC_007298 | 1 | 2014 |
| 551331c60d878525404e7d24 | Proceeding | 10.1117/12.893450 | | | CP000089 | 1 | 2011 |
| 551331c60d878525404e7d24 | Article | 10.1007/s11157-010-9219-2 | | | CP000089 | 1 | 2010 |
| 551331c60d878525404e7d24 | Article | 10.1261/rna.1056108 | 18676618 | PMC2525955 | CP000089 | 1 | 2008 |
| 551331c60d878525404e7d24 | Article | 10.1261/rna.896108 | 18441051 | PMC2390787 | NC_007298 | 1 | 2008 |
| 551331c60d878525404e7d24 | Article | 10.3184/003685013x13818570960538 | 24547671 | | NC_007298 | 1 | 2013 |
| 551331c60d878525404e7d24 | Article | 10.1016/j.soilbio.2008.02.015 | | | CP000089 | 1 | 2008 |
| 551331c60d878525404e7d24 | Article | 10.1111/j.1747-0765.2010.00453.x | | | NC_007298 | 1 | 2010 |
| 551331c60d878525404e7d24 | Article | 10.1038/ismej.2013.189 | 24152716 | PMC3960533 | CP000089 | 1 | 2013 |
| 551331c60d878525404e7d24 | Article | 10.1038/s41396-019-0508-7 | 31562384 | PMC6908604 | NC_007298 | 1 | 2019 |
| 551331c60d878525404e7d24 | Article | 10.3390/w12082220 | | | CP000089 | 1 | 2020 |
| 551331c60d878525404e7d24 | Chapter | 10.1016/b978-012373944-5.00171-1 | | | CP000089 | 1 | 2009 |
| 551331c60d878525404e7d24 | Chapter | 10.1007/978-3-319-44535-9_16-1 | | | NC_007298 | 1 | 2016 |
| 551331c60d878525404e7d24 | Chapter | 10.1007/978-3-540-77587-4_148 | | | NC_007298 | 1 | 2010 |
| 551331c60d878525404e7d24 | Proceeding | 10.1109/cac.2015.7382641 | | | NC_007298 | 1 | 2015 |
| 551331c60d878525404e7d24 | Article | 10.15083/00005528 | | | NC_007298 | 1 | 2012 |
| 594d60887ded5e4e5bbd6acc | Article | 10.1016/j.anaerobe.2015.11.006 | 26612007 | | CP001684 | 1 | 2015 |
| 594d60887ded5e4e5bbd6acc | Article | 10.1007/s10482-014-0212-6 | 24948086 | | CP001684 | 1 | 2014 |
| 594d60887ded5e4e5bbd6acc | Article | 10.4056/sigs.3246665 | 24019984 | PMC3764928 | CP001684 | 1 | 2013 |
| 594d60887ded5e4e5bbd6acc | Article | 10.4056/sigs.37633 | 21304663 | PMC3035243 | CP001684 | 1 | 2009 |
| 594d60887ded5e4e5bbd6acc | Article | 10.4056/sigs.37633 | 21304663 | PMC3035243 | Gc01094 | 1 | 2009 |
| 594d60887ded5e4e5bbd6acc | Article | 10.3389/fmicb.2018.02007 | 30186281 | PMC6113628 | SAMN02598438 | 1 | 2018 |
| 594d60887ded5e4e5bbd6acc | Article | 10.1099/ijsem.0.002917 | 30010523 | | CP001684 | 1 | 2018 |
| 594d60887ded5e4e5bbd6acc | Article | 10.1099/mic.0.038257-0 | 20093288 | | CP001684 | 1 | 2010 |
| 594d60887ded5e4e5bbd6acc | Article | 10.1016/j.yqres.2013.10.006 | | | CP001684 | 1 | 2014 |
| 594d60887ded5e4e5bbd6acc | Chapter | 10.1007/978-3-642-30138-4_343 | | | CP001684 | 1 | 2014 |
| 59a156907ded5e41edd859a6 | Article | 10.1002/mbo3.966 | 31743595 | PMC7002103 | PZZW00000000 | 1 | 2019 |
| 530d651b49607a1be0054c3e | Article | 10.1186/s40793-015-0119-5 | 26664655 | PMC4674904 | ATTR00000000 | 1 | 2015 |
| 529019eb067c013e2b0604cb | Article | 10.3389/fmicb.2020.00008 | 32038594 | PMC6985074 | KB903995 | 1 | 2020 |
| 529019eb067c013e2b0604cb | Article | 10.3389/fmicb.2020.00008 | 32038594 | PMC6985074 | KB904038 | 1 | 2020 |
| 529019eb067c013e2b0604cb | Article | 10.3389/fmicb.2020.00008 | 32038594 | PMC6985074 | NZ_KB903969 | 1 | 2020 |
| 529019eb067c013e2b0604cb | Article | 10.3389/fmicb.2020.00008 | 32038594 | PMC6985074 | NZ_KB904006 | 1 | 2020 |
| 529019eb067c013e2b0604cb | Article | 10.1016/j.ygeno.2020.03.023 | 32240724 | | ARBV00000000 | 1 | 2020 |
| 5621c9240d878540fd707442 | Article | 10.3389/fmicb.2018.02007 | 30186281 | PMC6113628 | SAMN04490356 | 1 | 2018 |
| 530d64a149607a1be0054bb0 | Article | 10.1007/s10482-020-01424-3 | 32399714 | PMC7716859 | CP003364 | 1 | 2020 |
| 530d64a149607a1be0054bb0 | Article | 10.1007/s10482-020-01471-w | 32936355 | | CP003364 | 1 | 2020 |
| 530d64a149607a1be0054bb0 | Article | 10.1007/s10482-020-01424-3 | 32399714 | PMC7716859 | CP003367 | 1 | 2020 |
| 530d64a149607a1be0054bb0 | Article | 10.1186/1944-3277-9-10 | 25780503 | PMC4334474 | CP003364 | 1 | 2014 |
| 530d64a149607a1be0054bb0 | Article | 10.1186/1944-3277-9-10 | 25780503 | PMC4334474 | CP003367 | 1 | 2014 |
| 530d64a149607a1be0054bb0 | Article | 10.1099/ijsem.0.002846 | 29873630 | | CP003364 | 1 | 2018 |
| 52901c1b067c013e2b0607d7 | Article | 10.1186/s40793-015-0072-3 | 26478785 | PMC4609093 | ARCY00000000 | 1 | 2015 |
| 594d60a07ded5e4e5bbd6ae5 | Article | 10.4056/sigs.2144922 | 22180816 | PMC3236038 | CP002017 | 1 | 2011 |
| 594d60a07ded5e4e5bbd6ae5 | Article | 10.4056/sigs.2144922 | 22180816 | PMC3236038 | Gc01268 | 1 | 2011 |
| 594d60a07ded5e4e5bbd6ae5 | Article | 10.1111/1574-6941.12346 | 24784780 | | CP002017 | 1 | 2014 |
| 594d60a07ded5e4e5bbd6ae5 | Article | 10.1099/ijs.0.053199-0 | 23912720 | | CP002017 | 1 | 2013 |
| 594d60a07ded5e4e5bbd6ae5 | Article | 10.1099/ijs.0.061929-0 | 24801156 | | CP002017 | 1 | 2014 |
| 594d60a07ded5e4e5bbd6ae5 | Article | 10.1111/nph.17365 | 33780002 | | CP002017 | 1 | 2021 |
| 594d60a07ded5e4e5bbd6ae5 | Article | 10.1016/j.syapm.2018.10.007 | 30447886 | | CP002017 | 1 | 2018 |
| 594d60a07ded5e4e5bbd6ae5 | Chapter | 10.1002/9781118960608.gbm01615 | | | CP002017 | 1 | 2019 |
| 594d60a07ded5e4e5bbd6ae5 | Chapter | 10.1128/9781555819323.ch1 | | | CP002017 | 1 | 2016 |
| 555518d30d8785178e712cb4 | Article | 10.1128/genomea.00825-16 | 27516514 | PMC4982293 | LIOL00000000 | 1 | 2016 |
| 555518d30d8785178e712cb4 | Chapter | 10.1007/978-981-13-0329-6_2 | | | LIOL00000000 | 1 | 2018 |
| 5314a27b49607a1be00565eb | Article | 10.3389/fmicb.2020.572053 | 33193169 | PMC7641034 | ARGN00000000 | 1 | 2020 |
| 529016ec067c013e2b05fdaa | Article | 10.1186/gb-2013-14-11-r130 | 24286338 | PMC4053759 | AQUH00000000 | 1 | 2013 |
| 529016ec067c013e2b05fdaa | Article | 10.1128/msystems.00232-18 | 30443603 | PMC6234284 | AQUH00000000 | 1 | 2018 |
| 530d64c749607a1be0054bd9 | Article | 10.1186/1944-3277-10-15 | 26203328 | PMC4511579 | CP007033 | 1 | 2015 |
| 530d64c749607a1be0054bd9 | Article | 10.3389/fmicb.2014.00751 | 25610435 | PMC4285132 | CP007033 | 1 | 2014 |
| 530d64c749607a1be0054bd9 | Article | 10.3389/fmicb.2016.00100 | 26903979 | PMC4751268 | CP007033 | 1 | 2016 |
| 530d64c749607a1be0054bd9 | Article | 10.1038/srep33660 | 27641516 | PMC5027565 | CP007033 | 1 | 2016 |
| 530d64c749607a1be0054bd9 | Article | 10.1016/j.syapm.2016.12.001 | 28292625 | | CP007033 | 1 | 2016 |
| 530d64c749607a1be0054bd9 | Chapter | 10.1007/978-3-662-49875-0_15 | | | CP007033 | 1 | 2016 |
| 530d64c749607a1be0054bd9 | Chapter | 10.1007/978-981-10-8542-0_3 | | | CP007033 | 1 | 2018 |
| 52901bfd067c013e2b060790 | Article | 10.1186/1471-2164-13-533 | 23035691 | PMC3496567 | AFYK00000000 | 1 | 2012 |

| | | | | | | | |
|---|---|---|---|---|---|---|---|
| 52901bfd067c013e2b060790 | Article | 10.3389/fmicb.2018.02415 | 30386310 | PMC6200037 | AFYK00000000 | 1 | 2018 |
| 530d647e49607a1be0054b88 | Article | 10.1016/j.gdata.2016.12.014 | 28116241 | PMC5228091 | NC_016584 | 1 | 2016 |
| 530d647e49607a1be0054b88 | Article | 10.1186/1944-3277-10-15 | 26203328 | PMC4511579 | CP003108 | 1 | 2015 |
| 530d647e49607a1be0054b88 | Article | 10.1186/s40793-015-0072-3 | 26478785 | PMC4609093 | Gi08873 | 1 | 2015 |
| 530d647e49607a1be0054b88 | Article | 10.1016/j.ibiod.2017.06.014 | | | NC_016584 | 1 | 2018 |
| 530d647e49607a1be0054b88 | Article | 10.1099/ijsem.0.001494 | 27613234 | | NC_016584 | 1 | 2016 |
| 530d647e49607a1be0054b88 | Article | 10.1128/jb.01392-12 | 23105050 | PMC3486391 | CP003108 | 1 | 2012 |
| 530d647e49607a1be0054b88 | Article | 10.1134/s0026261715050112 | | | CP003108 | 1 | 2015 |
| 530d647e49607a1be0054b88 | Chapter | 10.1128/9781555819323.ch1 | | | CP003108 | 1 | 2016 |
| 530d640e49607a1be0054afe | Article | 10.1038/s41426-018-0061-x | 29618738 | PMC5884818 | CM001437 | 1 | 2018 |
| 530d640e49607a1be0054afe | Article | 10.1099/ijs.0.000054 | 25563922 | | CM001437 | 1 | 2015 |
| 530d640e49607a1be0054afe | Article | 10.1099/ijs.0.054312-0 | 24453232 | | CM001437 | 1 | 2014 |
| 530d640e49607a1be0054afe | Article | 10.1099/ijsem.0.000530 | 26276159 | | CM001437 | 1 | 2015 |
| 530d640e49607a1be0054afe | Article | 10.1099/ijsem.0.001347 | 27469138 | | CM001437 | 1 | 2016 |
| 530d640e49607a1be0054afe | Article | 10.1099/ijsem.0.001920 | 28671526 | | CM001437 | 1 | 2017 |
| 530d640e49607a1be0054afe | Article | 10.1099/ijsem.0.002539 | 29297847 | | CM001437 | 1 | 2018 |
| 530d640e49607a1be0054afe | Article | 10.1186/s13071-014-0517-3 | 25630498 | PMC4329651 | CM001437 | 1 | 2015 |
| 533c5e9149607a0614353b36 | Article | 10.1155/2018/7609847 | 30210264 | PMC6120340 | JHWX00000000 | 1 | 2018 |
| 57d6f1027ded5e3135ba5293 | Article | 10.1128/genomea.01452-17 | 29437089 | PMC5794936 | FNLG00000000 | 1 | 2018 |
| 5314a26b49607a1be00565dd | Article | 10.1038/srep17609 | 26639610 | PMC4671022 | AREW00000000 | 1 | 2015 |
| 53920c660d878575e64354e2 | Article | 10.3390/min8120596 | | | JQKF00000000 | 1 | 2018 |
| 53920c660d878575e64354e2 | Article | 10.1016/j.resmic.2016.06.007 | 27394987 | | JQKF00000000 | 1 | 2016 |
| 53a2651d0d878514c2d0f169 | Article | 10.1128/genomea.00765-16 | 27491994 | PMC4974315 | JPOO00000000 | 1 | 2016 |
| 535290a10d87855a8277bd91 | Article | 10.1186/s12864-020-06808-3 | 32532247 | PMC7291426 | CP007181 | 1 | 2020 |
| 535290a10d87855a8277bd91 | Article | 10.3389/fmicb.2019.00107 | 30804905 | PMC6371046 | CP007181 | 1 | 2019 |
| 535290a10d87855a8277bd91 | Article | 10.3389/fmicb.2018.02167 | 30258424 | PMC6145009 | CP007182 | 1 | 2018 |
| 5ad0ea0864d0b33747708267 | Article | 10.1007/s10096-020-04003-6 | 32808110 | | CP022695 | 1 | 2020 |
| 5ad0ea0864d0b33747708267 | Article | 10.7717/peerj.4210 | 29312831 | PMC5756455 | SAMN07452765 | 1 | 2018 |
| 5af649bc64d0b3374774c1d5 | Article | 10.3389/fmicb.2020.605952 | 33343549 | PMC7738440 | CP033130 | 1 | 2020 |
| 5af649bc64d0b3374774c1d5 | Article | 10.3389/fmicb.2020.578020 | 33042094 | PMC7530245 | CP033131 | 1 | 2020 |
| 5af649bc64d0b3374774c1d5 | Article | 10.2147/idr.s236200 | 32273730 | PMC7106997 | CP033122 | 1 | 2020 |
| 535290d00d87855a8277be0e | Article | 10.1128/aac.00016-06 | 16723550 | PMC1479134 | AB088224 | 1 | 2006 |
| 535290d00d87855a8277be0e | Article | 10.1007/s00253-011-3128-3 | 21336932 | | AB088224 | 1 | 2011 |
| 535290d00d87855a8277be0e | Article | 10.1007/s00203-010-0548-x | 20177662 | | AB088224 | 1 | 2010 |
| 535290d00d87855a8277be0e | Article | 10.1016/j.chembiol.2005.01.009 | 15734652 | | AB088224 | 1 | 2005 |
| 535290d00d87855a8277be0e | Article | 10.1046/j.1365-2958.2003.03523.x | 12791134 | | AB088224 | 1 | 2003 |
| 535290d00d87855a8277be0e | Article | 10.1111/mmi.12904 | 25495952 | | AB088224 | 1 | 2015 |
| 535290d00d87855a8277be0e | Article | 10.1073/pnas.1315492110 | 24191063 | PMC3839717 | AB088224 | 1 | 2013 |
| 535290d00d87855a8277be0e | Article | 10.1038/s41598-019-47406-y | 31358803 | PMC6662830 | AB088224 | 1 | 2019 |
| 545aa65f0d8785528489020f | Article | 10.1371/journal.pone.0100426 | 24963920 | PMC4070988 | NC_017981 | 1 | 2014 |
| 52901752067c013e2b05fea5 | Article | 10.1186/1471-2164-12-523 | 22026465 | PMC3227697 | CP002641 | 1 | 2011 |
| 52901752067c013e2b05fea5 | Article | 10.1186/s12864-016-3267-0 | 28198678 | PMC5310280 | CP002641 | 1 | 2017 |
| 52901752067c013e2b05fea5 | Article | 10.1093/database/baw084 | 27270714 | PMC4911789 | NC_017620 | 1 | 2016 |
| 52901752067c013e2b05fea5 | Article | 10.1601/sigs.2404675 | | PMC3235512 | CP002641 | 1 | 2011 |
| 52901752067c013e2b05fea5 | Article | 10.4056/sigs.2404675 | | | CP002641 | 1 | 2011 |
| 52901752067c013e2b05fea5 | Article | 10.3389/fcimb.2014.00109 | 25161960 | PMC4129442 | NC_017620 | 1 | 2014 |
| 52901752067c013e2b05fea5 | Article | 10.3389/fmicb.2014.00746 | 25620959 | PMC4288039 | CP002641 | 1 | 2015 |
| 52901752067c013e2b05fea5 | Article | 10.3389/fmicb.2018.01910 | 30186253 | PMC6110895 | NC_017620 | 1 | 2018 |
| 5791b7817ded5e31abaa1e27 | Article | 10.1155/2019/6015730 | 30775379 | PMC6350579 | CP012027 | 1 | 2019 |
| 5791b7817ded5e31abaa1e27 | Article | 10.4172/2157-7420.1000310 | | | CP012027 | 1 | 2018 |
| 5791b7817ded5e31abaa1e27 | Article | 10.1128/genomea.01052-15 | 26358608 | PMC4566190 | CP012027 | 1 | 2015 |
| 5791aee87ded5e31abaa12ef | Article | 10.1128/genomea.01363-16 | 27932655 | PMC5146447 | CP015005 | 1 | 2016 |
| 5791aee87ded5e31abaa12ef | Article | 10.1128/genomea.01363-16 | 27932655 | PMC5146447 | CP015006 | 1 | 2016 |
| 5791aee87ded5e31abaa12ef | Article | 10.1128/genomea.01363-16 | 27932655 | PMC5146447 | CP015007 | 1 | 2016 |
| 5791aee87ded5e31abaa12ef | Article | 10.1128/genomea.01363-16 | 27932655 | PMC5146447 | CP015008 | 1 | 2016 |
| 5791aee87ded5e31abaa12ef | Article | 10.1128/genomea.01363-16 | 27932655 | PMC5146447 | CP015009 | 1 | 2016 |
| 5791aee87ded5e31abaa12ef | Article | 10.1111/nph.17365 | 33780002 | | CP015007 | 1 | 2021 |
| 5791aee87ded5e31abaa12ef | Chapter | 10.1002/9781118960608.fbm00374 | | | CP015005 | 1 | 2020 |
| 573f7be37ded5e7796380528 | Article | 10.3389/fpubh.2019.00066 | 31139608 | PMC6519141 | SAMN02741394 | 1 | 2019 |
| 58d413727ded5e7a2afc1175 | Article | 10.1053/j.gastro.2012.07.013 | 22796240 | PMC4236003 | CP010150 | 0 | 2012 |
| 5913849a7ded5e1e49ffea7a | Article | 10.1128/genomea.01411-16 | 27979954 | PMC5159587 | LNCM00000000 | 1 | 2016 |
| 5621c95a0d878540fd7076d1 | Article | 10.1186/s12864-015-2026-y | 26537657 | PMC4634907 | CP011325 | 1 | 2015 |
| 5621c95a0d878540fd7076d1 | Article | 10.1186/s12864-018-4951-z | 30055586 | PMC6064168 | CP011325 | 1 | 2018 |

| | | | | | | | |
|---|---|---|---|---|---|---|---|
| 5621c95a0d878540fd7076d1 | Article | 10.1016/j.fsi.2015.06.014 | 26087276 | | CP011325 | 1 | 2015 |
| 59138ea57ded5e1e49fff93b | Article | 10.3389/fmicb.2016.01902 | 27965637 | PMC5127849 | CP017006 | 1 | 2016 |
| 592475e67ded5e4e5bbaee8d | Article | 10.1128/mra.01232-18 | 30533757 | PMC6256492 | CP030288 | 1 | 2018 |
| 592475e67ded5e4e5bbaee8d | Article | 10.1128/mra.01232-18 | 30533757 | PMC6256492 | CP030289 | 1 | 2018 |
| 592475e67ded5e4e5bbaee8d | Article | 10.1128/mra.01232-18 | 30533757 | PMC6256492 | CP030290 | 1 | 2018 |
| 592475e67ded5e4e5bbaee8d | Article | 10.1128/mra.01232-18 | 30533757 | PMC6256492 | CP030291 | 1 | 2018 |
| 592475e67ded5e4e5bbaee8d | Article | 10.1128/genomea.01718-15 | 26941156 | PMC4777767 | LHSU00000000 | 1 | 2016 |
| 5e96fa0976b4483f14ad6e92 | Article | 10.1038/s41598-019-38789-z | 30886188 | PMC6423291 | QDGB00000000 | 1 | 2019 |
| 52904140067c013e2b065c17 | Article | 10.1007/s00203-020-01892-1 | 32388821 | | AEEC00000000 | 1 | 2020 |
| 52904140067c013e2b065c17 | Article | 10.3389/fmicb.2013.00168 | 23825472 | PMC3695564 | AEEC00000000 | 1 | 2013 |
| 52904140067c013e2b065c17 | Chapter | 10.1007/978-1-4614-9203-0_9 | | | AEEC00000000 | 1 | 2014 |
| 56af96620d878559e28716c2 | Article | 10.3389/fmicb.2017.00297 | 28298905 | PMC5331189 | CP013690 | 1 | 2017 |
| 56af96620d878559e28716c2 | Article | 10.1016/j.jbiotec.2017.02.003 | 28174040 | | CP013690 | 1 | 2017 |
| 56af96620d878559e28716c2 | Article | 10.1016/j.jbiotec.2017.02.003 | 28174040 | | CP013691 | 1 | 2017 |
| 56af96620d878559e28716c2 | Article | 10.1016/j.micpath.2017.09.012 | 28916321 | | CP013690 | 1 | 2017 |
| 56af96620d878559e28716c2 | Article | 10.1016/j.micpath.2017.09.012 | 28916321 | | CP013691 | 1 | 2017 |
| 56af96620d878559e28716c2 | Article | 10.1007/s00438-016-1261-5 | 27796642 | | CP013690 | 1 | 2016 |
| 56af96620d878559e28716c2 | Article | 10.1007/s00438-016-1261-5 | 27796642 | | CP013691 | 1 | 2016 |
| 5e27aac37776dfea0bd9d085 | Article | 10.1186/s12864-017-3804-5 | 28558786 | PMC5450258 | SAMN06324250 | 1 | 2017 |
| 5e27aac37776dfea0bd9d085 | Article | 10.1128/genomea.00637-17 | 28705983 | PMC5511922 | CP019755 | 1 | 2017 |
| 5e27aac37776dfea0bd9d085 | Article | 10.1128/genomea.00637-17 | 28705983 | PMC5511922 | CP019756 | 1 | 2017 |
| 5e27aac37776dfea0bd9d085 | Article | 10.1128/genomea.00637-17 | 28705983 | PMC5511922 | CP019757 | 1 | 2017 |
| 5e27aac37776dfea0bd9d085 | Article | 10.1128/genomea.00637-17 | 28705983 | PMC5511922 | CP019758 | 1 | 2017 |
| 5e27aac37776dfea0bd9d085 | Article | 10.1128/genomea.00637-17 | 28705983 | PMC5511922 | CP019759 | 1 | 2017 |
| 5e27aac37776dfea0bd9d085 | Article | 10.1128/genomea.00637-17 | 28705983 | PMC5511922 | CP019760 | 1 | 2017 |
| 5e27aac37776dfea0bd9d085 | Article | 10.1128/genomea.00637-17 | 28705983 | PMC5511922 | CP019761 | 1 | 2017 |
| 5e27aac37776dfea0bd9d085 | Article | 10.1128/genomea.00637-17 | 28705983 | PMC5511922 | CP019762 | 1 | 2017 |
| 5e27aac37776dfea0bd9d085 | Article | 10.1128/genomea.00637-17 | 28705983 | PMC5511922 | CP019763 | 1 | 2017 |
| 5e27aac37776dfea0bd9d085 | Article | 10.1128/genomea.00637-17 | 28705983 | PMC5511922 | CP019764 | 1 | 2017 |
| 5e27aac37776dfea0bd9d085 | Article | 10.1128/genomea.00637-17 | 28705983 | PMC5511922 | CP019765 | 1 | 2017 |
| 5e27aac37776dfea0bd9d085 | Article | 10.1128/genomea.00637-17 | 28705983 | PMC5511922 | CP019766 | 1 | 2017 |
| 5e27aac37776dfea0bd9d085 | Article | 10.1128/genomea.00637-17 | 28705983 | PMC5511922 | CP019767 | 1 | 2017 |
| 5bd0f21c46d1e64cf8db3f10 | Article | 10.1007/s13762-019-02474-5 | | | CP024160 | 1 | 2019 |
| 5bd0f21c46d1e64cf8db3f10 | Article | 10.1128/genomea.01361-17 | 29167267 | PMC5701492 | CP024160 | 1 | 2017 |
| 52903f70067c013e2b0657de | Article | 10.1007/s10658-014-0403-z | | | AKVS00000000 | 1 | 2014 |
| 52903f70067c013e2b0657de | Article | 10.3389/fpls.2013.00191 | 23781227 | PMC3678301 | AKVS00000000 | 1 | 2013 |
| 52903f70067c013e2b0657de | Article | 10.1007/s10709-014-9793-2 | 25297844 | | AKVS00000000 | 1 | 2014 |
| 52903f70067c013e2b0657de | Article | 10.1371/journal.ppat.1003013 | 23133391 | PMC3486870 | AKVS00000000 | 1 | 2012 |
| 52903f70067c013e2b0657de | Article | 10.1016/j.syapm.2017.11.005 | 29325987 | | AKVS00000000 | 1 | 2017 |
| 57be094b7ded5e0c87136aa3 | Article | 10.1128/genomea.01631-15 | 26868384 | PMC4751308 | BCMS00000000 | 1 | 2016 |
| 598bf25c7ded5e41edd726ad | Article | 10.1128/genomea.00895-16 | 27587814 | PMC5009971 | JQYK00000000 | 1 | 2016 |
| 5aedadb664d0b33747739460 | Article | 10.1038/srep40443 | 28074866 | PMC5225428 | FAXD00000000 | 1 | 2017 |
| 56af92290d878559e2870dd8 | Article | 10.3389/fmicb.2017.00961 | 28611758 | PMC5447721 | JQAO00000000 | 1 | 2017 |
| 56af92290d878559e2870dd8 | Article | 10.3389/fmicb.2018.01408 | 29997608 | PMC6029419 | JQAO00000000 | 1 | 2018 |
| 56af92290d878559e2870dd8 | Article | 10.1128/mra.00901-18 | 30533939 | PMC6256531 | GCA_001029495 | 1 | 2018 |
| 56af92290d878559e2870dd8 | Article | 10.1371/journal.pone.0130296 | 26151935 | PMC4495038 | JQAO00000000 | 1 | 2015 |
| 59138fd17ded5e1e49fffaab | Article | 10.1128/genomea.00392-16 | 27198026 | PMC4888993 | LUAU00000000 | 1 | 2016 |
| 5e96f69776b4483f14ad4886 | Article | 10.1038/s41396-018-0177-y | 29880910 | PMC6092422 | CP027845 | 1 | 2018 |
| 538020d80d87852a04c5bec3 | Article | 10.1371/journal.pone.0076142 | 24155888 | PMC3796525 | APJG00000000 | 1 | 2013 |
| 5621c96d0d878540fd707843 | Article | 10.3390/proceedings2110650 | | | CP010905 | 1 | 2018 |
| 5621c96d0d878540fd707843 | Article | 10.3389/fmicb.2016.01531 | 27733845 | PMC5039349 | CP010905 | 1 | 2016 |
| 5621c96d0d878540fd707843 | Article | 10.1099/ijsem.0.001548 | 27902180 | PMC5244501 | CP010905 | 1 | 2016 |
| 5621c96d0d878540fd707843 | Article | 10.1111/1751-7915.13270 | 29761637 | PMC6011938 | CP010905 | 1 | 2018 |
| 5621c96d0d878540fd707843 | Article | 10.1128/genomea.00276-15 | 25858846 | PMC4392158 | CP010905 | 1 | 2015 |
| 5621c96d0d878540fd707843 | Article | 10.1371/journal.pone.0206484 | 31509535 | PMC6738582 | CP010905 | 1 | 2019 |
| 5621c96d0d878540fd707843 | Article | 10.1038/srep43555 | 28262711 | PMC5337907 | CP010905 | 1 | 2017 |
| 573f76f17ded5e779637fea7 | Article | 10.1038/s41598-019-46588-9 | 31300709 | PMC6625978 | CP009025 | 1 | 2019 |
| 545aa6270d8785528488fe07 | Article | 10.1016/j.meegid.2018.07.004 | 30033383 | | NC_019517 | 1 | 2018 |
| 545aa6270d8785528488fe07 | Article | 10.1371/journal.pone.0080142 | 24278249 | PMC3836978 | NC_019517 | 1 | 2013 |
| 545aa6270d8785528488fe07 | Article | 10.1371/journal.pone.0117163 | 25629728 | PMC4309531 | NC_019517 | 1 | 2015 |
| 545aa6270d8785528488fe07 | Article | 10.1371/journal.pone.0156773 | 27280590 | PMC4900583 | NC_019517 | 1 | 2016 |
| 58d414ad7ded5e7a2afc1332 | Article | 10.1093/cid/ciy691 | 30423046 | | CP017650 | 1 | 2018 |
| 58d40b197ded5e7a2afc056a | Article | 10.1007/s10750-019-04175-z | | | CP018914 | 1 | 2020 |

| | | | | | | | |
|---|---|---|---|---|---|---|---|
| 58d40b197ded5e7a2afc056a | Article | 10.1128/jb.00826-16 | 28031280 | PMC5331673 | CP018914 | 1 | 2017 |
| 58d40b197ded5e7a2afc056a | Article | 10.1093/jme/tjz194 | 31693136 | | CP018914 | 1 | 2019 |
| 58d40b197ded5e7a2afc056a | Article | 10.1186/s13071-019-3377-z | 30909949 | PMC6434777 | CP018914 | 1 | 2019 |
| 58d40b197ded5e7a2afc056a | Article | 10.1007/s00436-018-6036-y | 30078071 | | CP018914 | 1 | 2018 |
| 58d40b197ded5e7a2afc056a | Article | 10.1007/s00436-019-06442-3 | 31473856 | | CP018914 | 1 | 2019 |
| 58d40b197ded5e7a2afc056a | Article | 10.1038/s41598-018-20138-1 | 29382871 | PMC5789838 | CP018914 | 1 | 2018 |
| 58d40b197ded5e7a2afc056a | Article | 10.1016/j.ttbdis.2018.01.005 | 29371125 | | CP018914 | 1 | 2018 |
| 570215f87ded5e7f7b944ab0 | Article | 10.1590/1678-4685-gmb-2016-0228 | 28767122 | PMC5596368 | SAMN02469464 | 1 | 2017 |
| 575cfb337ded5e5e73213be4 | Article | 10.1002/biot.201600120 | 27676587 | | CP014352 | 1 | 2016 |
| 575cfb337ded5e5e73213be4 | Article | 10.1128/genomea.00248-16 | 27198010 | PMC4888987 | CP014352 | 1 | 2016 |
| 575cfb337ded5e5e73213be4 | Article | 10.1128/genomea.00248-16 | 27198010 | PMC4888987 | CP014353 | 1 | 2016 |
| 52901722067c013e2b05fe2d | Article | 10.1007/s00203-017-1370-5 | 28378142 | | CP002914 | 1 | 2017 |
| 52901722067c013e2b05fe2d | Article | 10.1093/bioinformatics/btt273 | 23665771 | PMC3702249 | NC_016010 | 1 | 2013 |
| 52901722067c013e2b05fe2d | Article | 10.1007/s10658-012-0070-x | | | CP002914 | 1 | 2012 |
| 52901722067c013e2b05fe2d | Article | 10.1007/s10658-014-0497-3 | | | CP002914 | 1 | 2014 |
| 52901722067c013e2b05fe2d | Article | 10.1128/jb.05603-11 | 22056938 | PMC3256661 | CP002914 | 1 | 2011 |
| 52901722067c013e2b05fe2d | Article | 10.1128/jb.05777-11 | 21908674 | PMC3209208 | CP002914 | 1 | 2011 |
| 52901722067c013e2b05fe2d | Article | 10.1007/s42161-019-00294-7 | | | CP002914 | 1 | 2019 |
| 52901722067c013e2b05fe2d | Article | 10.3390/microorganisms9030536 | 33807692 | PMC8002079 | CP002914 | 1 | 2021 |
| 52901722067c013e2b05fe2d | Article | 10.1016/j.pmpp.2013.06.002 | | | CP002914 | 1 | 2013 |
| 52901722067c013e2b05fe2d | Article | 10.1016/j.pmpp.2018.07.005 | | | CP002914 | 1 | 2019 |
| 52901722067c013e2b05fe2d | Article | 10.5423/ppj.oa.01.2018.0014 | 30140181 | PMC6097821 | CP002914 | 1 | 2018 |
| 52901722067c013e2b05fe2d | Article | 10.1371/journal.pone.0079704 | 24278159 | PMC3838355 | CP002914 | 1 | 2013 |
| 52901722067c013e2b05fe2d | Article | 10.1371/journal.pone.0084995 | 24416331 | PMC3887016 | CP002914 | 1 | 2014 |
| 52901722067c013e2b05fe2d | Article | 10.1371/journal.pone.0098129 | 24897119 | PMC4045669 | CP002914 | 1 | 2014 |
| 52901722067c013e2b05fe2d | Article | 10.1371/journal.pone.0145035 | 26673755 | PMC4682653 | NC_016010 | 1 | 2015 |
| 52901722067c013e2b05fe2d | Article | 10.1016/j.soilbio.2012.12.010 | | | NC_016010 | 1 | 2013 |
| 52901722067c013e2b05fe2d | Chapter | 10.1007/978-1-4614-9203-0_7 | | | CP002914 | 1 | 2014 |
| 560c3c4b0d878540fd6fcc3f | Article | 10.1128/genomea.00477-15 | 25999572 | PMC4440952 | JZJL00000000 | 1 | 2015 |
| 58ebfa437ded5e52c4b053e3 | Article | 10.1128/genomea.00528-16 | 27284141 | PMC4901232 | CP014349 | 1 | 2016 |
| 58ebfa437ded5e52c4b053e3 | Article | 10.1128/genomea.00528-16 | 27284141 | PMC4901232 | CP014350 | 1 | 2016 |
| 58ebfa437ded5e52c4b053e3 | Article | 10.1128/genomea.00528-16 | 27284141 | PMC4901232 | CP014351 | 1 | 2016 |
| 58ebfa437ded5e52c4b053e3 | Article | 10.1128/genomea.00528-16 | 27284141 | PMC4901232 | CP014792 | 1 | 2016 |
| 58ebfa437ded5e52c4b053e3 | Article | 10.1128/genomea.00528-16 | 27284141 | PMC4901232 | CP014871 | 1 | 2016 |
| 58ebfa437ded5e52c4b053e3 | Article | 10.1128/genomea.00528-16 | 27284141 | PMC4901232 | CP015331 | 1 | 2016 |
| 58ebfa437ded5e52c4b053e3 | Article | 10.1128/genomea.00528-16 | 27284141 | PMC4901232 | CP015332 | 1 | 2016 |
| 58ebfa437ded5e52c4b053e3 | Article | 10.1128/genomea.00528-16 | 27284141 | PMC4901232 | CP015333 | 1 | 2016 |
| 58ebfa437ded5e52c4b053e3 | Article | 10.1128/genomea.00528-16 | 27284141 | PMC4901232 | CP015334 | 1 | 2016 |
| 58ebfa437ded5e52c4b053e3 | Article | 10.1128/genomea.00528-16 | 27284141 | PMC4901232 | CP015335 | 1 | 2016 |
| 58ebfa437ded5e52c4b053e3 | Article | 10.1128/genomea.00528-16 | 27284141 | PMC4901232 | CP015336 | 1 | 2016 |
| 58ebfa437ded5e52c4b053e3 | Article | 10.1128/genomea.00018-17 | 28302772 | PMC5356049 | SAMN04481062 | 1 | 2017 |
| 58ebfa437ded5e52c4b053e3 | Article | 10.1128/genomea.00528-16 | 27284141 | PMC4901232 | SAMN04481062 | 1 | 2016 |
| 58ebfa437ded5e52c4b053e3 | Article | 10.1371/journal.pone.0127997 | 26061173 | PMC4465626 | CP010419 | 1 | 2015 |
| 58ebfa437ded5e52c4b053e3 | Article | 10.3390/vetsci3030019 | 29056727 | PMC5606581 | SAMN04481062 | 1 | 2016 |
| 58ebfa437ded5e52c4b053e3 | Chapter | 10.1002/9781118960608.fbm00308 | | | CP014349 | 1 | 2018 |
| 58ebfa437ded5e52c4b053e3 | Chapter | 10.1002/9781118960608.gbm01246.pub2 | | | CP014349 | 1 | 2018 |
| 58ebfa437ded5e52c4b053e3 | Chapter | 10.1002/9781118960608.gbm01525 | | | CP014349 | 1 | 2019 |
| 545aa6630d878552848902a1 | Article | 10.1186/1471-2164-15-1184 | 25547158 | PMC4464726 | NC_022770 | 1 | 2014 |
| 545aa6630d878552848902a1 | Article | 10.1186/1471-2164-15-855 | 25280881 | PMC4197329 | NC_022770 | 1 | 2014 |
| 545aa6630d878552848902a1 | Article | 10.1128/jvi.01364-14 | 25100842 | PMC4178739 | NC_022770 | 1 | 2014 |
| 545aa6630d878552848902a1 | Article | 10.1128/genomea.00358-15 | 25953175 | PMC4424291 | NC_022770 | 1 | 2015 |
| 545aa6630d878552848902a1 | Article | 10.1128/genomea.00993-15 | 26337888 | PMC4559737 | NC_022770 | 1 | 2015 |
| 545aa6630d878552848902a1 | Article | 10.1128/genomea.01429-14 | 25635028 | PMC4319499 | NC_022770 | 1 | 2015 |
| 545aa6630d878552848902a1 | Article | 10.1128/genomea.01432-14 | 25635030 | PMC4319501 | NC_022770 | 1 | 2015 |
| 52904034067c013e2b06599e | Article | 10.1128/genomea.00280-13 | 23766401 | PMC3707572 | AORI00000000 | 1 | 2013 |
| 5a3732d47ded5e35e94f6ced | Article | 10.1128/genomea.00037-18 | 29472326 | PMC5823998 | CP014862 | 1 | 2018 |
| 5a3732d47ded5e35e94f6ced | Article | 10.1128/genomea.00037-18 | 29472326 | PMC5823998 | CP014863 | 1 | 2018 |
| 5c17fd5b46d1e6422dc41547 | Article | 10.1016/j.gdata.2017.04.002 | 28491497 | PMC5415549 | CP020570 | 1 | 2017 |
| 5c17fd5b46d1e6422dc41547 | Article | 10.3390/microorganisms7090360 | 31533290 | PMC6780108 | CP020570 | 1 | 2019 |
| 5421eabe0d87857459fb9908 | Article | 10.3923/aje.2017.89.100 | | | CP007539 | 1 | 2017 |
| 5421eabe0d87857459fb9908 | Article | 10.3389/fmicb.2018.01531 | 30042755 | PMC6048232 | CP007539 | 1 | 2018 |
| 5421eabe0d87857459fb9908 | Article | 10.1016/j.ibiod.2014.11.012 | | | CP007539 | 1 | 2015 |
| 52904631067c013e2b066737 | Article | 10.1128/genomea.00209-13 | 23661484 | PMC3650443 | AMXK00000000 | 1 | 2013 |

| | | | | | | | |
|---|---|---|---|---|---|---|---|
| 52904631067c013e2b066737 | Article | 10.1111/nph.15182 | 29726587 | | AMXK00000000 | 1 | 2018 |
| 5e573340de209cf12aa18c9d | Article | 10.1099/mgen.0.000234 | 30461375 | PMC6321869 | CP031112 | 1 | 2018 |
| 529015cb067c013e2b05fb19 | Article | 10.3390/genes2040957 | 24710300 | PMC3927595 | GL384839 | 1 | 2011 |
| 529015cb067c013e2b05fb19 | Article | 10.1111/j.1364-3703.2011.00713.x | 21726380 | PMC6640474 | AEAH00000000 | 1 | 2011 |
| 529015cb067c013e2b05fb19 | Article | 10.1371/journal.ppat.1003807 | 24391493 | PMC3879358 | Gi07003 | 1 | 2014 |
| 5d51985bc5ac66db46ea48c1 | Article | 10.1186/s12864-020-6591-3 | 32151246 | PMC7063779 | MOLZ00000000 | 1 | 2020 |
| 5d51985bc5ac66db46ea48c1 | Article | 10.1038/s41598-018-37398-6 | 30718739 | PMC6362013 | MOLZ00000000 | 1 | 2019 |
| 58a4bdcd7ded5e341aa820d5 | Article | 10.1128/genomea.00165-15 | 25814599 | PMC4384139 | JXLT00000000 | 1 | 2015 |
| 560c3d000d878540fd6fd621 | Article | 10.3389/fmicb.2017.02036 | 29097994 | PMC5654357 | CP009241 | 1 | 2017 |
| 530d69ae49607a1be0054f6c | Article | 10.1007/s10482-018-1041-9 | 29484518 | | CP006651 | 1 | 2018 |
| 530d69ae49607a1be0054f6c | Article | 10.1186/1471-2164-15-124 | 24517536 | PMC3925955 | CP006650 | 1 | 2014 |
| 530d69ae49607a1be0054f6c | Article | 10.1186/1471-2164-15-124 | 24517536 | PMC3925955 | CP006651 | 1 | 2014 |
| 530d69ae49607a1be0054f6c | Article | 10.1186/1471-2164-15-124 | 24517536 | PMC3925955 | CP006652 | 1 | 2014 |
| 530d69ae49607a1be0054f6c | Article | 10.1186/1471-2164-15-124 | 24517536 | PMC3925955 | CP006653 | 1 | 2014 |
| 530d69ae49607a1be0054f6c | Article | 10.1186/1471-2164-15-124 | 24517536 | PMC3925955 | CP006654 | 1 | 2014 |
| 530d69ae49607a1be0054f6c | Article | 10.1186/1471-2164-15-124 | 24517536 | PMC3925955 | CP006655 | 1 | 2014 |
| 530d69ae49607a1be0054f6c | Article | 10.3389/fmicb.2015.00852 | 26347732 | PMC4543880 | CP006650 | 1 | 2015 |
| 530d69ae49607a1be0054f6c | Article | 10.3389/fmicb.2015.00852 | 26347732 | PMC4543880 | CP006654 | 1 | 2015 |
| 530d69ae49607a1be0054f6c | Article | 10.3389/fmicb.2015.00852 | 26347732 | PMC4543880 | CP006655 | 1 | 2015 |
| 530d69ae49607a1be0054f6c | Article | 10.1007/s12088-014-0506-4 | 25805905 | PMC4363252 | CP006650 | 1 | 2014 |
| 530d69ae49607a1be0054f6c | Article | 10.1099/ijsem.0.001018 | 26971128 | | CP006651 | 0 | 2016 |
| 530d69ae49607a1be0054f6c | Article | 10.1099/ijsem.0.001222 | 27267079 | | CP006651 | 1 | 2016 |
| 530d69ae49607a1be0054f6c | Article | 10.1099/ijsem.0.001874 | 28598318 | | CP006651 | 1 | 2017 |
| 530d69ae49607a1be0054f6c | Article | 10.1099/ijsem.0.001976 | 28758621 | | CP006651 | 1 | 2017 |
| 530d69ae49607a1be0054f6c | Article | 10.1099/ijsem.0.001990 | 28741997 | | CP006651 | 1 | 2017 |
| 530d69ae49607a1be0054f6c | Article | 10.1099/ijsem.0.001993 | 28809137 | | CP006651 | 1 | 2017 |
| 530d69ae49607a1be0054f6c | Article | 10.1099/ijsem.0.002289 | 28933319 | | CP006651 | 1 | 2017 |
| 530d69ae49607a1be0054f6c | Article | 10.1099/ijsem.0.002793 | 29722645 | | CP006651 | 1 | 2018 |
| 530d69ae49607a1be0054f6c | Article | 10.1099/ijsem.0.003121 | 30457514 | | CP006651 | 1 | 2018 |
| 530d69ae49607a1be0054f6c | Article | 10.1099/ijsem.0.003226 | 30663955 | | CP006651 | 1 | 2019 |
| 530d69ae49607a1be0054f6c | Article | 10.1099/ijsem.0.003265 | 30735117 | | CP006651 | 1 | 2019 |
| 530d69ae49607a1be0054f6c | Article | 10.1099/ijsem.0.003344 | 30888313 | | CP006651 | 1 | 2019 |
| 530d69ae49607a1be0054f6c | Article | 10.1099/ijsem.0.003373 | 30942687 | | CP006651 | 1 | 2019 |
| 530d69ae49607a1be0054f6c | Article | 10.1099/ijsem.0.003561 | 31251720 | | CP006651 | 1 | 2019 |
| 530d69ae49607a1be0054f6c | Article | 10.1007/s42690-020-00310-9 | | | CP006651 | 1 | 2020 |
| 530d69ae49607a1be0054f6c | Article | 10.1002/jctb.5842 | | | CP006650 | 1 | 2018 |
| 530d69ae49607a1be0054f6c | Article | 10.1016/j.margen.2015.01.005 | 25637653 | | CP006651 | 1 | 2015 |
| 530d69ae49607a1be0054f6c | Article | 10.1111/nph.17365 | 33780002 | | CP006652 | 1 | 2021 |
| 530d69ae49607a1be0054f6c | Article | 10.1016/j.plasmid.2015.02.003 | 25752994 | | CP006650 | 1 | 2015 |
| 530d69ae49607a1be0054f6c | Article | 10.1016/j.plasmid.2015.02.003 | 25752994 | | NC_022041 | 1 | 2015 |
| 530d69ae49607a1be0054f6c | Article | 10.1038/s41598-019-44460-4 | 31133656 | PMC6536676 | NC_022041 | 1 | 2019 |
| 5654b8f00d878531d71ea27a | Article | 10.3389/fmicb.2015.00523 | 26074905 | PMC4444821 | CP010415 | 1 | 2015 |
| 5654b8f00d878531d71ea27a | Article | 10.1007/s11033-019-05133-7 | 31659690 | | CP010415 | 1 | 2019 |
| 5654b8f00d878531d71ea27a | Article | 10.1371/journal.pone.0127997 | 26061173 | PMC4465626 | CP010415 | 1 | 2015 |
| 5654b8f00d878531d71ea27a | Article | 10.1371/journal.pone.0127997 | 26061173 | PMC4465626 | CP010416 | 1 | 2015 |
| 5654b8f00d878531d71ea27a | Article | 10.1371/journal.pone.0127997 | 26061173 | PMC4465626 | CP010417 | 1 | 2015 |
| 5654b8f00d878531d71ea27a | Article | 10.1371/journal.pone.0127997 | 26061173 | PMC4465626 | CP010418 | 1 | 2015 |
| 5654b8f00d878531d71ea27a | Article | 10.1371/journal.pone.0127997 | 26061173 | PMC4465626 | CP010420 | 1 | 2015 |
| 5654b8f00d878531d71ea27a | Article | 10.1371/journal.pone.0127997 | 26061173 | PMC4465626 | CP010421 | 1 | 2015 |
| 5340ec9c49607a06143562a7 | Article | 10.1186/1471-2164-11-500 | 20846431 | PMC2996996 | AARU00000000 | 1 | 2010 |
| 5340ec9c49607a06143562a7 | Article | 10.3389/fmicb.2016.01138 | 27499751 | PMC4956650 | AARU02000000 | 1 | 2016 |
| 5340ec9c49607a06143562a7 | Article | 10.1186/s13099-019-0307-8 | 31198443 | PMC6558679 | AARU02000000 | 1 | 2019 |
| 535291080d87855a8277be9c | Article | 10.1002/cbic.201300389 | 24166757 | | EU124663 | 1 | 2013 |
| 535291080d87855a8277be9c | Article | 10.1007/s12223-008-0060-8 | 19085073 | | EU124663 | 1 | 2008 |
| 535291080d87855a8277be9c | Article | 10.1007/s12223-014-0339-x | 25128200 | PMC4194701 | EU124663 | 1 | 2014 |
| 535291080d87855a8277be9c | Article | 10.1039/c7np00047b | 29517100 | | EU124663 | 1 | 2018 |
| 535291080d87855a8277be9c | Article | 10.1371/journal.pone.0084902 | 24386435 | PMC3874040 | EU124663 | 1 | 2013 |
| 535291080d87855a8277be9c | Article | 10.1371/journal.pone.0118850 | 25741696 | PMC4351081 | EU124663 | 1 | 2015 |
| 535291080d87855a8277be9c | Article | 10.1073/pnas.2009306117 | 32958639 | PMC7547251 | EU124663 | 1 | 2020 |
| 5783f7617ded5e34bd91cae8 | Article | 10.1128/mbio.00403-16 | 27247229 | PMC4895104 | CP013838 | 1 | 2016 |
| 5783f7617ded5e34bd91cae8 | Article | 10.1093/ofid/ofx042 | 28470020 | PMC5407211 | CP013838 | 1 | 2017 |
| 5e572f44de209cf12aa16430 | Article | 10.1007/s10468-019-09937-w | | | CP019081 | 0 | 2019 |
| 5e572f44de209cf12aa16430 | Article | 10.3389/fmicb.2018.01408 | 29997608 | PMC6029419 | CP019081 | 1 | 2018 |

| | | | | | | | |
|---|---|---|---|---|---|---|---|
| 5e572f44de209cf12aa16430 | Article | 10.1016/j.jalgebra.2017.02.012 | | | CP019081 | 0 | 2017 |
| 5e572f44de209cf12aa16430 | Article | 10.1016/j.jalgebra.2017.03.020 | | | CP019081 | 0 | 2017 |
| 5e572f44de209cf12aa16430 | Article | 10.1016/j.micpath.2018.12.013 | 30550841 | | CP019081 | 1 | 2018 |
| 5e572f44de209cf12aa16430 | Article | 10.1128/genomea.00161-17 | 28408677 | PMC5391415 | CP019081 | 1 | 2017 |
| 52901dfe067c013e2b060c0b | Article | 10.1128/genomea.00446-13 | 23833139 | PMC3703600 | CP003906 | 1 | 2013 |
| 52901dfe067c013e2b060c0b | Article | 10.7717/peerj.6221 | 30648020 | PMC6330956 | NC_018938 | 1 | 2019 |
| 52901dfe067c013e2b060c0b | Article | 10.1371/journal.pone.0120659 | 25799515 | PMC4370579 | CP003906 | 1 | 2015 |
| 52901dfe067c013e2b060c0b | Article | 10.1371/journal.pone.0083177 | 24358262 | PMC3866246 | NC_018938 | 1 | 2013 |
| 56f1949e7ded5e7f7b938988 | Article | 10.1186/s40793-016-0155-9 | 27182430 | PMC4866403 | CP011564 | 1 | 2016 |
| 56f1949e7ded5e7f7b938988 | Chapter | 10.1002/9781118960608.gbm01496 | | | CP011564 | 1 | 2017 |
| 58ebfb3a7ded5e52c4b05539 | Article | 10.1128/aac.01572-17 | 28874374 | PMC5655077 | CP017149 | 1 | 2017 |
| 58ebfb3a7ded5e52c4b05539 | Article | 10.3389/fmicb.2019.02287 | 31632384 | PMC6779809 | CP017149 | 1 | 2019 |
| 569c16050d8785737066dc2c | Article | 10.1128/genomea.01488-14 | 25657274 | PMC4319599 | JSXW00000000 | 1 | 2015 |
| 569c16050d8785737066dc2c | Article | 10.1080/21505594.2018.1532243 | 30298790 | PMC7000205 | JSXW00000000 | 1 | 2018 |
| 56006e130d8785306f96b0b6 | Article | 10.3390/genes6030714 | 26213974 | PMC4584326 | JZIM00000000 | 1 | 2015 |
| 56af91660d878559e2870c44 | Article | 10.3390/ani10071128 | 32630808 | PMC7401587 | LBCS00000000 | 1 | 2020 |
| 56af91660d878559e2870c44 | Article | 10.1128/genomea.00461-15 | 26159520 | PMC4498107 | LBCS00000000 | 1 | 2015 |
| 58d40e5c7ded5e7a2afc0a7f | Article | 10.1128/aac.00859-17 | 28674050 | PMC5571358 | CP017633 | 1 | 2017 |
| 58d40e5c7ded5e7a2afc0a7f | Article | 10.3389/fmicb.2018.00011 | 29403463 | PMC5786510 | CP017632 | 1 | 2018 |
| 58d40e5c7ded5e7a2afc0a7f | Article | 10.1016/j.jgar.2020.01.009 | 32006752 | | CP017633 | 1 | 2020 |
| 58d40e5c7ded5e7a2afc0a7f | Article | 10.1128/msphere.00137-18 | 30021873 | PMC6052338 | CP017633 | 1 | 2018 |
| 58d40e5c7ded5e7a2afc0a7f | Article | 10.1038/s41598-018-37125-1 | 30679636 | PMC6346057 | CP017632 | 1 | 2019 |
| 58d40e5c7ded5e7a2afc0a7f | Article | 10.1016/j.vetmic.2020.108619 | 32273005 | | CP017633 | 1 | 2020 |
| 545aa65e0d878552848901f9 | Article | 10.3389/fmicb.2018.02021 | 30210484 | PMC6123377 | NC_019933 | 1 | 2018 |
| 545aa65e0d878552848901f9 | Article | 10.1128/jb.01080-13 | 24214944 | PMC3911242 | NC_019933 | 1 | 2013 |
| 545aa65e0d878552848901f9 | Article | 10.1016/j.vetmic.2014.10.015 | 25465667 | | NC_019933 | 1 | 2014 |
| 56af98140d878559e2871a64 | Article | 10.1186/s13068-016-0439-8 | 26839588 | PMC4736469 | CAEJ00000000 | 1 | 2016 |
| 56af98140d878559e2871a64 | Article | 10.1186/1471-2164-15-404 | 24884520 | PMC4070556 | CAEJ00000000 | 1 | 2014 |
| 5e572fe1de209cf12aa168fa | Article | 10.1093/gbe/evx255 | 29220487 | PMC5739047 | LT009748 | 1 | 2017 |
| 5e572fe1de209cf12aa168fa | Article | 10.1093/gbe/evx255 | 29220487 | PMC5739047 | LT009751 | 1 | 2017 |
| 57a9240e7ded5e31abab5e30 | Article | 10.1128/genomea.01662-15 | 26823594 | PMC4732347 | CP013245 | 1 | 2016 |
| 545aa6690d87855284890333 | Article | 10.1590/0074-02760200153 | 32785421 | PMC7416640 | NC_022103 | 1 | 2020 |
| 545aa6690d87855284890333 | Article | 10.3906/zoo-2004-16 | | | NC_022103 | 1 | 2020 |
| 545aa6690d87855284890333 | Article | 10.1111/zph.12362 | 28418192 | PMC5811810 | NC_022103 | 1 | 2017 |
| 545aa6690d87855284890333 | Chapter | 10.1002/9781118818824.ch5 | | | NC_022103 | 1 | 2015 |
| 545aa6690d87855284890333 | Chapter | 10.1007/978-981-13-1562-6_15 | | PMC7120312 | NC_022103 | 1 | 2018 |
| 5726e11f7ded5e4fbeb83730 | Article | 10.1016/j.dib.2019.104257 | 31384648 | PMC6661503 | KQ089875 | 1 | 2019 |
| 573f75667ded5e779637fc97 | Article | 10.1186/1471-2164-15-679 | 25124552 | PMC4153887 | CM001443 | 1 | 2014 |
| 573f75667ded5e779637fc97 | Article | 10.1093/gbe/evt128 | 23985970 | PMC3814188 | CM001443 | 1 | 2013 |
| 573f75667ded5e779637fc97 | Article | 10.1038/ncomms9933 | 26573375 | PMC4660358 | CM001443 | 1 | 2015 |
| 5a15ce5b7ded5e41c307494c | Article | 10.3390/pathogens8010022 | 30781742 | PMC6471051 | NC_031929 | 1 | 2019 |
| 59cc0fd47ded5e2f18693a26 | Article | 10.1134/s0026261718010095 | | | CP016268 | 1 | 2018 |
| 548f5cc60d8785565d46b886 | Article | 10.1186/s13568-018-0646-8 | 30019301 | PMC6049849 | CP009072 | 1 | 2018 |
| 548f5cc60d8785565d46b886 | Article | 10.1016/j.ajog.2018.10.018 | 30832984 | PMC6733039 | CP009072 | 1 | 2019 |
| 548f5cc60d8785565d46b886 | Article | 10.1128/aac.01603-16 | 27671071 | PMC5119044 | CP009072 | 1 | 2016 |
| 548f5cc60d8785565d46b886 | Article | 10.1111/apm.13013 | 31755584 | | CP009072 | 1 | 2020 |
| 548f5cc60d8785565d46b886 | Article | 10.1007/s00253-015-7123-y | 26563552 | | CP009072 | 1 | 2015 |
| 548f5cc60d8785565d46b886 | Article | 10.1186/s12879-018-3449-2 | 30497396 | PMC6267907 | CP009072 | 1 | 2018 |
| 548f5cc60d8785565d46b886 | Article | 10.1016/j.egg.2019.100039 | | | CP009074 | 1 | 2019 |
| 548f5cc60d8785565d46b886 | Article | 10.3389/fmicb.2016.00135 | 26913025 | PMC4753304 | CP009072 | 1 | 2016 |
| 548f5cc60d8785565d46b886 | Article | 10.3389/fmicb.2017.00228 | 28261184 | PMC5306282 | CP009072 | 1 | 2017 |
| 548f5cc60d8785565d46b886 | Article | 10.3389/fvets.2021.636715 | 33718473 | PMC7952442 | CP009072 | 1 | 2021 |
| 548f5cc60d8785565d46b886 | Article | 10.4155/fsoa-2017-0156 | 29796297 | PMC5961450 | CP009072 | 1 | 2018 |
| 548f5cc60d8785565d46b886 | Article | 10.3390/genes11121504 | 33327465 | PMC7764966 | CP009074 | 1 | 2020 |
| 548f5cc60d8785565d46b886 | Article | 10.1186/s13099-017-0204-y | 28943892 | PMC5607484 | CP009072 | 1 | 2017 |
| 548f5cc60d8785565d46b886 | Article | 10.1186/s13099-017-0204-y | 28943892 | PMC5607484 | GCA_000743255 | 1 | 2017 |
| 548f5cc60d8785565d46b886 | Article | 10.2147/idr.s157847 | 29674848 | PMC5898886 | CP009072 | 1 | 2018 |
| 548f5cc60d8785565d46b886 | Article | 10.1016/j.ijmm.2017.09.007 | 29031453 | PMC5792705 | CP009072 | 1 | 2017 |
| 548f5cc60d8785565d46b886 | Article | 10.1016/j.jiac.2020.05.013 | 32571646 | | CP009072 | 1 | 2020 |
| 548f5cc60d8785565d46b886 | Article | 10.1099/jmm.0.000634 | 29143727 | | CP009072 | 1 | 2017 |
| 548f5cc60d8785565d46b886 | Article | 10.1515/jpm-2019-0398 | 31927525 | PMC7147952 | CP009072 | 1 | 2020 |
| 548f5cc60d8785565d46b886 | Article | 10.1128/genomea.00969-14 | 25291776 | PMC4175212 | CP009072 | 1 | 2014 |
| 548f5cc60d8785565d46b886 | Article | 10.1186/s40168-020-00856-3 | 32591016 | PMC7320585 | CP009072 | 1 | 2020 |

| | | | | | | | |
|---|---|---|---|---|---|---|---|
| 548f5cc60d8785565d46b886 | Article | 10.1128/msphere.00210-20 | 32376701 | PMC7203455 | CP009072 | 1 | 2020 |
| 548f5cc60d8785565d46b886 | Article | 10.1038/nature17971 | 27193680 | | CP009072 | 1 | 2016 |
| 548f5cc60d8785565d46b886 | Article | 10.1016/j.plasmid.2018.09.007 | 30248363 | | CP009074 | 1 | 2018 |
| 548f5cc60d8785565d46b886 | Article | 10.1371/journal.pone.0173559 | 28278280 | PMC5344439 | CP009072 | 1 | 2017 |
| 548f5cc60d8785565d46b886 | Article | 10.1371/journal.pone.0178888 | 28636609 | PMC5479536 | CP009072 | 1 | 2017 |
| 548f5cc60d8785565d46b886 | Article | 10.1371/journal.pone.0206484 | 31509535 | PMC6738582 | CP009072 | 1 | 2019 |
| 548f5cc60d8785565d46b886 | Article | 10.1007/s40278-020-83757-6 | | | CP009072 | 1 | 2020 |
| 548f5cc60d8785565d46b886 | Article | 10.1038/s41598-019-38925-9 | 30787338 | PMC6382887 | CP009072 | 1 | 2019 |
| 548f5cc60d8785565d46b886 | Article | 10.1038/s41598-020-64616-x | 32376847 | PMC7203151 | CP009072 | 0 | 2020 |
| 548f5cc60d8785565d46b886 | Article | 10.1038/srep42097 | 28165067 | PMC5292719 | CP009072 | 1 | 2017 |
| 548f5cc60d8785565d46b886 | Article | 10.11606/d.9.2019.tde-02072019-182022 | | | CP009072 | 1 | 2019 |
| 5ab98c1764d0b30c13702b4f | Article | 10.1128/genomea.01073-16 | 27811087 | PMC5095457 | MCNV00000000 | 1 | 2016 |
| 545aa6340d8785528488ff4d | Article | 10.1007/s00705-020-04926-7 | 33559747 | | NC_019925 | 1 | 2021 |
| 545aa6340d8785528488ff4d | Chapter | 10.1007/978-1-4939-7343-9_18 | 29134600 | | NC_019925 | 1 | 2017 |
| 545aa6340d8785528488ff4d | Article | 10.1128/genomea.00521-14 | 25212610 | PMC4161739 | NC_019925 | 1 | 2014 |
| 545aa6340d8785528488ff4d | Article | 10.1073/pnas.1714812115 | 29555775 | PMC5889632 | NC_019925 | 1 | 2018 |
| 560c3c720d878540fd6fce46 | Article | 10.1007/s10482-016-0674-9 | 26979511 | | CP009285 | 1 | 2016 |
| 560c3c720d878540fd6fce46 | Article | 10.1007/s10482-018-1117-6 | 29971704 | | CP009285 | 1 | 2018 |
| 560c3c720d878540fd6fce46 | Article | 10.1134/s0003683817020041 | | | CP009285 | 1 | 2017 |
| 560c3c720d878540fd6fce46 | Article | 10.1099/ijsem.0.001151 | 27188601 | | CP009285 | 1 | 2016 |
| 560c3c720d878540fd6fce46 | Article | 10.1099/ijsem.0.001378 | 27498790 | | CP009285 | 1 | 2016 |
| 560c3c720d878540fd6fce46 | Article | 10.1099/ijsem.0.001404 | 27514529 | | CP009285 | 1 | 2016 |
| 560c3c720d878540fd6fce46 | Article | 10.1099/ijsem.0.001901 | 28671536 | | CP009285 | 1 | 2017 |
| 560c3c720d878540fd6fce46 | Article | 10.1099/ijsem.0.002269 | 29039306 | | CP009285 | 1 | 2017 |
| 560c3c720d878540fd6fce46 | Article | 10.1099/ijsem.0.002356 | 28950930 | | CP009285 | 1 | 2017 |
| 560c3c720d878540fd6fce46 | Article | 10.1099/ijsem.0.002444 | 29068277 | | CP009285 | 1 | 2017 |
| 560c3c720d878540fd6fce46 | Article | 10.1099/ijsem.0.002967 | 30129919 | | CP009285 | 1 | 2018 |
| 560c3c720d878540fd6fce46 | Article | 10.1007/s00248-015-0722-4 | 26714966 | | CP009285 | 1 | 2015 |
| 560c3c720d878540fd6fce46 | Article | 10.1016/j.nmni.2016.01.011 | 26958345 | PMC4773451 | CP009285 | 1 | 2016 |
| 560c3c720d878540fd6fce46 | Article | 10.1016/j.nmni.2016.01.013 | 26958346 | PMC4773480 | CP009285 | 1 | 2016 |
| 560c3c720d878540fd6fce46 | Article | 10.1016/j.nmni.2017.07.004 | 28912952 | PMC5583396 | CP009285 | 1 | 2017 |
| 560c3c720d878540fd6fce46 | Chapter | 10.1007/978-3-319-22171-7_12 | | | CP009285 | 1 | 2015 |
| 5784012d7ded5e34bd91d7d6 | Article | 10.1128/genomea.00770-17 | 29217787 | PMC5721132 | LKIR00000000 | 1 | 2017 |
| 53adb5820d878514c2d128dd | Article | 10.1007/s00253-019-09757-4 | 30972460 | | CP007566 | 1 | 2019 |
| 53adb5820d878514c2d128dd | Article | 10.1111/are.12931 | | | CP007566 | 1 | 2015 |
| 53adb5820d878514c2d128dd | Article | 10.3389/fmicb.2019.02132 | 31572337 | PMC6751286 | CP007566 | 1 | 2019 |
| 53adb5820d878514c2d128dd | Article | 10.1128/genomea.00450-14 | 24855300 | PMC4031339 | CP007566 | 1 | 2014 |
| 53adb5820d878514c2d128dd | Article | 10.1371/journal.pone.0117663 | 25768732 | PMC4358887 | CP007566 | 1 | 2015 |
| 53adb5820d878514c2d128dd | Article | 10.1038/s41598-020-72484-8 | 32968153 | PMC7512022 | CP007566 | 1 | 2020 |
| 53adb5820d878514c2d128dd | Article | 10.1038/srep09833 | 26014286 | PMC4444815 | CP007566 | 1 | 2015 |
| 5340eccb49607a06143562fb | Article | 10.4056/sigs.5399696 | 25197489 | PMC4149003 | ABXH00000000 | 1 | 2014 |
| 5340eccb49607a06143562fb | Article | 10.3389/fmicb.2018.02007 | 30186281 | PMC6113628 | SAMN00008813 | 1 | 2018 |
| 5340eccb49607a06143562fb | Article | 10.1002/mbo3.580 | 29900684 | PMC6182551 | ABXH00000000 | 1 | 2018 |
| 5340eccb49607a06143562fb | Chapter | 10.1007/978-3-642-30138-4_343 | | | ABXH00000000 | 1 | 2014 |
| 5340e14549607a061435562f | Article | 10.1128/jb.01528-12 | 23144373 | PMC3497484 | ALWQ00000000 | 1 | 2012 |
| 538022800d87852a04c5c2cd | Chapter | 10.1007/978-3-030-11461-9_1 | | | AYTE00000000 | 1 | 2019 |
| 538022800d87852a04c5c2cd | Article | 10.1128/genomea.01230-13 | 24459282 | PMC3900914 | AYTE00000000 | 1 | 2014 |
| 538022800d87852a04c5c2cd | Article | 10.1128/genomea.01047-17 | 29025939 | PMC5637499 | Ga0041805 | 1 | 2017 |
| 545aa64a0d8785528489017b | Article | 10.1186/s13071-018-3030-2 | 30081963 | PMC6090806 | NC_020899 | 1 | 2018 |
| 58d40ca17ded5e7a2afc07ac | Article | 10.3389/fmicb.2021.645860 | 33767684 | PMC7985530 | CP012739 | 1 | 2021 |
| 58d40ca17ded5e7a2afc07ac | Article | 10.3389/fmicb.2021.645860 | 33767684 | PMC7985530 | CP012740 | 1 | 2021 |
| 58d40ca17ded5e7a2afc07ac | Article | 10.1186/s13099-016-0104-6 | 27325916 | PMC4913425 | CP012739 | 1 | 2016 |
| 58d40ca17ded5e7a2afc07ac | Article | 10.1186/s13099-016-0104-6 | 27325916 | PMC4913425 | CP012740 | 1 | 2016 |
| 58d40ca17ded5e7a2afc07ac | Article | 10.1186/s13099-016-0104-6 | 27325916 | PMC4913425 | CP012741 | 1 | 2016 |
| 58d40ca17ded5e7a2afc07ac | Article | 10.1016/j.micpath.2017.02.029 | 28235640 | | CP012739 | 1 | 2017 |
| 529f9d55067c0121bf0d9f8a | Article | 10.3389/fmicb.2018.02501 | 30405564 | PMC6207643 | AEQS00000000 | 1 | 2018 |
| 529f9d55067c0121bf0d9f8a | Article | 10.1128/genomea.00786-13 | 24092784 | PMC3790088 | AEQS00000000 | 1 | 2013 |
| 529f9d55067c0121bf0d9f8a | Article | 10.1038/ismej.2011.19 | 21451583 | PMC3160677 | AEQS00000000 | 1 | 2011 |
| 5340e58949607a0614355d91 | Article | 10.1128/aac.01112-15 | 26349828 | PMC4604371 | CP006900 | 1 | 2015 |
| 5340e58949607a0614355d91 | Article | 10.3389/fmicb.2019.00033 | 30761094 | PMC6361800 | CP006900 | 1 | 2019 |
| 5340e58949607a0614355d91 | Article | 10.2217/fmb-2019-0038 | 31762328 | | CP006900 | 1 | 2019 |
| 5340e58949607a0614355d91 | Article | 10.1128/genomea.00427-14 | 24812228 | PMC4014696 | CP006900 | 1 | 2014 |
| 5e3d6af448feb56cc51ca959 | Article | 10.1186/s12864-015-2324-4 | 26694728 | PMC4687380 | JWHU00000000 | 1 | 2015 |

| | | | | | | | |
|---|---|---|---|---|---|---|---|
| 5e3d6af448feb56cc51ca959 | Article | 10.3389/fmicb.2015.00834 | 26441845 | PMC4566036 | JWHU00000000 | 1 | 2015 |
| 5e3d6af448feb56cc51ca959 | Article | 10.1099/mic.0.000053 | 25678547 | | JWHU00000000 | 1 | 2015 |
| 56362d620d87852aea5e5dea | Article | 10.1128/genomea.01020-15 | 26358601 | PMC4566183 | LFXA00000000 | 1 | 2015 |
| 57721a077ded5e34d82975ff | Article | 10.1016/j.gdata.2016.10.003 | 27766204 | PMC5065635 | LVVO00000000 | 1 | 2016 |
| 57721a077ded5e34d82975ff | Article | 10.1186/s40793-017-0223-9 | 28163826 | PMC5282799 | Gp0147190 | 1 | 2017 |
| 57721a077ded5e34d82975ff | Article | 10.1186/s40793-017-0223-9 | 28163826 | PMC5282799 | LVVO00000000 | 1 | 2017 |
| 59a1ac657ded5e41edd85fd2 | Article | 10.1128/genomea.00562-16 | 27313303 | PMC4911482 | LMCA00000000 | 1 | 2016 |
| 55791d4c0d878529e7cb15af | Article | 10.1007/s00284-020-02112-1 | 32661679 | | JRHH00000000 | 1 | 2020 |
| 55791d4c0d878529e7cb15af | Article | 10.1186/s40793-016-0159-5 | 27313837 | PMC4910214 | JRHH00000000 | 1 | 2016 |
| 55791d4c0d878529e7cb15af | Article | 10.1007/s12275-019-9194-4 | 31562606 | | JRHH00000000 | 1 | 2019 |
| 58ebfa447ded5e52c4b053e4 | Article | 10.1128/genomea.00528-16 | 27284141 | PMC4901232 | CP014349 | 1 | 2016 |
| 58ebfa447ded5e52c4b053e4 | Article | 10.1128/genomea.00528-16 | 27284141 | PMC4901232 | CP014350 | 1 | 2016 |
| 58ebfa447ded5e52c4b053e4 | Article | 10.1128/genomea.00528-16 | 27284141 | PMC4901232 | CP014351 | 1 | 2016 |
| 58ebfa447ded5e52c4b053e4 | Article | 10.1128/genomea.00528-16 | 27284141 | PMC4901232 | CP014792 | 1 | 2016 |
| 58ebfa447ded5e52c4b053e4 | Article | 10.1128/genomea.00528-16 | 27284141 | PMC4901232 | CP014871 | 1 | 2016 |
| 58ebfa447ded5e52c4b053e4 | Article | 10.1128/genomea.00528-16 | 27284141 | PMC4901232 | CP015331 | 1 | 2016 |
| 58ebfa447ded5e52c4b053e4 | Article | 10.1128/genomea.00528-16 | 27284141 | PMC4901232 | CP015332 | 1 | 2016 |
| 58ebfa447ded5e52c4b053e4 | Article | 10.1128/genomea.00528-16 | 27284141 | PMC4901232 | CP015333 | 1 | 2016 |
| 58ebfa447ded5e52c4b053e4 | Article | 10.1128/genomea.00528-16 | 27284141 | PMC4901232 | CP015334 | 1 | 2016 |
| 58ebfa447ded5e52c4b053e4 | Article | 10.1128/genomea.00528-16 | 27284141 | PMC4901232 | CP015335 | 1 | 2016 |
| 58ebfa447ded5e52c4b053e4 | Article | 10.1128/genomea.00528-16 | 27284141 | PMC4901232 | CP015336 | 1 | 2016 |
| 58ebfa447ded5e52c4b053e4 | Article | 10.1128/genomea.00018-17 | 28302772 | PMC5356049 | SAMN04481062 | 1 | 2017 |
| 58ebfa447ded5e52c4b053e4 | Article | 10.1128/genomea.00528-16 | 27284141 | PMC4901232 | SAMN04481062 | 1 | 2016 |
| 58ebfa447ded5e52c4b053e4 | Article | 10.3390/vetsci3030019 | 29056727 | PMC5606581 | SAMN04481062 | 1 | 2016 |
| 58ebfa447ded5e52c4b053e4 | Chapter | 10.1002/9781118960608.fbm00308 | | | CP014349 | 1 | 2018 |
| 58ebfa447ded5e52c4b053e4 | Chapter | 10.1002/9781118960608.gbm01246.pub2 | | | CP014349 | 1 | 2018 |
| 58ebfa447ded5e52c4b053e4 | Chapter | 10.1002/9781118960608.gbm01525 | | | CP014349 | 1 | 2019 |
| 57a92aca7ded5e31abab66a7 | Article | 10.1371/journal.ppat.1008348 | 32150591 | PMC7082065 | CP014239 | 1 | 2020 |
| 5af6454164d0b3374774bae5 | Article | 10.1099/mgen.0.000255 | 30810518 | PMC7067039 | SAMN05591547 | 1 | 2019 |

### Table S3: Evaluation materials for DCE results retrieved through PubMed & PubMed Central

| jamo_id | pmid | valid | s_key | s_key_type | t_key | t_key_type | final_db | sample_group |
|---|---|---|---|---|---|---|---|---|
| 51d4d121067c014cd6eb5bbe | 26251504 | 1 | PRJNA157399 | BPA | PRJNA157399 | BPA | NCBI BioProject Database | 1 |
| 5a15d1107ded5e41c3074f6f | 26988035 | 1 | PRJNA195770 | BPA | PRJNA195770 | BPA | NCBI BioProject Database | 1 |
| 51d4caf8067c014cd6eb0566 | 21501500 | 1 | PRJNA19983 | BPA | PRJNA19983 | BPA | NCBI BioProject Database | 1 |
| 51d4caf8067c014cd6eb0566 | 30181924 | 1 | Gi09583 | GiID | ABDF00000000 | GBA | PubMed Central | 1 |
| 51d4caf8067c014cd6eb0566 | 32216757 | 1 | Gi09583 | GiID | SAMN02744059 | BSA | PubMed Central | 1 |
| 51d4c149067c014cd6ea7ede | 21475585 | 1 | PRJNA42009 | BPA | PRJNA42009 | BPA | NCBI BioProject Database | 1 |
| 51d4c149067c014cd6ea7ede | 25736980 | 1 | Gc01549 | GcID | CP002345 | GBA | PubMed Central | 1 |
| 5486590c0d87850ddcd6ce52 | 26203337 | 1 | PRJNA255585 | BPA | PRJNA255585 | BPA | NCBI BioProject Database | 1 |
| 5a8386c364d0b326cdd1dd2d | 31572496 | 1 | PRJNA250595 | BPA | PRJNA250595 | BPA | NCBI BioProject Database | 1 |
| 523b4f2c067c013937080197 | 26203340 | 1 | PRJNA174982 | BPA | PRJNA174982 | BPA | NCBI BioProject Database | 1 |
| 51d4c3e3067c014cd6eaa5e7 | 22180813 | 1 | PRJNA45817 | BPA | PRJNA45817 | BPA | NCBI BioProject Database | 1 |
| 59cd8bed7ded5e2f18696426 | 29679020 | 1 | PRJNA243951 | BPA | PRJNA243951 | BPA | NCBI BioProject Database | 1 |
| 51d4bf7c067c014cd6ea6afe | 21750257 | 1 | PRJNA65273 | BPA | PRJNA65273 | BPA | NCBI BioProject Database | 1 |
| 51d493d6067c014cd6ea1960 | 21475589 | 1 | PRJNA41529 | BPA | PRJNA41529 | BPA | NCBI BioProject Database | 1 |
| 5486590a0d87850ddcd6ce39 | 26203337 | 1 | PRJNA255578 | BPA | PRJNA255578 | BPA | NCBI BioProject Database | 1 |
| 51d4bdc6067c014cd6ea5e5c | 21760885 | 1 | Gi02584 | GiID | ACXX00000000 | GBA | PubMed Central | 1 |
| 51d4bdc6067c014cd6ea5e5c | 29467821 | 1 | Gi02584 | GiID | ACXX00000000 | GBA | PubMed Central | 1 |
| 51d4bdc6067c014cd6ea5e5c | 31338125 | 1 | Gi02584 | GiID | ACXX00000000 | GBA | PubMed Central | 1 |
| 5621c9320d878540fd70755e | 31063502 | 1 | CP003704 | GBA | CP003704 | GBA | PubMed Central | 1 |
| 5621c9320d878540fd70755e | 31063502 | 1 | CP003707 | GBA | CP003707 | GBA | PubMed Central | 1 |
| 51d4c5d0067c014cd6eac2bd | 20672053 | 1 | Gc01100 | GcID | CP001688 | GBA | PubMed Central | 1 |
| 51d4c5d0067c014cd6eac2bd | 21304651 | 1 | Gc01100 | GcID | CP001688 | GBA | PubMed Central | 1 |
| 51d4c5d0067c014cd6eac2bd | 21304667 | 1 | PRJNA27945 | BPA | PRJNA27945 | BPA | NCBI BioProject Database | 1 |
| 51d4c5d0067c014cd6eac2bd | 22396526 | 1 | Gc01100 | GcID | CP001688 | GBA | PubMed Central | 1 |
| 51d4c5d0067c014cd6eac2bd | 22848480 | 1 | Gc01100 | GcID | CP001688 | GBA | PubMed Central | 1 |
| 51d4c5d0067c014cd6eac2bd | 22978470 | 1 | Gc01100 | GcID | CP001688 | GBA | PubMed Central | 1 |
| 51d4c5d0067c014cd6eac2bd | 24003073 | 1 | Gc01100 | GcID | CP001688 | GBA | PubMed Central | 1 |
| 51d4c5d0067c014cd6eac2bd | 31486766 | 1 | Gc01100 | GcID | CP001688 | GBA | PubMed Central | 1 |
| 51d490e8067c014cd6e9fdbd | 23236275 | 1 | PRJNA74753 | BPA | PRJNA74753 | BPA | NCBI BioProject Database | 1 |
| 51d4bdac067c014cd6ea5d94 | 21304718 | 1 | PRJNA20097 | BPA | PRJNA20097 | BPA | NCBI BioProject Database | 1 |
| 51d4bdac067c014cd6ea5d94 | 23409217 | 1 | Gc01039 | GcID | Gc01039 | GcID | PubMed Central | 1 |
| 51d4bdac067c014cd6ea5d94 | 28163823 | 1 | Gc01039 | GcID | CP001622 | GBA | PubMed Central | 1 |
| 51d4bdac067c014cd6ea5d94 | 29104370 | 1 | Gc01039 | GcID | CP001622 | GBA | PubMed Central | 1 |
| 51d4bdac067c014cd6ea5d94 | 29220487 | 1 | Gc01039 | GcID | CP001622 | GBA | PubMed Central | 1 |
| 51d4c14b067c014cd6ea7ef2 | 21886862 | 1 | PRJNA48289 | BPA | PRJNA48289 | BPA | NCBI BioProject Database | 1 |
| 59cace557ded5e2f18690ef6 | 26950333 | 1 | PRJNA262390 | BPA | PRJNA262390 | BPA | NCBI BioProject Database | 1 |
| 51d4c514067c014cd6eab6e6 | 21304733 | 1 | PRJNA33371 | BPA | PRJNA33371 | BPA | NCBI BioProject Database | 1 |
| 5592b1fb0d87854fee5ff0e4 | 23929491 | 1 | CM001240 | GBA | CM001240 | GBA | NCBI Nucleotide Database | 1 |
| 5592b1fb0d87854fee5ff0e4 | 23929491 | 1 | PRJNA42093 | BPA | PRJNA42093 | BPA | NCBI BioProject Database | 1 |
| 548659020d87850ddcd6cdcd | 26203337 | 1 | PRJNA255589 | BPA | PRJNA255589 | BPA | NCBI BioProject Database | 1 |
| 5a1d0c217ded5e41c3078a8e | 30398645 | 1 | PRJNA200618 | BPA | PRJNA200618 | BPA | NCBI BioProject Database | 1 |
| 5486590c0d87850ddcd6ce55 | 26203337 | 1 | PRJNA255608 | BPA | PRJNA255608 | BPA | NCBI BioProject Database | 1 |
| 51d4c81b067c014cd6eae393 | 21304711 | 1 | PRJNA29431 | BPA | PRJNA29431 | BPA | NCBI BioProject Database | 1 |
| 51d492a2067c014cd6ea10ec | 21304702 | 1 | PRJNA37275 | BPA | PRJNA37275 | BPA | NCBI BioProject Database | 1 |
| 53a2653d0d878514c2d0f1b3 | 26112795 | 1 | PRJNA234788 | BPA | PRJNA234788 | BPA | NCBI BioProject Database | 1 |
| 53a2653d0d878514c2d0f1b3 | 30186281 | 1 | SAMN02745412 | BSA | SAMN02745412 | BSA | PubMed Central | 1 |
| 51d4bc76067c014cd6ea5585 | 25977457 | 1 | PRJNA67455 | BPA | PRJNA67455 | BPA | NCBI BioProject Database | 1 |
| 51d4bf8a067c014cd6ea6b8e | 21304646 | 1 | PRJNA19711 | BPA | PRJNA19711 | BPA | NCBI BioProject Database | 1 |
| 51d4bf8a067c014cd6ea6b8e | 22479445 | 1 | Gc01087 | GcID | CP001819 | GBA | PubMed Central | 1 |
| 51d4bf8a067c014cd6ea6b8e | 23991262 | 1 | Gc01087 | GcID | CP001819 | GBA | PubMed Central | 1 |
| 51d4bf8a067c014cd6ea6b8e | 30186281 | 1 | Gc01087 | GcID | SAMN02598426 | BSA | PubMed Central | 1 |
| 51d4ca02067c014cd6eaf9a9 | 21886866 | 1 | PRJNA51777 | BPA | PRJNA51777 | BPA | NCBI BioProject Database | 1 |
| 51d4ca02067c014cd6eaf9a9 | 22347219 | 1 | Gc01720 | GcID | CP002629 | GBA | PubMed Central | 1 |
| 51d4ca02067c014cd6eaf9a9 | 23316187 | 1 | Gc01720 | GcID | CP002629 | GBA | PubMed Central | 1 |
| 51d4c6fe067c014cd6ead5e1 | 21475582 | 1 | PRJNA38283 | BPA | PRJNA38283 | BPA | NCBI BioProject Database | 1 |

| | | | | | | | | |
|---|---|---|---|---|---|---|---|---|
| 51d4c6fe067c014cd6ead5e1 | 31635256 | 1 | Gc01535 | GcID | CP002305 | GBA | PubMed Central | 1 |
| 51d4be7a067c014cd6ea6341 | 22470573 | 1 | Gc01666 | GcID | CP002536 | GBA | PubMed Central | 1 |
| 51d4be7a067c014cd6ea6341 | 22768367 | 1 | PRJNA41911 | BPA | PRJNA41911 | BPA | NCBI BioProject Database | 1 |
| 51d4be7a067c014cd6ea6341 | 31608019 | 1 | SAMN00016729 | BSA | SAMN00016729 | BSA | PubMed Central | 1 |
| 51d4a6bc067c014cd6ea2510 | 21886860 | 1 | PRJNA42243 | BPA | PRJNA42243 | BPA | NCBI BioProject Database | 1 |
| 593fd2bd7ded5e4e5bbc4e40 | 30619145 | 1 | PRJNA444022 | BPA | PRJNA444022 | BPA | NCBI BioProject Database | 1 |
| 51d4910b067c014cd6e9ff55 | 21677856 | 1 | PRJNA40055 | BPA | PRJNA40055 | BPA | NCBI BioProject Database | 1 |
| 51d4910b067c014cd6e9ff55 | 31509535 | 1 | Gc01665 | GcID | CP002530 | GBA | PubMed Central | 1 |
| 59138c6e7ded5e1e49fff62e | 27047476 | 1 | Gs0090290 | GsID | Gs0090290 | GsID | PubMed Central | 1 |
| 59138c6e7ded5e1e49fff62e | 28270584 | 1 | PRJNA269163 | BPA | PRJNA269163 | BPA | NCBI BioProject Database | 1 |
| 59138c6e7ded5e1e49fff62e | 28350381 | 1 | MTOR00000000 | GBA | MTOR00000000 | GBA | NCBI Nucleotide Database | 1 |
| 59138c6e7ded5e1e49fff62e | 28396823 | 1 | SAMN03166137 | BSA | SAMN03166137 | BSA | PubMed Central | 1 |
| 51d4cde0067c014cd6eb2dc8 | 21677852 | 1 | PRJNA37901 | BPA | PRJNA37901 | BPA | NCBI BioProject Database | 1 |
| 51d4ce4c067c014cd6eb3334 | 21475588 | 1 | PRJNA32825 | BPA | PRJNA32825 | BPA | NCBI BioProject Database | 1 |
| 51d4ce4c067c014cd6eb3334 | 25780503 | 1 | Gc01591 | GcID | CP002353 | GBA | PubMed Central | 1 |
| 51d4ce4c067c014cd6eb3334 | 28360896 | 1 | Gc01591 | GcID | CP002353 | GBA | PubMed Central | 1 |
| 51d4ce4c067c014cd6eb3334 | 32399714 | 1 | Gc01591 | GcID | CP002353 | GBA | PubMed Central | 1 |
| 51d4ce4c067c014cd6eb3334 | 33414774 | 1 | Gc01591 | GcID | CP002353 | GBA | PubMed Central | 1 |
| 59cd8a497ded5e2f18696152 | 29551995 | 1 | PRJNA234811 | BPA | PRJNA234811 | BPA | NCBI BioProject Database | 1 |
| 51d4ca26067c014cd6eafab5 | 25189582 | 1 | PRJNA63567 | BPA | PRJNA63567 | BPA | NCBI BioProject Database | 1 |
| 51d4ca8a067c014cd6eaff9d | 21677850 | 1 | PRJNA41547 | BPA | PRJNA41547 | BPA | NCBI BioProject Database | 1 |
| 58fa8fcc7ded5e1e49fe17a8 | 27774985 | 1 | MGII00000000 | GBA | MGII00000000 | GBA | NCBI Nucleotide Database | 1 |
| 52f566b4067c011a2114007f | 23469353 | 1 | PRJNA200605 | BPA | PRJNA200605 | BPA | NCBI BioProject Database | 1 |
| 51d4c4a0067c014cd6eab0a9 | 21695235 | 1 | PRJNA19047 | BPA | PRJNA19047 | BPA | NCBI BioProject Database | 1 |
| 51d55e7a067c014cd6f12f7c | 17803764 | 1 | Gc00590 | GcID | CP000738 | GBA | PubMed Central | 1 |
| 51d55e7a067c014cd6f12f7c | 19758436 | 1 | Gc00590 | GcID | CP000738 | GBA | PubMed Central | 1 |
| 51d55e7a067c014cd6f12f7c | 21304680 | 1 | PRJNA16304 | BPA | PRJNA16304 | BPA | NCBI BioProject Database | 1 |
| 51d55e7a067c014cd6f12f7c | 22419837 | 1 | Gc00590 | GcID | CP000738 | GBA | PubMed Central | 1 |
| 51d55e7a067c014cd6f12f7c | 23409217 | 1 | Gc00590 | GcID | Gc00590 | GcID | PubMed Central | 1 |
| 51d55e7a067c014cd6f12f7c | 23432981 | 1 | Gc00590 | GcID | CP000738 | GBA | PubMed Central | 1 |
| 51d55e7a067c014cd6f12f7c | 24736785 | 1 | Gc00590 | GcID | CP000738 | GBA | PubMed Central | 1 |
| 51d55e7a067c014cd6f12f7c | 29104370 | 1 | Gc00590 | GcID | CP000738 | GBA | PubMed Central | 1 |
| 51d55e7a067c014cd6f12f7c | 29220487 | 1 | Gc00590 | GcID | CP000738 | GBA | PubMed Central | 1 |
| 51d55e7a067c014cd6f12f7c | 33519831 | 1 | Gc00590 | GcID | CP000738 | GBA | PubMed Central | 1 |
| 51d68ef5067c014cd6012202 | 17803764 | 1 | Gc00618 | GcID | CP000769 | GBA | PubMed Central | 1 |
| 51d68ef5067c014cd6012202 | 25280065 | 1 | Gc00618 | GcID | CP000769 | GBA | PubMed Central | 1 |
| 51d68ef5067c014cd6012202 | 25614562 | 1 | PRJNA17729 | BPA | PRJNA17729 | BPA | NCBI BioProject Database | 1 |
| 51d4cec7067c014cd6eb399a | 22768359 | 1 | PRJNA40057 | BPA | PRJNA40057 | BPA | NCBI BioProject Database | 1 |
| 51d4910c067c014cd6e9ff5f | 21677860 | 1 | PRJNA40779 | BPA | PRJNA40779 | BPA | NCBI BioProject Database | 1 |
| 51d4c88f067c014cd6eae972 | 26441909 | 1 | PRJNA32711 | BPA | PRJNA32711 | BPA | NCBI BioProject Database | 1 |
| 51d4a5f6067c014cd6ea22aa | 22675589 | 1 | PRJNA51139 | BPA | PRJNA51139 | BPA | NCBI BioProject Database | 1 |
| 51d4a536067c014cd6ea2051 | 23407703 | 1 | PRJNA40777 | BPA | PRJNA40777 | BPA | NCBI BioProject Database | 1 |
| 59cad6347ded5e2f186918a7 | 27601008 | 1 | PRJNA196014 | BPA | PRJNA196014 | BPA | NCBI BioProject Database | 1 |
| 533f325349607a0614354bf3 | 25977428 | 1 | PRJNA215341 | BPA | PRJNA215341 | BPA | NCBI BioProject Database | 1 |
| 51d4c2ca067c014cd6ea9493 | 21475586 | 1 | PRJNA41913 | BPA | PRJNA41913 | BPA | NCBI BioProject Database | 1 |
| 51d4c2ca067c014cd6ea9493 | 27408745 | 1 | Gc01593 | GcID | CP002352 | GBA | PubMed Central | 1 |
| 51d4c2ca067c014cd6ea9493 | 29321938 | 1 | Gc01593 | GcID | CP002352 | GBA | PubMed Central | 1 |
| 51d4c2ca067c014cd6ea9493 | 31509535 | 1 | Gc01593 | GcID | CP002352 | GBA | PubMed Central | 1 |
| 523b41ce067c01393707f440 | 26767090 | 1 | PRJNA163045 | BPA | PRJNA163045 | BPA | NCBI BioProject Database | 1 |
| 51d4cec5067c014cd6eb3988 | 22768366 | 1 | PRJNA33163 | BPA | PRJNA33163 | BPA | NCBI BioProject Database | 1 |
| 51d4cec6067c014cd6eb3991 | 21304735 | 1 | PRJNA32577 | BPA | PRJNA32577 | BPA | NCBI BioProject Database | 1 |
| 51d4cec6067c014cd6eb3991 | 28116104 | 1 | Gc01413 | GcID | CP002281 | GBA | PubMed Central | 1 |
| 51d4ccae067c014cd6eb1d56 | 21304695 | 1 | PRJNA20741 | BPA | PRJNA20741 | BPA | NCBI BioProject Database | 1 |
| 51d4caba067c014cd6eb0239 | 21475591 | 1 | Gc01553 | GcID | CP002361 | GBA | PubMed Central | 1 |
| 51d4caba067c014cd6eb0239 | 21677858 | 1 | PRJNA40223 | BPA | PRJNA40223 | BPA | NCBI BioProject Database | 1 |
| 51d4caba067c014cd6eb0239 | 31608019 | 1 | SAMN00138957 | BSA | SAMN00138957 | BSA | PubMed Central | 1 |
| 51d4bd7d067c014cd6ea5c2e | 21677853 | 1 | PRJNA43461 | BPA | PRJNA43461 | BPA | NCBI BioProject Database | 1 |
| 51d4bd7d067c014cd6ea5c2e | 31608019 | 1 | Gc01597 | GcID | SAMN00713579 | BSA | PubMed Central | 1 |

| | | | | | | | | | |
|---|---|---|---|---|---|---|---|---|---|
| 51d4a462067c014cd6ea1dc2 | 22675590 | 1 | PRJNA51497 | BPA | PRJNA51497 | BPA | NCBI BioProject Database | | 1 |
| 51d493b9067c014cd6ea1912 | 21475592 | 1 | PRJNA32993 | BPA | PRJNA32993 | BPA | NCBI BioProject Database | | 1 |
| 51d493b9067c014cd6ea1912 | 26116678 | 1 | Gc01599 | GcID | CP002364 | GBA | PubMed Central | | 1 |
| 51d493b9067c014cd6ea1912 | 28217124 | 1 | Gc01599 | GcID | CP002364 | GBA | PubMed Central | | 1 |
| 51d493b9067c014cd6ea1912 | 32060020 | 1 | Gc01599 | GcID | CP002364 | GBA | PubMed Central | | 1 |
| 546bcfa70d8785221c60842b | 30297742 | 1 | PRJNA421608 | BPA | PRJNA421608 | BPA | NCBI BioProject Database | | 1 |
| 51d4c31b067c014cd6ea9999 | 21677851 | 1 | PRJNA41989 | BPA | PRJNA41989 | BPA | NCBI BioProject Database | | 1 |
| 51d4c31b067c014cd6ea9999 | 22768127 | 1 | Gc01548 | GcID | CP002346 | GBA | PubMed Central | | 1 |
| 51d4c31b067c014cd6ea9999 | 25303276 | 1 | Gc01548 | GcID | CP002346 | GBA | PubMed Central | | 1 |
| 51d4c31b067c014cd6ea9999 | 26107936 | 1 | Gc01548 | GcID | CP002346 | GBA | PubMed Central | | 1 |
| 51d4c31b067c014cd6ea9999 | 26928424 | 1 | Gc01548 | GcID | CP002346 | GBA | PubMed Central | | 1 |
| 51d4c31b067c014cd6ea9999 | 27373315 | 1 | Gc01548 | GcID | CP002346 | GBA | PubMed Central | | 1 |
| 51d4c31b067c014cd6ea9999 | 27577199 | 1 | Gc01548 | GcID | CP002346 | GBA | PubMed Central | | 1 |
| 51d4ca46067c014cd6eafc58 | 21677859 | 1 | PRJNA50743 | BPA | PRJNA50743 | BPA | NCBI BioProject Database | | 1 |
| 51d4ca46067c014cd6eafc58 | 26745366 | 1 | Gc01668 | GcID | CP002534 | GBA | PubMed Central | | 1 |
| 59cad2eb7ded5e2f186913fd | 24926052 | 1 | PRJNA196019 | BPA | PRJNA196019 | BPA | NCBI BioProject Database | | 1 |
| 548659000d87850ddcd6cdb1 | 26203337 | 1 | PRJNA255603 | BPA | PRJNA255603 | BPA | NCBI BioProject Database | | 1 |
| 55e07f090d878556782d9b3f | 26837824 | 1 | SAMN03203005 | BSA | SAMN03203005 | BSA | PubMed Central | | 1 |
| 51d492be067c014cd6ea1224 | 30519708 | 1 | 640281010 | IMG_Tax | NC_009042 | GBA | PubMed Central | | 1 |
| 51d4cebc067c014cd6eb3913 | 21304734 | 1 | PRJNA43527 | BPA | PRJNA43527 | BPA | NCBI BioProject Database | | 1 |
| 51d4cebc067c014cd6eb3913 | 30186281 | 1 | Gc01572 | GcID | SAMN00713569 | BSA | PubMed Central | | 1 |
| 59cad9b37ded5e2f18691d7e | 25838471 | 1 | PRJNA250607 | BPA | PRJNA250607 | BPA | NCBI BioProject Database | | 1 |
| 51d4bc77067c014cd6ea5592 | 25977457 | 1 | PRJNA68631 | BPA | PRJNA68631 | BPA | NCBI BioProject Database | | 1 |
| 51d4c72f067c014cd6ead843 | 21886858 | 1 | PRJNA32591 | BPA | PRJNA32591 | BPA | NCBI BioProject Database | | 1 |
| 51d4c72f067c014cd6ead843 | 28497061 | 1 | Gc01415 | GcID | CP002175 | GBA | PubMed Central | | 1 |
| 51d4a62a067c014cd6ea2340 | 27602057 | 1 | Gc01579 | GcID | CP002416 | GBA | PubMed Central | | 1 |
| 51d4a62a067c014cd6ea2340 | 28053665 | 1 | Gc01579 | GcID | CP002416 | GBA | PubMed Central | | 1 |
| 51d4a62a067c014cd6ea2340 | 28230109 | 1 | Gc01579 | GcID | CP002416 | GBA | PubMed Central | | 1 |
| 51d4a62a067c014cd6ea2340 | 29142822 | 1 | Gc01579 | GcID | CP002416 | GBA | PubMed Central | | 1 |
| 51d4a62a067c014cd6ea2340 | 30202437 | 1 | Gc01579 | GcID | CP002416 | GBA | PubMed Central | | 1 |
| 51d4a62a067c014cd6ea2340 | 30741948 | 1 | Gc01579 | GcID | CP002416 | GBA | PubMed Central | | 1 |
| 51d4a62a067c014cd6ea2340 | 31367231 | 1 | Gc01579 | GcID | CP002416 | GBA | PubMed Central | | 1 |
| 51d4a62a067c014cd6ea2340 | 31890588 | 1 | Gc01579 | GcID | CP002416 | GBA | PubMed Central | | 1 |
| 523b585c067c0139370815a9 | 24179119 | 1 | PRJNA188903 | BPA | PRJNA188903 | BPA | NCBI BioProject Database | | 1 |
| 59cad9a17ded5e2f18691d60 | 27125481 | 1 | PRJNA308023 | BPA | PRJNA308023 | BPA | NCBI BioProject Database | | 1 |
| 5a984c6764d0b326cdd31e58 | 31332186 | 1 | Gp0266600 | GpID | Gp0266600 | GpID | PubMed Central | | 1 |
| 5a984c6764d0b326cdd31e58 | 31332186 | 1 | SAMN09201852 | BSA | SAMN09201852 | BSA | PubMed Central | | 1 |
| 587d27047ded5e4229d8aa30 | 27774985 | 1 | MGYK00000000 | GBA | MGYK00000000 | GBA | NCBI Nucleotide Database | | 1 |
| 523b4d61067c01393707fddd | 25197480 | 1 | Gi11554 | GiID | Gi11554 | GiID | PubMed Central | | 1 |
| 52f3e4c0067c011a2113fd1c | 28832647 | 0 | Gi11069 | GiID | Gi11069 | GiID | PubMed Central | | 1 |
| 523b429a067c01393707f602 | 26380638 | 1 | Gi11346 | GiID | Gp0013740 | GpID | PubMed Central | | 1 |
| 540648ec0d878557fd3be7ef | 30186281 | 1 | SAMN02440801 | BSA | SAMN02440801 | BSA | PubMed Central | | 1 |
| 523b41da067c01393707f45c | 27721915 | 1 | Gi11895 | GiID | Gp0013295 | GpID | PubMed Central | | 1 |
| 52e04cc1067c017f29056847 | 28764658 | 1 | JHWM00000000 | GBA | JHWM00000000 | GBA | PubMed Central | | 1 |
| 558b70860d87854fee5fd571 | 30296267 | 1 | SAMN04487977 | BSA | SAMN04487977 | BSA | PubMed Central | | 1 |
| 58581f837ded5e78cff8e824 | 27774985 | 1 | MFXP00000000 | GBA | MFXP00000000 | GBA | NCBI Nucleotide Database | | 1 |
| 51d5317f067c014cd6efa7d0 | 30923245 | 1 | AUJX00000000 | GBA | AUJX00000000 | GBA | PubMed Central | | 1 |
| 530d650849607a1be0054c26 | 30186281 | 1 | SAMN02745916 | BSA | SAMN02745916 | BSA | PubMed Central | | 1 |
| 58ea58357ded5e52c4b0447d | 30219103 | 1 | Gs0110119 | GsID | Gs0110119 | GsID | PubMed Central | | 1 |
| 57d6ee757ded5e3135ba4f33 | 31608019 | 1 | SAMN05443667 | BSA | SAMN05443667 | BSA | PubMed Central | | 1 |
| 51d4cb53067c014cd6eb0adf | 23239474 | 1 | Gc01740 | GcID | CP002637 | GBA | PubMed Central | | 1 |
| 51d4cb53067c014cd6eb0adf | 23577086 | 1 | Gc01740 | GcID | CP002637 | GBA | PubMed Central | | 1 |
| 51d4cb53067c014cd6eb0adf | 23950883 | 1 | Gc01740 | GcID | CP002637 | GBA | PubMed Central | | 1 |
| 5abf1fcc64d0b30c137085c9 | 31332186 | 1 | Gp0266599 | GpID | Gp0266599 | GpID | PubMed Central | | 1 |
| 5abf1fcc64d0b30c137085c9 | 31332186 | 1 | SAMN10350899 | BSA | SAMN10350899 | BSA | PubMed Central | | 1 |
| 58ccefca7ded5e7a2afbbe64 | 30219103 | 1 | Gs0110119 | GsID | Gs0110119 | GsID | PubMed Central | | 1 |
| 51d53736067c014cd6efefa1 | 31608019 | 1 | SAMN02440754 | BSA | SAMN02440754 | BSA | PubMed Central | | 1 |
| 59122b2c7ded5e1e49ffdd4e | 30219103 | 1 | Gs0110119 | GsID | Gs0110119 | GsID | PubMed Central | | 1 |

| | | | | | | | | | |
|---|---|---|---|---|---|---|---|---|---|
| 53051c9349607a64d3cac728 | 27471578 | 1 | Gi05849 | GiID | Gp0006506 | GpID | PubMed Central | | 1 |
| 51d53185067c014cd6efa827 | 29348922 | 1 | AUJC00000000 | GBA | AUJC00000000 | GBA | PubMed Central | | 1 |
| 588b90677ded5e770a297a24 | 27774985 | 1 | MFQS00000000 | GBA | MFQS00000000 | GBA | NCBI Nucleotide Database | | 1 |
| 5b35710464d0b3516a5eb4ad | 30186281 | 1 | SAMN04489716 | BSA | SAMN04489716 | BSA | PubMed Central | | 1 |
| 51edff15067c014b2c663b35 | 29081911 | 1 | AXVL00000000 | GBA | AXVL00000000 | GBA | PubMed Central | | 1 |
| 52901e3b067c013e2b060c9d | 30993215 | 1 | GCA_000378445 | GBAsA | KB899704 | GBA | PubMed Central | | 2 |
| 52901e3b067c013e2b060c9d | 30993215 | 1 | GCA_000378445 | GBAsA | KB899704 | GBA | PubMed Central | | 2 |
| 52901e3b067c013e2b060c9d | 30993215 | 1 | GCA_000378445 | GBAsA | NZ_KB899704 | GBA | PubMed Central | | 2 |
| 52901e3b067c013e2b060c9d | 31608019 | 1 | ARKH00000000 | GBA | SAMN02440995 | BSA | PubMed Central | | 2 |
| 58010ebd7ded5e3135bc4d29 | 29709896 | 1 | LT670845 | GBA | LT670845 | GBA | PubMed Central | | 2 |
| 52901805067c013e2b060034 | 22624012 | 1 | CP003341 | GBA | CP003341 | GBA | PubMed Central | | 2 |
| 52901805067c013e2b060034 | 24039725 | 1 | CP003341 | GBA | CP003341 | GBA | PubMed Central | | 2 |
| 52901805067c013e2b060034 | 25011617 | 1 | CP003341 | GBA | CP003341 | GBA | PubMed Central | | 2 |
| 52901805067c013e2b060034 | 25084969 | 1 | CP003341 | GBA | CP003341 | GBA | PubMed Central | | 2 |
| 52901805067c013e2b060034 | 25271771 | 1 | CP003341 | GBA | CP003341 | GBA | PubMed Central | | 2 |
| 52901805067c013e2b060034 | 26652857 | 1 | CP003341 | GBA | CP003341 | GBA | PubMed Central | | 2 |
| 52901805067c013e2b060034 | 26866478 | 1 | CP003341 | GBA | CP003341 | GBA | PubMed Central | | 2 |
| 52901805067c013e2b060034 | 27089251 | 1 | CP003341 | GBA | CP003341 | GBA | PubMed Central | | 2 |
| 52901805067c013e2b060034 | 27767913 | 1 | CP003341 | GBA | CP003341 | GBA | PubMed Central | | 2 |
| 52901805067c013e2b060034 | 29047217 | 1 | GCA_000284195 | GBAsA | NC_017044 | GBA | PubMed Central | | 2 |
| 52901805067c013e2b060034 | 29692912 | 1 | GCA_000284195 | GBAsA | NC_017044 | GBA | PubMed Central | | 2 |
| 52901805067c013e2b060034 | 29723287 | 1 | CP003341 | GBA | CP003341 | GBA | PubMed Central | | 2 |
| 52901805067c013e2b060034 | 30882330 | 1 | CP003341 | GBA | CP003341 | GBA | PubMed Central | | 2 |
| 52901805067c013e2b060034 | 31490924 | 1 | CP003341 | GBA | CP003341 | GBA | PubMed Central | | 2 |
| 52901805067c013e2b060034 | 31961304 | 1 | CP003341 | GBA | CP003341 | GBA | PubMed Central | | 2 |
| 52901805067c013e2b060034 | 32697888 | 1 | CP003341 | GBA | CP003341 | GBA | PubMed Central | | 2 |
| 52901805067c013e2b060034 | 32748891 | 1 | CP003341 | GBA | CP003341 | GBA | PubMed Central | | 2 |
| 530d64d049607a1be0054be2 | 25789552 | 1 | CP007055 | GBA | CP007055 | GBA | PubMed Central | | 2 |
| 530d64d049607a1be0054be2 | 26564049 | 1 | CP007056 | GBA | CP007056 | GBA | PubMed Central | | 2 |
| 530d64d049607a1be0054be2 | 26564049 | 1 | CP007057 | GBA | CP007057 | GBA | PubMed Central | | 2 |
| 530d64d049607a1be0054be2 | 26564049 | 1 | CP007058 | GBA | CP007058 | GBA | PubMed Central | | 2 |
| 530d64d049607a1be0054be2 | 26564049 | 1 | CP007059 | GBA | CP007059 | GBA | PubMed Central | | 2 |
| 530d649f49607a1be0054bac | 31509535 | 1 | CP007034 | GBA | CP007034 | GBA | PubMed Central | | 2 |
| 594d609a7ded5e4e5bbd6ade | 21304745 | 1 | 646311962 | IMG_Tax | CP001825 | GBA | NCBI Nucleotide Database | | 2 |
| 594d609a7ded5e4e5bbd6ade | 21304745 | 1 | 646311962 | IMG_Tax | Gc01150 | GcID | PubMed Central | | 2 |
| 594d609a7ded5e4e5bbd6ade | 22768368 | 1 | 646311962 | IMG_Tax | CP001825 | GBA | PubMed Central | | 2 |
| 594d609a7ded5e4e5bbd6ade | 27340512 | 1 | 646311962 | IMG_Tax | CP001825 | GBA | PubMed Central | | 2 |
| 530d68b749607a1be0054ee5 | 26634758 | 1 | 195828 | BPA | 195828 | BPA | NCBI BioProject Database | | 2 |
| 530d68b749607a1be0054ee5 | 26634758 | 1 | JAGA00000000 | GBA | JAGA00000000 | GBA | NCBI Nucleotide Database | | 2 |
| 530d68b749607a1be0054ee5 | 27340512 | 1 | JAGA00000000 | GBA | JAGA00000000 | GBA | PubMed Central | | 2 |
| 530d68b749607a1be0054ee5 | 30186281 | 1 | JAGA00000000 | GBA | SAMN02584935 | BSA | PubMed Central | | 2 |
| 55791c510d878529e7cb084b | 27516518 | 1 | LIPW00000000 | GBA | LIPW00000000 | GBA | PubMed Central | | 2 |
| 55791c510d878529e7cb084b | 30364313 | 1 | LIPW00000000 | GBA | LIPW00000000 | GBA | PubMed Central | | 2 |
| 5513329c0d878525404e829e | 29788499 | 1 | CP010426 | GBA | CP010426 | GBA | PubMed Central | | 2 |
| 5513329c0d878525404e829e | 29884244 | 1 | CP010426 | GBA | CP010426 | GBA | PubMed Central | | 2 |
| 530d63a049607a1be0054a72 | 23432981 | 1 | CP002279 | GBA | CP002279 | GBA | PubMed Central | | 2 |
| 530d63a049607a1be0054a72 | 24019991 | 1 | CP002279 | GBA | CP002279 | GBA | PubMed Central | | 2 |
| 530d63a049607a1be0054a72 | 24906389 | 1 | GCA_000176035 | GBAsA | NC_015675 | GBA | PubMed Central | | 2 |
| 530d63a049607a1be0054a72 | 24976886 | 1 | CP002279 | GBA | CP002279 | GBA | PubMed Central | | 2 |
| 530d63a049607a1be0054a72 | 24976886 | 1 | Gc01853 | GcID | Gc01853 | GcID | PubMed Central | | 2 |
| 530d63a049607a1be0054a72 | 25780499 | 1 | Gc01853 | GcID | Gc01853 | GcID | PubMed Central | | 2 |
| 530d63a049607a1be0054a72 | 25780500 | 1 | Gc01853 | GcID | Gc01853 | GcID | PubMed Central | | 2 |
| 530d63a049607a1be0054a72 | 29220487 | 1 | CP002279 | GBA | CP002279 | GBA | PubMed Central | | 2 |
| 530d651d49607a1be0054c40 | 26203342 | 1 | Gp0010091 | GpID | Gp0010091 | GpID | PubMed Central | | 2 |
| 530d652b49607a1be0054c54 | 30777812 | 1 | ARRT00000000 | GBA | ARRT00000000 | GBA | PubMed Central | | 2 |
| 537cf6b90d87852a04c5a36c | 29126393 | 1 | JHYX00000000 | GBA | JHYX00000000 | GBA | PubMed Central | | 2 |
| 537cf6b90d87852a04c5a36c | 32974501 | 1 | Gp0040000 | GpID | Gp0040000 | GpID | PubMed Central | | 2 |
| 5340e53b49607a0614355d27 | 26172151 | 1 | JHZN00000000 | GBA | JHZN00000000 | GBA | PubMed Central | | 2 |

| | | | | | | | | | |
|---|---|---|---|---|---|---|---|---|---|
| 5340e53b49607a0614355d27 | 31753948 | 1 | GCA_000620965 | GBAsA | GCA_000620965 | GBAsA | PubMed Central | | 2 |
| 53bc8ab50d8785134026e064 | 26913187 | 1 | JQMU00000000 | GBA | JQMU00000000 | GBA | PubMed Central | | 2 |
| 52902b74067c013e2b062942 | 30533813 | 1 | AUFR00000000 | GBA | AUFR00000000 | GBA | PubMed Central | | 2 |
| 52902b74067c013e2b062942 | 31011429 | 1 | AUFR00000000 | GBA | AUFR00000000 | GBA | PubMed Central | | 2 |
| 56362eeb0d87852aea5e6f30 | 32322618 | 1 | Gp0120258 | GpID | 4489800 | BSA | PubMed Central | | 2 |
| 56362eeb0d87852aea5e6f30 | 32322618 | 1 | FNUD00000000 | GBA | SAMN04489800 | BSA | PubMed Central | | 2 |
| 5314a2eb49607a1be005664f | 27833590 | 1 | AUDI00000000 | GBA | AUDI00000000 | GBA | PubMed Central | | 2 |
| 52902808067c013e2b0621fb | 28074122 | 1 | AUFT00000000 | GBA | AUFT00000000 | GBA | PubMed Central | | 2 |
| 52902808067c013e2b0621fb | 28074122 | 1 | Gi11889 | GiID | Gi11889 | GiID | PubMed Central | | 2 |
| 5938bbc87ded5e4e5bbbeffd | 31380022 | 1 | Gp0211493 | GpID | SRX3939697 | SRA | PubMed Central | | 2 |
| 5702184a7ded5e7f7b944d9f | 30459722 | 1 | FMCP00000000 | GBA | SAMN04883176 | BSA | PubMed Central | | 2 |
| 535290280d87855a8277bc67 | 31215393 | 1 | JMLM00000000 | GBA | JMLM00000000 | GBA | PubMed Central | | 2 |
| 55132fc60d878525404e72db | 15059260 | 1 | GCA_000011465 | GBAsA | NC_005072 | GBA | PubMed Central | | 2 |
| 55132fc60d878525404e72db | 15693943 | 1 | GCA_000011465 | GBAsA | NC_005072 | GBA | PubMed Central | | 2 |
| 55132fc60d878525404e72db | 15828858 | 1 | BX548174 | GBA | BX548174 | GBA | PubMed Central | | 2 |
| 55132fc60d878525404e72db | 17941988 | 1 | GCA_000011465 | GBAsA | NC_005072 | GBA | PubMed Central | | 2 |
| 55132fc60d878525404e72db | 18045455 | 1 | BX548174 | GBA | BX548174 | GBA | PubMed Central | | 2 |
| 55132fc60d878525404e72db | 18203741 | 1 | GCA_000011465 | GBAsA | NC_005072 | GBA | PubMed Central | | 2 |
| 55132fc60d878525404e72db | 18507822 | 1 | BX548174 | GBA | BX548174 | GBA | PubMed Central | | 2 |
| 55132fc60d878525404e72db | 18534010 | 1 | BX548174 | GBA | BX548174 | GBA | PubMed Central | | 2 |
| 55132fc60d878525404e72db | 18769676 | 1 | BX548174 | GBA | BX548174 | GBA | PubMed Central | | 2 |
| 55132fc60d878525404e72db | 19850757 | 1 | BX548174 | GBA | BX548174 | GBA | PubMed Central | | 2 |
| 55132fc60d878525404e72db | 20345942 | 1 | BX548174 | GBA | BX548174 | GBA | PubMed Central | | 2 |
| 55132fc60d878525404e72db | 21255351 | 1 | Gc00152 | GcID | Gc00152 | GcID | PubMed Central | | 2 |
| 55132fc60d878525404e72db | 21799937 | 1 | GCA_000011465 | GBAsA | NC_005072 | GBA | PubMed Central | | 2 |
| 55132fc60d878525404e72db | 22768379 | 1 | BX548174 | GBA | BX548174 | GBA | PubMed Central | | 2 |
| 55132fc60d878525404e72db | 22894826 | 1 | BX548174 | GBA | BX548174 | GBA | PubMed Central | | 2 |
| 55132fc60d878525404e72db | 22941652 | 1 | BX548174 | GBA | BX548174 | GBA | PubMed Central | | 2 |
| 55132fc60d878525404e72db | 23185592 | 1 | GCA_000011465 | GBAsA | NC_005072 | GBA | PubMed Central | | 2 |
| 55132fc60d878525404e72db | 24438106 | 1 | GCA_000011465 | GBAsA | NC_005072 | GBA | PubMed Central | | 2 |
| 55132fc60d878525404e72db | 24822028 | 1 | GCA_000011465 | GBAsA | NC_005072 | GBA | PubMed Central | | 2 |
| 55132fc60d878525404e72db | 25977791 | 1 | BX548174 | GBA | BX548174 | GBA | PubMed Central | | 2 |
| 55132fc60d878525404e72db | 26244890 | 1 | BX548174 | GBA | BX548174 | GBA | PubMed Central | | 2 |
| 55132fc60d878525404e72db | 27703668 | 1 | GCA_000011465 | GBAsA | GCA_000011465 | GBAsA | PubMed Central | | 2 |
| 55132fc60d878525404e72db | 27788196 | 1 | BX548174 | GBA | BX548174 | GBA | PubMed Central | | 2 |
| 55132fc60d878525404e72db | 27868089 | 1 | BX548174 | GBA | BX548174 | GBA | PubMed Central | | 2 |
| 55132fc60d878525404e72db | 29915114 | 1 | BX548174 | GBA | BX548174 | GBA | PubMed Central | | 2 |
| 55132fc60d878525404e72db | 30134835 | 1 | GCA_000011465 | GBAsA | NC_005072 | GBA | PubMed Central | | 2 |
| 55132fc60d878525404e72db | 30609821 | 1 | GCA_000011465 | GBAsA | NC_005072 | GBA | PubMed Central | | 2 |
| 55132fc60d878525404e72db | 32346084 | 1 | BX548174 | GBA | BX548174 | GBA | PubMed Central | | 2 |
| 530d650249607a1be0054c1f | 25780499 | 1 | AZAM00000000 | GBA | AZAM00000000 | GBA | PubMed Central | | 2 |
| 530d650249607a1be0054c1f | 25780499 | 1 | Gi08825 | GiID | Gi08825 | GiID | PubMed Central | | 2 |
| 5314a25f49607a1be00565d3 | 26568374 | 1 | AQVC00000000 | GBA | AQVC00000000 | GBA | PubMed Central | | 2 |
| 5726d57c7ded5e4fbeb8283f | 31506266 | 1 | GCA_900111835 | GBAsA | GCF_900111835 | RSAsA | PubMed Central | | 2 |
| 5726d57c7ded5e4fbeb8283f | 31506266 | 1 | GCA_900111835 | GBAsA | GCF_900111835 | RSAsA | PubMed Central | | 2 |
| 5726d57c7ded5e4fbeb8283f | 32299875 | 1 | FOJP00000000 | GBA | FOJP00000000 | GBA | PubMed Central | | 2 |
| 5726d57c7ded5e4fbeb8283f | 32657018 | 1 | FOJP00000000 | GBA | FOJP00000000 | GBA | PubMed Central | | 2 |
| 5bcb97c046d1e64cf8dadc28 | 31619684 | 1 | Gp0323843 | GpID | SRP185225 | SRA | PubMed Central | | 2 |
| 5af1795664d0b3374773f264 | 27590813 | 1 | Gp0057369 | GpID | SRX2014391 | SRA | PubMed Central | | 2 |
| 5af1795664d0b3374773f264 | 31474963 | 1 | Gs0085736 | GsID | Gs0085736 | GsID | PubMed Central | | 2 |
| 5290433f067c013e2b066087 | 24874669 | 1 | 218809 | BPA | 218809 | BPA | NCBI BioProject Database | | 2 |
| 5290433f067c013e2b066087 | 24874669 | 1 | JDWH00000000 | GBA | JDWH00000000 | GBA | NCBI Nucleotide Database | | 2 |
| 52901eb8067c013e2b060dd5 | 17504473 | 1 | CP000606 | GBA | CP000606 | GBA | PubMed Central | | 2 |
| 52901eb8067c013e2b060dd5 | 18534010 | 1 | GCA_000016065 | GBAsA | NC_009092 | GBA | PubMed Central | | 2 |
| 52901eb8067c013e2b060dd5 | 19758436 | 1 | CP000606 | GBA | CP000606 | GBA | PubMed Central | | 2 |
| 52901eb8067c013e2b060dd5 | 21261910 | 1 | GCA_000016065 | GBAsA | NC_009092 | GBA | PubMed Central | | 2 |
| 52901eb8067c013e2b060dd5 | 21569439 | 1 | GCA_000016065 | GBAsA | NC_009092 | GBA | PubMed Central | | 2 |
| 52901eb8067c013e2b060dd5 | 23110226 | 1 | CP000606 | GBA | CP000606 | GBA | PubMed Central | | 2 |

| | | | | | | | | |
|---|---|---|---|---|---|---|---|---|
| 52901eb8067c013e2b060dd5 | 23516532 | 1 | GCA_000016065 | GBAsA | NC_009092 | GBA | PubMed Central | 2 |
| 52901eb8067c013e2b060dd5 | 25768732 | 1 | CP000606 | GBA | CP000606 | GBA | PubMed Central | 2 |
| 52901eb8067c013e2b060dd5 | 27206730 | 1 | CP000606 | GBA | CP000606 | GBA | PubMed Central | 2 |
| 52901eb8067c013e2b060dd5 | 30367596 | 1 | GCA_000016065 | GBAsA | NC_009092 | GBA | PubMed Central | 2 |
| 533c5e8e49607a0614353b33 | 30186281 | 1 | JHWQ00000000 | GBA | SAMN02743880 | BSA | PubMed Central | 2 |
| 530d646049607a1be0054b64 | 28293682 | 1 | CP003153 | GBA | CP003153 | GBA | PubMed Central | 2 |
| 530d63e849607a1be0054acd | 27379049 | 1 | CP003029 | GBA | CP003029 | GBA | PubMed Central | 2 |
| 530d63e849607a1be0054acd | 27471502 | 1 | CP003029 | GBA | CP003029 | GBA | PubMed Central | 2 |
| 530d63e849607a1be0054acd | 30105201 | 1 | CP003029 | GBA | CP003029 | GBA | PubMed Central | 2 |
| 530d63dd49607a1be0054abe | 21886865 | 1 | 48579 | BPA | 48579 | BPA | NCBI BioProject Database | 2 |
| 530d63dd49607a1be0054abe | 21886865 | 1 | CP002838 | GBA | CP002838 | GBA | NCBI Nucleotide Database | 2 |
| 530d63dd49607a1be0054abe | 25789552 | 1 | GCA_000223395 | GBAsA | NC_015931 | GBA | PubMed Central | 2 |
| 530d659749607a1be0054cdd | 26251504 | 1 | 157399 | BPA | 157399 | BPA | NCBI BioProject Database | 2 |
| 530d659749607a1be0054cdd | 26251504 | 1 | LANG00000000 | GBA | LANG00000000 | GBA | NCBI Nucleotide Database | 2 |
| 530d659749607a1be0054cdd | 31333602 | 1 | LANG00000000 | GBA | LANG00000000 | GBA | PubMed Central | 2 |
| 5314a28749607a1be00565f9 | 25914684 | 1 | ARJT00000000 | GBA | ARJT00000000 | GBA | PubMed Central | 2 |
| 5314a28749607a1be00565f9 | 30147685 | 1 | GCA_000378185 | GBAsA | KB899391 | GBA | PubMed Central | 2 |
| 5314a28749607a1be00565f9 | 30147685 | 1 | GCA_000378185 | GBAsA | KB899391 | GBA | PubMed Central | 2 |
| 5314a28749607a1be00565f9 | 30147685 | 1 | GCA_000378185 | GBAsA | NZ_KB899391 | GBA | PubMed Central | 2 |
| 573f749b7ded5e779637fb77 | 29692909 | 1 | FNDX00000000 | GBA | FNDX00000000 | GBA | PubMed Central | 2 |
| 530d63ee49607a1be0054ad4 | 23458837 | 1 | 52545 | BPA | 52545 | BPA | NCBI BioProject Database | 2 |
| 530d63ee49607a1be0054ad4 | 23458837 | 1 | CP002771 | GBA | CP002771 | GBA | NCBI Nucleotide Database | 2 |
| 530d63ee49607a1be0054ad4 | 28989277 | 1 | GCA_000214215 | GBAsA | NC_015559 | GBA | PubMed Central | 2 |
| 56362f3f0d87852aea5e72b2 | 30894631 | 1 | LT629699 | GBA | LT629699 | GBA | PubMed Central | 2 |
| 529029e6067c013e2b0625f6 | 24363793 | 0 | Gi21295 | GiID | Gi21295 | GiID | PubMed Central | 2 |
| 55e07de50d878556782d8fad | 26379647 | 1 | Gp0087948 | GpID | Gp0087948 | GpID | PubMed Central | 2 |
| 52901b6c067c013e2b0606cd | 26039074 | 1 | ASWY00000000 | GBA | ASWY00000000 | GBA | PubMed Central | 2 |
| 5290253a067c013e2b061bc0 | 28764658 | 1 | ATVB00000000 | GBA | ATVB00000000 | GBA | PubMed Central | 2 |
| 5290253a067c013e2b061bc0 | 30186281 | 1 | ATVB00000000 | GBA | SAMN02441248 | BSA | PubMed Central | 2 |
| 537d25820d87852a04c5b200 | 23577086 | 1 | AAWL00000000 | GBA | AAWL00000000 | GBA | PubMed Central | 2 |
| 5314a28749607a1be00565fa | 28070336 | 1 | ARJO00000000 | GBA | ARJO00000000 | GBA | PubMed Central | 2 |
| 52901d73067c013e2b060ae4 | 28184216 | 1 | ARJN00000000 | GBA | ARJN00000000 | GBA | PubMed Central | 2 |
| 52cb210e067c0120babfc21c | 31749830 | 1 | Gp0056908 | GpID | SRR3989263 | SRA | PubMed Central | 2 |
| 545aa66d0d878552848903a1 | 28491240 | 1 | Gp0103631 | GpID | Gp0103631 | GpID | PubMed Central | 2 |
| 545aa66d0d878552848903a1 | 32974515 | 1 | GCA_900108115 | GBAsA | FTPU01000000 | GBA | PubMed Central | 2 |
| 594d60937ded5e4e5bbd6ad8 | 21304676 | 1 | 646311953 | IMG_Tax | CP001823 | GBA | NCBI Nucleotide Database | 2 |
| 594d60937ded5e4e5bbd6ad8 | 22768368 | 1 | 646311953 | IMG_Tax | CP001823 | GBA | PubMed Central | 2 |
| 594d60937ded5e4e5bbd6ad8 | 30186281 | 1 | 646311953 | IMG_Tax | SAMN02598446 | BSA | PubMed Central | 2 |
| 559c853b0d87852b215059d1 | 28935744 | 1 | FRAU00000000 | GBA | FRAU00000000 | GBA | PubMed Central | 2 |
| 559c853b0d87852b215059d1 | 31608019 | 1 | Gp0113017 | GpID | 4488087 | BSA | PubMed Central | 2 |
| 559c853b0d87852b215059d1 | 31608019 | 1 | FRAU00000000 | GBA | SAMN04488087 | BSA | PubMed Central | 2 |
| 52902786067c013e2b0620e1 | 26478786 | 1 | AUFE00000000 | GBA | AUFE00000000 | GBA | PubMed Central | 2 |
| 52902786067c013e2b0620e1 | 26478786 | 1 | Gp0009812 | GpID | Gp0009812 | GpID | PubMed Central | 2 |
| 52901d77067c013e2b060aec | 24003073 | 1 | ARKK00000000 | GBA | ARKK00000000 | GBA | PubMed Central | 2 |
| 52901d77067c013e2b060aec | 25678942 | 1 | ARKK00000000 | GBA | ARKK00000000 | GBA | PubMed Central | 2 |
| 52901d77067c013e2b060aec | 25678942 | 1 | Gi11553 | GiID | Gi11553 | GiID | PubMed Central | 2 |
| 5314a27c49607a1be00565ed | 27257493 | 1 | ARGO00000000 | GBA | ARGO00000000 | GBA | PubMed Central | 2 |
| 5314a27c49607a1be00565ed | 28912952 | 1 | ARGO00000000 | GBA | ARGO00000000 | GBA | PubMed Central | 2 |
| 56abd2900d878559e286f3ae | 31551998 | 1 | Gp0111916 | GpID | SRP117915 | SRA | PubMed Central | 2 |
| 594d60b07ded5e4e5bbd6af7 | 21357773 | 1 | 646564587 | IMG_Tax | CP001966 | GBA | PubMed Central | 2 |
| 594d60b07ded5e4e5bbd6af7 | 21886861 | 1 | 646564587 | IMG_Tax | CP001966 | GBA | NCBI Nucleotide Database | 2 |
| 594d60b07ded5e4e5bbd6af7 | 21886861 | 1 | 646564587 | IMG_Tax | Gc01341 | GcID | PubMed Central | 2 |
| 594d60b07ded5e4e5bbd6af7 | 24266988 | 1 | 646564587 | IMG_Tax | CP001966 | GBA | PubMed Central | 2 |
| 594d60b07ded5e4e5bbd6af7 | 30186281 | 1 | 646564587 | IMG_Tax | SAMN00002597 | BSA | PubMed Central | 2 |
| 594d60b07ded5e4e5bbd6af7 | 31456790 | 1 | 646564587 | IMG_Tax | CP001966 | GBA | PubMed Central | 2 |
| 594d609c7ded5e4e5bbd6ae0 | 21304678 | 1 | 646311965 | IMG_Tax | CP001820 | GBA | NCBI Nucleotide Database | 2 |
| 594d609c7ded5e4e5bbd6ae0 | 21304678 | 1 | 646311965 | IMG_Tax | Gc01152 | GcID | PubMed Central | 2 |
| 594d609c7ded5e4e5bbd6ae0 | 23577086 | 1 | 646311965 | IMG_Tax | CP001820 | GBA | PubMed Central | 2 |

| | | | | | | | | |
|---|---|---|---|---|---|---|---|---|
| 594d609c7ded5e4e5bbd6ae0 | 24466089 | 1 | 646311965 | IMG_Tax | CP001820 | GBA | PubMed Central | 2 |
| 594d609c7ded5e4e5bbd6ae0 | 28713344 | 1 | 646311965 | IMG_Tax | CP001820 | GBA | PubMed Central | 2 |
| 594d609c7ded5e4e5bbd6ae0 | 31509535 | 1 | 646311965 | IMG_Tax | CP001820 | GBA | PubMed Central | 2 |
| 530d650d49607a1be0054c2c | 25780497 | 1 | ATTL00000000 | GBA | ATTL00000000 | GBA | PubMed Central | 2 |
| 530d650d49607a1be0054c2c | 25780497 | 1 | Gi08905 | GiID | Gi08905 | GiID | PubMed Central | 2 |
| 569c14480d8785737066d8b5 | 23468637 | 1 | GCA_000166115 | GBAsA | NC_017310 | GBA | PubMed Central | 2 |
| 569c14480d8785737066d8b5 | 23577209 | 1 | CP002297 | GBA | CP002297 | GBA | PubMed Central | 2 |
| 569c14480d8785737066d8b5 | 32461582 | 1 | CP002297 | GBA | CP002297 | GBA | PubMed Central | 2 |
| 529018c4067c013e2b060210 | 25225493 | 1 | ATZE00000000 | GBA | ATZE00000000 | GBA | PubMed Central | 2 |
| 529018c4067c013e2b060210 | 27721915 | 1 | ATZE00000000 | GBA | ATZE00000000 | GBA | PubMed Central | 2 |
| 529018c4067c013e2b060210 | 27721915 | 1 | Gp0013295 | GpID | Gp0013295 | GpID | PubMed Central | 2 |
| 55c388220d87855594c278db | 29765358 | 1 | Gp0106908 | GpID | 4487782 | BSA | PubMed Central | 2 |
| 55c388220d87855594c278db | 29765358 | 1 | FOZF00000000 | GBA | SAMN04487782 | BSA | PubMed Central | 2 |
| 530d674b49607a1be0054e08 | 26205857 | 1 | 89155 | BPA | 89155 | BPA | NCBI BioProject Database | 2 |
| 530d674b49607a1be0054e08 | 26205857 | 1 | ASAA00000000 | GBA | ASAA00000000 | GBA | NCBI Nucleotide Database | 2 |
| 551331c60d878525404e7d24 | 19533120 | 1 | GCA_000012425 | GBAsA | NC_007298 | GBA | PubMed Central | 2 |
| 551331c60d878525404e7d24 | 21750120 | 1 | CP000089 | GBA | CP000089 | GBA | PubMed Central | 2 |
| 551331c60d878525404e7d24 | 23825601 | 1 | CP000089 | GBA | CP000089 | GBA | PubMed Central | 2 |
| 551331c60d878525404e7d24 | 24475028 | 1 | GCA_000012425 | GBAsA | NC_007298 | GBA | PubMed Central | 2 |
| 551331c60d878525404e7d24 | 31275250 | 1 | CP000089 | GBA | CP000089 | GBA | PubMed Central | 2 |
| 551331c60d878525404e7d24 | 31562384 | 1 | GCA_000012425 | GBAsA | NC_007298 | GBA | PubMed Central | 2 |
| 551331c60d878525404e7d24 | 31922045 | 1 | GCA_000012425 | GBAsA | NC_007298 | GBA | PubMed Central | 2 |
| 594d60887ded5e4e5bbd6acc | 21304663 | 1 | 644736405 | IMG_Tax | CP001684 | GBA | NCBI Nucleotide Database | 2 |
| 594d60887ded5e4e5bbd6acc | 21304663 | 1 | 644736405 | IMG_Tax | Gc01094 | GcID | PubMed Central | 2 |
| 594d60887ded5e4e5bbd6acc | 28326685 | 1 | 644736405 | IMG_Tax | CP001684 | GBA | PubMed Central | 2 |
| 594d60887ded5e4e5bbd6acc | 30186281 | 1 | 644736405 | IMG_Tax | SAMN02598438 | BSA | PubMed Central | 2 |
| 59a156907ded5e41edd859a6 | 31743595 | 1 | PZZW00000000 | GBA | PZZW00000000 | GBA | PubMed Central | 2 |
| 53bc8abe0d8785134026e071 | 28337181 | 1 | JQMX00000000 | GBA | JQMX00000000 | GBA | PubMed Central | 2 |
| 530d651b49607a1be0054c3e | 26664655 | 1 | 165339 | BPA | 165339 | BPA | NCBI BioProject Database | 2 |
| 530d651b49607a1be0054c3e | 26664655 | 1 | ATTR00000000 | GBA | ATTR00000000 | GBA | NCBI Nucleotide Database | 2 |
| 529019eb067c013e2b0604cb | 26659677 | 1 | ARBV00000000 | GBA | ARBV00000000 | GBA | PubMed Central | 2 |
| 529019eb067c013e2b0604cb | 28442708 | 1 | ARBV00000000 | GBA | ARBV00000000 | GBA | PubMed Central | 2 |
| 529019eb067c013e2b0604cb | 32038594 | 1 | GCA_000379885 | GBAsA | KB903969 | GBA | PubMed Central | 2 |
| 529019eb067c013e2b0604cb | 32038594 | 1 | GCA_000379885 | GBAsA | KB903969 | GBA | PubMed Central | 2 |
| 529019eb067c013e2b0604cb | 32038594 | 1 | GCA_000379885 | GBAsA | KB903995 | GBA | PubMed Central | 2 |
| 529019eb067c013e2b0604cb | 32038594 | 1 | GCA_000379885 | GBAsA | KB903995 | GBA | PubMed Central | 2 |
| 529019eb067c013e2b0604cb | 32038594 | 1 | GCA_000379885 | GBAsA | KB904006 | GBA | PubMed Central | 2 |
| 529019eb067c013e2b0604cb | 32038594 | 1 | GCA_000379885 | GBAsA | KB904006 | GBA | PubMed Central | 2 |
| 529019eb067c013e2b0604cb | 32038594 | 1 | GCA_000379885 | GBAsA | KB904038 | GBA | PubMed Central | 2 |
| 529019eb067c013e2b0604cb | 32038594 | 1 | GCA_000379885 | GBAsA | NZ_KB903969 | GBA | PubMed Central | 2 |
| 529019eb067c013e2b0604cb | 32038594 | 1 | GCA_000379885 | GBAsA | NZ_KB903995 | GBA | PubMed Central | 2 |
| 529019eb067c013e2b0604cb | 32038594 | 1 | GCA_000379885 | GBAsA | NZ_KB904006 | GBA | PubMed Central | 2 |
| 5621c9240d878540fd707442 | 30186281 | 1 | Gp0117059 | GpID | 4490356 | BSA | PubMed Central | 2 |
| 5621c9240d878540fd707442 | 30186281 | 1 | FNST00000000 | GBA | SAMN04490356 | BSA | PubMed Central | 2 |
| 529015da067c013e2b05fb39 | 27151933 | 1 | AZXR00000000 | GBA | AZXR00000000 | GBA | PubMed Central | 2 |
| 530d64a149607a1be0054bb0 | 25780503 | 1 | CP003364 | GBA | CP003364 | GBA | PubMed Central | 2 |
| 530d64a149607a1be0054bb0 | 25780503 | 1 | CP003367 | GBA | CP003367 | GBA | PubMed Central | 2 |
| 530d64a149607a1be0054bb0 | 28360896 | 1 | CP003364 | GBA | CP003364 | GBA | PubMed Central | 2 |
| 530d64a149607a1be0054bb0 | 28360896 | 1 | CP003365 | GBA | CP003365 | GBA | PubMed Central | 2 |
| 530d64a149607a1be0054bb0 | 32399714 | 1 | CP003364 | GBA | CP003364 | GBA | PubMed Central | 2 |
| 530d64a149607a1be0054bb0 | 32399714 | 1 | CP003367 | GBA | CP003367 | GBA | PubMed Central | 2 |
| 530d64a149607a1be0054bb0 | 33414774 | 1 | CP003364 | GBA | CP003364 | GBA | PubMed Central | 2 |
| 52901c1b067c013e2b0607d7 | 26478785 | 1 | ARCY00000000 | GBA | ARCY00000000 | GBA | PubMed Central | 2 |
| 52901c1b067c013e2b0607d7 | 26478785 | 1 | Gi08873 | GiID | Gi08873 | GiID | PubMed Central | 2 |
| 594d60a07ded5e4e5bbd6ae5 | 22180816 | 1 | 646564511 | IMG_Tax | CP002017 | GBA | NCBI Nucleotide Database | 2 |
| 594d60a07ded5e4e5bbd6ae5 | 22180816 | 1 | 646564511 | IMG_Tax | Gc01268 | GcID | PubMed Central | 2 |
| 5ad0e8de64d0b33747707fad | 30732610 | 1 | Ga0236286 | GaID | Ga0236286 | GaID | PubMed Central | 2 |
| 555518d30d8785178e712cb4 | 27516514 | 1 | LIOL00000000 | GBA | LIOL00000000 | GBA | PubMed Central | 2 |

| | | | | | | | | | |
|---|---|---|---|---|---|---|---|---|---|
| 555518d30d8785178e712cb4 | 30497377 | 1 | Ga0061061 | GaID | Ga0061061 | GaID | PubMed Central | | 2 |
| 5314a27b49607a1be00565eb | 28649989 | 1 | GCA_000374805 | GBAsA | KB895358 | GBA | PubMed Central | | 2 |
| 5314a27b49607a1be00565eb | 33193169 | 1 | ARGN00000000 | GBA | ARGN00000000 | GBA | PubMed Central | | 2 |
| 529016ec067c013e2b05fdaa | 24286338 | 1 | AQUH00000000 | GBA | AQUH00000000 | GBA | PubMed Central | | 2 |
| 529016ec067c013e2b05fdaa | 28116058 | 0 | Gi21891 | GiID | Gi21891 | GiID | PubMed Central | | 2 |
| 529016ec067c013e2b05fdaa | 30443603 | 1 | AQUH00000000 | GBA | AQUH00000000 | GBA | PubMed Central | | 2 |
| 530d64c749607a1be0054bd9 | 25610435 | 1 | CP007033 | GBA | CP007033 | GBA | PubMed Central | | 2 |
| 530d64c749607a1be0054bd9 | 26903979 | 1 | CP007033 | GBA | CP007033 | GBA | PubMed Central | | 2 |
| 530d64c749607a1be0054bd9 | 27641516 | 1 | CP007033 | GBA | CP007033 | GBA | PubMed Central | | 2 |
| 52901bfd067c013e2b060790 | 23035691 | 1 | 68459 | BPA | 68459 | BPA | NCBI BioProject Database | | 2 |
| 52901bfd067c013e2b060790 | 23035691 | 1 | AFYK00000000 | GBA | AFYK00000000 | GBA | NCBI Nucleotide Database | | 2 |
| 52901bfd067c013e2b060790 | 30386310 | 1 | AFYK00000000 | GBA | AFYK00000000 | GBA | PubMed Central | | 2 |
| 530d647e49607a1be0054b88 | 28116241 | 1 | GCA_000235605 | GBAsA | NC_016584 | GBA | PubMed Central | | 2 |
| 530d647e49607a1be0054b88 | 29106396 | 1 | CP003108 | GBA | CP003108 | GBA | PubMed Central | | 2 |
| 530d640e49607a1be0054afe | 29618738 | 1 | CM001437 | GBA | CM001437 | GBA | PubMed Central | | 2 |
| 533c5e9149607a0614353b36 | 30210264 | 1 | JHWX00000000 | GBA | JHWX00000000 | GBA | PubMed Central | | 2 |
| 57d6f1027ded5e3135ba5293 | 29437089 | 1 | FNLG00000000 | GBA | FNLG00000000 | GBA | PubMed Central | | 2 |
| 5314a26b49607a1be00565dd | 26639610 | 1 | AREW00000000 | GBA | AREW00000000 | GBA | PubMed Central | | 2 |
| 53920c660d878575e64354e2 | 26074887 | 1 | JQKF00000000 | GBA | JQKF00000000 | GBA | PubMed Central | | 2 |
| 53a2651d0d878514c2d0f169 | 27491994 | 1 | JPOO00000000 | GBA | JPOO00000000 | GBA | PubMed Central | | 2 |
| 53a2651d0d878514c2d0f169 | 30643889 | 1 | JPOO00000000 | GBA | JPOO00000000 | GBA | PubMed Central | | 2 |
| 535290a10d87855a8277bd91 | 27127589 | 1 | CP007181 | GBA | CP007181 | GBA | PubMed Central | | 3 |
| 535290a10d87855a8277bd91 | 28790203 | 1 | CP007182 | GBA | CP007182 | GBA | PubMed Central | | 3 |
| 535290a10d87855a8277bd91 | 30230632 | 1 | CP007181 | GBA | CP007181 | GBA | PubMed Central | | 3 |
| 535290a10d87855a8277bd91 | 30258424 | 1 | CP007182 | GBA | CP007182 | GBA | PubMed Central | | 3 |
| 535290a10d87855a8277bd91 | 30914053 | 1 | CP007181 | GBA | CP007181 | GBA | PubMed Central | | 3 |
| 535290a10d87855a8277bd91 | 32111893 | 1 | CP007181 | GBA | CP007181 | GBA | PubMed Central | | 3 |
| 535290a10d87855a8277bd91 | 32152417 | 1 | CP007181 | GBA | CP007181 | GBA | PubMed Central | | 3 |
| 535290a10d87855a8277bd91 | 32532247 | 1 | CP007181 | GBA | CP007181 | GBA | PubMed Central | | 3 |
| 5ad0ea0864d0b33747708267 | 29312831 | 1 | CP022695 | GBA | SAMN07452765 | BSA | PubMed Central | | 3 |
| 5ad0ea0864d0b33747708267 | 29312831 | 1 | CP022695 | GBA | SAMN07452765 | BSA | PubMed Central | | 3 |
| 5ad0ea0864d0b33747708267 | 29312831 | 1 | CP022696 | GBA | SAMN07452765 | BSA | PubMed Central | | 3 |
| 5ad0ea0864d0b33747708267 | 29312831 | 1 | CP022696 | GBA | SAMN07452765 | BSA | PubMed Central | | 3 |
| 5ad0ea0864d0b33747708267 | 29312831 | 1 | CP022697 | GBA | SAMN07452765 | BSA | PubMed Central | | 3 |
| 5ad0ea0864d0b33747708267 | 29312831 | 1 | CP022697 | GBA | SAMN07452765 | BSA | PubMed Central | | 3 |
| 5ad0ea0864d0b33747708267 | 29312831 | 1 | CP022698 | GBA | SAMN07452765 | BSA | PubMed Central | | 3 |
| 5ad0ea0864d0b33747708267 | 29312831 | 1 | CP022698 | GBA | SAMN07452765 | BSA | PubMed Central | | 3 |
| 5ad0ea0864d0b33747708267 | 30782635 | 1 | CP022695 | GBA | CP022695 | GBA | PubMed Central | | 3 |
| 5ad0ea0864d0b33747708267 | 32763937 | 1 | CP022695 | GBA | CP022695 | GBA | PubMed Central | | 3 |
| 5af649bc64d0b3374774c1d5 | 32076091 | 1 | GCA_002165345 | GBAsA | CP033133 | GBA | PubMed Central | | 3 |
| 5af649bc64d0b3374774c1d5 | 32273730 | 1 | GCA_002165345 | GBAsA | CP033122 | GBA | PubMed Central | | 3 |
| 5af649bc64d0b3374774c1d5 | 33042094 | 1 | GCA_002165345 | GBAsA | CP033131 | GBA | PubMed Central | | 3 |
| 5af649bc64d0b3374774c1d5 | 33343549 | 1 | GCA_002165345 | GBAsA | CP033130 | GBA | PubMed Central | | 3 |
| 535290d00d87855a8277be0e | 31358803 | 1 | AB088224 | GBA | AB088224 | GBA | PubMed Central | | 3 |
| 545aa65f0d8785528489020f | 24963920 | 1 | NC_017981 | GBA | NC_017981 | GBA | PubMed Central | | 3 |
| 52901752067c013e2b05fea5 | 22026465 | 1 | 65435 | BPA | 65435 | BPA | NCBI BioProject Database | | 3 |
| 52901752067c013e2b05fea5 | 22026465 | 1 | CP002641 | GBA | CP002641 | GBA | NCBI Nucleotide Database | | 3 |
| 52901752067c013e2b05fea5 | 25161960 | 1 | GCA_000231885 | GBAsA | NC_017620 | GBA | PubMed Central | | 3 |
| 52901752067c013e2b05fea5 | 25620959 | 1 | CP002641 | GBA | CP002641 | GBA | PubMed Central | | 3 |
| 52901752067c013e2b05fea5 | 26870017 | 1 | CP002641 | GBA | CP002641 | GBA | PubMed Central | | 3 |
| 52901752067c013e2b05fea5 | 27270714 | 1 | GCA_000231885 | GBAsA | NC_017620 | GBA | PubMed Central | | 3 |
| 52901752067c013e2b05fea5 | 28198678 | 1 | CP002641 | GBA | CP002641 | GBA | PubMed Central | | 3 |
| 52901752067c013e2b05fea5 | 30186253 | 1 | GCA_000231885 | GBAsA | NC_017620 | GBA | PubMed Central | | 3 |
| 5791b7817ded5e31abaa1e27 | 26358608 | 1 | 278367 | BPA | 278367 | BPA | NCBI BioProject Database | | 3 |
| 5791b7817ded5e31abaa1e27 | 26358608 | 1 | CP012027 | GBA | CP012027 | GBA | NCBI Nucleotide Database | | 3 |
| 5791b7817ded5e31abaa1e27 | 26358608 | 1 | CP012027 | GBA | CP012027 | GBA | NCBI Nucleotide Database | | 3 |
| 5791b7817ded5e31abaa1e27 | 28348872 | 1 | CP012027 | GBA | CP012027 | GBA | PubMed Central | | 3 |
| 5791b7817ded5e31abaa1e27 | 30775379 | 1 | CP012027 | GBA | CP012027 | GBA | PubMed Central | | 3 |

| | | | | | | | | |
|---|---|---|---|---|---|---|---|---|
| 5791b7817ded5e31abaa1e27 | 31293838 | 1 | CP012027 | GBA | CP012027 | GBA | PubMed Central | 3 |
| 5791b7817ded5e31abaa1e27 | 31293838 | 1 | CP012027 | GBA | CP012027 | GBA | PubMed Central | 3 |
| 5791b7817ded5e31abaa1e27 | 31358980 | 1 | CP012027 | GBA | CP012027 | GBA | PubMed Central | 3 |
| 5791b7817ded5e31abaa1e27 | 32461582 | 1 | CP012027 | GBA | CP012027 | GBA | PubMed Central | 3 |
| 5791aee87ded5e31abaa12ef | 27932655 | 1 | CP015005 | GBA | CP015005 | GBA | PubMed Central | 3 |
| 5791aee87ded5e31abaa12ef | 27932655 | 1 | CP015006 | GBA | CP015006 | GBA | PubMed Central | 3 |
| 5791aee87ded5e31abaa12ef | 27932655 | 1 | CP015007 | GBA | CP015007 | GBA | PubMed Central | 3 |
| 5791aee87ded5e31abaa12ef | 27932655 | 1 | CP015008 | GBA | CP015008 | GBA | PubMed Central | 3 |
| 5791aee87ded5e31abaa12ef | 27932655 | 1 | CP015009 | GBA | CP015009 | GBA | PubMed Central | 3 |
| 573f7be37ded5e7796380528 | 28330888 | 1 | 245882 | BPA | 245882 | BPA | NCBI BioProject Database | 3 |
| 573f7be37ded5e7796380528 | 31139608 | 1 | JNUM00000000 | GBA | SAMN02741394 | BSA | PubMed Central | 3 |
| 58d413727ded5e7a2afc1175 | 19190628 | 0 | CP010150 | GBA | CP010150 | GBA | PubMed Central | 3 |
| 58d413727ded5e7a2afc1175 | 20551958 | 0 | CP010150 | GBA | CP010150 | GBA | PubMed Central | 3 |
| 58d413727ded5e7a2afc1175 | 22413839 | 0 | CP010150 | GBA | CP010150 | GBA | PubMed Central | 3 |
| 58d413727ded5e7a2afc1175 | 28803888 | 0 | CP010150 | GBA | CP010150 | GBA | PubMed Central | 3 |
| 58d413727ded5e7a2afc1175 | 29980701 | 1 | CP010150 | GBA | CP010150 | GBA | PubMed Central | 3 |
| 58d413727ded5e7a2afc1175 | 32420029 | 1 | CP010150 | GBA | CP010150 | GBA | PubMed Central | 3 |
| 5913849a7ded5e1e49ffea7a | 27979954 | 1 | LNCM00000000 | GBA | LNCM00000000 | GBA | PubMed Central | 3 |
| 5621c95a0d878540fd7076d1 | 26537657 | 1 | 258321 | BPA | 258321 | BPA | NCBI BioProject Database | 3 |
| 5621c95a0d878540fd7076d1 | 26537657 | 1 | CP011325 | GBA | CP011325 | GBA | NCBI Nucleotide Database | 3 |
| 5621c95a0d878540fd7076d1 | 26537657 | 1 | CP011325 | GBA | CP011325 | GBA | NCBI Nucleotide Database | 3 |
| 5621c95a0d878540fd7076d1 | 30055586 | 1 | CP011325 | GBA | CP011325 | GBA | PubMed Central | 3 |
| 5621c95a0d878540fd7076d1 | 31449567 | 1 | CP011325 | GBA | CP011325 | GBA | PubMed Central | 3 |
| 5621c95a0d878540fd7076d1 | 32423070 | 1 | CP011325 | GBA | CP011325 | GBA | PubMed Central | 3 |
| 59138ea57ded5e1e49fff93b | 27965637 | 1 | CP017006 | GBA | CP017006 | GBA | PubMed Central | 3 |
| 59138ea57ded5e1e49fff93b | 27965637 | 1 | CP017006 | GBA | CP017006 | GBA | PubMed Central | 3 |
| 592475e67ded5e4e5bbaee8d | 26941156 | 1 | 177212 | BPA | 177212 | BPA | NCBI BioProject Database | 3 |
| 592475e67ded5e4e5bbaee8d | 26941156 | 1 | LHSU00000000 | GBA | LHSU00000000 | GBA | NCBI Nucleotide Database | 3 |
| 592475e67ded5e4e5bbaee8d | 30533757 | 1 | GCA_001448665 | GBAsA | CP030288 | GBA | PubMed Central | 3 |
| 592475e67ded5e4e5bbaee8d | 30533757 | 1 | GCA_001448665 | GBAsA | CP030289 | GBA | PubMed Central | 3 |
| 592475e67ded5e4e5bbaee8d | 30533757 | 1 | GCA_001448665 | GBAsA | CP030290 | GBA | PubMed Central | 3 |
| 592475e67ded5e4e5bbaee8d | 30533757 | 1 | GCA_001448665 | GBAsA | CP030291 | GBA | PubMed Central | 3 |
| 5e96fa0976b4483f14ad6e92 | 30886188 | 1 | QDGB00000000 | GBA | QDGB00000000 | GBA | PubMed Central | 3 |
| 52904140067c013e2b065c17 | 23825472 | 1 | 50373 | BPA | 50373 | BPA | NCBI BioProject Database | 3 |
| 52904140067c013e2b065c17 | 23825472 | 1 | AEEC00000000 | GBA | AEEC00000000 | GBA | NCBI Nucleotide Database | 3 |
| 56af96620d878559e28716c2 | 28298905 | 1 | CP013690 | GBA | CP013690 | GBA | PubMed Central | 3 |
| 56af96620d878559e28716c2 | 29312264 | 1 | CP013690 | GBA | CP013690 | GBA | PubMed Central | 3 |
| 56af96620d878559e28716c2 | 29312264 | 1 | CP013690 | GBA | CP013690 | GBA | PubMed Central | 3 |
| 56af96620d878559e28716c2 | 29797432 | 1 | CP013690 | GBA | CP013690 | GBA | PubMed Central | 3 |
| 56af96620d878559e28716c2 | 29797432 | 1 | CP013691 | GBA | CP013691 | GBA | PubMed Central | 3 |
| 56af96620d878559e28716c2 | 32464640 | 1 | CP013691 | GBA | CP013691 | GBA | PubMed Central | 3 |
| 5e27aac37776dfea0bd9d085 | 28558786 | 1 | CP019755 | GBA | SAMN06324250 | BSA | PubMed Central | 3 |
| 5e27aac37776dfea0bd9d085 | 28558786 | 1 | CP019755 | GBA | SAMN06324250 | BSA | PubMed Central | 3 |
| 5e27aac37776dfea0bd9d085 | 28558786 | 1 | CP019756 | GBA | SAMN06324250 | BSA | PubMed Central | 3 |
| 5e27aac37776dfea0bd9d085 | 28558786 | 1 | CP019756 | GBA | SAMN06324250 | BSA | PubMed Central | 3 |
| 5e27aac37776dfea0bd9d085 | 28558786 | 1 | CP019757 | GBA | SAMN06324250 | BSA | PubMed Central | 3 |
| 5e27aac37776dfea0bd9d085 | 28558786 | 1 | CP019757 | GBA | SAMN06324250 | BSA | PubMed Central | 3 |
| 5e27aac37776dfea0bd9d085 | 28558786 | 1 | CP019758 | GBA | SAMN06324250 | BSA | PubMed Central | 3 |
| 5e27aac37776dfea0bd9d085 | 28558786 | 1 | CP019758 | GBA | SAMN06324250 | BSA | PubMed Central | 3 |
| 5e27aac37776dfea0bd9d085 | 28558786 | 1 | CP019759 | GBA | SAMN06324250 | BSA | PubMed Central | 3 |
| 5e27aac37776dfea0bd9d085 | 28558786 | 1 | CP019759 | GBA | SAMN06324250 | BSA | PubMed Central | 3 |
| 5e27aac37776dfea0bd9d085 | 28558786 | 1 | CP019760 | GBA | SAMN06324250 | BSA | PubMed Central | 3 |
| 5e27aac37776dfea0bd9d085 | 28558786 | 1 | CP019760 | GBA | SAMN06324250 | BSA | PubMed Central | 3 |
| 5e27aac37776dfea0bd9d085 | 28558786 | 1 | CP019761 | GBA | SAMN06324250 | BSA | PubMed Central | 3 |
| 5e27aac37776dfea0bd9d085 | 28558786 | 1 | CP019761 | GBA | SAMN06324250 | BSA | PubMed Central | 3 |
| 5e27aac37776dfea0bd9d085 | 28558786 | 1 | CP019762 | GBA | SAMN06324250 | BSA | PubMed Central | 3 |
| 5e27aac37776dfea0bd9d085 | 28558786 | 1 | CP019762 | GBA | SAMN06324250 | BSA | PubMed Central | 3 |
| 5e27aac37776dfea0bd9d085 | 28558786 | 1 | CP019763 | GBA | SAMN06324250 | BSA | PubMed Central | 3 |

| | | | | | | | | |
|---|---|---|---|---|---|---|---|---|
| 5e27aac37776dfea0bd9d085 | 28558786 | 1 | CP019763 | GBA | SAMN06324250 | BSA | PubMed Central | 3 |
| 5e27aac37776dfea0bd9d085 | 28558786 | 1 | CP019764 | GBA | SAMN06324250 | BSA | PubMed Central | 3 |
| 5e27aac37776dfea0bd9d085 | 28558786 | 1 | CP019764 | GBA | SAMN06324250 | BSA | PubMed Central | 3 |
| 5e27aac37776dfea0bd9d085 | 28558786 | 1 | CP019765 | GBA | SAMN06324250 | BSA | PubMed Central | 3 |
| 5e27aac37776dfea0bd9d085 | 28558786 | 1 | CP019765 | GBA | SAMN06324250 | BSA | PubMed Central | 3 |
| 5e27aac37776dfea0bd9d085 | 28558786 | 1 | CP019766 | GBA | SAMN06324250 | BSA | PubMed Central | 3 |
| 5e27aac37776dfea0bd9d085 | 28558786 | 1 | CP019766 | GBA | SAMN06324250 | BSA | PubMed Central | 3 |
| 5e27aac37776dfea0bd9d085 | 28558786 | 1 | CP019767 | GBA | SAMN06324250 | BSA | PubMed Central | 3 |
| 5e27aac37776dfea0bd9d085 | 28558786 | 1 | CP019767 | GBA | SAMN06324250 | BSA | PubMed Central | 3 |
| 5e27aac37776dfea0bd9d085 | 28705983 | 1 | CP019755 | GBA | CP019755 | GBA | PubMed Central | 3 |
| 5e27aac37776dfea0bd9d085 | 28705983 | 1 | CP019756 | GBA | CP019756 | GBA | PubMed Central | 3 |
| 5e27aac37776dfea0bd9d085 | 28705983 | 1 | CP019757 | GBA | CP019757 | GBA | PubMed Central | 3 |
| 5e27aac37776dfea0bd9d085 | 28705983 | 1 | CP019758 | GBA | CP019758 | GBA | PubMed Central | 3 |
| 5e27aac37776dfea0bd9d085 | 28705983 | 1 | CP019759 | GBA | CP019759 | GBA | PubMed Central | 3 |
| 5e27aac37776dfea0bd9d085 | 28705983 | 1 | CP019760 | GBA | CP019760 | GBA | PubMed Central | 3 |
| 5e27aac37776dfea0bd9d085 | 28705983 | 1 | CP019761 | GBA | CP019761 | GBA | PubMed Central | 3 |
| 5e27aac37776dfea0bd9d085 | 28705983 | 1 | CP019762 | GBA | CP019762 | GBA | PubMed Central | 3 |
| 5e27aac37776dfea0bd9d085 | 28705983 | 1 | CP019763 | GBA | CP019763 | GBA | PubMed Central | 3 |
| 5e27aac37776dfea0bd9d085 | 28705983 | 1 | CP019764 | GBA | CP019764 | GBA | PubMed Central | 3 |
| 5e27aac37776dfea0bd9d085 | 28705983 | 1 | CP019765 | GBA | CP019765 | GBA | PubMed Central | 3 |
| 5e27aac37776dfea0bd9d085 | 28705983 | 1 | CP019766 | GBA | CP019766 | GBA | PubMed Central | 3 |
| 5e27aac37776dfea0bd9d085 | 28705983 | 1 | CP019767 | GBA | CP019767 | GBA | PubMed Central | 3 |
| 5e27aac37776dfea0bd9d085 | 30986208 | 1 | CP019767 | GBA | CP019767 | GBA | PubMed Central | 3 |
| 5e27aac37776dfea0bd9d085 | 32941463 | 1 | CP019767 | GBA | CP019767 | GBA | PubMed Central | 3 |
| 5e27aac37776dfea0bd9d085 | 33092135 | 1 | CP019767 | GBA | CP019767 | GBA | PubMed Central | 3 |
| 5bd0f21c46d1e64cf8db3f10 | 29167267 | 1 | CP024160 | GBA | CP024160 | GBA | PubMed Central | 3 |
| 52903f70067c013e2b0657de | 23133391 | 1 | 159625 | BPA | 159625 | BPA | NCBI BioProject Database | 3 |
| 52903f70067c013e2b0657de | 23133391 | 1 | AKVS00000000 | GBA | AKVS00000000 | GBA | NCBI Nucleotide Database | 3 |
| 52903f70067c013e2b0657de | 23781227 | 1 | AKVS00000000 | GBA | AKVS00000000 | GBA | PubMed Central | 3 |
| 57be094b7ded5e0c87136aa3 | 26868384 | 1 | BCMS00000000 | GBA | BCMS00000000 | GBA | NCBI Nucleotide Database | 3 |
| 598bf25c7ded5e41edd726ad | 27417830 | 1 | 254816 | BPA | 254816 | BPA | NCBI BioProject Database | 3 |
| 598bf25c7ded5e41edd726ad | 27587814 | 1 | JQYK00000000 | GBA | JQYK00000000 | GBA | PubMed Central | 3 |
| 5aedadb664d0b33747739460 | 28074866 | 1 | FAXD00000000 | GBA | FAXD00000000 | GBA | PubMed Central | 3 |
| 56af92290d878559e2870dd8 | 26151935 | 1 | 244049 | BPA | 244049 | BPA | NCBI BioProject Database | 3 |
| 56af92290d878559e2870dd8 | 26151935 | 1 | JQAO00000000 | GBA | JQAO00000000 | GBA | NCBI Nucleotide Database | 3 |
| 56af92290d878559e2870dd8 | 28611758 | 1 | JQAO00000000 | GBA | JQAO00000000 | GBA | PubMed Central | 3 |
| 56af92290d878559e2870dd8 | 29997608 | 1 | JQAO00000000 | GBA | JQAO00000000 | GBA | PubMed Central | 3 |
| 56af92290d878559e2870dd8 | 30533939 | 1 | GCA_001029495 | GBAsA | GCA_001029495 | GBAsA | PubMed Central | 3 |
| 59138fd17ded5e1e49fffaab | 27198026 | 1 | LUAU00000000 | GBA | LUAU00000000 | GBA | PubMed Central | 3 |
| 5e96f69776b4483f14ad4886 | 29880910 | 1 | CP027845 | GBA | CP027845 | GBA | PubMed Central | 3 |
| 538020d80d87852a04c5bec3 | 24155888 | 1 | 191808 | BPA | 191808 | BPA | NCBI BioProject Database | 3 |
| 538020d80d87852a04c5bec3 | 24155888 | 1 | APJG00000000 | GBA | APJG00000000 | GBA | NCBI Nucleotide Database | 3 |
| 5621c96d0d878540fd707843 | 25858846 | 1 | CP010905 | GBA | CP010905 | GBA | NCBI Nucleotide Database | 3 |
| 5621c96d0d878540fd707843 | 25858846 | 1 | CP010905 | GBA | CP010905 | GBA | NCBI Nucleotide Database | 3 |
| 5621c96d0d878540fd707843 | 27733845 | 1 | CP010905 | GBA | CP010905 | GBA | PubMed Central | 3 |
| 5621c96d0d878540fd707843 | 28262711 | 1 | CP010905 | GBA | CP010905 | GBA | PubMed Central | 3 |
| 5621c96d0d878540fd707843 | 29291715 | 1 | CP010905 | GBA | CP010905 | GBA | PubMed Central | 3 |
| 5621c96d0d878540fd707843 | 29761637 | 1 | CP010905 | GBA | CP010905 | GBA | PubMed Central | 3 |
| 5621c96d0d878540fd707843 | 30224751 | 1 | CP010905 | GBA | CP010905 | GBA | PubMed Central | 3 |
| 5621c96d0d878540fd707843 | 31148548 | 1 | CP010905 | GBA | CP010905 | GBA | PubMed Central | 3 |
| 5621c96d0d878540fd707843 | 31509535 | 1 | CP010905 | GBA | CP010905 | GBA | PubMed Central | 3 |
| 5621c96d0d878540fd707843 | 31552001 | 1 | CP010905 | GBA | CP010905 | GBA | PubMed Central | 3 |
| 5e16320de08d44553ef6cd41 | 31395630 | 1 | CP039642 | GBA | CP039642 | GBA | PubMed Central | 3 |
| 5e16320de08d44553ef6cd41 | 31395630 | 1 | CP039643 | GBA | CP039643 | GBA | PubMed Central | 3 |
| 5e16320de08d44553ef6cd41 | 31395630 | 1 | CP039644 | GBA | CP039644 | GBA | PubMed Central | 3 |
| 5e16320de08d44553ef6cd41 | 31395630 | 1 | CP039645 | GBA | CP039645 | GBA | PubMed Central | 3 |
| 5e16320de08d44553ef6cd41 | 31395630 | 1 | CP039646 | GBA | CP039646 | GBA | PubMed Central | 3 |
| 5e16320de08d44553ef6cd41 | 31395630 | 1 | CP039647 | GBA | CP039647 | GBA | PubMed Central | 3 |

| | | | | | | | | |
|---|---|---|---|---|---|---|---|---|
| 5e16320de08d44553ef6cd41 | 31395630 | 1 | CP039648 | GBA | CP039648 | GBA | PubMed Central | 3 |
| 5e16320de08d44553ef6cd41 | 31395630 | 1 | CP039649 | GBA | CP039649 | GBA | PubMed Central | 3 |
| 5e16320de08d44553ef6cd41 | 31395630 | 1 | CP039650 | GBA | CP039650 | GBA | PubMed Central | 3 |
| 5e16320de08d44553ef6cd41 | 31395630 | 1 | CP039651 | GBA | CP039651 | GBA | PubMed Central | 3 |
| 573f76f17ded5e779637fea7 | 31300709 | 1 | CP009025 | GBA | CP009025 | GBA | PubMed Central | 3 |
| 573f76f17ded5e779637fea7 | 31500575 | 1 | CP009025 | GBA | CP009025 | GBA | PubMed Central | 3 |
| 545aa6270d8785528488fe07 | 24278249 | 1 | NC_019517 | GBA | NC_019517 | GBA | PubMed Central | 3 |
| 545aa6270d8785528488fe07 | 25629728 | 1 | NC_019517 | GBA | NC_019517 | GBA | PubMed Central | 3 |
| 545aa6270d8785528488fe07 | 27280590 | 1 | NC_019517 | GBA | NC_019517 | GBA | PubMed Central | 3 |
| 5844917c7ded5e2d305ca9ff | 27138938 | 1 | NC_030115 | GBA | NC_030115 | GBA | NCBI Nucleotide Database | 3 |
| 58d414ad7ded5e7a2afc1332 | 29631621 | 1 | CP017650 | GBA | CP017650 | GBA | PubMed Central | 3 |
| 58d414ad7ded5e7a2afc1332 | 31194814 | 1 | CP017650 | GBA | CP017650 | GBA | PubMed Central | 3 |
| 58d40b197ded5e7a2afc056a | 29382871 | 1 | CP018914 | GBA | CP018914 | GBA | PubMed Central | 3 |
| 58d40b197ded5e7a2afc056a | 29692912 | 1 | CP018914 | GBA | CP018914 | GBA | PubMed Central | 3 |
| 58d40b197ded5e7a2afc056a | 29692912 | 1 | CP018914 | GBA | CP018914 | GBA | PubMed Central | 3 |
| 58d40b197ded5e7a2afc056a | 30475889 | 1 | CP018914 | GBA | CP018914 | GBA | PubMed Central | 3 |
| 58d40b197ded5e7a2afc056a | 30909949 | 1 | CP018914 | GBA | CP018914 | GBA | PubMed Central | 3 |
| 58d40b197ded5e7a2afc056a | 33126449 | 1 | CP018914 | GBA | CP018914 | GBA | PubMed Central | 3 |
| 570215f87ded5e7f7b944ab0 | 28767122 | 1 | AJPW00000000 | GBA | SAMN02469464 | BSA | PubMed Central | 3 |
| 575cfb337ded5e5e73213be4 | 27198010 | 1 | CP014352 | GBA | CP014352 | GBA | PubMed Central | 3 |
| 575cfb337ded5e5e73213be4 | 27198010 | 1 | CP014353 | GBA | CP014353 | GBA | PubMed Central | 3 |
| 52901722067c013e2b05fe2d | 23408961 | 1 | CP002914 | GBA | CP002914 | GBA | PubMed Central | 3 |
| 52901722067c013e2b05fe2d | 23665771 | 1 | GCA_000225915 | GBAsA | NC_016010 | GBA | PubMed Central | 3 |
| 52901722067c013e2b05fe2d | 24274055 | 1 | GCA_000225915 | GBAsA | NC_016010 | GBA | PubMed Central | 3 |
| 52901722067c013e2b05fe2d | 24278159 | 1 | CP002914 | GBA | CP002914 | GBA | PubMed Central | 3 |
| 52901722067c013e2b05fe2d | 24416331 | 1 | CP002914 | GBA | CP002914 | GBA | PubMed Central | 3 |
| 52901722067c013e2b05fe2d | 24897119 | 1 | CP002914 | GBA | CP002914 | GBA | PubMed Central | 3 |
| 52901722067c013e2b05fe2d | 26673755 | 1 | GCA_000225915 | GBAsA | NC_016010 | GBA | PubMed Central | 3 |
| 52901722067c013e2b05fe2d | 30140181 | 1 | CP002914 | GBA | CP002914 | GBA | PubMed Central | 3 |
| 52901722067c013e2b05fe2d | 31164878 | 1 | CP002914 | GBA | CP002914 | GBA | PubMed Central | 3 |
| 52901722067c013e2b05fe2d | 31623235 | 1 | GCA_000225915 | GBAsA | NC_016010 | GBA | PubMed Central | 3 |
| 560c3c4b0d878540fd6fcc3f | 25999572 | 1 | 277476 | BPA | 277476 | BPA | NCBI BioProject Database | 3 |
| 560c3c4b0d878540fd6fcc3f | 25999572 | 1 | JZJL00000000 | GBA | JZJL00000000 | GBA | NCBI Nucleotide Database | 3 |
| 58ebfa437ded5e52c4b053e3 | 27284141 | 1 | 311246 | BPA | 311246 | BPA | NCBI BioProject Database | 3 |
| 58ebfa437ded5e52c4b053e3 | 27284141 | 1 | CP014349 | GBA | CP014349 | GBA | NCBI Nucleotide Database | 3 |
| 58ebfa437ded5e52c4b053e3 | 27284141 | 1 | CP014349 | GBA | CP014349 | GBA | NCBI Nucleotide Database | 3 |
| 58ebfa437ded5e52c4b053e3 | 27284141 | 1 | CP014350 | GBA | CP014350 | GBA | NCBI Nucleotide Database | 3 |
| 58ebfa437ded5e52c4b053e3 | 27284141 | 1 | CP014350 | GBA | CP014350 | GBA | NCBI Nucleotide Database | 3 |
| 58ebfa437ded5e52c4b053e3 | 27284141 | 1 | CP014351 | GBA | CP014351 | GBA | NCBI Nucleotide Database | 3 |
| 58ebfa437ded5e52c4b053e3 | 27284141 | 1 | CP014351 | GBA | CP014351 | GBA | NCBI Nucleotide Database | 3 |
| 58ebfa437ded5e52c4b053e3 | 27284141 | 1 | CP014792 | GBA | CP014792 | GBA | NCBI Nucleotide Database | 3 |
| 58ebfa437ded5e52c4b053e3 | 27284141 | 1 | CP014792 | GBA | CP014792 | GBA | NCBI Nucleotide Database | 3 |
| 58ebfa437ded5e52c4b053e3 | 27284141 | 1 | CP014871 | GBA | CP014871 | GBA | NCBI Nucleotide Database | 3 |
| 58ebfa437ded5e52c4b053e3 | 27284141 | 1 | CP014871 | GBA | CP014871 | GBA | NCBI Nucleotide Database | 3 |
| 58ebfa437ded5e52c4b053e3 | 27284141 | 1 | CP015331 | GBA | CP015331 | GBA | NCBI Nucleotide Database | 3 |
| 58ebfa437ded5e52c4b053e3 | 27284141 | 1 | CP015331 | GBA | CP015331 | GBA | NCBI Nucleotide Database | 3 |
| 58ebfa437ded5e52c4b053e3 | 27284141 | 1 | CP015332 | GBA | CP015332 | GBA | NCBI Nucleotide Database | 3 |
| 58ebfa437ded5e52c4b053e3 | 27284141 | 1 | CP015332 | GBA | CP015332 | GBA | NCBI Nucleotide Database | 3 |
| 58ebfa437ded5e52c4b053e3 | 27284141 | 1 | CP015333 | GBA | CP015333 | GBA | NCBI Nucleotide Database | 3 |
| 58ebfa437ded5e52c4b053e3 | 27284141 | 1 | CP015333 | GBA | CP015333 | GBA | NCBI Nucleotide Database | 3 |
| 58ebfa437ded5e52c4b053e3 | 27284141 | 1 | CP015334 | GBA | CP015334 | GBA | NCBI Nucleotide Database | 3 |
| 58ebfa437ded5e52c4b053e3 | 27284141 | 1 | CP015334 | GBA | CP015334 | GBA | NCBI Nucleotide Database | 3 |
| 58ebfa437ded5e52c4b053e3 | 27284141 | 1 | CP015335 | GBA | CP015335 | GBA | NCBI Nucleotide Database | 3 |
| 58ebfa437ded5e52c4b053e3 | 27284141 | 1 | CP015335 | GBA | CP015335 | GBA | NCBI Nucleotide Database | 3 |
| 58ebfa437ded5e52c4b053e3 | 27284141 | 1 | CP015336 | GBA | CP015336 | GBA | NCBI Nucleotide Database | 3 |
| 58ebfa437ded5e52c4b053e3 | 27284141 | 1 | CP015336 | GBA | CP015336 | GBA | NCBI Nucleotide Database | 3 |
| 58ebfa437ded5e52c4b053e3 | 28302772 | 1 | CP014349 | GBA | SAMN04481062 | BSA | PubMed Central | 3 |
| 58ebfa437ded5e52c4b053e3 | 28302772 | 1 | CP014349 | GBA | SAMN04481062 | BSA | PubMed Central | 3 |

| | | | | | | | | |
|---|---|---|---|---|---|---|---|---|
| 58ebfa437ded5e52c4b053e3 | 28302772 | 1 | CP014350 | GBA | SAMN04481062 | BSA | PubMed Central | 3 |
| 58ebfa437ded5e52c4b053e3 | 28302772 | 1 | CP014350 | GBA | SAMN04481062 | BSA | PubMed Central | 3 |
| 58ebfa437ded5e52c4b053e3 | 28302772 | 1 | CP014351 | GBA | SAMN04481062 | BSA | PubMed Central | 3 |
| 58ebfa437ded5e52c4b053e3 | 28302772 | 1 | CP014351 | GBA | SAMN04481062 | BSA | PubMed Central | 3 |
| 58ebfa437ded5e52c4b053e3 | 28302772 | 1 | CP014792 | GBA | SAMN04481062 | BSA | PubMed Central | 3 |
| 58ebfa437ded5e52c4b053e3 | 28302772 | 1 | CP014792 | GBA | SAMN04481062 | BSA | PubMed Central | 3 |
| 58ebfa437ded5e52c4b053e3 | 28302772 | 1 | CP014871 | GBA | SAMN04481062 | BSA | PubMed Central | 3 |
| 58ebfa437ded5e52c4b053e3 | 28302772 | 1 | CP014871 | GBA | SAMN04481062 | BSA | PubMed Central | 3 |
| 58ebfa437ded5e52c4b053e3 | 28302772 | 1 | CP015331 | GBA | SAMN04481062 | BSA | PubMed Central | 3 |
| 58ebfa437ded5e52c4b053e3 | 28302772 | 1 | CP015331 | GBA | SAMN04481062 | BSA | PubMed Central | 3 |
| 58ebfa437ded5e52c4b053e3 | 28302772 | 1 | CP015332 | GBA | SAMN04481062 | BSA | PubMed Central | 3 |
| 58ebfa437ded5e52c4b053e3 | 28302772 | 1 | CP015332 | GBA | SAMN04481062 | BSA | PubMed Central | 3 |
| 58ebfa437ded5e52c4b053e3 | 28302772 | 1 | CP015333 | GBA | SAMN04481062 | BSA | PubMed Central | 3 |
| 58ebfa437ded5e52c4b053e3 | 28302772 | 1 | CP015333 | GBA | SAMN04481062 | BSA | PubMed Central | 3 |
| 58ebfa437ded5e52c4b053e3 | 28302772 | 1 | CP015334 | GBA | SAMN04481062 | BSA | PubMed Central | 3 |
| 58ebfa437ded5e52c4b053e3 | 28302772 | 1 | CP015334 | GBA | SAMN04481062 | BSA | PubMed Central | 3 |
| 58ebfa437ded5e52c4b053e3 | 28302772 | 1 | CP015335 | GBA | SAMN04481062 | BSA | PubMed Central | 3 |
| 58ebfa437ded5e52c4b053e3 | 28302772 | 1 | CP015335 | GBA | SAMN04481062 | BSA | PubMed Central | 3 |
| 58ebfa437ded5e52c4b053e3 | 28302772 | 1 | CP015336 | GBA | SAMN04481062 | BSA | PubMed Central | 3 |
| 58ebfa437ded5e52c4b053e3 | 28302772 | 1 | CP015336 | GBA | SAMN04481062 | BSA | PubMed Central | 3 |
| 58ebfa437ded5e52c4b053e3 | 28656008 | 1 | CP014349 | GBA | SAMN04481062 | BSA | PubMed Central | 3 |
| 58ebfa437ded5e52c4b053e3 | 28656008 | 1 | CP014349 | GBA | SAMN04481062 | BSA | PubMed Central | 3 |
| 58ebfa437ded5e52c4b053e3 | 28656008 | 1 | CP014350 | GBA | SAMN04481062 | BSA | PubMed Central | 3 |
| 58ebfa437ded5e52c4b053e3 | 28656008 | 1 | CP014350 | GBA | SAMN04481062 | BSA | PubMed Central | 3 |
| 58ebfa437ded5e52c4b053e3 | 28656008 | 1 | CP014351 | GBA | SAMN04481062 | BSA | PubMed Central | 3 |
| 58ebfa437ded5e52c4b053e3 | 28656008 | 1 | CP014351 | GBA | SAMN04481062 | BSA | PubMed Central | 3 |
| 58ebfa437ded5e52c4b053e3 | 28656008 | 1 | CP014792 | GBA | SAMN04481062 | BSA | PubMed Central | 3 |
| 58ebfa437ded5e52c4b053e3 | 28656008 | 1 | CP014792 | GBA | SAMN04481062 | BSA | PubMed Central | 3 |
| 58ebfa437ded5e52c4b053e3 | 28656008 | 1 | CP014871 | GBA | SAMN04481062 | BSA | PubMed Central | 3 |
| 58ebfa437ded5e52c4b053e3 | 28656008 | 1 | CP014871 | GBA | SAMN04481062 | BSA | PubMed Central | 3 |
| 58ebfa437ded5e52c4b053e3 | 28656008 | 1 | CP015331 | GBA | SAMN04481062 | BSA | PubMed Central | 3 |
| 58ebfa437ded5e52c4b053e3 | 28656008 | 1 | CP015331 | GBA | SAMN04481062 | BSA | PubMed Central | 3 |
| 58ebfa437ded5e52c4b053e3 | 28656008 | 1 | CP015332 | GBA | SAMN04481062 | BSA | PubMed Central | 3 |
| 58ebfa437ded5e52c4b053e3 | 28656008 | 1 | CP015332 | GBA | SAMN04481062 | BSA | PubMed Central | 3 |
| 58ebfa437ded5e52c4b053e3 | 28656008 | 1 | CP015333 | GBA | SAMN04481062 | BSA | PubMed Central | 3 |
| 58ebfa437ded5e52c4b053e3 | 28656008 | 1 | CP015333 | GBA | SAMN04481062 | BSA | PubMed Central | 3 |
| 58ebfa437ded5e52c4b053e3 | 28656008 | 1 | CP015334 | GBA | SAMN04481062 | BSA | PubMed Central | 3 |
| 58ebfa437ded5e52c4b053e3 | 28656008 | 1 | CP015334 | GBA | SAMN04481062 | BSA | PubMed Central | 3 |
| 58ebfa437ded5e52c4b053e3 | 28656008 | 1 | CP015335 | GBA | SAMN04481062 | BSA | PubMed Central | 3 |
| 58ebfa437ded5e52c4b053e3 | 28656008 | 1 | CP015335 | GBA | SAMN04481062 | BSA | PubMed Central | 3 |
| 58ebfa437ded5e52c4b053e3 | 28656008 | 1 | CP015336 | GBA | SAMN04481062 | BSA | PubMed Central | 3 |
| 58ebfa437ded5e52c4b053e3 | 28656008 | 1 | CP015336 | GBA | SAMN04481062 | BSA | PubMed Central | 3 |
| 58ebfa437ded5e52c4b053e3 | 29056727 | 1 | CP014349 | GBA | SAMN04481062 | BSA | PubMed Central | 3 |
| 58ebfa437ded5e52c4b053e3 | 29056727 | 1 | CP014349 | GBA | SAMN04481062 | BSA | PubMed Central | 3 |
| 58ebfa437ded5e52c4b053e3 | 29056727 | 1 | CP014350 | GBA | SAMN04481062 | BSA | PubMed Central | 3 |
| 58ebfa437ded5e52c4b053e3 | 29056727 | 1 | CP014350 | GBA | SAMN04481062 | BSA | PubMed Central | 3 |
| 58ebfa437ded5e52c4b053e3 | 29056727 | 1 | CP014351 | GBA | SAMN04481062 | BSA | PubMed Central | 3 |
| 58ebfa437ded5e52c4b053e3 | 29056727 | 1 | CP014351 | GBA | SAMN04481062 | BSA | PubMed Central | 3 |
| 58ebfa437ded5e52c4b053e3 | 29056727 | 1 | CP014792 | GBA | SAMN04481062 | BSA | PubMed Central | 3 |
| 58ebfa437ded5e52c4b053e3 | 29056727 | 1 | CP014792 | GBA | SAMN04481062 | BSA | PubMed Central | 3 |
| 58ebfa437ded5e52c4b053e3 | 29056727 | 1 | CP014871 | GBA | SAMN04481062 | BSA | PubMed Central | 3 |
| 58ebfa437ded5e52c4b053e3 | 29056727 | 1 | CP014871 | GBA | SAMN04481062 | BSA | PubMed Central | 3 |
| 58ebfa437ded5e52c4b053e3 | 29056727 | 1 | CP015331 | GBA | SAMN04481062 | BSA | PubMed Central | 3 |
| 58ebfa437ded5e52c4b053e3 | 29056727 | 1 | CP015331 | GBA | SAMN04481062 | BSA | PubMed Central | 3 |
| 58ebfa437ded5e52c4b053e3 | 29056727 | 1 | CP015332 | GBA | SAMN04481062 | BSA | PubMed Central | 3 |
| 58ebfa437ded5e52c4b053e3 | 29056727 | 1 | CP015332 | GBA | SAMN04481062 | BSA | PubMed Central | 3 |
| 58ebfa437ded5e52c4b053e3 | 29056727 | 1 | CP015333 | GBA | SAMN04481062 | BSA | PubMed Central | 3 |
| 58ebfa437ded5e52c4b053e3 | 29056727 | 1 | CP015333 | GBA | SAMN04481062 | BSA | PubMed Central | 3 |

| | | | | | | | | |
|---|---|---|---|---|---|---|---|---|
| 58ebfa437ded5e52c4b053e3 | 29056727 | 1 | CP015334 | GBA | SAMN04481062 | BSA | PubMed Central | 3 |
| 58ebfa437ded5e52c4b053e3 | 29056727 | 1 | CP015334 | GBA | SAMN04481062 | BSA | PubMed Central | 3 |
| 58ebfa437ded5e52c4b053e3 | 29056727 | 1 | CP015335 | GBA | SAMN04481062 | BSA | PubMed Central | 3 |
| 58ebfa437ded5e52c4b053e3 | 29056727 | 1 | CP015335 | GBA | SAMN04481062 | BSA | PubMed Central | 3 |
| 58ebfa437ded5e52c4b053e3 | 29056727 | 1 | CP015336 | GBA | SAMN04481062 | BSA | PubMed Central | 3 |
| 58ebfa437ded5e52c4b053e3 | 29056727 | 1 | CP015336 | GBA | SAMN04481062 | BSA | PubMed Central | 3 |
| 58ebfa437ded5e52c4b053e3 | 30586413 | 1 | CP014349 | GBA | CP014349 | GBA | PubMed Central | 3 |
| 58ebfa437ded5e52c4b053e3 | 30586413 | 1 | CP014349 | GBA | CP014349 | GBA | PubMed Central | 3 |
| 545aa6630d878552848902a1 | 24309727 | 1 | NC_022770 | GBA | NC_022770 | GBA | NCBI Nucleotide Database | 3 |
| 545aa6630d878552848902a1 | 25280881 | 1 | NC_022770 | GBA | NC_022770 | GBA | PubMed Central | 3 |
| 545aa6630d878552848902a1 | 25547158 | 1 | NC_022770 | GBA | NC_022770 | GBA | PubMed Central | 3 |
| 545aa6630d878552848902a1 | 25635028 | 1 | NC_022770 | GBA | NC_022770 | GBA | PubMed Central | 3 |
| 545aa6630d878552848902a1 | 25635030 | 1 | NC_022770 | GBA | NC_022770 | GBA | PubMed Central | 3 |
| 545aa6630d878552848902a1 | 25953175 | 1 | NC_022770 | GBA | NC_022770 | GBA | PubMed Central | 3 |
| 545aa6630d878552848902a1 | 26337888 | 1 | NC_022770 | GBA | NC_022770 | GBA | PubMed Central | 3 |
| 52904034067c013e2b06599e | 23766401 | 1 | 164747 | BPA | 164747 | BPA | NCBI BioProject Database | 3 |
| 52904034067c013e2b06599e | 23766401 | 1 | AORI00000000 | GBA | AORI00000000 | GBA | NCBI Nucleotide Database | 3 |
| 5a3732d47ded5e35e94f6ced | 29472326 | 1 | CP014862 | GBA | CP014862 | GBA | PubMed Central | 3 |
| 5a3732d47ded5e35e94f6ced | 29472326 | 1 | CP014863 | GBA | CP014863 | GBA | PubMed Central | 3 |
| 5c17fd5b46d1e6422dc41547 | 28491497 | 1 | CP020570 | GBA | CP020570 | GBA | PubMed Central | 3 |
| 5c17fd5b46d1e6422dc41547 | 31533290 | 1 | CP020570 | GBA | CP020570 | GBA | PubMed Central | 3 |
| 5421eabe0d87857459fb9908 | 29931813 | 1 | CP007539 | GBA | CP007539 | GBA | PubMed Central | 3 |
| 5421eabe0d87857459fb9908 | 30042755 | 1 | CP007539 | GBA | CP007539 | GBA | PubMed Central | 3 |
| 5421eabe0d87857459fb9908 | 31346170 | 1 | 231221 | BPA | 231221 | BPA | NCBI BioProject Database | 3 |
| 5392119f0d878575e6435cb3 | 26413042 | 1 | JAAO00000000 | GBA | JAAO00000000 | GBA | PubMed Central | 3 |
| 52904631067c013e2b066737 | 23661484 | 1 | 169408 | BPA | 169408 | BPA | NCBI BioProject Database | 3 |
| 52904631067c013e2b066737 | 23661484 | 1 | AMXK00000000 | GBA | AMXK00000000 | GBA | NCBI Nucleotide Database | 3 |
| 5e573340de209cf12aa18c9d | 30461375 | 1 | CP031112 | GBA | CP031112 | GBA | PubMed Central | 3 |
| 529015cb067c013e2b05fb19 | 21799664 | 1 | 33209 | BPA | 33209 | BPA | NCBI BioProject Database | 3 |
| 529015cb067c013e2b05fb19 | 21799664 | 1 | AEAH00000000 | GBA | AEAH00000000 | GBA | NCBI Nucleotide Database | 3 |
| 529015cb067c013e2b05fb19 | 24391493 | 1 | Gi07003 | GiID | Gi07003 | GiID | PubMed Central | 3 |
| 529015cb067c013e2b05fb19 | 24710300 | 1 | GCA_000145785 | GBAsA | GL384839 | GBA | PubMed Central | 3 |
| 5d51985bc5ac66db46ea48c1 | 30718739 | 1 | MOLZ00000000 | GBA | MOLZ00000000 | GBA | PubMed Central | 3 |
| 5d51985bc5ac66db46ea48c1 | 32151246 | 1 | MOLZ00000000 | GBA | MOLZ00000000 | GBA | PubMed Central | 3 |
| 58a4bdcd7ded5e341aa820d5 | 25814599 | 1 | 270599 | BPA | 270599 | BPA | NCBI BioProject Database | 3 |
| 58a4bdcd7ded5e341aa820d5 | 25814599 | 1 | JXLT00000000 | GBA | JXLT00000000 | GBA | NCBI Nucleotide Database | 3 |
| 560c3d000d878540fd6fd621 | 29097994 | 1 | CP009241 | GBA | CP009241 | GBA | PubMed Central | 3 |
| 530d69ae49607a1be0054f6c | 24517536 | 1 | 212980 | BPA | 212980 | BPA | NCBI BioProject Database | 3 |
| 530d69ae49607a1be0054f6c | 24517536 | 1 | CP006650 | GBA | CP006650 | GBA | NCBI Nucleotide Database | 3 |
| 530d69ae49607a1be0054f6c | 24517536 | 1 | CP006651 | GBA | CP006651 | GBA | NCBI Nucleotide Database | 3 |
| 530d69ae49607a1be0054f6c | 24517536 | 1 | CP006652 | GBA | CP006652 | GBA | NCBI Nucleotide Database | 3 |
| 530d69ae49607a1be0054f6c | 24517536 | 1 | CP006653 | GBA | CP006653 | GBA | NCBI Nucleotide Database | 3 |
| 530d69ae49607a1be0054f6c | 24517536 | 1 | CP006654 | GBA | CP006654 | GBA | NCBI Nucleotide Database | 3 |
| 530d69ae49607a1be0054f6c | 24517536 | 1 | CP006655 | GBA | CP006655 | GBA | NCBI Nucleotide Database | 3 |
| 530d69ae49607a1be0054f6c | 26347732 | 1 | CP006650 | GBA | CP006650 | GBA | PubMed Central | 3 |
| 530d69ae49607a1be0054f6c | 26347732 | 1 | CP006654 | GBA | CP006654 | GBA | PubMed Central | 3 |
| 530d69ae49607a1be0054f6c | 26347732 | 1 | CP006655 | GBA | CP006655 | GBA | PubMed Central | 3 |
| 530d69ae49607a1be0054f6c | 31133656 | 1 | GCA_000444995 | GBAsA | NC_022041 | GBA | PubMed Central | 3 |
| 5654b8f00d878531d71ea27a | 26061173 | 1 | 200545 | BPA | 200545 | BPA | NCBI BioProject Database | 3 |
| 5654b8f00d878531d71ea27a | 26061173 | 1 | CP010415 | GBA | CP010415 | GBA | NCBI Nucleotide Database | 3 |
| 5654b8f00d878531d71ea27a | 26061173 | 1 | CP010415 | GBA | CP010415 | GBA | NCBI Nucleotide Database | 3 |
| 5654b8f00d878531d71ea27a | 26061173 | 1 | CP010416 | GBA | CP010416 | GBA | NCBI Nucleotide Database | 3 |
| 5654b8f00d878531d71ea27a | 26061173 | 1 | CP010416 | GBA | CP010416 | GBA | NCBI Nucleotide Database | 3 |
| 5654b8f00d878531d71ea27a | 26061173 | 1 | CP010417 | GBA | CP010417 | GBA | NCBI Nucleotide Database | 3 |
| 5654b8f00d878531d71ea27a | 26061173 | 1 | CP010417 | GBA | CP010417 | GBA | NCBI Nucleotide Database | 3 |
| 5654b8f00d878531d71ea27a | 26061173 | 1 | CP010418 | GBA | CP010418 | GBA | NCBI Nucleotide Database | 3 |
| 5654b8f00d878531d71ea27a | 26061173 | 1 | CP010418 | GBA | CP010418 | GBA | NCBI Nucleotide Database | 3 |
| 5654b8f00d878531d71ea27a | 26061173 | 1 | CP010419 | GBA | CP010419 | GBA | NCBI Nucleotide Database | 3 |

| | | | | | | | | | |
|---|---|---|---|---|---|---|---|---|---|
| 5654b8f00d878531d71ea27a | 26061173 | 1 | CP010419 | GBA | CP010419 | GBA | NCBI Nucleotide Database | | 3 |
| 5654b8f00d878531d71ea27a | 26061173 | 1 | CP010420 | GBA | CP010420 | GBA | NCBI Nucleotide Database | | 3 |
| 5654b8f00d878531d71ea27a | 26061173 | 1 | CP010420 | GBA | CP010420 | GBA | NCBI Nucleotide Database | | 3 |
| 5654b8f00d878531d71ea27a | 26061173 | 1 | CP010421 | GBA | CP010421 | GBA | NCBI Nucleotide Database | | 3 |
| 5654b8f00d878531d71ea27a | 26061173 | 1 | CP010421 | GBA | CP010421 | GBA | NCBI Nucleotide Database | | 3 |
| 5654b8f00d878531d71ea27a | 27460437 | 1 | CP010420 | GBA | CP010420 | GBA | PubMed Central | | 3 |
| 5340ec9c49607a06143562a7 | 19014550 | 1 | AARU00000000 | GBA | AARU00000000 | GBA | PubMed Central | | 3 |
| 5340ec9c49607a06143562a7 | 20846431 | 1 | AARU00000000 | GBA | AARU00000000 | GBA | PubMed Central | | 3 |
| 5340ec9c49607a06143562a7 | 23825666 | 1 | AARU00000000 | GBA | AARU00000000 | GBA | PubMed Central | | 3 |
| 5340ec9c49607a06143562a7 | 27499751 | 1 | GCA_000168615 | GBAsA | AARU02000000 | GBA | PubMed Central | | 3 |
| 5340ec9c49607a06143562a7 | 27520821 | 1 | AARU00000000 | GBA | AARU00000000 | GBA | PubMed Central | | 3 |
| 5340ec9c49607a06143562a7 | 31198443 | 1 | GCA_000168615 | GBAsA | AARU02000000 | GBA | PubMed Central | | 3 |
| 535291080d87855a8277be9c | 24324587 | 1 | EU124663 | GBA | EU124663 | GBA | PubMed Central | | 3 |
| 535291080d87855a8277be9c | 24386435 | 1 | EU124663 | GBA | EU124663 | GBA | PubMed Central | | 3 |
| 535291080d87855a8277be9c | 25128200 | 1 | EU124663 | GBA | EU124663 | GBA | PubMed Central | | 3 |
| 535291080d87855a8277be9c | 25741696 | 1 | EU124663 | GBA | EU124663 | GBA | PubMed Central | | 3 |
| 5783f7617ded5e34bd91cae8 | 27247229 | 1 | 167314 | BPA | 167314 | BPA | NCBI BioProject Database | | 3 |
| 5783f7617ded5e34bd91cae8 | 27247229 | 1 | CP013838 | GBA | CP013838 | GBA | NCBI Nucleotide Database | | 3 |
| 5783f7617ded5e34bd91cae8 | 27247229 | 1 | CP013838 | GBA | CP013838 | GBA | NCBI Nucleotide Database | | 3 |
| 5783f7617ded5e34bd91cae8 | 28470020 | 1 | CP013838 | GBA | CP013838 | GBA | PubMed Central | | 3 |
| 5783f7617ded5e34bd91cae8 | 29927994 | 1 | CP013838 | GBA | CP013838 | GBA | PubMed Central | | 3 |
| 5e572f44de209cf12aa16430 | 28408677 | 1 | CP019081 | GBA | CP019081 | GBA | PubMed Central | | 3 |
| 5e572f44de209cf12aa16430 | 29997608 | 1 | CP019081 | GBA | CP019081 | GBA | PubMed Central | | 3 |
| 52901dfe067c013e2b060c0b | 23833139 | 1 | 175625 | BPA | 175625 | BPA | NCBI BioProject Database | | 3 |
| 52901dfe067c013e2b060c0b | 23833139 | 1 | CP003906 | GBA | CP003906 | GBA | NCBI Nucleotide Database | | 3 |
| 52901dfe067c013e2b060c0b | 24358262 | 1 | GCA_000307835 | GBAsA | NC_018938 | GBA | PubMed Central | | 3 |
| 52901dfe067c013e2b060c0b | 25799515 | 1 | CP003906 | GBA | CP003906 | GBA | PubMed Central | | 3 |
| 52901dfe067c013e2b060c0b | 30648020 | 1 | GCA_000307835 | GBAsA | NC_018938 | GBA | PubMed Central | | 3 |
| 5e572da6de209cf12aa15254 | 31672739 | 1 | GCA_002023715 | GBAsA | GCA_002023715 | GBAsA | PubMed Central | | 3 |
| 56f1949e7ded5e7f7b938988 | 27182430 | 1 | 284332 | BPA | 284332 | BPA | NCBI BioProject Database | | 3 |
| 56f1949e7ded5e7f7b938988 | 30832293 | 1 | CP011564 | GBA | CP011564 | GBA | PubMed Central | | 3 |
| 56f1949e7ded5e7f7b938988 | 30832293 | 1 | CP011564 | GBA | CP011564 | GBA | PubMed Central | | 3 |
| 58ebfb3a7ded5e52c4b05539 | 29608703 | 1 | CP017149 | GBA | CP017149 | GBA | PubMed Central | | 3 |
| 58ebfb3a7ded5e52c4b05539 | 31632384 | 1 | CP017149 | GBA | CP017149 | GBA | PubMed Central | | 3 |
| 569c16050d8785737066dc2c | 25657274 | 1 | 265862 | BPA | 265862 | BPA | NCBI BioProject Database | | 3 |
| 569c16050d8785737066dc2c | 25657274 | 1 | JSXW00000000 | GBA | JSXW00000000 | GBA | NCBI Nucleotide Database | | 3 |
| 569c16050d8785737066dc2c | 30298790 | 1 | JSXW00000000 | GBA | JSXW00000000 | GBA | PubMed Central | | 3 |
| 56006e130d8785306f96b0b6 | 26213974 | 1 | JZIM00000000 | GBA | JZIM00000000 | GBA | NCBI Nucleotide Database | | 3 |
| 56af91660d878559e2870c44 | 26159520 | 1 | LBCS00000000 | GBA | LBCS00000000 | GBA | NCBI Nucleotide Database | | 3 |
| 56af91660d878559e2870c44 | 29051238 | 0 | 280939 | BPA | 280939 | BPA | NCBI BioProject Database | | 3 |
| 56af91660d878559e2870c44 | 32630808 | 1 | LBCS00000000 | GBA | LBCS00000000 | GBA | PubMed Central | | 3 |
| 58d40e5c7ded5e7a2afc0a7f | 29403463 | 1 | CP017632 | GBA | CP017632 | GBA | PubMed Central | | 3 |
| 58d40e5c7ded5e7a2afc0a7f | 30021873 | 1 | CP017633 | GBA | CP017633 | GBA | PubMed Central | | 3 |
| 58d40e5c7ded5e7a2afc0a7f | 30679636 | 1 | CP017632 | GBA | CP017632 | GBA | PubMed Central | | 3 |
| 58d40e5c7ded5e7a2afc0a7f | 31828082 | 1 | CP017632 | GBA | CP017632 | GBA | PubMed Central | | 3 |
| 58d40e5c7ded5e7a2afc0a7f | 32010114 | 1 | CP017632 | GBA | CP017632 | GBA | PubMed Central | | 3 |
| 545aa65e0d878552848901f9 | 30210484 | 1 | NC_019933 | GBA | NC_019933 | GBA | PubMed Central | | 3 |
| 56af98140d878559e2871a64 | 24571088 | 1 | CAEJ00000000 | GBA | CAEJ00000000 | GBA | PubMed Central | | 3 |
| 56af98140d878559e2871a64 | 24884520 | 1 | CAEJ00000000 | GBA | CAEJ00000000 | GBA | PubMed Central | | 3 |
| 56af98140d878559e2871a64 | 26839588 | 1 | CAEJ00000000 | GBA | CAEJ00000000 | GBA | PubMed Central | | 3 |
| 5e572fe1de209cf12aa168fa | 29220487 | 1 | GCA_900013515 | GBAsA | LT009748 | GBA | PubMed Central | | 3 |
| 5e572fe1de209cf12aa168fa | 29220487 | 1 | GCA_900013515 | GBAsA | LT009751 | GBA | PubMed Central | | 3 |
| 57a9240e7ded5e31abab5e30 | 26823594 | 1 | 302926 | BPA | 302926 | BPA | NCBI BioProject Database | | 3 |
| 57a9240e7ded5e31abab5e30 | 26823594 | 1 | CP013245 | GBA | CP013245 | GBA | NCBI Nucleotide Database | | 3 |
| 57a9240e7ded5e31abab5e30 | 26823594 | 1 | CP013245 | GBA | CP013245 | GBA | NCBI Nucleotide Database | | 3 |
| 545aa6690d87855284890333 | 29299531 | 1 | NC_022103 | GBA | NC_022103 | GBA | PubMed Central | | 3 |
| 545aa6690d87855284890333 | 32785421 | 1 | NC_022103 | GBA | NC_022103 | GBA | PubMed Central | | 3 |
| 5726e11f7ded5e4fbeb83730 | 31384648 | 1 | GCA_001037585 | GBAsA | KQ089875 | GBA | PubMed Central | | 3 |

| | | | | | | | | |
|---|---|---|---|---|---|---|---|---|
| 573f75667ded5e779637fc97 | 24923324 | 1 | CM001443 | GBA | CM001443 | GBA | PubMed Central | 3 |
| 573f75667ded5e779637fc97 | 25124552 | 1 | CM001443 | GBA | CM001443 | GBA | PubMed Central | 3 |
| 573f75667ded5e779637fc97 | 26573375 | 1 | CM001443 | GBA | CM001443 | GBA | PubMed Central | 3 |
| 5a15ce5b7ded5e41c307494c | 26934590 | 0 | NC_031929 | GBA | NC_031929 | GBA | NCBI Nucleotide Database | 3 |
| 5a15ce5b7ded5e41c307494c | 30781742 | 1 | NC_031929 | GBA | NC_031929 | GBA | PubMed Central | 3 |
| 59cc0fd47ded5e2f18693a26 | 31988474 | 1 | CP016268 | GBA | CP016268 | GBA | PubMed Central | 3 |
| 548f5cc60d8785565d46b886 | 25291776 | 1 | CP009072 | GBA | CP009072 | GBA | NCBI Nucleotide Database | 3 |
| 548f5cc60d8785565d46b886 | 25291776 | 1 | CP009072 | GBA | CP009072 | GBA | NCBI Nucleotide Database | 3 |
| 548f5cc60d8785565d46b886 | 26824353 | 1 | CP009072 | GBA | CP009072 | GBA | PubMed Central | 3 |
| 548f5cc60d8785565d46b886 | 26913025 | 1 | CP009072 | GBA | CP009072 | GBA | PubMed Central | 3 |
| 548f5cc60d8785565d46b886 | 28165067 | 1 | CP009072 | GBA | CP009072 | GBA | PubMed Central | 3 |
| 548f5cc60d8785565d46b886 | 28261184 | 1 | CP009072 | GBA | CP009072 | GBA | PubMed Central | 3 |
| 548f5cc60d8785565d46b886 | 28278280 | 1 | CP009072 | GBA | CP009072 | GBA | PubMed Central | 3 |
| 548f5cc60d8785565d46b886 | 28592550 | 1 | CP009072 | GBA | CP009072 | GBA | PubMed Central | 3 |
| 548f5cc60d8785565d46b886 | 28636609 | 1 | CP009072 | GBA | CP009072 | GBA | PubMed Central | 3 |
| 548f5cc60d8785565d46b886 | 28943892 | 1 | CP009072 | GBA | CP009072 | GBA | PubMed Central | 3 |
| 548f5cc60d8785565d46b886 | 28943892 | 1 | GCA_000743255 | GBAsA | GCA_000743255 | GBAsA | PubMed Central | 3 |
| 548f5cc60d8785565d46b886 | 29674848 | 1 | CP009072 | GBA | CP009072 | GBA | PubMed Central | 3 |
| 548f5cc60d8785565d46b886 | 29796297 | 1 | CP009072 | GBA | CP009072 | GBA | PubMed Central | 3 |
| 548f5cc60d8785565d46b886 | 30019301 | 1 | CP009072 | GBA | CP009072 | GBA | PubMed Central | 3 |
| 548f5cc60d8785565d46b886 | 30497396 | 1 | CP009072 | GBA | CP009072 | GBA | PubMed Central | 3 |
| 548f5cc60d8785565d46b886 | 30564756 | 1 | CP009072 | GBA | CP009072 | GBA | PubMed Central | 3 |
| 548f5cc60d8785565d46b886 | 30787338 | 1 | CP009072 | GBA | CP009072 | GBA | PubMed Central | 3 |
| 548f5cc60d8785565d46b886 | 30852163 | 1 | CP009072 | GBA | CP009072 | GBA | PubMed Central | 3 |
| 548f5cc60d8785565d46b886 | 31191496 | 1 | CP009072 | GBA | CP009072 | GBA | PubMed Central | 3 |
| 548f5cc60d8785565d46b886 | 31354315 | 1 | CP009072 | GBA | CP009072 | GBA | PubMed Central | 3 |
| 548f5cc60d8785565d46b886 | 31508405 | 1 | CP009072 | GBA | CP009072 | GBA | PubMed Central | 3 |
| 548f5cc60d8785565d46b886 | 31509535 | 0 | CP009072 | GBA | CP009072 | GBA | PubMed Central | 3 |
| 548f5cc60d8785565d46b886 | 31896625 | 1 | CP009072 | GBA | CP009072 | GBA | PubMed Central | 3 |
| 548f5cc60d8785565d46b886 | 32102944 | 1 | CP009072 | GBA | CP009072 | GBA | PubMed Central | 3 |
| 548f5cc60d8785565d46b886 | 32205351 | 1 | CP009072 | GBA | CP009072 | GBA | PubMed Central | 3 |
| 548f5cc60d8785565d46b886 | 32376701 | 1 | CP009072 | GBA | CP009072 | GBA | PubMed Central | 3 |
| 548f5cc60d8785565d46b886 | 32376847 | 0 | CP009072 | GBA | CP009072 | GBA | PubMed Central | 3 |
| 548f5cc60d8785565d46b886 | 32591016 | 1 | CP009072 | GBA | CP009072 | GBA | PubMed Central | 3 |
| 548f5cc60d8785565d46b886 | 32971800 | 1 | CP009072 | GBA | CP009072 | GBA | PubMed Central | 3 |
| 548f5cc60d8785565d46b886 | 33176691 | 1 | CP009072 | GBA | CP009072 | GBA | PubMed Central | 3 |
| 548f5cc60d8785565d46b886 | 33327465 | 1 | CP009072 | GBA | CP009074 | GBA | PubMed Central | 3 |
| 5ab98c1764d0b30c13702b4f | 27811087 | 1 | MCNV00000000 | GBA | MCNV00000000 | GBA | PubMed Central | 3 |
| 545aa6340d8785528488ff4d | 22413005 | 1 | NC_019925 | GBA | NC_019925 | GBA | NCBI Nucleotide Database | 3 |
| 545aa6340d8785528488ff4d | 25212610 | 1 | NC_019925 | GBA | NC_019925 | GBA | PubMed Central | 3 |
| 545aa6340d8785528488ff4d | 29555775 | 1 | NC_019925 | GBA | NC_019925 | GBA | PubMed Central | 3 |
| 5ae77fc964d0b337477333d8 | 31582435 | 1 | CP023664 | GBA | CP023664 | GBA | PubMed Central | 3 |
| 560c3c720d878540fd6fce46 | 26958345 | 1 | CP009285 | GBA | CP009285 | GBA | PubMed Central | 3 |
| 560c3c720d878540fd6fce46 | 26958346 | 1 | CP009285 | GBA | CP009285 | GBA | PubMed Central | 3 |
| 560c3c720d878540fd6fce46 | 28912952 | 1 | CP009285 | GBA | CP009285 | GBA | PubMed Central | 3 |
| 560c3c720d878540fd6fce46 | 29692909 | 1 | CP009285 | GBA | CP009285 | GBA | PubMed Central | 3 |
| 5784012d7ded5e34bd91d7d6 | 29217787 | 1 | LKIR00000000 | GBA | LKIR00000000 | GBA | PubMed Central | 3 |
| 53adb5820d878514c2d128dd | 24855300 | 1 | 227037 | BPA | 227037 | BPA | NCBI BioProject Database | 3 |
| 53adb5820d878514c2d128dd | 24855300 | 1 | CP007566 | GBA | CP007566 | GBA | NCBI Nucleotide Database | 3 |
| 53adb5820d878514c2d128dd | 24855300 | 1 | CP007566 | GBA | CP007566 | GBA | NCBI Nucleotide Database | 3 |
| 53adb5820d878514c2d128dd | 25768732 | 1 | CP007566 | GBA | CP007566 | GBA | PubMed Central | 3 |
| 53adb5820d878514c2d128dd | 25768732 | 1 | CP007566 | GBA | CP007566 | GBA | PubMed Central | 3 |
| 53adb5820d878514c2d128dd | 25768732 | 1 | CP007566 | GBA | CP007566 | GBA | PubMed Central | 3 |
| 53adb5820d878514c2d128dd | 27803692 | 1 | CP007566 | GBA | CP007566 | GBA | PubMed Central | 3 |
| 53adb5820d878514c2d128dd | 31572337 | 1 | CP007566 | GBA | CP007566 | GBA | PubMed Central | 3 |
| 53adb5820d878514c2d128dd | 32968153 | 1 | CP007566 | GBA | CP007566 | GBA | PubMed Central | 3 |
| 5340eccb49607a06143562fb | 25197489 | 1 | ABXH00000000 | GBA | ABXH00000000 | GBA | PubMed Central | 3 |
| 5340eccb49607a06143562fb | 29204287 | 1 | ABXH00000000 | GBA | ABXH00000000 | GBA | PubMed Central | 3 |

| | | | | | | | | |
|---|---|---|---|---|---|---|---|---|
| 5340eccb49607a06143562fb | 29900684 | 1 | ABXH00000000 | GBA | ABXH00000000 | GBA | PubMed Central | 3 |
| 5340eccb49607a06143562fb | 30186281 | 1 | ABXH00000000 | GBA | SAMN00008813 | BSA | PubMed Central | 3 |
| 5340e14549607a061435562f | 24003073 | 1 | ALWQ00000000 | GBA | ALWQ00000000 | GBA | PubMed Central | 3 |
| 538022800d87852a04c5c2cd | 24459282 | 1 | 228956 | BPA | 228956 | BPA | NCBI BioProject Database | 3 |
| 538022800d87852a04c5c2cd | 24459282 | 1 | AYTE00000000 | GBA | AYTE00000000 | GBA | NCBI Nucleotide Database | 3 |
| 538022800d87852a04c5c2cd | 29025939 | 1 | AYTE00000000 | GBA | AYTE00000000 | GBA | PubMed Central | 3 |
| 538022800d87852a04c5c2cd | 29025939 | 1 | Ga0041805 | GaID | Ga0041805 | GaID | PubMed Central | 3 |
| 545aa64a0d8785528489017b | 30081963 | 1 | NC_020899 | GBA | NC_020899 | GBA | PubMed Central | 3 |
| 58d40ca17ded5e7a2afc07ac | 27325916 | 1 | 291949 | BPA | 291949 | BPA | NCBI BioProject Database | 3 |
| 58d40ca17ded5e7a2afc07ac | 27325916 | 1 | CP012739 | GBA | CP012739 | GBA | NCBI Nucleotide Database | 3 |
| 58d40ca17ded5e7a2afc07ac | 27325916 | 1 | CP012739 | GBA | CP012739 | GBA | NCBI Nucleotide Database | 3 |
| 58d40ca17ded5e7a2afc07ac | 27325916 | 1 | CP012740 | GBA | CP012740 | GBA | NCBI Nucleotide Database | 3 |
| 58d40ca17ded5e7a2afc07ac | 27325916 | 1 | CP012740 | GBA | CP012740 | GBA | NCBI Nucleotide Database | 3 |
| 58d40ca17ded5e7a2afc07ac | 27325916 | 1 | CP012741 | GBA | CP012741 | GBA | NCBI Nucleotide Database | 3 |
| 58d40ca17ded5e7a2afc07ac | 27325916 | 1 | CP012741 | GBA | CP012741 | GBA | NCBI Nucleotide Database | 3 |
| 58d40ca17ded5e7a2afc07ac | 31711432 | 1 | CP012739 | GBA | CP012739 | GBA | PubMed Central | 3 |
| 529f9d55067c0121bf0d9f8a | 24092784 | 1 | AEQS00000000 | GBA | AEQS00000000 | GBA | PubMed Central | 3 |
| 529f9d55067c0121bf0d9f8a | 26644037 | 1 | AEQS00000000 | GBA | AEQS00000000 | GBA | PubMed Central | 3 |
| 529f9d55067c0121bf0d9f8a | 30405564 | 1 | AEQS00000000 | GBA | AEQS00000000 | GBA | PubMed Central | 3 |
| 5340e58949607a0614355d91 | 24812228 | 1 | 226227 | BPA | 226227 | BPA | NCBI BioProject Database | 3 |
| 5340e58949607a0614355d91 | 24812228 | 1 | CP006900 | GBA | CP006900 | GBA | NCBI Nucleotide Database | 3 |
| 5340e58949607a0614355d91 | 28070515 | 1 | CP006900 | GBA | CP006900 | GBA | PubMed Central | 3 |
| 5340e58949607a0614355d91 | 30761094 | 1 | CP006900 | GBA | CP006900 | GBA | PubMed Central | 3 |
| 5340e58949607a0614355d91 | 32271799 | 1 | CP006900 | GBA | CP006900 | GBA | PubMed Central | 3 |
| 5e3d6af448feb56cc51ca959 | 26441845 | 1 | JWHU00000000 | GBA | JWHU00000000 | GBA | PubMed Central | 3 |
| 5e3d6af448feb56cc51ca959 | 26694728 | 1 | JWHU00000000 | GBA | JWHU00000000 | GBA | PubMed Central | 3 |
| 56362d620d87852aea5e5dea | 26358601 | 1 | LFXA00000000 | GBA | LFXA00000000 | GBA | NCBI Nucleotide Database | 3 |
| 57721a077ded5e34d82975ff | 27766204 | 1 | LVVO00000000 | GBA | LVVO00000000 | GBA | PubMed Central | 3 |
| 57721a077ded5e34d82975ff | 28163826 | 1 | Gp0147190 | GpID | Gp0147190 | GpID | PubMed Central | 3 |
| 57721a077ded5e34d82975ff | 28163826 | 1 | LVVO00000000 | GBA | LVVO00000000 | GBA | PubMed Central | 3 |
| 58ebf2597ded5e52c4b048f8 | 27774985 | 1 | MGZV00000000 | GBA | MGZV00000000 | GBA | NCBI Nucleotide Database | 3 |
| 58ebf2597ded5e52c4b048f8 | 30687241 | 1 | MGZV00000000 | GBA | MGZV00000000 | GBA | PubMed Central | 3 |
| 59a1ac657ded5e41edd85fd2 | 27313303 | 1 | 299439 | BPA | 299439 | BPA | NCBI BioProject Database | 3 |
| 59a1ac657ded5e41edd85fd2 | 27313303 | 1 | LMCA00000000 | GBA | LMCA00000000 | GBA | NCBI Nucleotide Database | 3 |
| 55791d4c0d878529e7cb15af | 26380634 | 1 | JRHH00000000 | GBA | JRHH00000000 | GBA | PubMed Central | 3 |
| 55791d4c0d878529e7cb15af | 27313837 | 1 | JRHH00000000 | GBA | JRHH00000000 | GBA | PubMed Central | 3 |
| 58ebfa447ded5e52c4b053e4 | 27284141 | 1 | 311246 | BPA | 311246 | BPA | NCBI BioProject Database | 3 |
| 58ebfa447ded5e52c4b053e4 | 27284141 | 1 | CP014349 | GBA | CP014349 | GBA | NCBI Nucleotide Database | 3 |
| 58ebfa447ded5e52c4b053e4 | 27284141 | 1 | CP014349 | GBA | CP014349 | GBA | NCBI Nucleotide Database | 3 |
| 58ebfa447ded5e52c4b053e4 | 27284141 | 1 | CP014350 | GBA | CP014350 | GBA | NCBI Nucleotide Database | 3 |
| 58ebfa447ded5e52c4b053e4 | 27284141 | 1 | CP014350 | GBA | CP014350 | GBA | NCBI Nucleotide Database | 3 |
| 58ebfa447ded5e52c4b053e4 | 27284141 | 1 | CP014351 | GBA | CP014351 | GBA | NCBI Nucleotide Database | 3 |
| 58ebfa447ded5e52c4b053e4 | 27284141 | 1 | CP014351 | GBA | CP014351 | GBA | NCBI Nucleotide Database | 3 |
| 58ebfa447ded5e52c4b053e4 | 27284141 | 1 | CP014792 | GBA | CP014792 | GBA | NCBI Nucleotide Database | 3 |
| 58ebfa447ded5e52c4b053e4 | 27284141 | 1 | CP014792 | GBA | CP014792 | GBA | NCBI Nucleotide Database | 3 |
| 58ebfa447ded5e52c4b053e4 | 27284141 | 1 | CP014871 | GBA | CP014871 | GBA | NCBI Nucleotide Database | 3 |
| 58ebfa447ded5e52c4b053e4 | 27284141 | 1 | CP014871 | GBA | CP014871 | GBA | NCBI Nucleotide Database | 3 |
| 58ebfa447ded5e52c4b053e4 | 27284141 | 1 | CP015331 | GBA | CP015331 | GBA | NCBI Nucleotide Database | 3 |
| 58ebfa447ded5e52c4b053e4 | 27284141 | 1 | CP015331 | GBA | CP015331 | GBA | NCBI Nucleotide Database | 3 |
| 58ebfa447ded5e52c4b053e4 | 27284141 | 1 | CP015332 | GBA | CP015332 | GBA | NCBI Nucleotide Database | 3 |
| 58ebfa447ded5e52c4b053e4 | 27284141 | 1 | CP015332 | GBA | CP015332 | GBA | NCBI Nucleotide Database | 3 |
| 58ebfa447ded5e52c4b053e4 | 27284141 | 1 | CP015333 | GBA | CP015333 | GBA | NCBI Nucleotide Database | 3 |
| 58ebfa447ded5e52c4b053e4 | 27284141 | 1 | CP015333 | GBA | CP015333 | GBA | NCBI Nucleotide Database | 3 |
| 58ebfa447ded5e52c4b053e4 | 27284141 | 1 | CP015334 | GBA | CP015334 | GBA | NCBI Nucleotide Database | 3 |
| 58ebfa447ded5e52c4b053e4 | 27284141 | 1 | CP015334 | GBA | CP015334 | GBA | NCBI Nucleotide Database | 3 |
| 58ebfa447ded5e52c4b053e4 | 27284141 | 1 | CP015335 | GBA | CP015335 | GBA | NCBI Nucleotide Database | 3 |
| 58ebfa447ded5e52c4b053e4 | 27284141 | 1 | CP015335 | GBA | CP015335 | GBA | NCBI Nucleotide Database | 3 |
| 58ebfa447ded5e52c4b053e4 | 27284141 | 1 | CP015336 | GBA | CP015336 | GBA | NCBI Nucleotide Database | 3 |

| | | | | | | | | |
|---|---|---|---|---|---|---|---|---|
| 58ebfa447ded5e52c4b053e4 | 27284141 | 1 | CP015336 | GBA | CP015336 | GBA | NCBI Nucleotide Database | 3 |
| 58ebfa447ded5e52c4b053e4 | 28302772 | 1 | CP014349 | GBA | SAMN04481062 | BSA | PubMed Central | 3 |
| 58ebfa447ded5e52c4b053e4 | 28302772 | 1 | CP014349 | GBA | SAMN04481062 | BSA | PubMed Central | 3 |
| 58ebfa447ded5e52c4b053e4 | 28302772 | 1 | CP014350 | GBA | SAMN04481062 | BSA | PubMed Central | 3 |
| 58ebfa447ded5e52c4b053e4 | 28302772 | 1 | CP014350 | GBA | SAMN04481062 | BSA | PubMed Central | 3 |
| 58ebfa447ded5e52c4b053e4 | 28302772 | 1 | CP014351 | GBA | SAMN04481062 | BSA | PubMed Central | 3 |
| 58ebfa447ded5e52c4b053e4 | 28302772 | 1 | CP014351 | GBA | SAMN04481062 | BSA | PubMed Central | 3 |
| 58ebfa447ded5e52c4b053e4 | 28302772 | 1 | CP014792 | GBA | SAMN04481062 | BSA | PubMed Central | 3 |
| 58ebfa447ded5e52c4b053e4 | 28302772 | 1 | CP014792 | GBA | SAMN04481062 | BSA | PubMed Central | 3 |
| 58ebfa447ded5e52c4b053e4 | 28302772 | 1 | CP014871 | GBA | SAMN04481062 | BSA | PubMed Central | 3 |
| 58ebfa447ded5e52c4b053e4 | 28302772 | 1 | CP014871 | GBA | SAMN04481062 | BSA | PubMed Central | 3 |
| 58ebfa447ded5e52c4b053e4 | 28302772 | 1 | CP015331 | GBA | SAMN04481062 | BSA | PubMed Central | 3 |
| 58ebfa447ded5e52c4b053e4 | 28302772 | 1 | CP015331 | GBA | SAMN04481062 | BSA | PubMed Central | 3 |
| 58ebfa447ded5e52c4b053e4 | 28302772 | 1 | CP015332 | GBA | SAMN04481062 | BSA | PubMed Central | 3 |
| 58ebfa447ded5e52c4b053e4 | 28302772 | 1 | CP015332 | GBA | SAMN04481062 | BSA | PubMed Central | 3 |
| 58ebfa447ded5e52c4b053e4 | 28302772 | 1 | CP015333 | GBA | SAMN04481062 | BSA | PubMed Central | 3 |
| 58ebfa447ded5e52c4b053e4 | 28302772 | 1 | CP015333 | GBA | SAMN04481062 | BSA | PubMed Central | 3 |
| 58ebfa447ded5e52c4b053e4 | 28302772 | 1 | CP015334 | GBA | SAMN04481062 | BSA | PubMed Central | 3 |
| 58ebfa447ded5e52c4b053e4 | 28302772 | 1 | CP015334 | GBA | SAMN04481062 | BSA | PubMed Central | 3 |
| 58ebfa447ded5e52c4b053e4 | 28302772 | 1 | CP015335 | GBA | SAMN04481062 | BSA | PubMed Central | 3 |
| 58ebfa447ded5e52c4b053e4 | 28302772 | 1 | CP015335 | GBA | SAMN04481062 | BSA | PubMed Central | 3 |
| 58ebfa447ded5e52c4b053e4 | 28302772 | 1 | CP015336 | GBA | SAMN04481062 | BSA | PubMed Central | 3 |
| 58ebfa447ded5e52c4b053e4 | 28302772 | 1 | CP015336 | GBA | SAMN04481062 | BSA | PubMed Central | 3 |
| 58ebfa447ded5e52c4b053e4 | 28656008 | 1 | CP014349 | GBA | SAMN04481062 | BSA | PubMed Central | 3 |
| 58ebfa447ded5e52c4b053e4 | 28656008 | 1 | CP014349 | GBA | SAMN04481062 | BSA | PubMed Central | 3 |
| 58ebfa447ded5e52c4b053e4 | 28656008 | 1 | CP014350 | GBA | SAMN04481062 | BSA | PubMed Central | 3 |
| 58ebfa447ded5e52c4b053e4 | 28656008 | 1 | CP014350 | GBA | SAMN04481062 | BSA | PubMed Central | 3 |
| 58ebfa447ded5e52c4b053e4 | 28656008 | 1 | CP014351 | GBA | SAMN04481062 | BSA | PubMed Central | 3 |
| 58ebfa447ded5e52c4b053e4 | 28656008 | 1 | CP014351 | GBA | SAMN04481062 | BSA | PubMed Central | 3 |
| 58ebfa447ded5e52c4b053e4 | 28656008 | 1 | CP014792 | GBA | SAMN04481062 | BSA | PubMed Central | 3 |
| 58ebfa447ded5e52c4b053e4 | 28656008 | 1 | CP014792 | GBA | SAMN04481062 | BSA | PubMed Central | 3 |
| 58ebfa447ded5e52c4b053e4 | 28656008 | 1 | CP014871 | GBA | SAMN04481062 | BSA | PubMed Central | 3 |
| 58ebfa447ded5e52c4b053e4 | 28656008 | 1 | CP014871 | GBA | SAMN04481062 | BSA | PubMed Central | 3 |
| 58ebfa447ded5e52c4b053e4 | 28656008 | 1 | CP015331 | GBA | SAMN04481062 | BSA | PubMed Central | 3 |
| 58ebfa447ded5e52c4b053e4 | 28656008 | 1 | CP015331 | GBA | SAMN04481062 | BSA | PubMed Central | 3 |
| 58ebfa447ded5e52c4b053e4 | 28656008 | 1 | CP015332 | GBA | SAMN04481062 | BSA | PubMed Central | 3 |
| 58ebfa447ded5e52c4b053e4 | 28656008 | 1 | CP015332 | GBA | SAMN04481062 | BSA | PubMed Central | 3 |
| 58ebfa447ded5e52c4b053e4 | 28656008 | 1 | CP015333 | GBA | SAMN04481062 | BSA | PubMed Central | 3 |
| 58ebfa447ded5e52c4b053e4 | 28656008 | 1 | CP015333 | GBA | SAMN04481062 | BSA | PubMed Central | 3 |
| 58ebfa447ded5e52c4b053e4 | 28656008 | 1 | CP015334 | GBA | SAMN04481062 | BSA | PubMed Central | 3 |
| 58ebfa447ded5e52c4b053e4 | 28656008 | 1 | CP015334 | GBA | SAMN04481062 | BSA | PubMed Central | 3 |
| 58ebfa447ded5e52c4b053e4 | 28656008 | 1 | CP015335 | GBA | SAMN04481062 | BSA | PubMed Central | 3 |
| 58ebfa447ded5e52c4b053e4 | 28656008 | 1 | CP015335 | GBA | SAMN04481062 | BSA | PubMed Central | 3 |
| 58ebfa447ded5e52c4b053e4 | 28656008 | 1 | CP015336 | GBA | SAMN04481062 | BSA | PubMed Central | 3 |
| 58ebfa447ded5e52c4b053e4 | 28656008 | 1 | CP015336 | GBA | SAMN04481062 | BSA | PubMed Central | 3 |
| 58ebfa447ded5e52c4b053e4 | 29056727 | 1 | CP014349 | GBA | SAMN04481062 | BSA | PubMed Central | 3 |
| 58ebfa447ded5e52c4b053e4 | 29056727 | 1 | CP014349 | GBA | SAMN04481062 | BSA | PubMed Central | 3 |
| 58ebfa447ded5e52c4b053e4 | 29056727 | 1 | CP014350 | GBA | SAMN04481062 | BSA | PubMed Central | 3 |
| 58ebfa447ded5e52c4b053e4 | 29056727 | 1 | CP014350 | GBA | SAMN04481062 | BSA | PubMed Central | 3 |
| 58ebfa447ded5e52c4b053e4 | 29056727 | 1 | CP014351 | GBA | SAMN04481062 | BSA | PubMed Central | 3 |
| 58ebfa447ded5e52c4b053e4 | 29056727 | 1 | CP014351 | GBA | SAMN04481062 | BSA | PubMed Central | 3 |
| 58ebfa447ded5e52c4b053e4 | 29056727 | 1 | CP014792 | GBA | SAMN04481062 | BSA | PubMed Central | 3 |
| 58ebfa447ded5e52c4b053e4 | 29056727 | 1 | CP014792 | GBA | SAMN04481062 | BSA | PubMed Central | 3 |
| 58ebfa447ded5e52c4b053e4 | 29056727 | 1 | CP014871 | GBA | SAMN04481062 | BSA | PubMed Central | 3 |
| 58ebfa447ded5e52c4b053e4 | 29056727 | 1 | CP014871 | GBA | SAMN04481062 | BSA | PubMed Central | 3 |
| 58ebfa447ded5e52c4b053e4 | 29056727 | 1 | CP015331 | GBA | SAMN04481062 | BSA | PubMed Central | 3 |
| 58ebfa447ded5e52c4b053e4 | 29056727 | 1 | CP015331 | GBA | SAMN04481062 | BSA | PubMed Central | 3 |
| 58ebfa447ded5e52c4b053e4 | 29056727 | 1 | CP015332 | GBA | SAMN04481062 | BSA | PubMed Central | 3 |

| | | | | | | | | |
|---|---|---|---|---|---|---|---|---|
| 58ebfa447ded5e52c4b053e4 | 29056727 | 1 | CP015332 | GBA | SAMN04481062 | BSA | PubMed Central | 3 |
| 58ebfa447ded5e52c4b053e4 | 29056727 | 1 | CP015333 | GBA | SAMN04481062 | BSA | PubMed Central | 3 |
| 58ebfa447ded5e52c4b053e4 | 29056727 | 1 | CP015333 | GBA | SAMN04481062 | BSA | PubMed Central | 3 |
| 58ebfa447ded5e52c4b053e4 | 29056727 | 1 | CP015334 | GBA | SAMN04481062 | BSA | PubMed Central | 3 |
| 58ebfa447ded5e52c4b053e4 | 29056727 | 1 | CP015334 | GBA | SAMN04481062 | BSA | PubMed Central | 3 |
| 58ebfa447ded5e52c4b053e4 | 29056727 | 1 | CP015335 | GBA | SAMN04481062 | BSA | PubMed Central | 3 |
| 58ebfa447ded5e52c4b053e4 | 29056727 | 1 | CP015335 | GBA | SAMN04481062 | BSA | PubMed Central | 3 |
| 58ebfa447ded5e52c4b053e4 | 29056727 | 1 | CP015336 | GBA | SAMN04481062 | BSA | PubMed Central | 3 |
| 58ebfa447ded5e52c4b053e4 | 29056727 | 1 | CP015336 | GBA | SAMN04481062 | BSA | PubMed Central | 3 |
| 58ebfa447ded5e52c4b053e4 | 30586413 | 1 | CP014349 | GBA | CP014349 | GBA | PubMed Central | 3 |
| 58ebfa447ded5e52c4b053e4 | 30586413 | 1 | CP014349 | GBA | CP014349 | GBA | PubMed Central | 3 |
| 57a92aca7ded5e31abab66a7 | 32150591 | 1 | CP014239 | GBA | CP014239 | GBA | PubMed Central | 3 |
| 5af6454164d0b3374774bae5 | 29242221 | 1 | NKDV00000000 | GBA | NKDV00000000 | GBA | NCBI Nucleotide Database | 3 |
| 5af6454164d0b3374774bae5 | 29724828 | 1 | 230969 | BPA | 230969 | BPA | NCBI BioProject Database | 3 |
| 5af6454164d0b3374774bae5 | 30810518 | 1 | NKDV00000000 | GBA | SAMN05591547 | BSA | PubMed Central | 3 |
